# Evolution of microRNAs in Amoebozoa and implications for the origin of multicellularity

**DOI:** 10.1101/2023.10.19.562841

**Authors:** Bart Edelbroek, Jonas Kjellin, Inna Biryukova, Zhen Liao, Torgny Lundberg, Angelika A. Noegel, Ludwig Eichinger, Marc R. Friedländer, Fredrik Söderbom

**Affiliations:** Department of Cell and Molecular Biology, Biomedical Centre, Uppsala University, 75124 Uppsala, Sweden; Science for Life Laboratory, Department of Molecular Biosciences, The Wenner-Gren Institute, Stockholm University, Stockholm, Sweden; Centre for Biochemistry, Medical Faculty, University of Cologne, 50931 Cologne, Germany; Génétique Moléculaire, Génomique, Microbiologie (GMGM), University of Strasbourg, 67000 Strasbourg, France

## Abstract

MicroRNAs (miRNAs) are important and ubiquitous regulators of gene expression in both plants and animals. They are thought to have evolved convergently in these lineages and hypothesized to have played a role in the evolution of multicellularity. In line with this hypothesis, miRNAs have so far only been described in few unicellular eukaryotes. Here, we investigate the presence and evolution of miRNAs in Amoebozoa, focusing on species belonging to *Acanthamoeba*, *Physarum*, and dictyostelid taxonomic groups, representing a range of unicellular and multicellular lifestyles. miRNAs that adhere to both the stringent plant and animal miRNA criteria were identified in all examined amoebae, greatly expanding the total number of protists harbouring miRNAs. We found conserved miRNAs between closely related species, but the majority of species feature only unique miRNAs. Our results show that miRNAs are rapidly lost and gained in Amoebozoa, and that miRNAs were not required for transition from uni- to multicellular life.

## Introduction

microRNAs (miRNAs) are small, ∼21 nucleotide (nt) non-coding RNAs, which control gene expression to create intricate regulatory networks^1^. The vast majority of miRNAs has been identified in animals and plants^2–7^. In both these lineages, miRNA biogenesis involves transcription of larger primary, hairpin structured precursors (pri-miRNAs) that are processed to pre-miRNAs before being matured into miRNAs^8^. This process is performed by a machinery derived from the ancestral RNA-interference (RNAi) machinery that dates back to the last eukaryotic ancestor and protects cells from viruses and mobilization of transposons^9–13^. Once processed to mature miRNAs, these small RNAs specify genes to be regulated based on binding to their target mRNA(s) via sequence complementarity, thereby guiding the RNA induced silencing complex (RISC) to perform translational silencing and/or induce target RNA degradation^14^. Even though miRNA maturation and formation of the RISC involve a number of associated proteins – to some extent differing between animal and plants – two proteins, Dicer and Argonaute, are central to a functional RNAi machinery regardless of organism. While Dicer binds to miRNA hairpin structures and cleaves out double stranded miRNAs (further matured to single stranded miRNAs in the RISC), Argonautes are the effector proteins, guided to the target RNA by bound miRNA^14^. Although the animal and plant miRNA machineries share many characteristics, miRNA regulation is generally thought to have evolved convergently. In particular, no miRNA sequences have been found to be conserved between plants and animals^2^. The convergent evolution of miRNAs in plants and animals is, however, still up for debate^15,16^.

It is perhaps no coincidence that the majority of miRNAs has been discovered in either plants or animals, where many organisms show a high degree of organismal complexity^14,17^. Both plants and animals independently evolved clonal multicellularity and regulatory genetic networks necessary for multicellular life^18^. It is possible that miRNA-mediated gene regulation was already present during the transition from uni- to multicellularity, especially since miRNAs have been found in unicellular organisms closely related to both lineages^19–22^. Hence, miRNAs may have contributed in the evolution of multicellularity, but whether or not this is the case remains unclear^19,23,24^.

To better address fundamental questions regarding the evolution of miRNAs and their contribution to multicellularity, stringent analysis of the small RNA populations in understudied eukaryotic lineages is needed^15^. One of these is the Amoebozoa taxonomic group, which belongs to the Amorphea supergroup that also includes animals and fungi^25^. The Amoebozoans exhibit diverse lifestyles and a varied level of developmental complexity. Some are strictly unicellular, such as the *Acanthamoebae*^26^. Others, such as *Physarum polycephalum*, are unicellular but can switch from a free-living amoeboid to a plasmodial lifestyle in which it grows without cytokinesis. This results in a large intricate multinucleate network that permits sensing and transport of nutrients^27^. Upon starvation, the plasmodium can differentiate to form sporangia^27^. Another group of Amoebozoans is the Dictyostelia, estimated to date back 600 million years^28^. Members of Dictyostelia switch from a unicellular life to an aggregative multicellular lifestyle upon starvation. Here, up to 100,000 cells stream together to form distinct multicellular structures such as migrating slugs, where cells can collaborate to move towards light^29^. Finally, cells culminate to form a fruiting body^29^. Even within the dictyostelids, the level of developmental complexity varies. Some, like *Acytostelium subglobosum*, form a fruiting body with one single cell type, while others, such as *Dictyostelium discoideum*, employ up to four major cell types.

Thus far, *D. discoideum* is the only Amoebozoan in which miRNAs have been convincingly identified^30–33^. The *D. discoideum* miRNAs are mostly expressed from single intergenic regions and are upregulated upon multicellular development. They rely on the Dicer-like protein, DrnB, and a double-stranded RNA binding domain-containing protein RbdB for their processing^32,33^. *D. discoideum* also encodes five Argonautes. Which one of these binds miRNAs is still unknown.

Here, we systematically investigated the small RNA repertoire of nine Amoebozoa species to approach the evolutionary history of miRNAs. We also address the hypothesized role of miRNAs as a driver for the evolution of multicellularity. The amoebae we used were six dictyostelids, the closely related *P. polycephalum*, and two species belonging to *Acanthamoebae*. By using stringent miRNA identification criteria, high-confidence miRNAs were identified in all the multicellular dictyostelids, but also in *P. polycephalum* and the strictly unicellular *Acanthamoebae* species. One of the identified miRNAs is conserved within the lineage of *Acanthamoebae* and another is conserved between group 4 dictyostelids, one of the four monophyletic groups of Dictyostelia. No other conserved miRNAs were detected, and nearly all miRNAs appear to be recent innovations. The importance of miRNAs for unicellular life is indicated by the expression and conservation of miRNAs in Amoebozoans that lack a multicellular lifestyle, and is corroborated by the increased generation time of cells depleted of miRNAs in *D. discoideum*. Further, phylogenetic analysis of RNAi components shows a rapidly evolving and flexible system with their numbers varying between species. Together, our results indicate that miRNAs have evolved recently and rapidly throughout Amoebozoa, and that the miRNA regulatory networks are not exclusive to Amoebozoa that exhibit aggregative multicellularity.

## Results

### microRNAs are present in six out of six dictyostelid social amoebae studied

Our previous discovery of miRNAs in *D. discoideum* prompted us to investigate whether these small RNAs are also present in other social amoebae^30,31^. If so, miRNAs may be important not only for cell differentiation in organisms exhibiting clonal multicellularity (e.g. animals and plants) but also for those that display aggregative multicellularity. To address this, we sequenced small RNAs from six species belonging to the four different phylogenetic groups of Dictyostelia^28,34^ (Fig. 1a). To facilitate capturing of temporally expressed miRNAs, RNA was isolated and sequenced from three distinct stages – vegetative growth and two stages of multicellular development (finger/slug stage and culminating fruiting bodies [Extended Data Fig. 1]). The sequenced small RNAs were mapped against the genomes of the respective amoebae and the *Escherichia coli* bacteria used as food source. Reads derived from bacteria and rRNA were removed from further analyses (Extended Data Fig. 2a, Methods). In agreement with the previous results from *D. discoideum*^30^, the small RNAs showed a distinct peak at 21 nt for all social amoebae (Extended Data Fig. 2b).

**Fig. 1.**
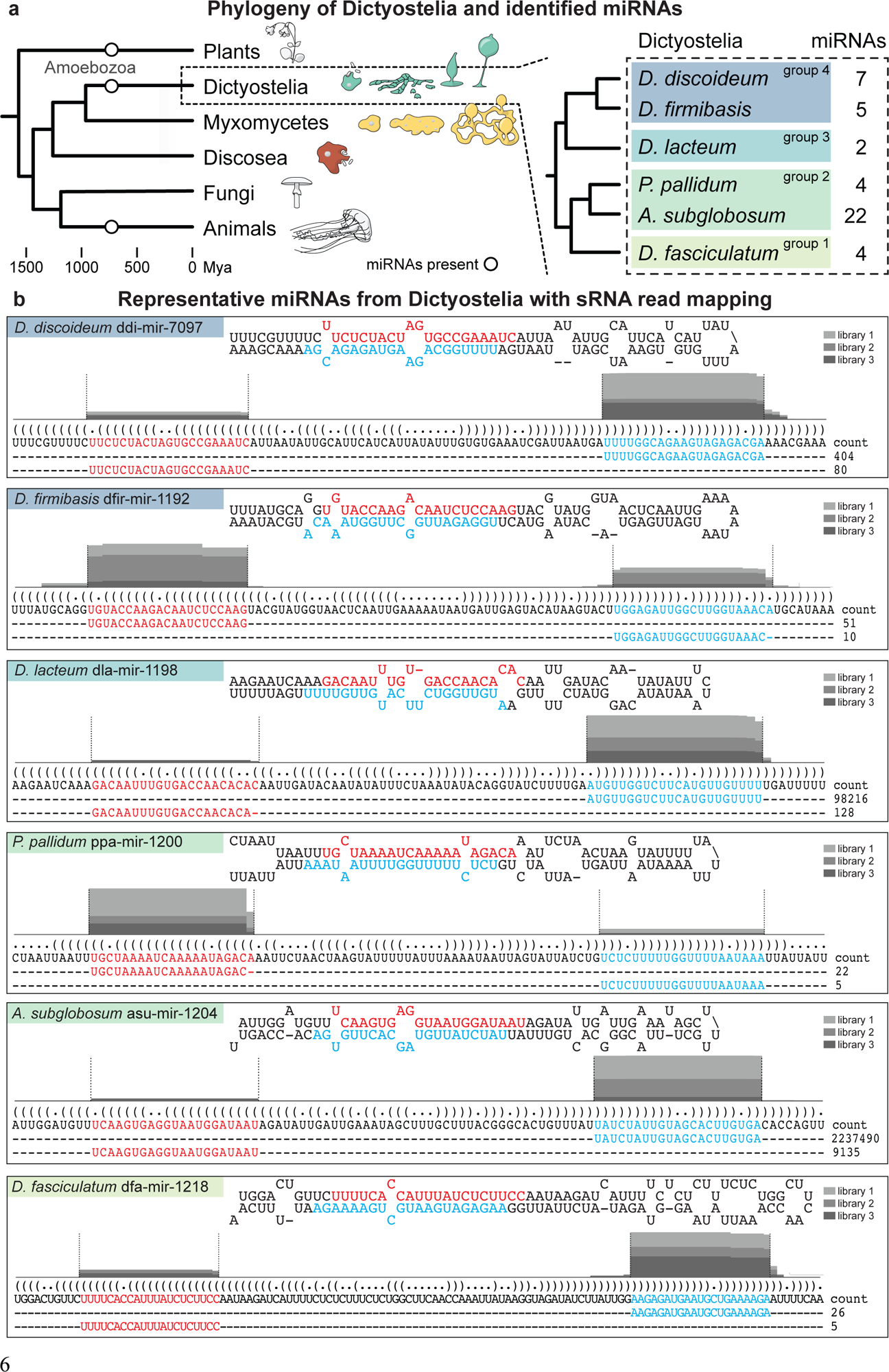
miRNAs identified in social amoebae. **a** Simplified phylogeny illustrating the position of Amoebozoa between Animals and Plants. The branches of Amoebozoa included in this study are shown with characteristic phenotypes: true multicellular aggregation for Dictyostelia, the multinucleate plasmodial stage in Myxomycetes, and single-cellular Discosea. The group of Dictyostelia is further enlarged to display the phylogeny of the four major groups of Dictyostelia social amoebae and the specific members included in this study. Number of miRNAs, identified in this study, for the different dictyostelids is indicated. Mya, million years ago **b** Representative examples of miRNAs from each of the studied Dictyostelia. pri-miRNAs are visualized with their predicted secondary structures, with the miRNA-5p and miRNA-3p marked in red and cyan, respectively. The DNA sequences representing the pri-miRNAs are displayed in bracket notation with their most abundant reads mapping to the miRNA-5p and miRNA-3p arms, together with their read-counts. Histogram above the pri-miRNA DNA sequence shows proportion of sRNA reads mapping to each nucleotide, divided into three shades of grey for each of the three biological replicates. The histograms for dla-mir-1198-5p and asu-mir-1204-5p are not to scale, but shown for clarity. For detailed information, see Supplementary Data 1.

Our earlier results in *D. discoideum* showed that the majority of small RNAs were small interfering (si)RNAs^30–32,35^. Furthermore, the biogenesis of *D. discoideum* miRNAs showed similarities to both plant and animal miRNA maturation, congruent with the phylogenetic position of this species^32,36^. Based on these observations, the sequenced small RNAs were subjected to stringent analyses, applying sets of criteria used to define miRNAs in plants, animals, or both^2,37–39^. To ensure that we would report only *bona fide* miRNAs, we exclusively considered candidate RNAs that adhered to at least two of the three sets of strict criteria (Extended Data Fig. 3). With this strategy, high-confidence miRNAs were identified in all analyzed dictyostelids (Supplementary Table 1). Examples of miRNAs, predicted pri-miRNA hairpins, and number of reads of the mir-5p and mir-3p for each organism, are illustrated in Fig. 1b. Supplementary Data 1 shows the details of the full set of identified miRNAs.

The analyses show that the number of different miRNAs detected in the different Dictyostelia species varies greatly, and that also expression levels seem highly variable (Supplementary Table 2). As in *D. discoideum*, most miRNAs in the other dictyostelids are expressed from a single, usually intergenic, region. For both asu-mir-1204 and dfa-mir-1220, however, we could detect several paralogs with the exact same miRNA-duplex in the *A. subglobosum* and *D. fasciculatum* genomes (Supplementary Table 2). It is still unclear from which loci these particular miRNAs originate. Besides these, no further intra- or interspecies homology of miRNAs was found. It is worth noting that, due to the high level of stringency, not all previously identified miRNAs in *D. discoideum* were detected by the present search approach. In sum, miRNAs are present in all phylogenetic groups of Dictyostelia, a lineage which originated 600 million years ago.

### miRNAs are present in true unicellular amoeba

The discovery of miRNAs in all tested dictyostelid social amoebae raised the question whether they are exclusive to aggregative multicellular amoebae or if they were already present in unicellular ancestors. The importance of miRNAs for developmental processes is indicated by their upregulation during *D. discoideum* development^30,31^ To address the question if unicellular amoebae have miRNAs, we first sequenced small RNAs from *Physarum polycephalum* (Fig. 2a). This Myxomycete is considered to be the closest sequenced unicellular relative to Dictyostelia and features a complex life cycle. It forms a multinucleate plasmodium but is not multicellular and lacks the social interaction specifying Dictyostelia^27,34^. Stringent analyses of the sequencing data showed that *P. polycephalum* encodes seven distinct miRNAs (Fig. 2b and Supplementary Table 3 - 4). Importantly, this indicates that the presence of miRNAs is not restricted to amoebae that feature multicellular aggregative development. In contrast to Dictyostelia, most of the small RNAs in *P. polycephalum* are 20 nt long (Extended Data Fig. 2). The identified *bona fide* miRNAs, however, are mostly 21 nt in length, just like the Dictyostelia miRNAs (Extended Data Fig. 4).

**Fig. 2.**
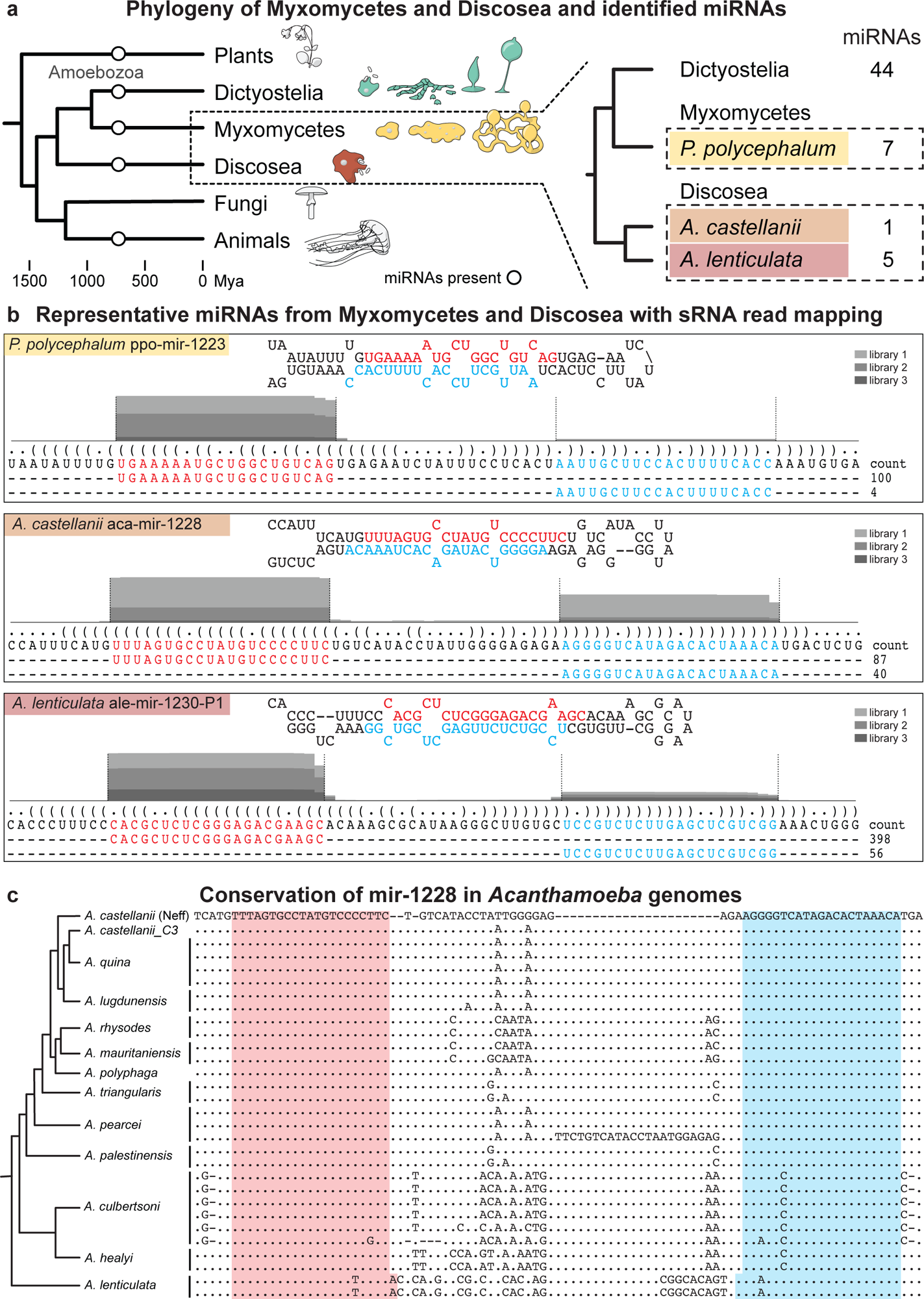
miRNAs identified in *Physarum polycephalum* and *Acanthamoeba* unicellular amoebae. **a** Simplified phylogeny of eukaryotes, focusing on the Amoebozoan groups Myxomycetes and Discosea. The specific organisms included in this study, with the number of discovered miRNAs in each species, is displayed to the right. **b** Examples of miRNAs from *P. polycephalum*, *A. castellanii* and *A. lenticulata*. For further description of panel b, see figure legend for Fig. 1b. For detailed information, see Supplementary Data 2. **c** Alignment of the predicted mir-1228 hairpin sequence identified in the indicated species. Predicted miRNA-5p and miRNA-3p positions are indicated by pink and cyan background, respectively. The start- and end-positions of the miRNA-5p and miRNA-3p are estimated based on the cleavage observed in *A. castellanii* (Neff strain), except for *A. lenticulata*, which has a 1-nt shifted cleavage. Simplified phylogenetic dendrogram is shown on the left, based on Corsaro, 2020^64^.

The presence of miRNAs in *P. polycephalum* shows that they are not exclusive to Dictyostelia in the group of Amoebozoa. Therefore, we investigated whether miRNAs were also present in true unicellular amoebae that are evolutionarily even more distant to dictyostelids^34^. We sequenced small RNAs from *Acanthamoeba castellanii* and *Acanthamoeba lenticulata* and identified one and five miRNAs, respectively (Fig. 2, Supplementary Data 2, Supplementary Table 3). In *A. lenticulata*, two paralogs could be identified for both ale-mir-1228 and ale-mir-1230, and additionally an ortholog of ale-mir-1228 in *A. castellanii*: aca-mir-1228 (Supplementary Table 4). To our knowledge, the miRNAs in *P. polycephalum*, *A. castellanii,* and *A. lenticulata* are the first to be reported in Amoebozoa outside of Dictyostelia. Taken together, our data suggests important roles for miRNAs in both uni- and multicellular life.

### miRNAs are conserved within *Acanthamoebae*

Using stringent search criteria for identification of miRNAs in small RNA sequence libraries is powerful. Adding conservation, when possible, gives an additional level of confidence^40^. To further investigate the conservation of the miRNAs found in *A. lenticulata* and *A. castellanii*, we searched the available *Acanthamoebae* genomes (see Methods). Orthologs of the miRNAs, as well as the predicted pre-miRNAs, were identified in most of the available genomes of species that phylogenetically lie between *A. castellanii* and *A. lenticulata* (Fig. 2c). However, we could not find miRNA or precursor sequences in more distant *Acanthamoeba* species. As expected, mir-5p and mir-3p sequences are well conserved while the connecting sequences tend to be more variable. Notably, the nucleotide change in mir-3p sequences in *Acanthamoeba healyi* (compared to *A. castellianii*) is placed in a predicted bulge in the miRNA duplex, and the two single nucleotide changes in the mir-5p and mir-3p in *A. lenticulata* are compensatory. Hence, these changes should have little impact on the pre-miRNA and miRNA duplex structure. The mir-1228 sequences in *Acanthamoeba cultbertsoni* feature the same nucleotide change as *A. healyi,* however, one of the sequences contains two more nucleotide changes in the miRNA duplex, predicted to cause additional bulges. In summary, we provide evidence that numerous *Acanthamoeba* species harbor conserved miRNAs.

### miRNAs are not required for multicellularity

The presence of miRNAs in true unicellular amoebae as well as in social amoebae that can go through aggregative multicellularity, indicate that miRNAs are not required for transition from uni- to multicellular life. This is supported by the observation that depleting miRNAs by knocking out *drnB*^30,32^ did not visibly affect development (Fig. 3a). In contrast, the *drnB^-^* cells have a slower growth rate in axenic media, suggesting that miRNAs have a regulatory role in the unicellular life stage of the amoebae (Fig. 3b). Hence, miRNAs do not appear to be required for the transition to multicellar life and have no crucial impact on multicellular development of *D. discoideum*.

**Fig. 3.**
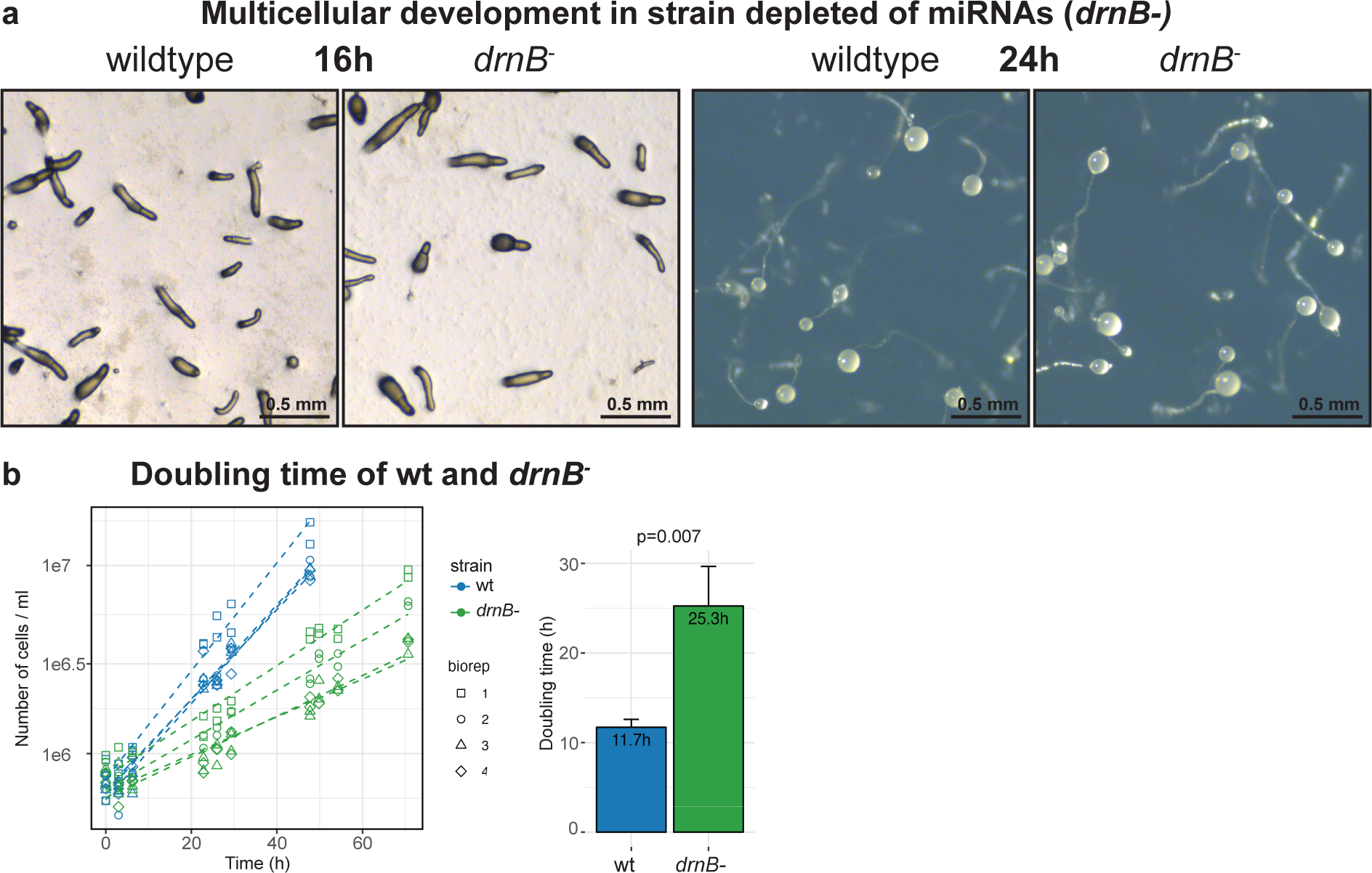
Effect of loss of miRNAs on multicellular development and growth in *D. discoideum*. **a** Images of development for *D. discoideum* cells where *drnB* has been disrupted (*drnB*^-^), which causes depletion of the miRNAs (Extended Data Fig. 5 and Liao *et al.*^32^ for detailed analysis). Images of development of the wildtype cells shown for comparison. Development was initiated 16h or 24h prior to imaging. Photos show slugs (16h) and fruiting bodies (24h). **b** Growth curves of wt and *drnB*^-^ *D. discoideum* cells grown in liquid medium. Four biological replicates for each strain are shown. Doubling times are calculated from the growth curves and summarised in the bargraph. P-value from two-tailed t-test comparing the two strains (n=4).

### miRNA conserved within group 4 Dictyostelia

The discovery of high-confidence miRNAs in all investigated Dictyostelia suggests that they are important regulatory small non-coding (nc)RNAs in these social amoebae. Even though miRNAs do not appear to have a major impact on overall *D. discoideum* development (Fig. 3), they could still have been involved in the evolution of the social behavior in dictyostelid amoebae. If so, we may expect some miRNAs to be conserved throughout Dictyostelia. Surprisingly, we initially failed to find evidence for miRNA gene conservation between any of the species. However, a closer inspection of the small RNA sequences from *D. firmibasis* revealed a homolog of the *D. discoideum* miRNA ddi-mir-1177-5p. We reasoned that our initial failure to identify this homology was due to the quality of the *D. firmibasis* genome assembly (Supplementary Table 5) that prevented mapping of the ddi-mir-1177-5p homolog to the genome. Therefore, we carried out *de novo* nanopore long-read sequencing of the genome to obtain high coverage also of intergenic regions, which is challenging due to high AT-content (86.2% in intergenic regions for *D. discoideum*) (see Methods). This reduced the number of contigs from 997 to 8 (the same number of contigs as for *D. discoideum*) and also has BUSCO scores similar to that of *D. discoideum*, with 92.9% and 90.2% single copy orthologs in *D. firmibasis* and *D. discoideum,* respectively (Supplementary Table 5). Upon re-mapping of the small RNAs to the new improved *D. firmibasis* genome sequence, the identified miRNAs increased from six to ten (Supplementary Table 6, Supplementary Data 3). As we had hypothesized, one newly mapped miRNA, dfi-mir-1177, was an ortholog of the *D. discoideum* miRNA ddi-mir-1177. Both of these miRNA genes lie in the intergenic region between (orthologs of) *D. discoideum* genes DDB_G0287869 and DDB_G0287713, an unresolved region in the previous version of the *D. firmibasis* genome. We next asked whether sequences for this miRNA were present in other group 4 dicyostelids, i.e. *Dictyostelium citrinum* and *Dictyostelium intermedium.* We searched their genomes for orthologs to DDB_G0287869 and DDB_G0287713 and PCR-amplified and sequenced the unresolved intergenic regions. In both cases, homologous sequences to ddi-mir-1177 and the miRNA hairpin were identified (Fig. 4a, Extended Data Fig. 6a). Further analyses showed that while the mir-1177-5p is perfectly conserved between the analyzed group 4 dictyostelids, the mir-1177-3p has one or two nucleotide changes as compared to ddi-mir-1177-3p in *D. discoideum* (Extended Data Fig. 6a). In analogy to the conserved *Acanthamoeba* miRNA, these changes are located in bulges or predicted to maintain base-pairing. Hence, they should not affect the structure of the miRNA-duplex. The remainder of the sequences, between mir-5p and mir-3p, are not as well conserved but still predicted to form pre-miRNA stem-loops (Fig. 4a). Expression of the miRNA from all four species was verified by Northern blot. By contrast, in line with RNA-seq results and genome analyses, no hybridization signal was detected from the more distantly related *Dictyostelium lacteum* and *A. subglobosum* (Fig. 4b). These results show that at least one miRNA appears to be conserved exclusively within group 4 dictyostelids.

**Fig. 4.**
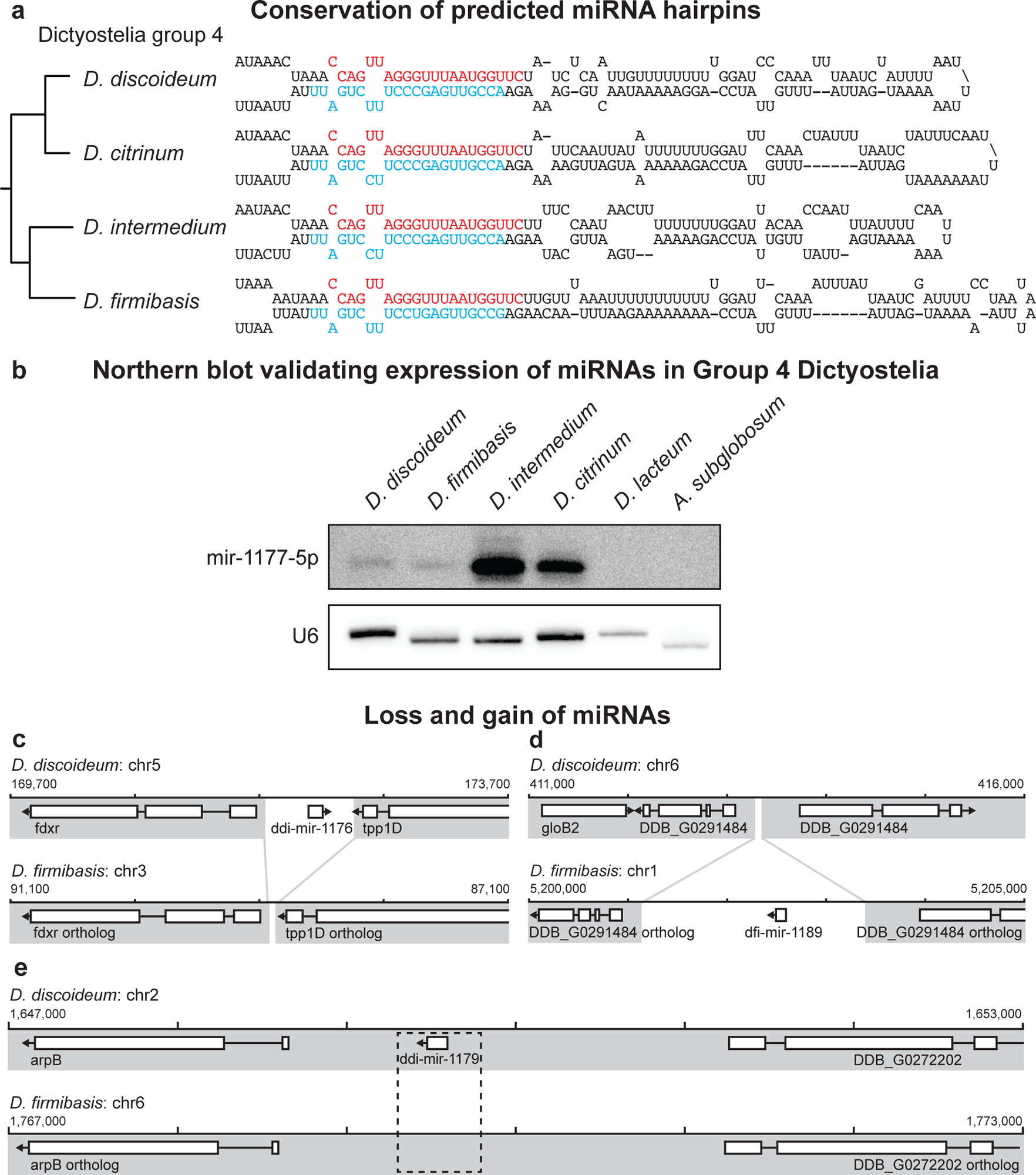
Conservation of mir-1177 in group 4 species of Dictyostelia. **a** Predicted secondary structure of the mir-1177 orthologs found in *D. discoideum*, *D. citrinum*, *D. intermedium*, and *D. firmibasis*,with a simplified phylogeny shown on the left, based on Schilde *et. al*, 2019^34^. The DNA sequences from *D. citrinum* and *D. intermedium* were resolved by sequencing the genomic region in which the ortholog is located. The *D. firmibasis* ortholog was resolved by whole genome sequencing. Positions of the mir-1177-5p and mir-1177-3p are indicated in red and cyan, respectively. **b** Expression of mir-1177-5p in the group 4 dictyostelids *D. discoideum*, *D. firmibasis*, *D. intermedium,* and *D. citrinum* verified by Northern blot. *D. lacteum* or *A. subglobosum,* outside group 4, showed no expression signal. Loading control by probing for U6 spliceosomal RNA. Both panels show different sections of the same membrane at different exposure times. **c-e** Detailed examples of syntenic regions containing miRNAs, suggesting different evolutionary histories. In some instances, the intergenic regions containing the miRNA are lacking in the other species, such as in **c** and **d**. In other synteny blocks, the size of the intergenic region is preserved, but extensive conservation of the miRNA sequence, or expression of sRNAs from this region, is not supported (as in **e)**.

### Evolution of miRNAs in *D. firmibasis* and *D. discoideum*

Considering the relatively close evolutionary positions of *D. discoideum* and *D. firmibasis*, we were surprised that, except for ddi-mir-1177, none of the other miRNAs were conserved between these dictyostelids. To approach the evolution of miRNAs in these species, the two genomes were compared to identify synteny blocks. Those that contained miRNA sequences in either species were then aligned for analyses (Extended Data Fig. 6b). Interestingly, for some *D. discoideum* miRNAs, the orthologous intergenic region in *D. firmibasis* is reduced in size indicating the absence of the whole miRNA locus (Fig. 4c). The same was observed for some *D. firmibasis* miRNAs, such as dfi-mir-1192 (Fig. 4d). Here, the orthologous intergenic region in *D. discoideum* is ∼2kb shorter, completely lacking an orthologous gene for the miRNA. In other cases, the orthologous intergenic regions are of similar length, and the miRNA seems to have evolved through multiple nucleotide substitutions, as opposed to insertion / deletion of the entire locus. This is illustrated by the *D. discoideum* specific miRNA ddi-mir-1179. Here, the orthologous intergenic region in *D. firmibasis* also has the potential to be transcribed into an imperfectly paired hairpin. This might suggest how this miRNA has emerged in *D. discoideum,* or alternatively how it has been degenerated in *D. firmibasis* (Fig. 4e, Extended Data Fig. 6c). Taken together, by comparing the evolutionarily relatively closely related genomes of *D. discoideum* and *D. firmibasis*, we gain insights into the evolutionary history of miRNAs, which seems to depend on gains and losses of miRNA “modules” as well as modification of stem-loop structures.

### Features of Amoebozoan miRNAs

The identified miRNAs in the different amoebae allowed for analyses of shared features. One general trend is the identity of the 5’ end, most frequently a uridine or, to a lesser extent, an adenosine or cytidine, as in plant and animal miRNAs (Fig. 5a). The majority of miRNAs in the amoebae are derived from intergenic regions (Extended Data Fig. 7a) which are AT-rich in social amoebae. However, the miRNAs are considerably less AT-rich, a pattern seen in other ncRNAs in the AT-rich genomes of dictyostelids^41,42^ (Fig. 5b). Although the majority of miRNAs originate from intergenic regions, miRNAs in *P. pallidum* are mostly intronic, and in both *A. subglobosum* and *D. fasciculatum*, some miRNA sequences are in antisense orientation to annotated coding regions (Extended Data Fig. 7a). The miRNAs are derived from predicted pre-miRNAs, varying in size within species but most often 75-125 nt in length, similar to the size of pre-miRNAs in plants (Fig. 5c, Extended Data Fig. 7b). As seen previously, not all small RNAs in amoebae are miRNAs. In *A. subglobosum,* the majority of the 21 nt RNAs are miRNAs, but in the other species, RNAs originating from e.g. transposons or unannotated intergenic regions dominate (Extended Data Fig. 7c).

**Fig. 5.**
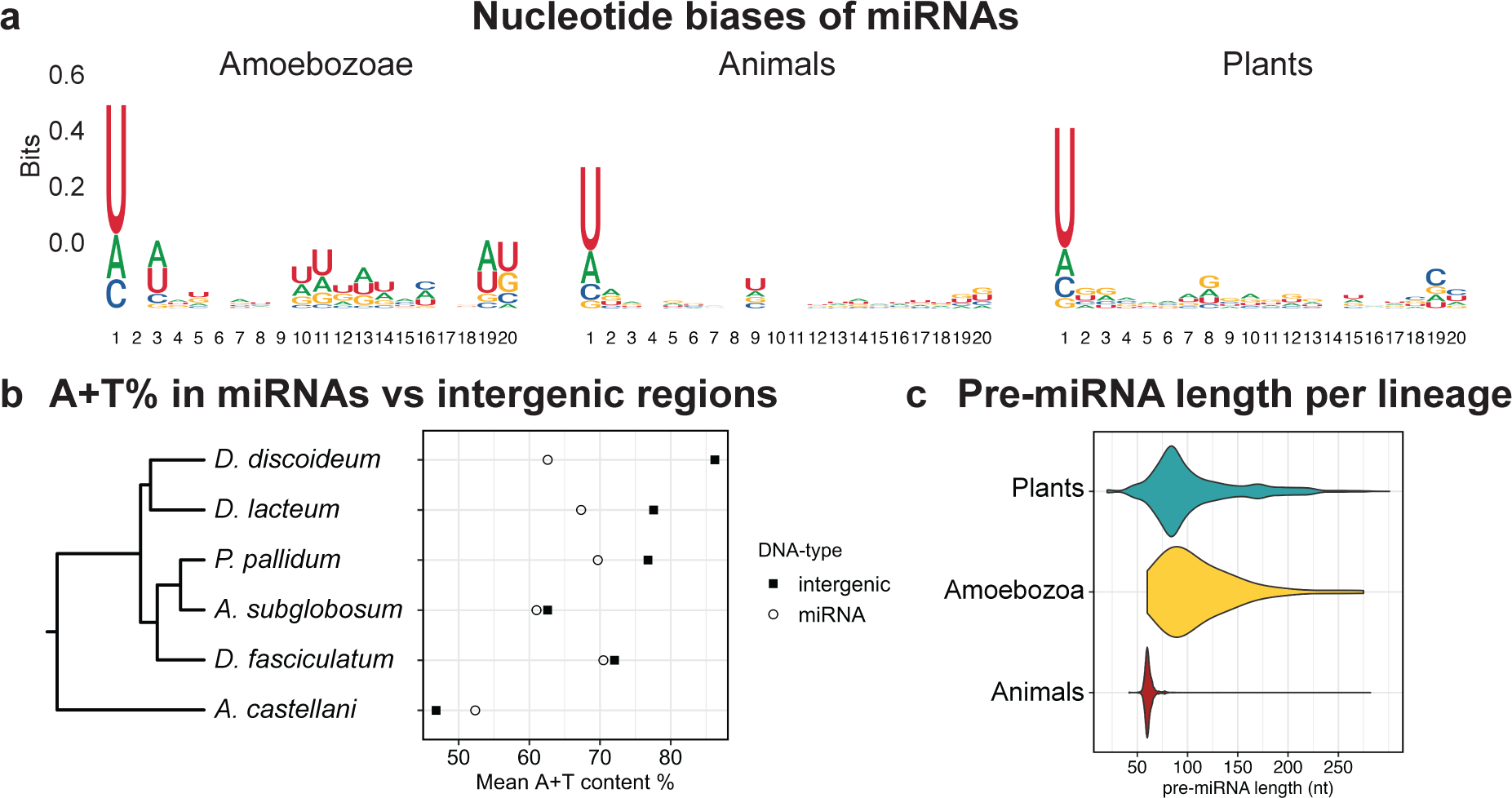
Characteristics of Amoebozoan miRNAs. **a** Sequence logo of the amoebozoan miRNAs identified in this study, as well as animal miRNAs accessed from MirGeneDB^65^, and plant miRNAs from PmiREN^66^. miRNAs from all three major phylogenetic groups show a 5’U preference. **b** AT-content of intergenic regions and of miRNA for the analyzed species for which genome annotations are available. **c** Violin plots of the pre-miRNA lengths in plants, Amoebozoae and animals. Pre-miRNA lengths for each of the amoebozoan species shown in Extended Data Fig. 7b.

### The RNAi machinery in Amoebozoa

Dicers and Argonautes are the most conserved components for processing and function of miRNAs^9,43^. We asked how the evolution of the RNAi machinery matched the evolution of miRNAs in the studied amoebae. In most species, we could identify two Dicers and two or more Argonautes (Fig. 6). Looking more closely at the evolution of Dicer proteins, a single copy of Dicer was likely present prior to the expansion of the dictyostelids, which then duplicated in Group 1, 2, and 4, but not 3, leaving *D. lacteum* with just a single Dicer (Extended Data Fig. 8; See also Kruse *et al.*^44^ for additional discussion). The *Acanthamoebae* appear to have experienced a duplication of Dicer prior to the *A. lenticulata – A. castellanii* split. Concerning the Argonautes, a single copy was most likely present in the last common ancestor of the Amoebozoae. A duplication event prior to the expansion of the dictyostelids resulted in two or more copies of the Argonautes in most of these species (Extended Data Fig. 9). A duplication of the Argonautes is also suggested in the *Acanthamoebae*, leaving both *A. lenticulata* and *A. castellanii* with two Argonaute paralogs. The expansion of the RNAi machinery in the *Acanthamoebae* points towards the appearance of a specialized miRNA processing machinery, which would be in line with the discovery of conserved miRNAs in the *Acanthamoebae*. Interestingly, while the RNA directed RNA polymerases seem to have expanded in the Dictyostelia and *P. polycephalum*, we could not identify any in the *Acanthamoebae* (Extended Data Fig. 10). Taken together, the RNAi machinery is present in all studied amoebozoans and has been expanded by duplications. The flexibility of this machinery is demonstrated by the variable numbers of RNAi-genes in different organisms, underscored by the observation that morphological complexity coincides with an expanded repertoire of RNAi-related genes.

**Fig. 6.**
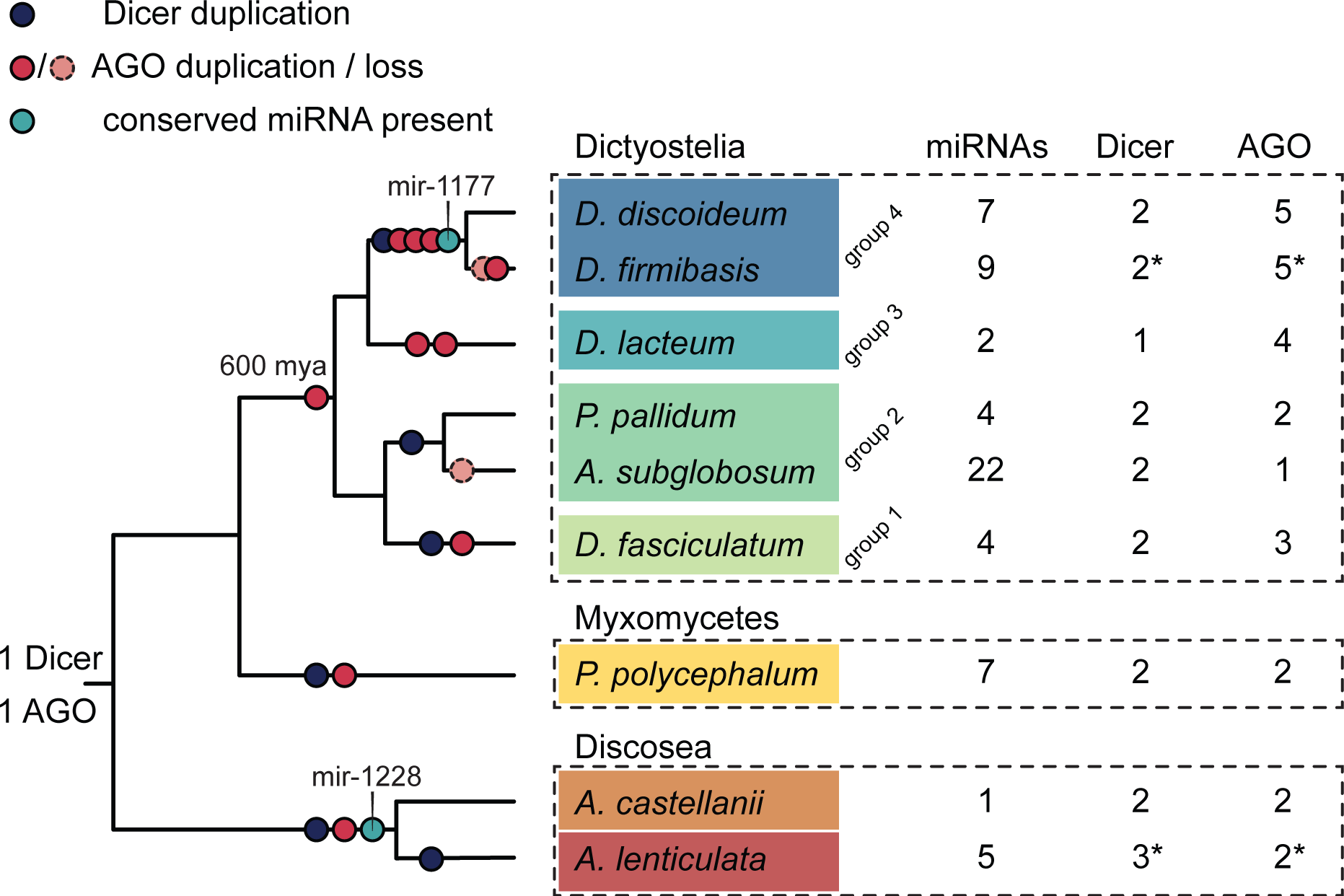
Model of the evolution of the miRNAs, Dicers, and Argonautes in Amoebozoa. Number of miRNAs, Dicers and Argonautes (AGO) identified in each of the species. Asterisk (*) indicates that genome annotations or protein sequences were not available and these orthologs were identified manually. Schematic phylogeny of the Amoebozoans with root of the Dictyostelia at 600 mya according to Heidel *et. al.*, 2011^28^. Duplications / losses of the Dicer and Argonaute machinery are indicated on the dendrogram based on parsimonious interpretation of Extended Data Fig. 8 and Extended Data Fig. 9. For miRNAs where we could detect conservation between species, i.e. mir-1177 and mir-1229, their presence prior to speciation is indicated.

## Discussion

The great majority of miRNAs have been discovered in plants and animals, where they have been estimated to regulate nearly all biological processes. In particular, it was also suggested that miRNAs played a role in the evolution of multicellularity in these lineages^17,24,45^. Hence, most of what we know about biogenesis, function, and evolution of miRNAs stems from studies in plants and animals. Although, to understand the commonality of miRNA regulation and their putative role in evolution of multicellularity, it is essential to investigate how widespread miRNAs are, i.e., in what other organisms they are present. In order to address this, we searched for miRNAs and their associated genes in Amoebozoa, a phylogenetic group positioned between plants and animals, in which a wide variety of both uni- and multicellular lifestyles can be found^25,36^. Until now, high confidence miRNAs had only been described in the social amoeba *D. discoideum*^30–33^. Here, we substantially expanded the number of amoebae carrying miRNAs by sequencing and stringently analyzing the small RNA repertoire of eight Amoebozoans in addition to *D. discoideum*. We hypothesized that miRNAs would be predominantly present in the multicellular social amoebae, supporting the evolution of their multicellular lifestyle. To study this, we included Amoebozoans with a range of lifestyles, including several social amoebae from the monophyletic group Dictyostelia, which all feature aggregative multicellularity, as well as strictly unicellular species. Surprisingly, we identified miRNAs in all sequenced amoebae, including those lacking any sort of multicellular lifestyle, revealing that miRNAs are more widespread than previously believed. Furthermore, the fact that depletion of the miRNAs did not affect aggregation and multicellularity, and that miRNAs are present in true unicellular amoebae show that miRNAs are not required *per se* for transition from uni- to multicellular life (Fig. 2, 3).

### Did miRNAs play a role in the evolution of multicellularity?

Our discovery of miRNAs in unicellular outgroups to Dictyostelia is analogous to the findings of miRNAs in unicellular organisms closely related to animal and plants. This suggests that the evolution of clonal multicellularity in plants and animals, as well as the evolution of aggregative multicellularity in Dictyostelia, did not coincide with the evolution of miRNAs and the miRNA machinery (Fig. 6)^19,20,22^. However, the importance of miRNAs in multicellular processes and development in animals is well documented. Many miRNAs are conserved in bilaterian animals, and mir-100/mir10, is found in almost all multicellular animals^38,46^. Also in plants, some miRNAs are highly conserved, indicating that their ancestor already had miRNAs^47^. Dysregulation of these ancient conserved miRNAs often causes developmental phenotypes, illustrating the significance of these miRNAs for proper multicellular development^48–50^. In contrast to the situation in animals and plants, we could not detect any conserved miRNAs throughout Dictyostelia. In addition, different studies have shown a correlation between the number of miRNAs and increasing complexity, though in plants this correlation is less clear^47,51,52^. We do not see an obvious correlation between the number of miRNAs and complexity in Amoebozoa (Fig. 6). Also, in a mutant *D. discoideum* strain depleted of miRNAs, no effect on multicellular development was observed (Fig. 3). In animals, most miRNAs only partially regulate their targets and rather act as buffers to, for instance, stabilize phenotypes and dampen variability^51^. Perhaps, this is also true for miRNA regulation in Dictyostelia, and more detailed studies of phenotypic effects are required to detect the influence of miRNAs. It would also be useful to investigate how other Dictyostelia respond to depletion of miRNAs.

### When did miRNAs evolve?

It is commonly believed that miRNAs evolved independently in plants and animals, following the expansion of their RNAi machinery^16,19^. Partly, our data agree with this since we observe a low level of conservation of miRNAs indicating that miRNAs evolved independently and several times in Amoebozoa. Also, we see expansions of the miRNA machinery, i.e. genes for Dicers and Argonautes, simultaneously with the appearance of novel miRNAs (Fig. 6). Given these findings, we may argue for the common model that RNAi was present in the last eukaryotic common ancestor where it protected against genomic invaders such as transposons and viruses^9^. Hence, there has been, and is, a strong evolutionary pressure to maintain the RNAi machinery, which could act as a scaffold from which miRNAs can readily evolve in different organisms^14–16,53^. This is further supported by our data showing that a minimal basic miRNA machinery (considering Argonautes and Dicers) can be sufficient, where some Amoebozoans feature only a single Dicer or a single Argonaute. Thus, the convergent evolutionary scenario is possible but the hypothesis of an ancient miRNA machinery cannot be ruled out and is discussed further below.

### How fast do miRNAs evolve?

While we found conservation of some miRNAs between unicellular members of *Acanthamoebae*, and also between the group 4 dictyostelids *D. discoideum* and *D. firmibasis*, the other identified miRNAs were all unique for each species. This indicates a rapid evolution of Amoebozoan miRNAs. To analyze in more detail how some of these miRNAs evolved, we investigated the two most closely related organisms in our dataset, *D. discoideum* and *D. firmibasis*. While some of the miRNAs seem to have been acquired or lost gradually, for others we could identify insertions or deletions of whole miRNA modules (Fig. 4). The fact that these two closely related organisms have such a diversified miRNA repertoire shows that broadly, their individual miRNAs are under weak evolutionary pressure, consistent with the minor phenotypic effects we observed upon depletion of miRNAs in *D. discoideum*. The rapid gain and loss of miRNAs between the two closely related species and the prescence of miRNAs in many eukaryotic lineages could support that miRNAs were present in an ancient eukaryotic ancestor and that the fast turn over of miRNA may obscure their origin. We have recently initiated a study where we will acquire high-quality genomes of closely related amoebae and sequence their small RNAs to obtain a high-resolution picture of mechanisms driving miRNA evolution in Dictyostelia.

Taken together, this work greatly expands the number of miRNAs in Amoebozoans, and the number of Amoebozoans, or protists in general, that feature miRNAs. The presence of miRNAs throughout the Amoebozoan phylogenetic group indicates that miRNAs are much more common outside animals and plants than previously believed, and argues against the idea that miRNAs drive the evolution of multicellularity.

## Methods

### Strains and cell culture

Social amoebae were grown at 22 °C on SM Agar/5 (Formedium) with 0.5% activated charcoal, in association with *Escherichia coli* 281. Cells were allowed to grow until they started clearing the plate from *E. coli*, but before starvation occurred, at which point they were harvested with a cell scraper (Thermo Scientific). Remaining *E. coli* were removed by four consecutive washes with KK2 buffer (2.2 g/l KH_2_PO_4_, 0.7 g/l K_2_HPO_4_) at 400xg, 5 min. One-third of the cells were resuspended in TRIzol Reagent (Invitrogen) and frozen at -20 °C until further processing. The remaining two-thirds of the cells were seeded on non-nutrient agar with 0.5% activated charcoal to allow for starvation and subsequent development of the cells. One-third of the cells were harvested at the sorogen or slug stage (13-20 h post induction of starvation) and one-third of the cells at the sorocarp of fruiting body stage (23-40 h post induction of starvation) into TRIzol Reagent (Invitrogen) and frozen at -20 °C. *Acanthamoebae* were first grown axenically in PYG w/additives (ATTC medium 712 at 30 °C) and subsequently grown and harvested similar to the social amoebae, on SM Agar/5 plates with charcoal, with *E. coli* 281 as food source. *Physarum polycephalum* was grown on oats on 2% agar. Parts of sclerotium were scraped from the plate and resuspended directly into TRIzol Reagent.

For the growth curves of *D. discoideum* wildtype (AX2) and *drnB*^-^ strains, cells were grown axenically in HL5-c (Formedium) with 100µg/ml Pen Strep (Gibco) at 22 °C, shaking at 180 rpm, and quantified in a hemocytometer. For strain numbers and access to strains see attributes file at SRA under BioProject accession number PRJNA972620.

### Small RNA sequencing

RNA isolation was performed according to the TRIzol Reagent user guide (Invitrogen), except with an additional 75% EtOH wash of the RNA pellet. For social amoebae, RNA from different developmental stages was pooled using equal amounts of total RNA. Small RNA sequencing libraries were generated according to the TruSeq Small RNA Library Prep Kit (Illumina). Libraries were sequenced on a NextSeq 500 System (Illumina). Development, RNA preparation and sequencing of the *drnB^-^* knockout strain versus wildtype was performed as in^32^. Differential gene expression analysis was done in R.

### miRNA identification

Quality control of raw sequencing reads was performed with miRTrace^54^, after which they were processed by trimming the sequencing adapters and selecting reads of size 18-35 nt with Cutadapt^55^. Small RNAs were mapped to the reference genome or *E. coli* genome, with a maximum of one mismatch, and multimapping reads placed according to the fractional weighted probability, using Shortstack v3.8.5^56^. Clusters with potential miRNA candidates were selected and further analyzed with an in-house python script available on https://github.com/Bart-Edelbroek/amoeba_miRNAevo. Briefly, in the first round, exploratory analysis was done by mapping reads with no mismatches to each selected cluster and folding the cluster using the ViennaRNA RNAlib-2.6.2 python package^57^. If the cluster could be folded into a hairpin-structure, the positions of the mir-5p and mir-3p were defined based on the mapped reads. Clusters that yielded a hairpin and had at least a 30% 5’ precision of mir-5p and mir-3p were selected for a he second round of analysis, similar to the first, but with additional plotting the read-mapping density of the cluster and a 2D representation of the hairpin structure. The characteristics of the hairpin as well as precision of the miRNA-duplex were calculated according to different miRNA-annotation criteria. miRNA-candidates were selected if they passed at least two out of three sets of stringent criteria, outlined in Extended Data Fig. 3. Candidates were manually inspected prior to annotation of *bona fide* miRNAs, to ensure the maximum level of confidence in the annotated miRNAs.

### Analysis of miRNAs

Analysis of the identified miRNAs was performed in R. Non-coding RNA regions were identified and annotated in the genome using Infernal 1.1.4 with the Rfam covariance model filtered to amoebozoa. Transposable elements were identified and annotated in the genome using tblastx (evalue cutoff 10^-15^) with reference transposable element sequences from Repbase (Genetic Information Research Institute) filtered to the studied organisms.

### Whole genome sequencing of *D. firmibasis*

For genomic DNA isolation, 6*10^8^ *D. firmibasis* cells were harvested from clearing SM Agar/5 plates and washed five times with 50ml PDF buffer [20 mM KCl, 9.2 mM K_2_HPO_4_, 13.2 mM KH_2_PO_4_, 1 mM CaCl_2_, 2.5 mM MgSO_4_, pH 6.4]. Genomic DNA was isolated with QIAGEN 100/g Genomic-tips, according to the ‘Cell cultures’ protocol, but with 0.2 mg/ml RNase A added to the G2 buffer. Short DNA was filtered out with the PacBio Short read eliminator kit. Library preparation was performed with 1.5 µg of filtered genomic DNA using the SQK-LSK112 nanopore sequencing kit. 100 ng of final prepared library was loaded on a MinION Mk1C with R10.4 flowcell and sequencing was performed for 72 hours. Basecalling, adapter trimming and read-splitting was performed with Guppy v6.3.2. The data was filtered using Filtlong v0.2.1 to retain the best 5 Gbp of data and remove reads smaller than 1000 bp. Assembly was done using Flye v2.9.1 in nano-hq mode^58^, and two rounds of long-read polishing were done using Medaka v1.7.2. The filtered reads used for assembly can be accessed from NCBI BioProject PRJNA972620. The assembly used for downstream sRNA mapping and miRNA discovery can be downloaded from https://doi.org/10.5281/zenodo.7937210.

### Sequencing of the conserved ddi-miR-1177 region

*D. citrinum* and *D. intermedium* cells were grown at 22 °C on SM Agar/5 (Formedium) with *Klebsiella aerogenes* as food source. DNA was isolated as described previously^59^. Primers for amplification of homologs of the ddi-mir-1177 region were designed based on orthologs of the genes up- and downstream of ddi-mir-1177 (DDB_G0287869 and DDB_G0287713 respectively). The intergenic regions homologous to ddi-mir-1177 were amplified by PCR containing 2 µl of lysate, 1U Taq polymerase (Thermo Scientific), 1x reaction buffer, 2 mM MgCl_2_, 0.2 mM dNTPs, 0.2 µM of p1301 (CCTAATGATAAACCAACAAC) and p1330 (ATACATACTGTTCCCAAAG) for *D. citrinum*, and 0.2 µM of p1331 (CCACAGATCAAAAACGTTCA) and p1332 (CCCAAAGACAAACCAACA) for *D. intermedium*, in a final volume of 40 µl. The PCR was performed as follows: 60 s at 95 °C; [15 s at 95 °C; 30 s at 46 °C or 50 °C for *D. citrinum* or *D. intermedium* respectively; 120 s at 62 °C] x35, followed by 7 min at 62 °C. The amplified products were gel purified (GeneJET Gel Extraction Kit; Thermo Scientific) and TA-cloned (InsTAclone PCR Cloning Kit; Thermo Scientific). Plasmid DNA from positive clones was isolated (GeneJET Plasmid Miniprep Kit; Thermo Scientific) and Sanger-sequenced (Macrogen). The resulting sequences were aligned to the ddi-mir-1177 and dfi-mir-1177 intergenic region.

### mir-1177 Northern

Dictyostelid strains were grown in association with *K. aerogenes* at 22°C on SM Agar/5 (Formedium). After the majority of bacteria had been consumed, upon onset of development, the plates were harvested into TRIzol Reagent (Invitrogen), and RNA was isolated according to the user guide. 30 µg RNA samples were denatured in 47.5% formamide at 70°C for 5min. RNA was separated on a 12.5% acrylamide gel (7 M Urea, 1x TBE) at 12W. In addition to the RNA samples, 0.5 µl of Decade RNA marker (Invitrogen) was loaded on the gel. Transfer, probing, and washing was performed as described before^60^, with probe p549 specific for miR-1177-5p (GAACCATTAAACCCTAACTGG) or probe p1339 for U6 snRNA (TGCTAATCTTCTCTGTATCGTT). Northern blots were exposed for up to 72h to BAS-IP MS Phosphorimaging plates (Cytiva) and imaged by Amersham Typhoon (Cytiva).

### Synteny search and alignment of miRNA synteny blocks

The *D. firmibasis* genome assembly was compared to the *D. discoideum* reference genome, and synteny blocks were identified using Satsuma2. Blocks within 5000 bp of each other were merged and the synteny blocks containing miRNAs were visualized in R. To more closely study the evolution of the miRNAs, the synteny blocks with miRNAs were aligned using Clustal Omega.

### Phylogeny of RNAi machinery

Orthofinder v2.5.2 was run with the protein fasta files available for the amoebae included in this study, to find orthogroups containing the Argonaute and Dicer proteins^61^. Using the identified homologs, the genomes of *D. firmibasis* and *A. lenticulata* were searched using tblastn with a maximum e-value of 10^-15^ to identify regions of interest containing Argonaute or Dicer homologs. Conserved domains in the homolog candidates were identified using PfamScan. Sequences were aligned with MAFFT v7.520^62^, trimmed with trimAl v1.4.rev15, and phylogenetic consensus trees were constructed using IQ-TREE v2.2.2.3 with the Blosum62 amino-acid exchange rate matrix^63^.

## Data availability

All sequencing data generated in this study can be accessed at NCBI BioProject PRJNA972620. sRNA sequencing data of different Amoebozoan species for miRNA discovery are available with BioSample accessions SAMN35084086 - SAMN35084094. sRNA sequencing data to compare the *D. discoideum drnB*^-^ knockout strain with wildtype are available with BioSample accessions SAMN35084095 - SAMN35084098. Third-generation sequencing data of *D. firmibasis*, used to assemble the genome, is available with BioSample accession SAMN35084099. The *D. firmibasis* genome assembly used in this study is deposited at https://doi.org/10.5281/zenodo.7937210. Data used for analysis of miRNAs, synteny between *D, firmibasis* and *D. discoideum*, and differential miRNA expression is available at https://github.com/Bart-Edelbroek/amoeba_miRNAevo/tree/main/data_in. Data generated such as miRNA fasta files and tables with miRNA characteristics, are available at https://github.com/Bart-Edelbroek/amoeba_miRNAevo/tree/main/data_out.

## Code availability

For initial curation of the miRNA candidates, we used a python script available at https://github.com/Bart-Edelbroek/amoeba_miRNAevo. A folder with scripts and sample data to perform an example run is provided in the repository. Downstream analysis of the miRNAs, genome synteny analysis, and differential miRNA expression analysis was performed in R and the code is provided at https://github.com/Bart-Edelbroek/amoeba_miRNAevo. Further provided in the repository are the plots generated in the analysis, and helper scripts used for generating input data.

## Acknowledgements

We would like to thank Jon Jerlström-Hultqvist for help with *D. firmibasis* sequencing and genome assembly; Gernot Glöckner (deceased) and Pauline Schaap for help with initiating this study; Wolfgang Marwan and Lionel Guy for providing *P*. *polycephalum* and *Acanthamoeba* strains, respectively. We are also grateful to Andrea Hinas, Pauline Schaap and Gerhart Wagner for critical reading and commenting on the manuscript. Sequencing was performed by the SNP&SEQ Technology Platform in Uppsala. The facility is part of the National Genomics Infrastructure (NGI) Sweden and Science for Life Laboratory. The SNP&SEQ Platform is also supported by the Swedish Research Council and the Knut and Alice Wallenberg Foundation. Computations and data storage were enabled by resources provided by the National Academic Infrastructure for Supercomputing in Sweden (NAISS) at UPPMAX, funded by the Swedish Research Council through grant agreement no. 2022-06725. This work was supported by grant no. 2021-05793 from Swedish Research Council (Vetenskapsrådet) to F.S.

## Author information

### Contributions

Author contributions: B.E., J.K., A.A.N., L.E., M.R.F. and F.S. participated in the design of the project. B.E. performed most experiments and data analysis. B.E. and I.B. prepared sRNA sequencing libraries, and I.B. performed the sequencing. Z.L. prepared RNA for mRNA and sRNA sequencing of the *drnB^-^* knockout strain. T.L. monitored growth of *drnB^-^* and wt cells. B.E. and J.K. generated *D. firmibasis* gDNA libraries and performed genome sequencing. J.K. assisted in bioinformatic analysis. M.R.F. and F.S. supervised the study. B.E. prepared figures, and together with F.S. drafted the manuscript.

### Corresponding authors

Correspondence to Bart Edelbroek or Fredrik Söderbom

## Competing Interests / Ethics declarations

The authors declare no competing interests

---

**Extended Data Fig. 1.**
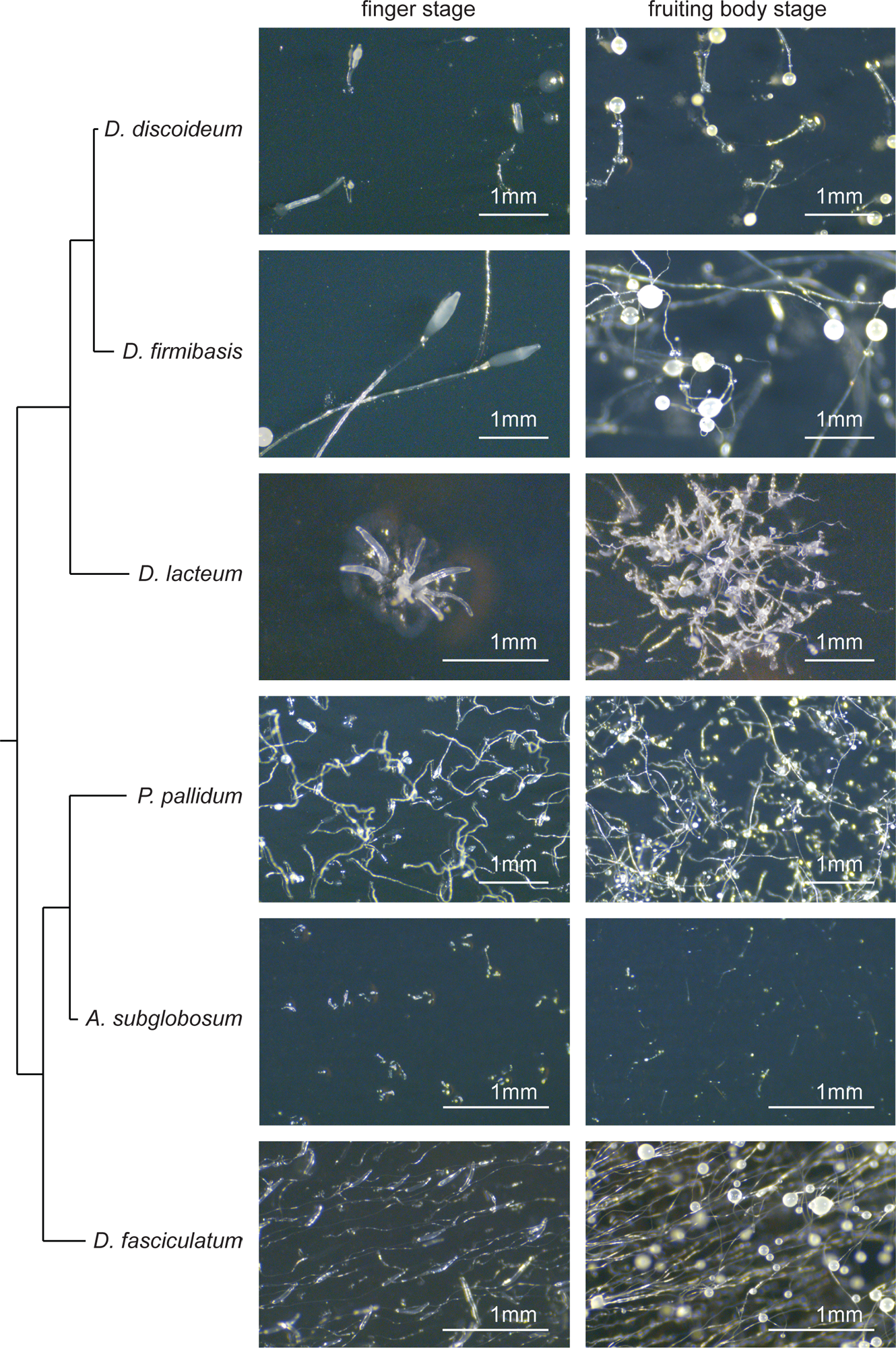
Phenotypes of studied Dictyostelia species at different stages of multicellular development. Small RNAs were sequenced from a mix of vegetative cells (not pictured) and during stages of multicellular development: the finger/slug stage (first column) and fruiting body stage (second column). Each of the social amoebae display distinct phenotypes at the different stages.

**Extended Data Fig. 2.**
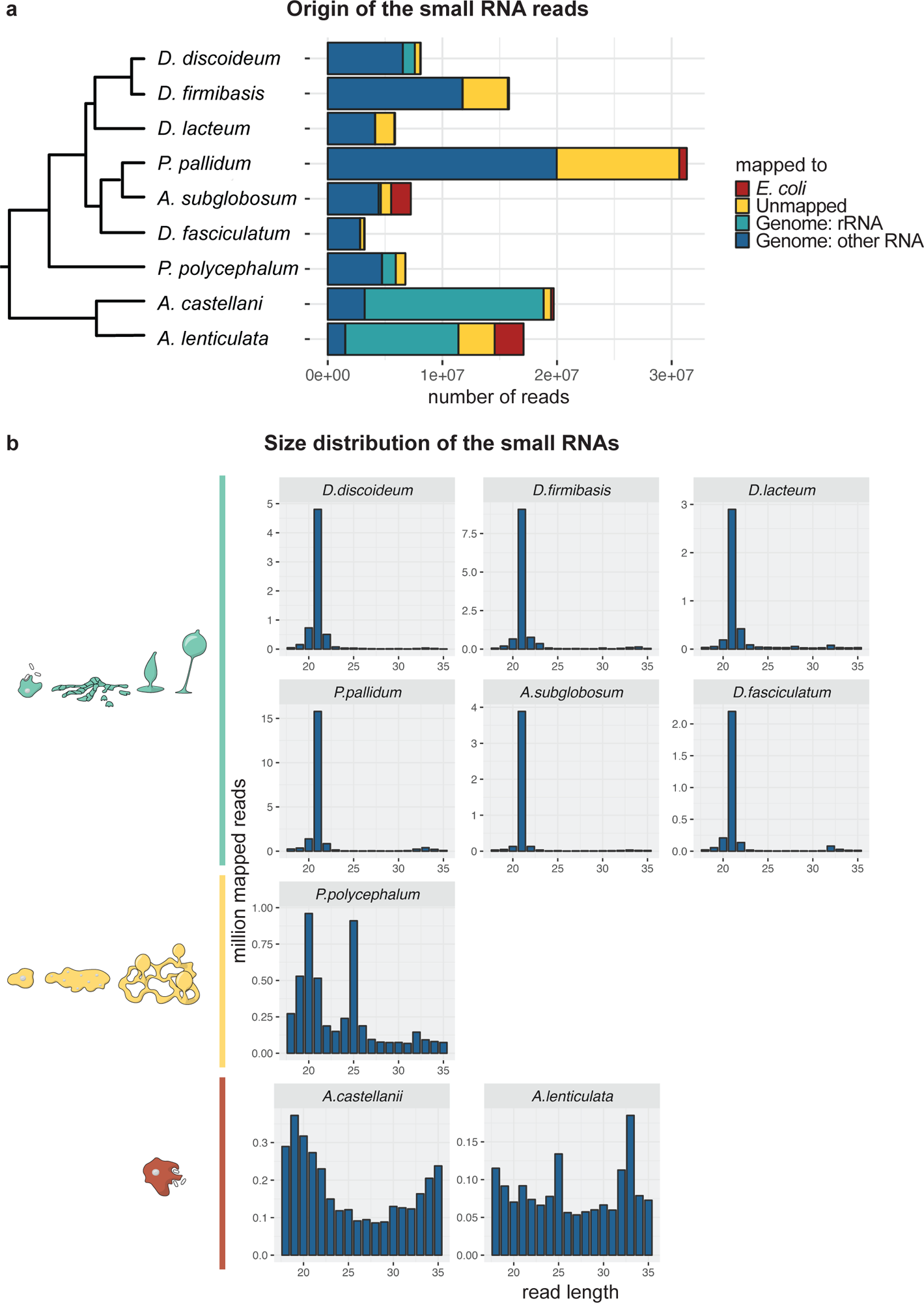
Origin and length profile of isolated small RNAs. **a** Number of reads from small RNA sequencing mapping with maximum one mismatch to the food bacterium *E. coli* or the genomes of the indicated amoebae for which the reads were further subdivided into sequences mapping to the ribosomal RNA (Genome_rRNA) or any other part of the genome (Genome_otherRNA). Reads that could not be mapped are marked as Unmapped. Number of reads are displayed in power of 10. **b** sRNA length profile of reads mapping to the genome, but not the ribosomal RNA (dark-blue in **a**). The characteristic phenotypes of the different amoebae included in the study are shown on the left, with Dictyostelia in green, Myxomycetes in yellow, and Discosea in red.

**Extended Data Fig. 3.**
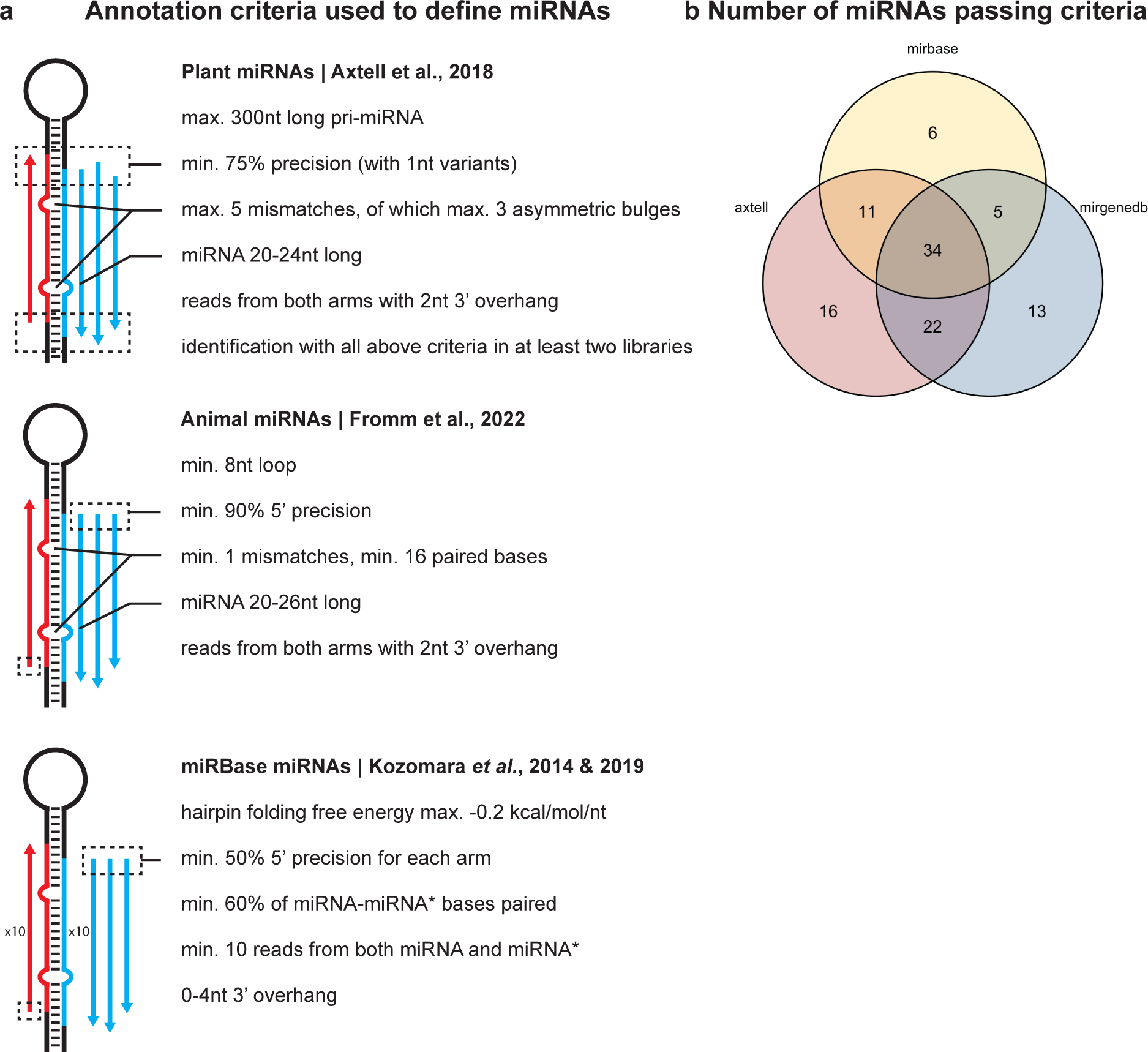
Criteria used for miRNA annotation. **a** Graphical summary of the different sets of miRNA annotation criteria used in this study. ‘Precision’ is the percentage of miRNA-5p and miRNA-3p reads (e.g. ‘precise reads’) versus total reads mapped on the miRNA-hairpin, and is calculated differently for the different sets of criteria. For plant miRNAs (Axtell et al., 2018^37^), all reads that map to the miRNA-5p or miRNA-3p with 1-nt variants for both the 5’ and 3’-ends are considered ‘precise’. For animal miRNAs (Fromm et al., 2022^65^), only reads that map precisely to the 5’-end of the miRNA-5p or miRNA-3p are included. 3’-end variation is allowed. For miRBase miRNAs (Kozomara et al., 2014 & 2019^2,39^), reads are also required to map precisely to the 5’-end of the miRNAs. Here however, calculations are done on the miRNA-5p-arm and miRNA-3p-arm separately, yielding two precision percentages per miRNA, both of which have to be larger than 50% **b** Venn diagram of number of miRNAs that passed all criteria for plant miRNAs (Axtell, red), animal miRNAs (mirgenedb, blue), or miRBase miRNAs (miRbase, yellow). Numbers based on data from *D. discoideum*, *D. firmibasis*, *D. lacteum*, *P. pallidum*, *A. subglobosum*, *P. polycephalum, A. castellanii*, and *A. lenticulata*.

**Extended Data Fig. 4.**
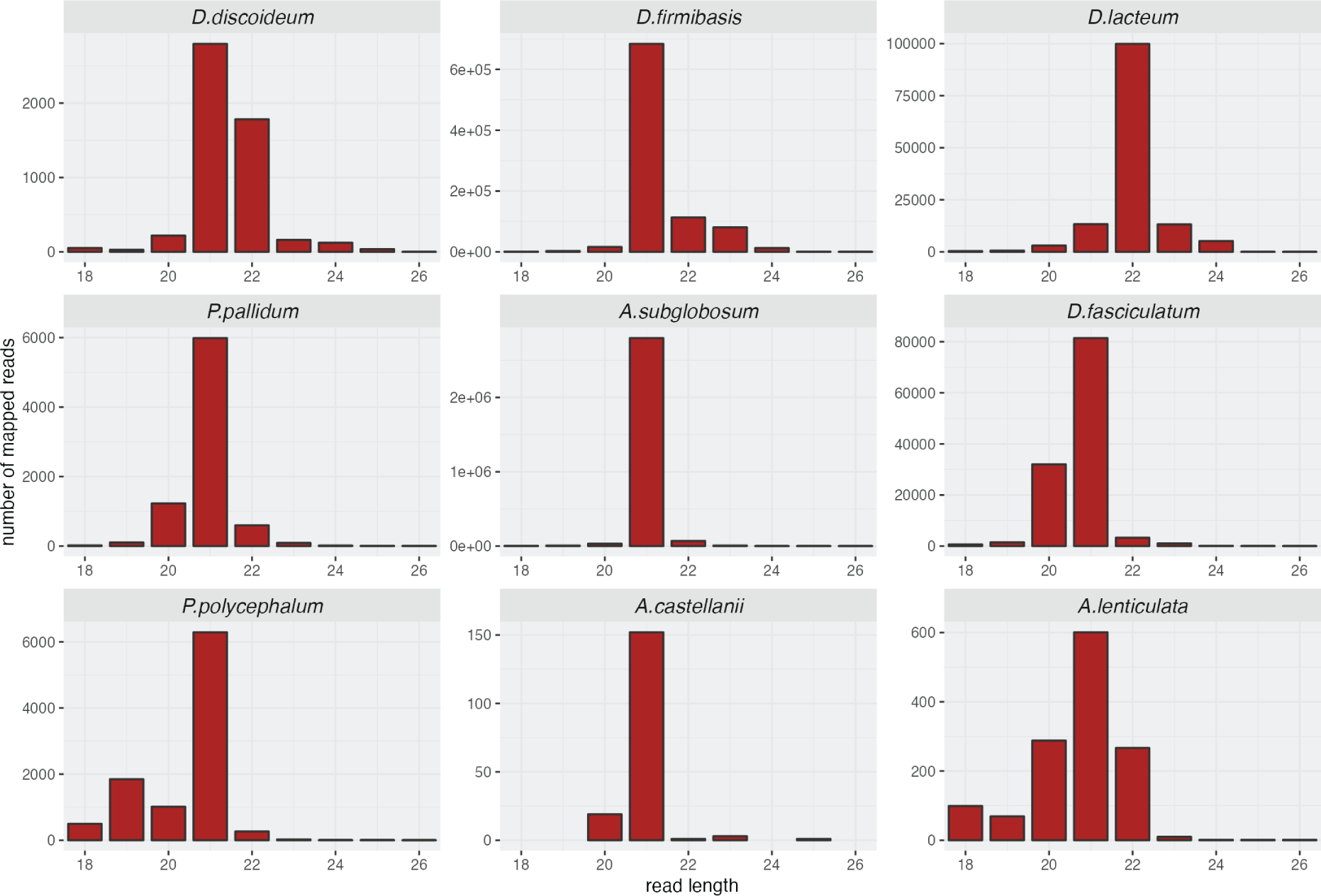
Length distribution of sRNAs mapping to miRNA-loci. Histogram of read lengths mapping to loci of miRNA-hairpins. For *D. firmibasis*, mapping was done to the de-novo sequenced genome.

**Extended Data Fig. 5.**
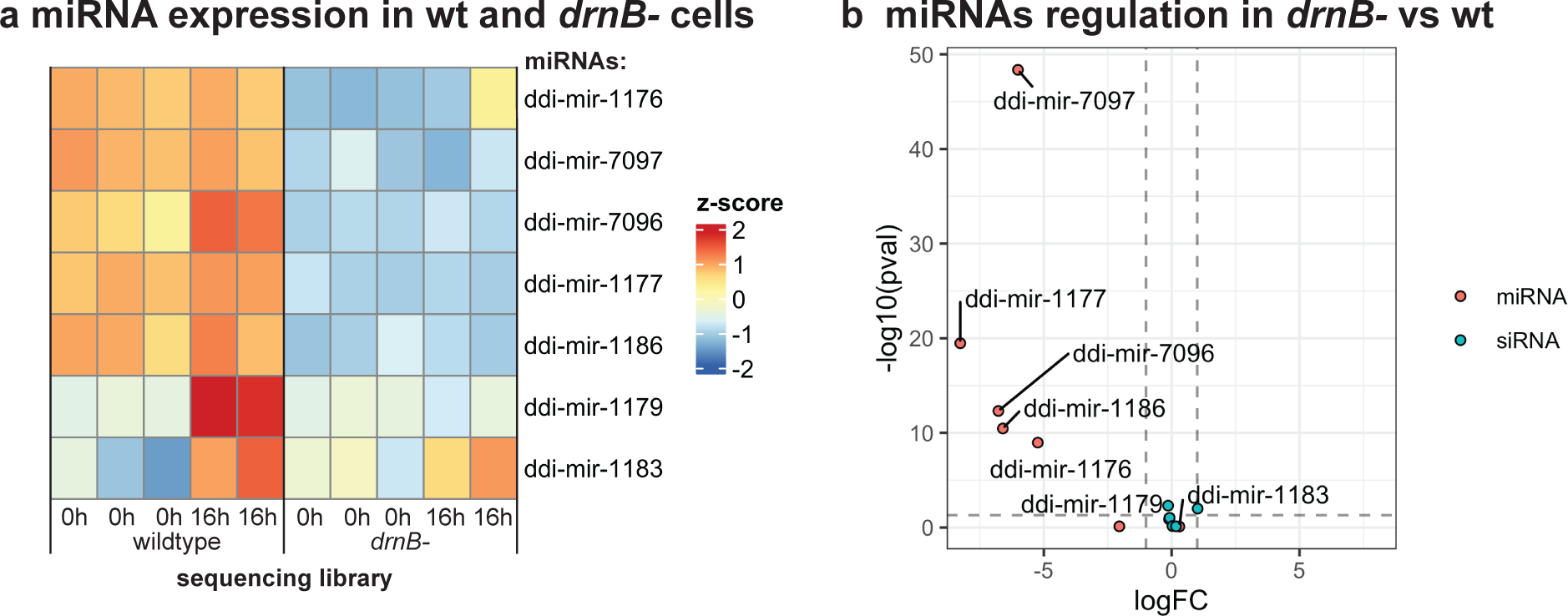
Analysis of loss of miRNAs in *drnB*^-^ strain. **a** Heatmap showing expression of miRNAs in *D. discoideum* wildtype (wt) and *drnB*-strains in vegetative cells (0h) and cells during multicellular development (16h). **b** Volcano plot of miRNAs and siRNAs in *drnB*^-^ cells versus wt. logFC indicates the extent to which the sRNAs are up- or down-regulated, with vertical dashed lines marking a fold change of 0.5 and 2, respectively. -log10(pval) shows the significance with higher values for more significantly regulated sRNAs. The horizonal dashed line marks a p-value of 0.05.

**Extended Data Fig. 6.**
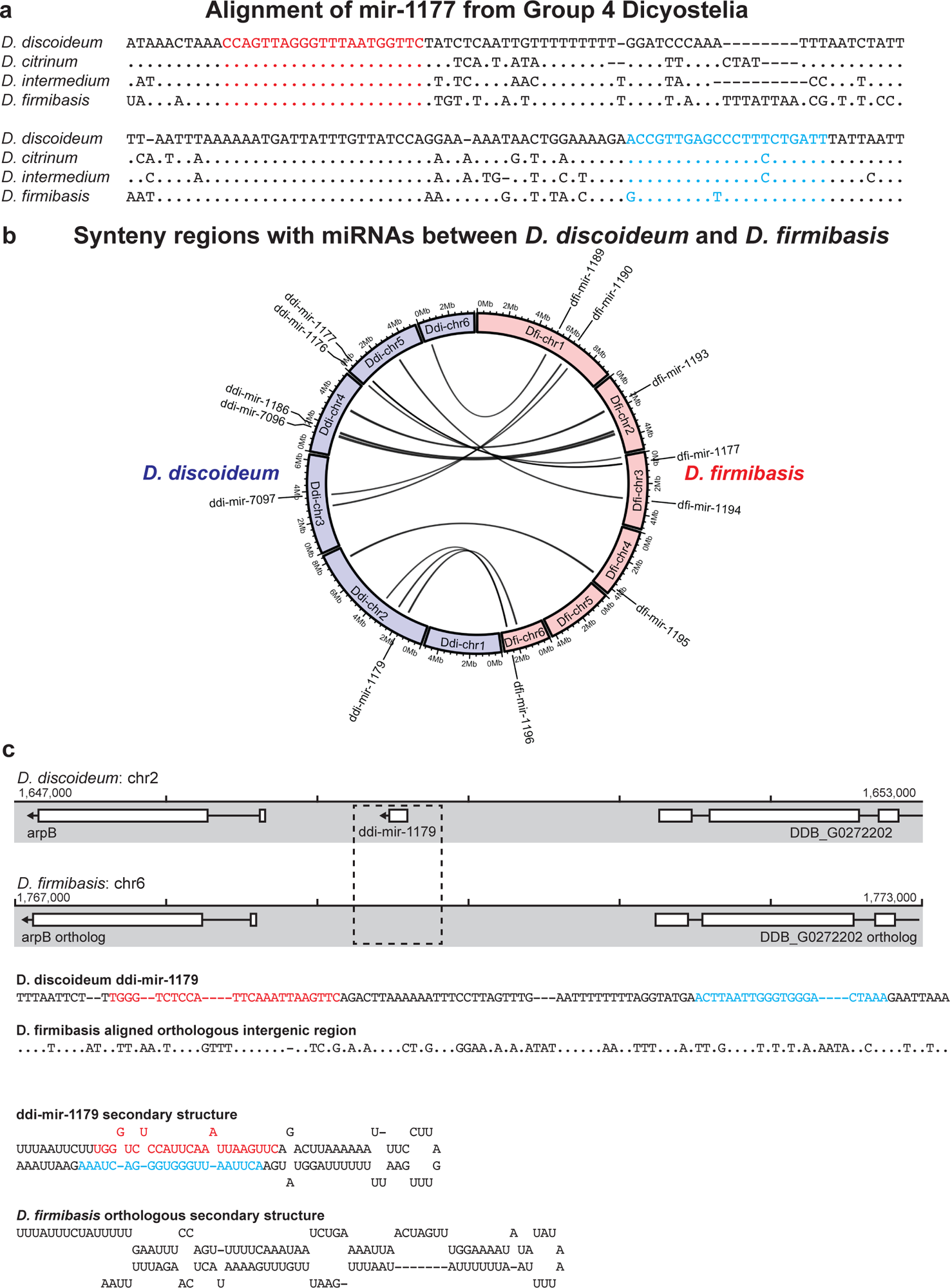
miRNA conservation in Group 4 Dictyostelia and miRNA evolution between *D. discoideum* and *D. firmibasis*. **a** Alignment of the mir-1177 orthologs found in *D. discoideum*, *D. citrinum*, *D. intermedium*, and *D. firmibasis*. The DNA sequences from *D. citrinum* and *D. intermedium* were resolved by sequencing the genomic region in which the ortholog is located. The *D. firmibasis* ortholog was resolved by whole genome sequencing. Positions of the mir-1177-5p and mir-1177-3p are indicated in red and cyan, respectively. **b** Circos plot of the *D. firmibasis* (pink) and *D. discoideum* (purple) main contigs, with links showing syntenic regions containing a miRNA in either *D. firmibasis* or *D. discoideum*. (Note that ddi-mir-1177 is the only miRNA identified to be conserved between the two dictyostelids). The name of the miRNAs present in the syntenic regions are shown on the outside of the plot. **c** Intergenic region containing ddi-mir-1179 in *D. discoideum*, and below the aligned syntenic region in *D. firmibasis*. The DNA sequence in the dashed box was further aligned and the hypothetical ncRNA folded in silico, which yielded the displayed hairpin-like structure. The positions of ddi-mir-1179-5p and ddi-mir-1179-3p are indicated in red and cyan, respectively, in both the DNA alignment and RNA secondary structure.

**Extended Data Fig. 7.**
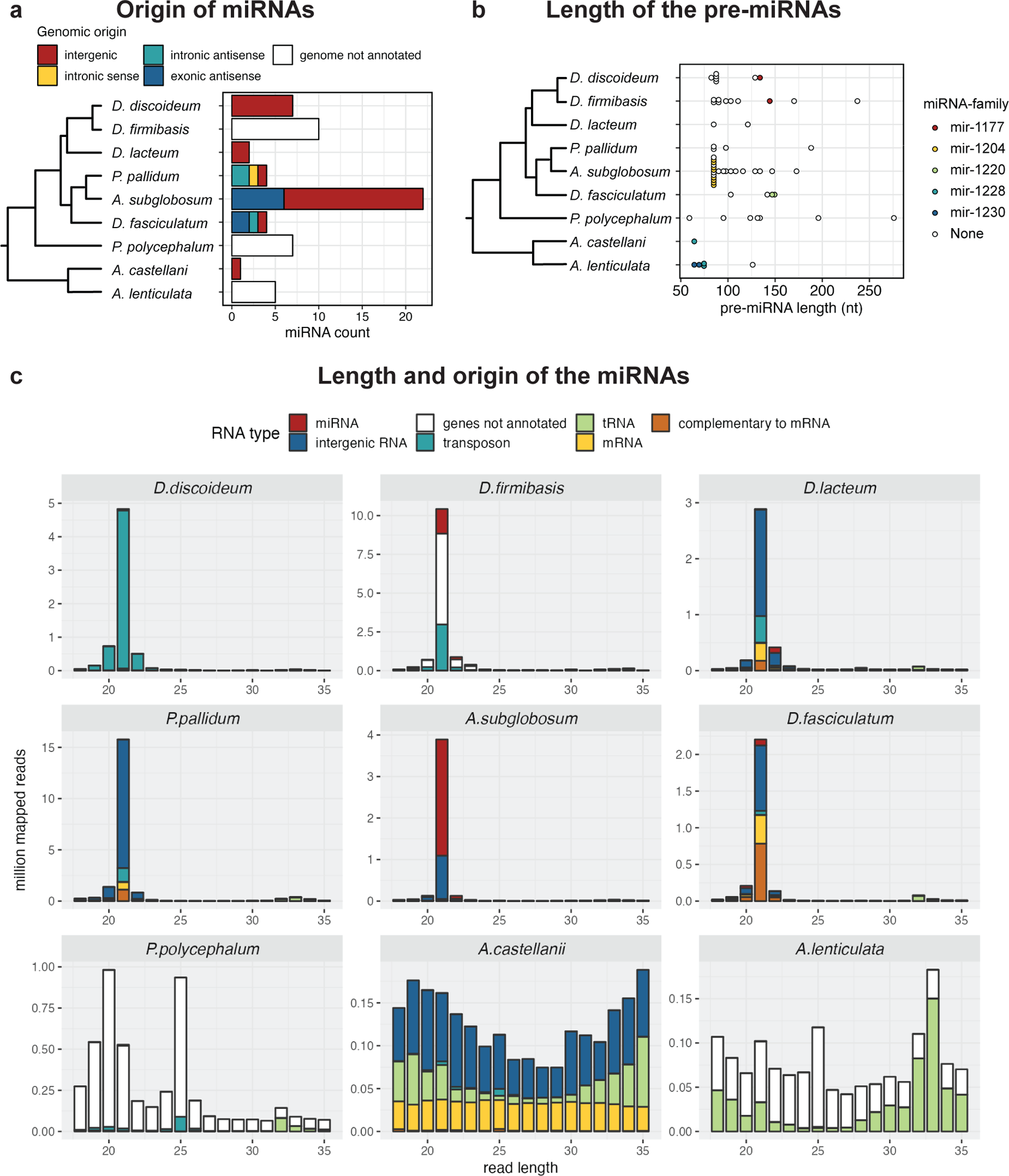
Genomic origin and size of the miRNAs and sRNAs in Amoebozoa. **a** Barchart displaying the number of all discovered miRNAs in this study and their origin in the respective amoebae, with a simplified phylogenetic dendrogram. Genome annotations are not available for *D. firmibasis*, *P. polycephalum* and *A. lenticulata.* **b** miRNAs plotted based on their pre-miRNA length. miRNAs of the same family are plotted with a single color. **c** Length distribution of the small RNAs sequenced from each of the studied amoebae, as in Extended Data Fig. 2b, but with annotations, showing where the sRNAs map to the genome. Annotation of transposons is based on blast search with transposable elements available from Repbase. Annotations of tRNAs and other known non-coding RNAs was performed using Infernal with models based on the Rfam database. Number of sRNAs mapping to spliceosomal RNAs and other ncRNAs were omitted since they were very low, but they can be accessed through GitHub.

**Extended Data Fig. 8.**
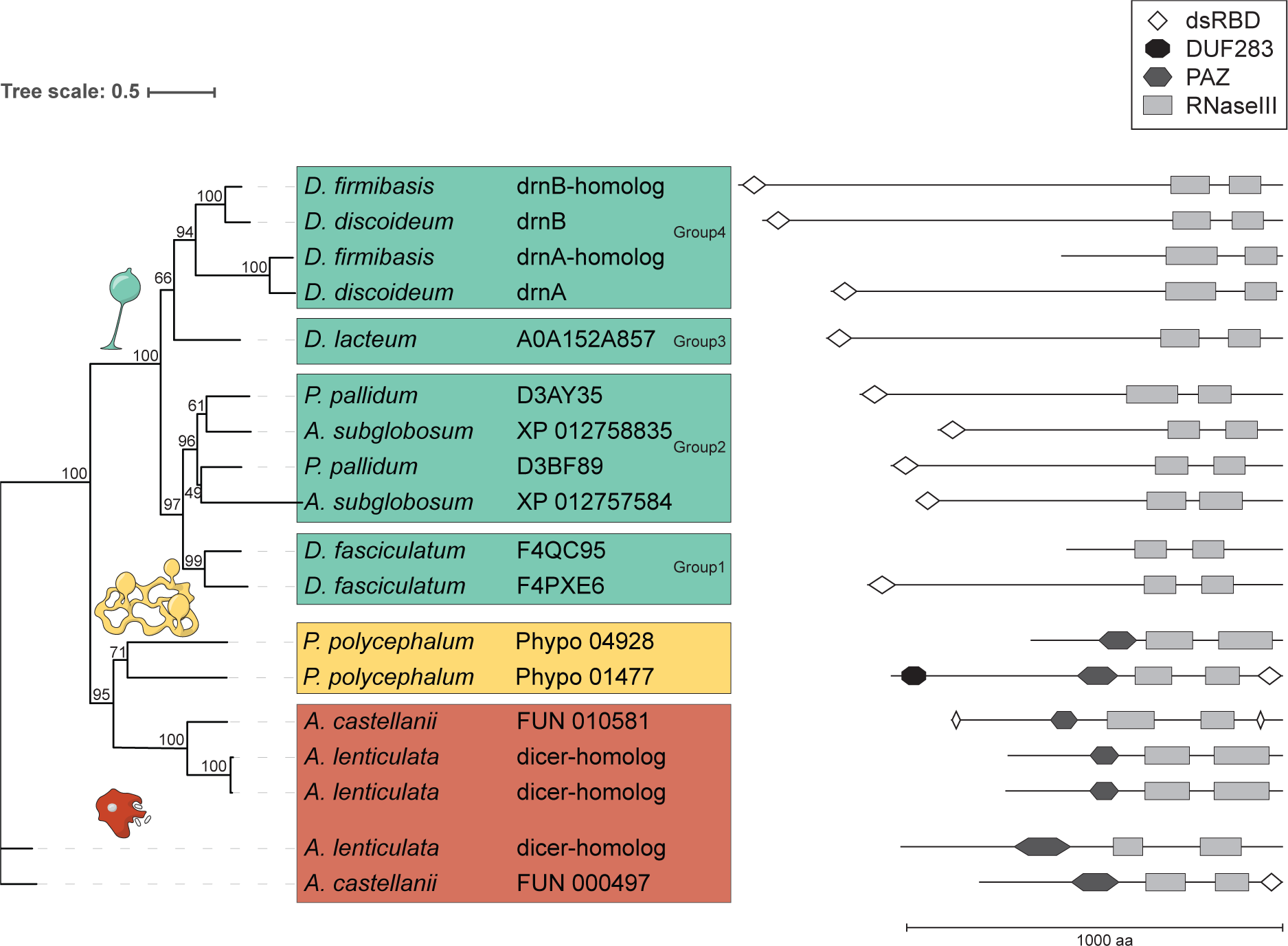
Phylogeny of the Dicer proteins in Amoebozoa. Dicer homologs identified through search with OrthoFinder^61^ on the proteomes of all included species, except *D. firmibasis* and *A. lenticulata* for which no proteomes are available. Homologs for *D. firmibasis* and *A. lenticulata* were annotated by an additional BLAST search. Numbers after species names are the protein identifiers. For *D. discoideum*, *D. lacteum*, *P. pallidum*, *A. subglobosum* and *D. fasciculatum*, proteins can be accessed at UniProt. The domain structure of the homologs is shown (right). All harbor two Ribonuclease III domains (RNase III) and additionally some contain PAZ and/or double stranded RNA-binding domains (dsRBD). The amino acid sequences were aligned, trimmed and used to build a phylogenetic tree. The consensus tree is shown with bootstrap supports (%) on the branches. The scale shows evolutionary distance, defined as the number of nucleotide substitutions per site. Amoebae (and their Dicer homologs) belonging to Dictyostelia, Myxomycetes, and Discosea are marked with cyan, yellow, and orange back ground, respectively.

**Extended Data Fig. 9.**
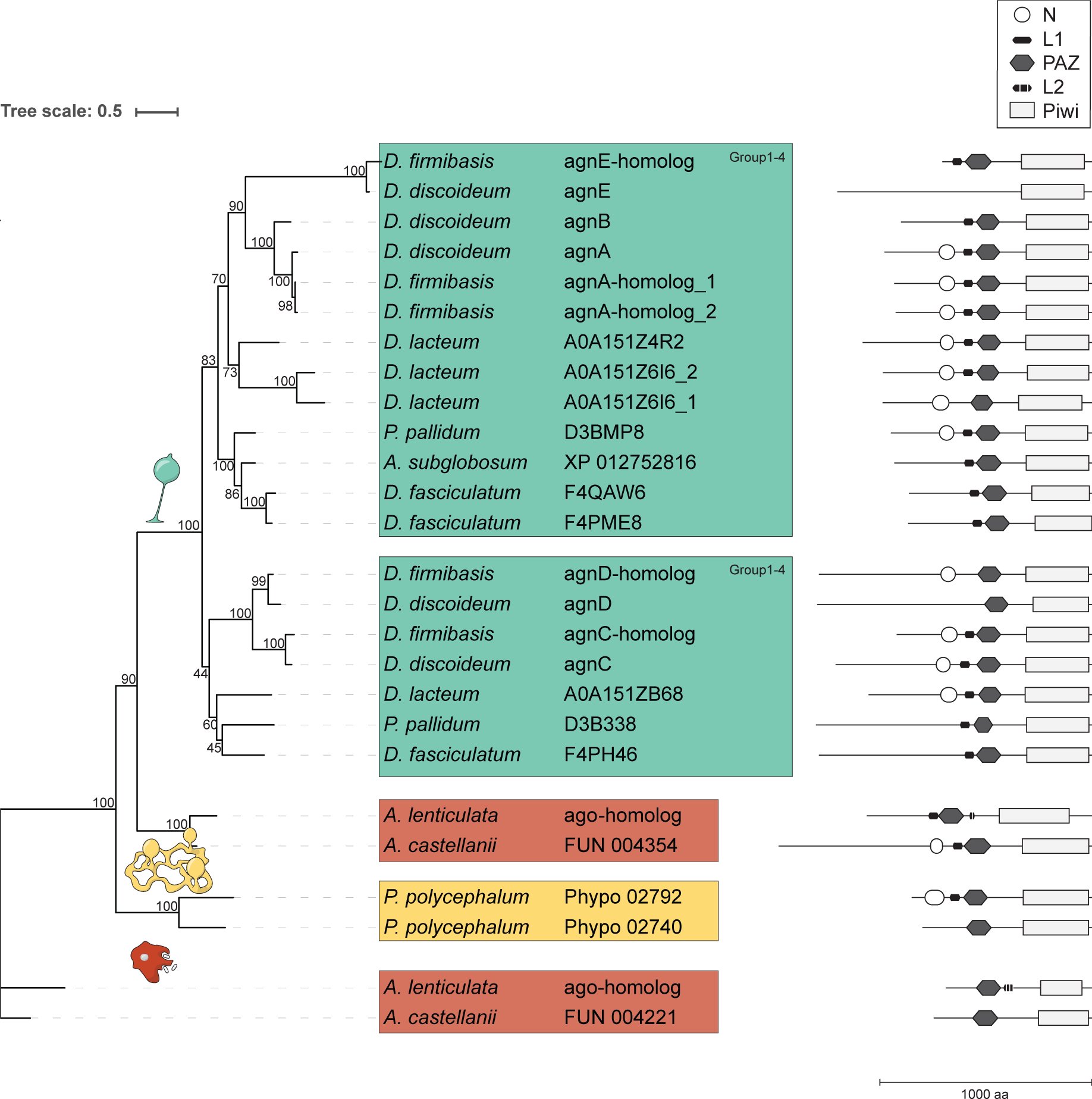
Phylogeny of the Argonaute proteins in Amoebozoa. Argonaute homologs identified through search with OrthoFinder^61^ on the proteomes of all included species, except *D. firmibasis* and *A. lenticulata* for which no proteomes are available. Homologs for *D. firmibasis* and *A. lenticulata* were annotated by an additional BLAST search. Numbers after species names are the protein identifiers. For *D. discoideum*, *D. lacteum*, *P. pallidum*, *A. subglobosum* and *D. fasciculatum*, proteins can be accessed at UniProt. The domain structure of the homologs is shown (right). All identified Argonautes feature a single PAZ domain, followed by a single Piwi domain. Identified Argonaute N-terminal domains (N) as well as linker domains (L1, L2) are shown as well. The amino acid sequences of the identified Argonautes were aligned, trimmed and used to build a phylogenetic tree. The consensus tree is shown with bootstrap supports (%) on the branches. The scale shows evolutionary distance, defined as the number of nucleotide substitutions per site. Dictyostelia feature two main Argonaute orthologs which are present in all analyzed species, and which were probably both present in their last common ancestor. Amoebae (and their Argonaute homologs) belonging to Dictyostelia, Myxomycetes, and Discosea are marked with cyan, yellow, and orange back ground, respectively.

**Extended Data Fig. 10.**
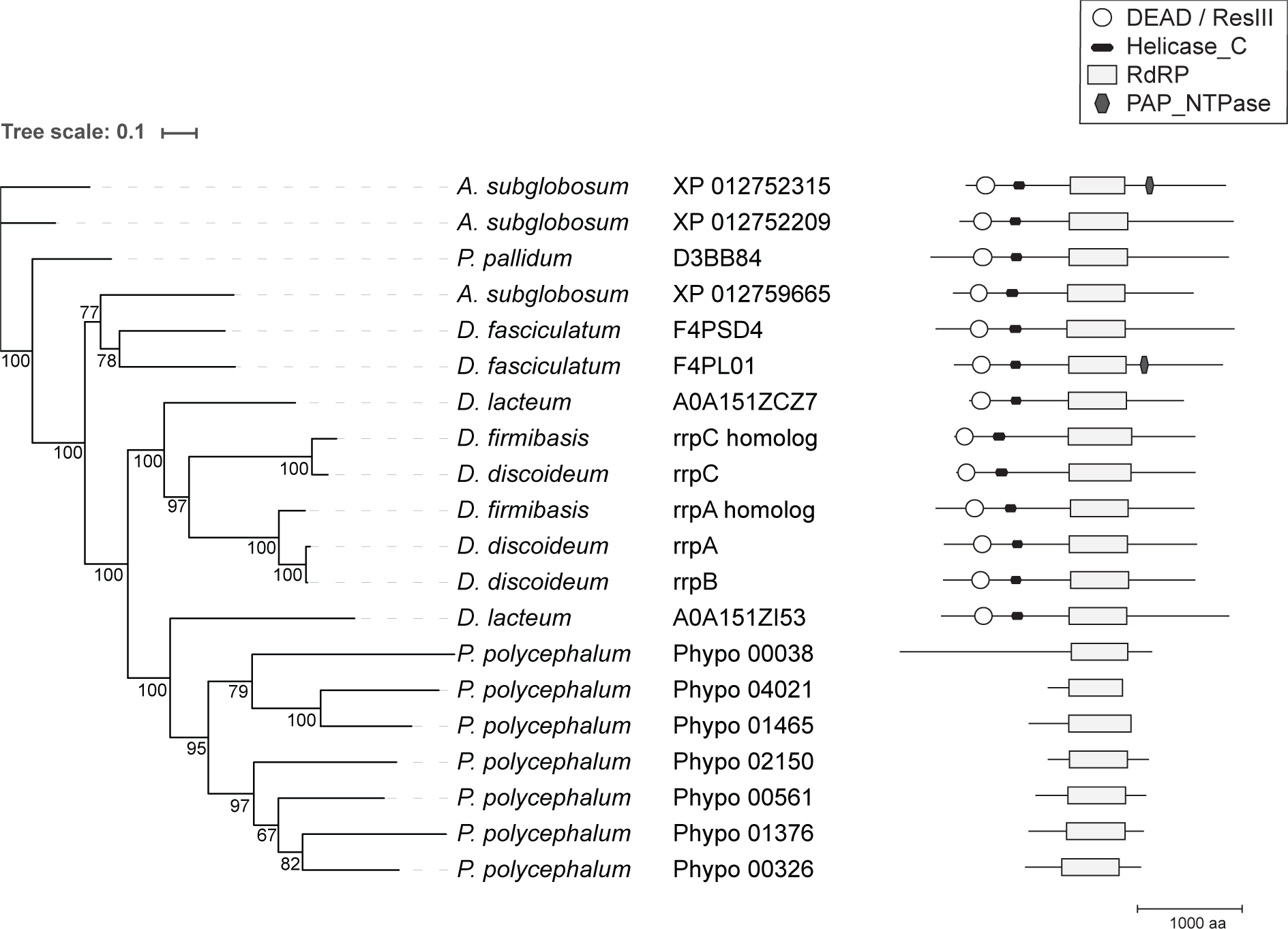
Phylogeny of the RNA-dependent RNA polymerases in Amoebozoa. RNA-dependent RNA polymerase (RdRP) homologs identified through search with OrthoFinder^61^ on the proteomes of all included species, except *D. firmibasis* and *A. lenticulata* for which no proteomes are available. Homologs for *D. firmibasis* were annotated by an additional BLAST search. No homologs could be identified for *A. castelllanii* and *A. lenticulata*. Numbers after species names are the protein identifiers. For *D. discoideum*, *D. lacteum*, *P. pallidum*, *A. subglobosum* and *D. fasciculatum*, proteins can be accessed at UniProt. The domain structure of the homologs is shown (right). All identified RdRPs feature a RdRP domain. Identified ResIII/DEAD box helicase domains (ResIII/DEAD) and Helicase c-terminal domains (Helicase_C) are shown as well. Additionallly, two RdRPs contain a poly(A) polymerase nucleotidyltransferase domain (PAP_NTPase). The amino acid sequences of the identified RdRPs were aligned, trimmed and used to build a phylogenetic tree. The consensus tree is shown with bootstrap supports (%) on the branches. The scale shows evolutionary distance, defined as the number of nucleotide substitutions per site.

