## Supplementary Data 1 for "Evolution of microRNAs in Amoebozoa and implications for the origin of multicellularity"

#### *D. discoideum* ddi-mir-1176

```
((((((((((.((((((((.((((((((.((((.(((((((((.(.(.......).).))))))))).)))).)))))))).)))))))).))))))))))  -70.6 kcal/mol
UAUCAGGUGGcCAAUUUUUAUCAAGGAAAGcUGUAUCAUCAAGUGCCUCCCUCUGUCUCUUGAUGAUUCAGCcUUCCUUGACAAAAAUUGCCcACCUGAUA  count    length
Libraries combined          Precision: [ total = 98.5% | 5p-arm = 98.4% | 3p-arm = 100.0% ]
----------CCAAUUUUUAUCAAGGAAAGC----------------------------------------------------------------------  694      21
------------------------------------------------------------------------CUUCCUUGACAAAAAUUGCCC--------  29       21
------------------------------------------------------------------------CUUCCUUGACAAAAAUUGCC---------  15       20
----------CCAAUUUUUAUCAAGGAAAG-----------------------------------------------------------------------  24       20
---------GCCAAUUUUUAUCAAGGAAAG-----------------------------------------------------------------------  3        21
-----------CAAUUUUUAUCAAGGAAAGC----------------------------------------------------------------------  5        20
-------UGGCCAAUUUUUAUCAAGGAAAG-----------------------------------------------------------------------  3        23
---------GCCAAUUUUUAUCAAGGAAAGC----------------------------------------------------------------------  1        22
----------CCAAUUUUUAUCAAGGAAAGCU---------------------------------------------------------------------  1        22
Library = 1        Precision: [ total = 99.0% | 5p-arm = 98.9% | 3p-arm = 100.0% ]
----------CCAAUUUUUAUCAAGGAAAGC----------------------------------------------------------------------  171      21
------------------------------------------------------------------------CUUCCUUGACAAAAAUUGCCC--------  15       21
------------------------------------------------------------------------CUUCCUUGACAAAAAUUGCC---------  6        20
----------CCAAUUUUUAUCAAGGAAAG-----------------------------------------------------------------------  5        20
---------GCCAAUUUUUAUCAAGGAAAG-----------------------------------------------------------------------  1        21
-----------CAAUUUUUAUCAAGGAAAGC----------------------------------------------------------------------  1        20
Library = 2        Precision: [ total = 98.8% | 5p-arm = 98.7% | 3p-arm = 100.0% ]
----------CCAAUUUUUAUCAAGGAAAGC----------------------------------------------------------------------  71       21
----------CCAAUUUUUAUCAAGGAAAG-----------------------------------------------------------------------  4        20
------------------------------------------------------------------------CUUCCUUGACAAAAAUUGCCC--------  3        21
------------------------------------------------------------------------CUUCCUUGACAAAAAUUGCC---------  2        20
---------GCCAAUUUUUAUCAAGGAAAG-----------------------------------------------------------------------  1        21
Library = 3        Precision: [ total = 98.2% | 5p-arm = 98.1% | 3p-arm = 100.0% ]
----------CCAAUUUUUAUCAAGGAAAGC----------------------------------------------------------------------  452      21
----------CCAAUUUUUAUCAAGGAAAG-----------------------------------------------------------------------  15       20
------------------------------------------------------------------------CUUCCUUGACAAAAAUUGCCC--------  11       21
------------------------------------------------------------------------CUUCCUUGACAAAAAUUGCC---------  7        20
-----------CAAUUUUUAUCAAGGAAAGC----------------------------------------------------------------------  4        20
-------UGGCCAAUUUUUAUCAAGGAAAG-----------------------------------------------------------------------  3        23
---------GCCAAUUUUUAUCAAGGAAAG-----------------------------------------------------------------------  1        21
---------GCCAAUUUUUAUCAAGGAAAGC----------------------------------------------------------------------  1        22
----------CCAAUUUUUAUCAAGGAAAGCU---------------------------------------------------------------------  1        22
```

```
          c        A        A    U         U C UCC
UAUCAGGUGG CAAUUUUU UCAAGGAA GcUG AUCAUCAAG G C   \
AUAGUCCAcC GUUAAAAA AGUUCCUU CGAC UAGUAGUUC C G   C
          C        C        c    U         U U UCU
```

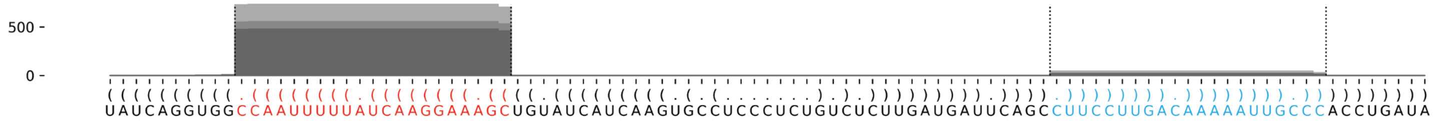

#### *D. discoideum* ddi-mir-1177

```
......((((.(((..((((((((((((((((.((.((.(((((((((((.((((..((((..(((((.(((((.......))))))))))))))..)))))))))))))))).))))..))))))))))))))))..))).))))......   -65.48 kcal/mol
AUAAACUAAAcCAGUUAGGGUUUAAUGGUUcUAUCUCAAUUGUUUUUUUUUGGAUCCCAAAUUUAAUCUAUUUUAAUUUAAAAAAUGAUUAUUUGUUAUCCAGGAAAAAUAACUGGAAAAGAaCCGUUGAGCCCUUUCUGAUuUAUUAAUU   count    length
Libraries combined          Precision: [ total = 98.7% | 5p-arm = 98.6% | 3p-arm = 99.2% ]
----------CCAGUUAGGGUUUAAUGGUUC------------------------------------------------------------------------------------------------------------------------   612      21
--------------------------------------------------------------------------------------------------------------------------ACCGUUGAGCCCUUUCUG-----------   28       18
----------CCAGUUAGGGUUUAAUGGUUCU-----------------------------------------------------------------------------------------------------------------------   30       22
--------------------------------------------------------------------------------------------------------------------------ACCGUUGAGCCCUUUCUGAU---------   70       20
----------CCAGUUAGGGUUUAAUGGUU-------------------------------------------------------------------------------------------------------------------------   5        20
-----------CAGUUAGGGUUUAAUGGUUC------------------------------------------------------------------------------------------------------------------------   8        20
--------------------------------------------------------------------------------------------------------------------------ACCGUUGAGCCCUUUCUGAUU--------   23       21
----------CCAGUUAGGGUUUAAUGGU--------------------------------------------------------------------------------------------------------------------------   1        19
----------CCAGUUAGGGUUUAAUGGUUCUA----------------------------------------------------------------------------------------------------------------------   1        23
------------AGUUAGGGUUUAAUGGUUCU-----------------------------------------------------------------------------------------------------------------------   1        20
-----------------------------------------------------------------------------------------------------AGGAAAAAUAACUGGAAAAGA-----------------------------   1        21
--------------------------------------------------------------------------------------------------------------------------ACCGUUGAGCCCUUUCUGA----------   6        19
--------------------------------------------------------------------------------------------------------------------------ACCGUUGAGCCCUUUCUGAUUU-------   1        22
Library = 1        Precision: [ total = 98.8% | 5p-arm = 98.8% | 3p-arm = 100.0% ]
----------CCAGUUAGGGUUUAAUGGUUC------------------------------------------------------------------------------------------------------------------------   152      21
--------------------------------------------------------------------------------------------------------------------------ACCGUUGAGCCCUUUCUG-----------   6        18
----------CCAGUUAGGGUUUAAUGGUUCU-----------------------------------------------------------------------------------------------------------------------   4        22
--------------------------------------------------------------------------------------------------------------------------ACCGUUGAGCCCUUUCUGAU---------   4        20
----------CCAGUUAGGGUUUAAUGGUU-------------------------------------------------------------------------------------------------------------------------   2        20
-----------CAGUUAGGGUUUAAUGGUUC------------------------------------------------------------------------------------------------------------------------   2        20
--------------------------------------------------------------------------------------------------------------------------ACCGUUGAGCCCUUUCUGAUU--------   2        21
Library = 2        Precision: [ total = 97.7% | 5p-arm = 97.5% | 3p-arm = 98.2% ]
----------CCAGUUAGGGUUUAAUGGUUC------------------------------------------------------------------------------------------------------------------------   150      21
--------------------------------------------------------------------------------------------------------------------------ACCGUUGAGCCCUUUCUGAU---------   30       20
--------------------------------------------------------------------------------------------------------------------------ACCGUUGAGCCCUUUCUGAUU--------   12       21
--------------------------------------------------------------------------------------------------------------------------ACCGUUGAGCCCUUUCUG-----------   9        18
----------CCAGUUAGGGUUUAAUGGUUCU-----------------------------------------------------------------------------------------------------------------------   4        22
--------------------------------------------------------------------------------------------------------------------------ACCGUUGAGCCCUUUCUGA----------   4        19
-----------CAGUUAGGGUUUAAUGGUUC------------------------------------------------------------------------------------------------------------------------   3        20
----------CCAGUUAGGGUUUAAUGGU--------------------------------------------------------------------------------------------------------------------------   1        19
------------AGUUAGGGUUUAAUGGUUCU-----------------------------------------------------------------------------------------------------------------------   1        20
-----------------------------------------------------------------------------------------------------AGGAAAAAUAACUGGAAAAGA-----------------------------   1        21
--------------------------------------------------------------------------------------------------------------------------ACCGUUGAGCCCUUUCUGAUUU-------   1        22
Library = 3        Precision: [ total = 99.2% | 5p-arm = 99.1% | 3p-arm = 100.0% ]
----------CCAGUUAGGGUUUAAUGGUUC------------------------------------------------------------------------------------------------------------------------   310      21
--------------------------------------------------------------------------------------------------------------------------ACCGUUGAGCCCUUUCUGAU---------   36       20
----------CCAGUUAGGGUUUAAUGGUUCU-----------------------------------------------------------------------------------------------------------------------   22       22
--------------------------------------------------------------------------------------------------------------------------ACCGUUGAGCCCUUUCUG-----------   13       18
--------------------------------------------------------------------------------------------------------------------------ACCGUUGAGCCCUUUCUGAUU--------   9        21
----------CCAGUUAGGGUUUAAUGGUU-------------------------------------------------------------------------------------------------------------------------   3        20
-----------CAGUUAGGGUUUAAUGGUUC------------------------------------------------------------------------------------------------------------------------   3        20
--------------------------------------------------------------------------------------------------------------------------ACCGUUGAGCCCUUUCUGA----------   2        19
----------CCAGUUAGGGUUUAAUGGUUCUA----------------------------------------------------------------------------------------------------------------------   1        23
```

```
AUAAAC    c   UU                A   U  A           U    CC    UU     U     AAU
      UAAA CAG  AGGGUUUAAUGGUUcU -UC CA UUGUUUUUUUU GGAU  CAAA  UAAUC AUUUU   \
      AUuU GUC  UCCCGAGUUGCCaAGA  AG-GU AAUAAAAAGGA-CCUA  GUUU--AUUAG-UAAAA   U
UUAAUU    A   UU                AA     C                UU                 AAU
```

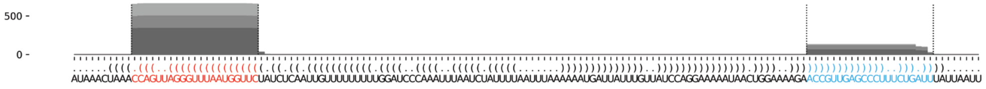

#### *D. discoideum* ddi-mir-1179

```
((((((((((((.((.((((((((.((((((((.((((((((((.(((........)))..)))))))))).)))))))))))))))))))))))))))))))))      -56.04 kcal/mol
UUUAAUUCUUuGGGUCUCCAUUCAAAUUAAGUUcAGACUUAAAAAAUUUCCUUAGUUUGAAUUUUUUUUAGGUAUGAaCUUAAUUGGGUGGGACUAAaGAAUUAAA      count    length
Libraries combined          Precision: [ total = 98.8% | 5p-arm = 100.0% | 3p-arm = 92.3% ]
----------UGGGUCUCCAUUCAAAUUAAGUUC------------------------------------------------------------------------      65       24
----------UGGGUCUCCAUUCAAAUUAAGUU-------------------------------------------------------------------------      6        23
-----------------------------------------------------------------------------ACUUAAUUGGGUGGGACUAAA--------      8        21
------------------------------------------------------------------------UAUGAACUUAAUUGGGUGGGACU-----------      1        23
-----------------------------------------------------------------------------ACUUAAUUGGGUGGGACUAA---------      4        20
Library = 1         Precision: [ total = 100.0% | 5p-arm = 100.0% | 3p-arm = 100.0% ]
----------UGGGUCUCCAUUCAAAUUAAGUUC------------------------------------------------------------------------      14       24
----------UGGGUCUCCAUUCAAAUUAAGUU-------------------------------------------------------------------------      2        23
-----------------------------------------------------------------------------ACUUAAUUGGGUGGGACUAAA--------      2        21
Library = 2         Precision: [ total = 96.9% | 5p-arm = 100.0% | 3p-arm = 83.3% ]
----------UGGGUCUCCAUUCAAAUUAAGUUC------------------------------------------------------------------------      24       24
-----------------------------------------------------------------------------ACUUAAUUGGGUGGGACUAAA--------      3        21
----------UGGGUCUCCAUUCAAAUUAAGUU-------------------------------------------------------------------------      2        23
-----------------------------------------------------------------------------ACUUAAUUGGGUGGGACUAA---------      2        20
------------------------------------------------------------------------UAUGAACUUAAUUGGGUGGGACU-----------      1        23
Library = 3         Precision: [ total = 100.0% | 5p-arm = 100.0% | 3p-arm = 100.0% ]
----------UGGGUCUCCAUUCAAAUUAAGUUC------------------------------------------------------------------------      27       24
-----------------------------------------------------------------------------ACUUAAUUGGGUGGGACUAAA--------      3        21
----------UGGGUCUCCAUUCAAAUUAAGUU-------------------------------------------------------------------------      2        23
-----------------------------------------------------------------------------ACUUAAUUGGGUGGGACUAA---------      2        20
```

```
             G  U        A         G          U    CUU
UUUAAUUCUUuGG UC CCAUUCAA UUAAGUUcA ACUUAAAAAA -UUC   A
AAAUUAAGaAAUC-AG-GGUGGGUUA-AUUCaAGU UGGAUUUUUU  AAG   G
                                   A          UU   UUU
```

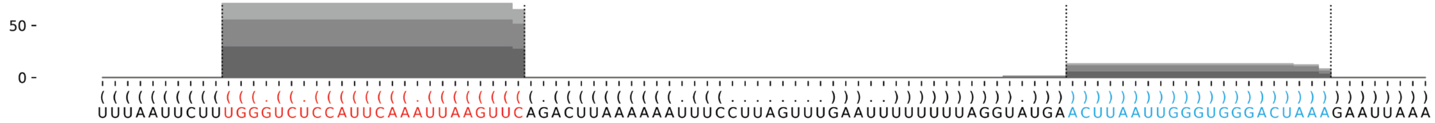

##### *D. discoideum* ddi-mir-7096

```
((((((((.(((((((((((((.((((((((((((((((((((((((..........)))).)))))))))))))))))))).))))))))))))).))))))))        -72.22 kcal/mol
UUAAACUGAGuCUGUGAACAAUCAAGAUUUgAUAAACACUAUUGAAAUUGAAAUUUUUUUCUAAUGGUGUUUAUCAaAUCUUAAUUGUUCACAGACcCAGUUUAA        count    length
Libraries combined         Precision: [ total = 99.5% | 5p-arm = 100.0% | 3p-arm = 99.5% ]
----------------------------------------------------------------------------AAUCUUAAUUGUUCACAGACC--------        386      21
----------------------------------------------------------------------------AAUCUUAAUUGUUCACAGACCC-------        15       22
----------UCUGUGAACAAUCAAGAUUUG--------------------------------------------------------------------------        7        21
----------------------------------------------------------------------------AAUCUUAAUUGUUCACAGACCCAG-----        1        24
----------------------------------------------------------------------------AAUCUUAAUUGUUCACAGAC---------        3        20
-------------------------------------------------------------------------------CUUAAUUGUUCACAGACC--------        1        18
Library = 1        Precision: [ total = 100.0% | 5p-arm = 100.0% | 3p-arm = 100.0% ]
----------------------------------------------------------------------------AAUCUUAAUUGUUCACAGACC--------        66       21
----------------------------------------------------------------------------AAUCUUAAUUGUUCACAGACCC-------        3        22
----------UCUGUGAACAAUCAAGAUUUG--------------------------------------------------------------------------        2        21
Library = 2        Precision: [ total = 100.0% | 5p-arm = 100.0% | 3p-arm = 100.0% ]
----------------------------------------------------------------------------AAUCUUAAUUGUUCACAGACC--------        72       21
----------------------------------------------------------------------------AAUCUUAAUUGUUCACAGACCC-------        5        22
----------UCUGUGAACAAUCAAGAUUUG--------------------------------------------------------------------------        2        21
Library = 3        Precision: [ total = 99.2% | 5p-arm = 100.0% | 3p-arm = 99.2% ]
----------------------------------------------------------------------------AAUCUUAAUUGUUCACAGACC--------        248      21
----------------------------------------------------------------------------AAUCUUAAUUGUUCACAGACCC-------        7        22
----------UCUGUGAACAAUCAAGAUUUG--------------------------------------------------------------------------        3        21
----------------------------------------------------------------------------AAUCUUAAUUGUUCACAGAC---------        3        20
----------------------------------------------------------------------------AAUCUUAAUUGUUCACAGACCCAG-----        1        24
-------------------------------------------------------------------------------CUUAAUUGUUCACAGACC--------        1        18
```

```
        A             C                               UUGA
UUAAACUG GuCUGUGAACAAU AAGAUUUgAUAAACACUAUU-GAAA    A
AAUUUGAC CAGACACUUGUUA UUCUAaACUAUUUGUGGUAA CUUU    A
        c             A                    U    UUUU
```

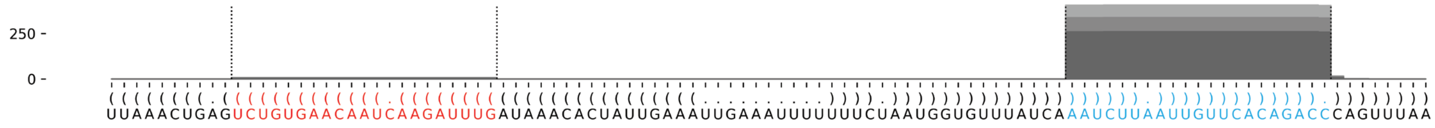

#### *D. discoideum* ddi-mir-1183

```
((((.((((((((((((((((((((((.((((((((.....((((((((.(((((.((.(((((.(((....))).))))).))((....))))))).)))))))))).....))))))).)))))))))))))))))))))).))))        -88.47 kcal/mol
UUAUACUAUAuAAGUGGUCGUUAAUUGCUGGcAGACCUCAGUUUACUCACACUAGACCACUGGCCGUCUUCUGGCUGCUAGCGGGCAAUGGCCUAGUAUGAGUAAAUGACCAUCUGCcACCAGUUAGCGGUCAUUUGUaUAGCAUGG        count    length
Libraries combined         Precision: [ total = 88.1% | 5p-arm = 88.1% | 3p-arm = 87.2% ]
----------UAAGUGGUCGUUAAUUGCUGGC-------------------------------------------------------------------------------------------------------------------        1471     22
-----------AAGUGGUCGUUAAUUGCUGGC-------------------------------------------------------------------------------------------------------------------        184      21
---------------------------------------------------------------------------------------------------------------------CACCAGUUAGCGGUCAUUUGUA--------        27       22
----------UAAGUGGUCGUUAAUUGC-----------------------------------------------------------------------------------------------------------------------        16       18
----------UAAGUGGUCGUUAAUUGCUGG--------------------------------------------------------------------------------------------------------------------        7        21
----------UAAGUGGUCGUUAAUUGCUG---------------------------------------------------------------------------------------------------------------------        7        20
--------------------------------AGACCUCAGUUUACUCACACU----------------------------------------------------------------------------------------------        5        21
--------UAUAAGUGGUCGUUAAUUGCU----------------------------------------------------------------------------------------------------------------------        3        21
----------UAAGUGGUCGUUAAUUGCU----------------------------------------------------------------------------------------------------------------------        8        19
------------AGUGGUCGUUAAUUGCUGGC-------------------------------------------------------------------------------------------------------------------        6        20
------------AGUGGUCGUUAAUUGCUGGCA------------------------------------------------------------------------------------------------------------------        2        21
----------------------------------------------------------------CGUCUUCUGGCUGCUAGC-----------------------------------------------------------------        1        18
----------------------------------------------------------------------UCUGGCUGCUAGCGGGCAAU---------------------------------------------------------        1        20
---------------------------------------------------------------------------------------------------------------------CACCAGUUAGCGGUCAUUUGU---------        5        21
---------------------------------------------------------------------------------------------------------------------CACCAGUUAGCGGUCAUU------------        2        18
--------------UGGUCGUUAAUUGCUGGC-------------------------------------------------------------------------------------------------------------------        1        18
-------------------------------CAGACCUCAGUUUACUCACACU----------------------------------------------------------------------------------------------        1        22
------------------------------------------------------------------------------------------------UAUGAGUAAAUGACCAUCUGC------------------------------        2        21
-------------------------------------------------------------------------------------------------AUGAGUAAAUGACCAUCUGCCACC--------------------------        1        24
---------------------------------------------------------------------------------------------------------------------------UUAGCGGUCAUUUGUAUAGC----        1        20
Library = 1        Precision: [ total = 86.6% | 5p-arm = 86.4% | 3p-arm = 93.8% ]
----------UAAGUGGUCGUUAAUUGCUGGC-------------------------------------------------------------------------------------------------------------------        497      22
-----------AAGUGGUCGUUAAUUGCUGGC-------------------------------------------------------------------------------------------------------------------        71       21
---------------------------------------------------------------------------------------------------------------------CACCAGUUAGCGGUCAUUUGUA--------        13       22
----------UAAGUGGUCGUUAAUUGC-----------------------------------------------------------------------------------------------------------------------        6        18
----------UAAGUGGUCGUUAAUUGCUGG--------------------------------------------------------------------------------------------------------------------        4        21
----------UAAGUGGUCGUUAAUUGCUG---------------------------------------------------------------------------------------------------------------------        4        20
--------------------------------AGACCUCAGUUUACUCACACU----------------------------------------------------------------------------------------------        4        21
--------UAUAAGUGGUCGUUAAUUGCU----------------------------------------------------------------------------------------------------------------------        2        21
----------UAAGUGGUCGUUAAUUGCU----------------------------------------------------------------------------------------------------------------------        2        19
------------AGUGGUCGUUAAUUGCUGGC-------------------------------------------------------------------------------------------------------------------        2        20
------------AGUGGUCGUUAAUUGCUGGCA------------------------------------------------------------------------------------------------------------------        1        21
----------------------------------------------------------------CGUCUUCUGGCUGCUAGC-----------------------------------------------------------------        1        18
----------------------------------------------------------------------UCUGGCUGCUAGCGGGCAAU---------------------------------------------------------        1        20
---------------------------------------------------------------------------------------------------------------------CACCAGUUAGCGGUCAUUUGU---------        1        21
---------------------------------------------------------------------------------------------------------------------CACCAGUUAGCGGUCAUU------------        1        18
Library = 2        Precision: [ total = 84.2% | 5p-arm = 84.7% | 3p-arm = 50.0% ]
----------UAAGUGGUCGUUAAUUGCUGGC-------------------------------------------------------------------------------------------------------------------        108      22
-----------AAGUGGUCGUUAAUUGCUGGC-------------------------------------------------------------------------------------------------------------------        18       21
----------UAAGUGGUCGUUAAUUGC-----------------------------------------------------------------------------------------------------------------------        2        18
----------UAAGUGGUCGUUAAUUGCU----------------------------------------------------------------------------------------------------------------------        1        19
------------AGUGGUCGUUAAUUGCUGGC-------------------------------------------------------------------------------------------------------------------        1        20
-------------------------------CAGACCUCAGUUUACUCACACU----------------------------------------------------------------------------------------------        1        22
---------------------------------------------------------------------------------------------------------------------CACCAGUUAGCGGUCAUUUGUA--------        1        22
---------------------------------------------------------------------------------------------------------------------------UUAGCGGUCAUUUGUAUAGC----        1        20
Library = 3        Precision: [ total = 89.6% | 5p-arm = 89.7% | 3p-arm = 85.7% ]
----------UAAGUGGUCGUUAAUUGCUGGC-------------------------------------------------------------------------------------------------------------------        866      22
-----------AAGUGGUCGUUAAUUGCUGGC-------------------------------------------------------------------------------------------------------------------        95       21
---------------------------------------------------------------------------------------------------------------------CACCAGUUAGCGGUCAUUUGUA--------        13       22
----------UAAGUGGUCGUUAAUUGC-----------------------------------------------------------------------------------------------------------------------        8        18
----------UAAGUGGUCGUUAAUUGCU----------------------------------------------------------------------------------------------------------------------        5        19
---------------------------------------------------------------------------------------------------------------------CACCAGUUAGCGGUCAUUUGU---------        4        21
----------UAAGUGGUCGUUAAUUGCUGG--------------------------------------------------------------------------------------------------------------------        3        21
```

#### *D. discoideum* ddi-mir-1183-continued

```
((((.((((((((((((((((((((((.(((((((.....(((((((((.(((((.((.(((((.(((....))).))))).))((....))))))).))))))))).....))))))).))))))))))))))))))))))).))))   -88.47 kcal/mol
UUAUACUAUAuAAGUGGUCGUUAAUUGCUGGcAGACCUCAGUUUACUCACACUAGACCACUGGCCGUCUUCUGGCUGCUAGCGGGCAAUGGCCUAGUAUGAGUAAAUGACCAUCUGCcACCAGUUAGCGGUCAUUUGUaUAGCAUGG   count   length
----------UAAGUGGUCGUUAAUUGCUG---------------------------------------------------------------------------------------------------------------------   3   20
------------AGUGGUCGUUAAUUGCUGGC-------------------------------------------------------------------------------------------------------------------   3   20
------------------------------------------------------------------------------------------------UAUGAGUAAAUGACCAUCUGC------------------------------   2   21
--------UAUAAGUGGUCGUUAAUUGCU----------------------------------------------------------------------------------------------------------------------   1   21
------------AGUGGUCGUUAAUUGCUGGCA------------------------------------------------------------------------------------------------------------------   1   21
--------------UGGUCGUUAAUUGCUGGC-------------------------------------------------------------------------------------------------------------------   1   18
--------------------------------AGACCUCAGUUUACUCACACU----------------------------------------------------------------------------------------------   1   21
-------------------------------------------------------------------------------------------------AUGAGUAAAUGACCAUCUGCCACC--------------------------   1   24
---------------------------------------------------------------------------------------------------------------------CACCAGUUAGCGGUCAUU------------   1   18
```

```
    A                      C       CCUCA         C     A    A     C   U
UUAU CUAUAuAAGUGGUCGUUAAUUG UGGcAGA     GUUUACUCA ACUAG --CC CUGGC GUC U
GGUA GAUaUGUUUACUGGCGAUUGAC AcCGUCU     UAAAUGAGU UGAUC   GG GAUCG CGG C
    C                      C       ACCAG         A     C G  C     U   U
                                                       G C
                                                      G   A
                                                       U-A
```

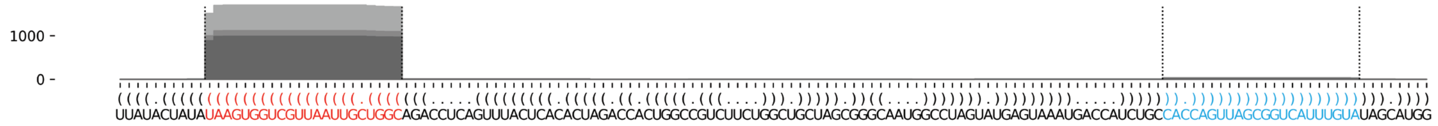


#### *D. discoideum* ddi-mir-7097

```
(((((((((.((((((((..(((((((((((((((..((((..((((.(((.......)))))))..)))))))))))))))))))..)))))))).)))))))))   -61.87 kcal/mol
UUUCGUUUUCuUCUCUACUAGUGCCGAAAUcAUUAAUAUUGCAUUCAUCAUUAUAUUUGUGUGAAAUCGAUUAAUGAuUUUGGCAGAAGUAGAGACGaAAACGAAA   count   length
Libraries combined          Precision: [ total = 91.2% | 5p-arm = 97.8% | 3p-arm = 90.1% ]
-----------------------------------------------------------------------------UUUUGGCAGAAGUAGAGACGA--------   404   21
----------UUCUCUACUAGUGCCGAAAUC---------------------------------------------------------------------------   80    21
-----------------------------------------------------------------------------UUUUGGCAGAAGUAGAGACG---------   36    20
-----------------------------------------------------------------------------UUUUGGCAGAAGUAGAGACGAAAA-----   46    24
-----------------------------------------------------------------------------UUUUGGCAGAAGUAGAGACGAAA------   40    23
-----------------------------------------------------------------------------UUUUGGCAGAAGUAGAGACGAA-------   25    22
----------UUCUCUACUAGUGCCGAAAU----------------------------------------------------------------------------   9     20
-----------------------------------------------------------------------------UUUUGGCAGAAGUAGAGAC----------   5     19
------------------------------------------------------------------------------UUUGGCAGAAGUAGAGACGA--------   3     20
---------CUUCUCUACUAGUGCCGAAAUC---------------------------------------------------------------------------   2     22
----------UUCUCUACUAGUGCCGAAA-----------------------------------------------------------------------------   1     19
----------------------------------------------------------------------------AUUUUGGCAGAAGUAGAGACGA--------   1     22
-----------------------------------------------------------------------------UUUUGGCAGAAGUAGAGACGAAAAC----   2     25
------------------------------------------------------------------------------UUUGGCAGAAGUAGAGACGAAA------   1     22
-------------------------------------------------------------------------------UUGGCAGAAGUAGAGACGA--------   1     19
--------------------------------------------------------------------------------UGGCAGAAGUAGAGACGAAAA-----   1     21
--------------------------------------------------------------------------------UGGCAGAAGUAGAGACGAA-------   1     19
Library = 1        Precision: [ total = 93.3% | 5p-arm = 97.4% | 3p-arm = 92.8% ]
-----------------------------------------------------------------------------UUUUGGCAGAAGUAGAGACGA--------   249   21
----------UUCUCUACUAGUGCCGAAAUC---------------------------------------------------------------------------   31    21
-----------------------------------------------------------------------------UUUUGGCAGAAGUAGAGACG---------   17    20
-----------------------------------------------------------------------------UUUUGGCAGAAGUAGAGACGAAAA-----   16    24
-----------------------------------------------------------------------------UUUUGGCAGAAGUAGAGACGAAA------   15    23
```

#### *D. discoideum* ddi-mir-7097-continued

```
((((((((((.((((((((..((((((((((((((..((((..((((.(((.......)))))))..))))))))))))))))))..)))))))).))))))))))      -61.87 kcal/mol
UUUCGUUUUCuUCUCUACUAGUGCCGAAAUcAUUAAUAUUGCAUUCAUCAUUAUAUUUGUGUGAAAUCGAUUAAUGAuUUUGGCAGAAGUAGAGACGaAAACGAAA      count   length
Libraries combined         Precision: [ total = 91.2% | 5p-arm = 97.8% | 3p-arm = 90.1% ]
-----------------------------------------------------------------------------UUUUGGCAGAAGUAGAGACGAA-------      12      22
----------UUCUCUACUAGUGCCGAAAU----------------------------------------------------------------------------      5       20
-----------------------------------------------------------------------------UUUUGGCAGAAGUAGAGAC----------      2       19
------------------------------------------------------------------------------UUUGGCAGAAGUAGAGACGA--------      2       20
---------CUUCUCUACUAGUGCCGAAAUC---------------------------------------------------------------------------      1       22
----------UUCUCUACUAGUGCCGAAA-----------------------------------------------------------------------------      1       19
----------------------------------------------------------------------------AUUUUGGCAGAAGUAGAGACGA--------      1       22
-----------------------------------------------------------------------------UUUUGGCAGAAGUAGAGACGAAAAC----      1       25
------------------------------------------------------------------------------UUUGGCAGAAGUAGAGACGAAA------      1       22
-------------------------------------------------------------------------------UUGGCAGAAGUAGAGACGA--------      1       19
--------------------------------------------------------------------------------UGGCAGAAGUAGAGACGAAAA-----      1       21
Library = 2        Precision: [ total = 89.7% | 5p-arm = 90.9% | 3p-arm = 89.4% ]
-----------------------------------------------------------------------------UUUUGGCAGAAGUAGAGACGA--------      31      21
----------UUCUCUACUAGUGCCGAAAUC---------------------------------------------------------------------------      9       21
-----------------------------------------------------------------------------UUUUGGCAGAAGUAGAGACGAAAA-----      5       24
-----------------------------------------------------------------------------UUUUGGCAGAAGUAGAGACGAAA------      4       23
-----------------------------------------------------------------------------UUUUGGCAGAAGUAGAGACG---------      3       20
-----------------------------------------------------------------------------UUUUGGCAGAAGUAGAGACGAA-------      3       22
---------CUUCUCUACUAGUGCCGAAAUC---------------------------------------------------------------------------      1       22
----------UUCUCUACUAGUGCCGAAAU----------------------------------------------------------------------------      1       20
-----------------------------------------------------------------------------UUUUGGCAGAAGUAGAGAC----------      1       19
Library = 3        Precision: [ total = 88.5% | 5p-arm = 100.0% | 3p-arm = 86.1% ]
-----------------------------------------------------------------------------UUUUGGCAGAAGUAGAGACGA--------      124     21
----------UUCUCUACUAGUGCCGAAAUC---------------------------------------------------------------------------      40      21
-----------------------------------------------------------------------------UUUUGGCAGAAGUAGAGACGAAAA-----      25      24
-----------------------------------------------------------------------------UUUUGGCAGAAGUAGAGACGAAA------      21      23
-----------------------------------------------------------------------------UUUUGGCAGAAGUAGAGACG---------      16      20
-----------------------------------------------------------------------------UUUUGGCAGAAGUAGAGACGAA-------      10      22
----------UUCUCUACUAGUGCCGAAAU----------------------------------------------------------------------------      3       20
-----------------------------------------------------------------------------UUUUGGCAGAAGUAGAGAC----------      2       19
-----------------------------------------------------------------------------UUUUGGCAGAAGUAGAGACGAAAAC----      1       25
------------------------------------------------------------------------------UUUGGCAGAAGUAGAGACGA--------      1       20
--------------------------------------------------------------------------------UGGCAGAAGUAGAGACGAA-------      1       19
```

```
          u        AG              AU    CA    U   UAU
UUUCGUUUUC UCUCUACU  UGCCGAAAUcAUUA  AUUG  UUCA CAU   \
AAAGCAAAaG AGAGAUGA  ACGGUUUuAGUAAU--UAGC  AAGU-GUG   A
          C        AG                    UA        UUU
```

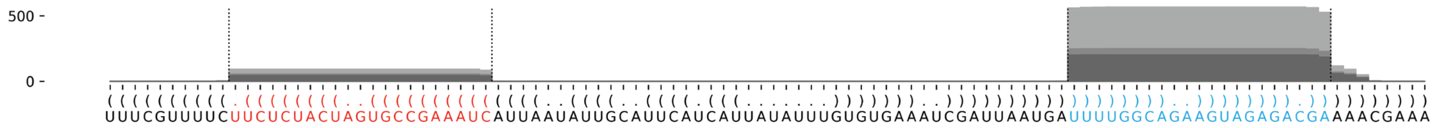

#### *D. discoideum* ddi-mir-1186 (ddi-mir-can_D1)

```
(((.(((((((((((((((((((((((((.((((((.((((.((((.((((.....)))).)))).)))).))))))))))))))))))))))))))))))))))).)))      -56.91 kcal/mol
UAUAGAAAUCuCUUUCUCUAAUUUCAUUUAuUAAAUUGUAAAUCACCCAUUUUAAUAAUGUGUGAGUUAUAUUUAAaAAUGAAAUUAGAGAAAGGGaUUUUCAUG      count    length
Libraries combined          Precision: [ total = 97.8% | 5p-arm = 100.0% | 3p-arm = 97.3% ]
----------------------------------------------------------------------------AAAUGAAAUUAGAGAAAGGGA--------      60       21
----------------------------------------------------------------------------AAAUGAAAUUAGAGAAAGGGAU-------      46       22
----------UCUUUCUCUAAUUUCAUUUA---------------------------------------------------------------------------      1        20
----------------------------------------------------------------------------AAAUGAAAUUAGAGAAAGGGAUU------      1        23
----------UCUUUCUCUAAUUUCAUUUAUU-------------------------------------------------------------------------      18       22
----------UCUUUCUCUAAUUUCAUUUAU--------------------------------------------------------------------------      2        21
----------------------------------------------------------------------------AAAUGAAAUUAGAGAAAGGG---------      3        20
-----------------------------------------------------------------------------AAUGAAAUUAGAGAAAGGGAU-------      3        21
Library = 1       Precision: [ total = 100.0% | 5p-arm = 100.0% | 3p-arm = 100.0% ]
----------------------------------------------------------------------------AAAUGAAAUUAGAGAAAGGGA--------      11       21
----------------------------------------------------------------------------AAAUGAAAUUAGAGAAAGGGAU-------      9        22
----------UCUUUCUCUAAUUUCAUUUA---------------------------------------------------------------------------      1        20
----------------------------------------------------------------------------AAAUGAAAUUAGAGAAAGGGAUU------      1        23
Library = 2       Precision: [ total = 93.8% | 5p-arm = 100.0% | 3p-arm = 92.3% ]
----------------------------------------------------------------------------AAAUGAAAUUAGAGAAAGGGA--------      8        21
----------------------------------------------------------------------------AAAUGAAAUUAGAGAAAGGGAU-------      4        22
----------UCUUUCUCUAAUUUCAUUUAUU-------------------------------------------------------------------------      3        22
-----------------------------------------------------------------------------AAUGAAAUUAGAGAAAGGGAU-------      1        21
Library = 3       Precision: [ total = 97.9% | 5p-arm = 100.0% | 3p-arm = 97.5% ]
----------------------------------------------------------------------------AAAUGAAAUUAGAGAAAGGGA--------      41       21
----------------------------------------------------------------------------AAAUGAAAUUAGAGAAAGGGAU-------      33       22
----------UCUUUCUCUAAUUUCAUUUAUU-------------------------------------------------------------------------      15       22
----------------------------------------------------------------------------AAAUGAAAUUAGAGAAAGGG---------      3        20
----------UCUUUCUCUAAUUUCAUUUAU--------------------------------------------------------------------------      2        21
-----------------------------------------------------------------------------AAUGAAAUUAGAGAAAGGGAU-------      2        21
```

```
   A                          A      U    A    C    UU
UAU GAAAUCuCUUUCUCUAAUUUCAUUU uUAAAU GUAA UCAC CAUU  \
GUA UUUUaGGGAAAGAGAUUAAAGUAAa-AAUUUA-UAUU AGUG GUAA  A
   C                                G    U    UA
```

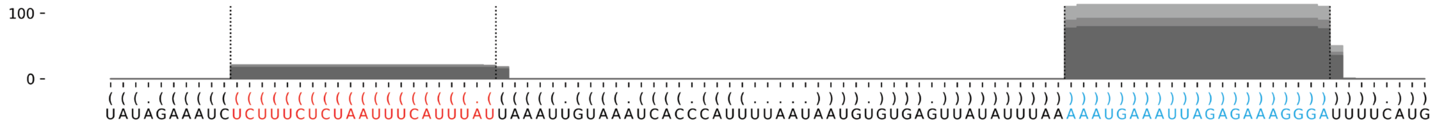

#### *D. firmibasis* dfi-mir-1190

(((((((((((((((((((((((((((((((((((((((((((((((((.(((((((((((((((...(((((..........(((((((((.......((((((((...)))))))))(((((.(((((.....(((((.......))))))))))).)))))))))))))))).......)))))..)))))))))))))))).))))))))))))))))))))))))))))))))))))))))))))))))) -142.84 kcal/mol

CAACCAAAUCaAAAUUAUCAAAUGUCCUCAGcAAGAUAAUUCACAUAAUUCUUUAUUAAAUUCUUUUUUAAAUGGGUUAACUCUGAUCUAAAAUAUUAACAAAAUUAUUACUAUAAUAAUUUAUAAAUUGACCAUAAUCAUUUCUUUUUAAAUGGGUUAUUUUAUAUUUAGAUCCAAAUAAUAUUUUAAAGAAUUUAAUAAACAAUUAUGUGAAUUAUCUUGCuGAGGACAUUUGAUAAUUUUGaUUUGGUUG

Libraries combined Precision: [ total = 76.9% | 5p-arm = 51.6% | 3p-arm = 78.1% ]

| read | count | length |
|---|---|---|
| UGAGGACAUUUGAUAAUUUUGA | 59509 | 22 |
| UGAGGACAUUUGAUAAUUUUG | 30890 | 21 |
| CUGAGGACAUUUGAUAAUUUU | 9961 | 21 |
| UGAGGACAUUUGAUAAUUUUGAU | 5278 | 23 |
| UGCUGAGGACAUUUGAUAAUU | 4367 | 21 |
| AAAAUUAUCAAAUGUCCUCAGC | 2990 | 22 |
| GCUGAGGACAUUUGAUAAUUU | 2755 | 21 |
| UUGCUGAGGACAUUUGAUAAU | 2676 | 21 |
| AAAUUAUCAAAUGUCCUCAGC | 2111 | 21 |
| UGAGGACAUUUGAUAAUUUU | 1774 | 20 |
| CUGAGGACAUUUGAUAAUUU | 1633 | 20 |
| CAAUUAUGUGAAUUAUCUUGC | 996 | 21 |
| CUGAGGACAUUUGAUAAUUUUG | 768 | 22 |
| UGAGGACAUUUGAUAAUUU | 657 | 19 |
| UGCUGAGGACAUUUGAUAAUUU | 568 | 22 |
| AUCUUGCUGAGGACAUUUGAU | 453 | 21 |
| UCUUGCUGAGGACAUUUGAUA | 370 | 21 |
| AAUUAUCAAAUGUCCUCAGC | 388 | 20 |
| UGAGGACAUUUGAUAAUUUUGAUU | 392 | 24 |
| ACAAUUAUGUGAAUUAUCUUGC | 231 | 22 |
| UUGCUGAGGACAUUUGAUAAUU | 337 | 22 |
| UGAGGACAUUUGAUAAUU | 208 | 18 |
| UGCUGAGGACAUUUGAUAAU | 254 | 20 |
| AAUUAUCAAAUGUCCUCAGCA | 230 | 21 |
| GAGGACAUUUGAUAAUUUUGA | 206 | 21 |
| GCUGAGGACAUUUGAUAAUU | 221 | 20 |
| CUGAGGACAUUUGAUAAUU | 142 | 19 |
| ACAAUUAUGUGAAUUAUCUUG | 152 | 21 |
| CUGAGGACAUUUGAUAAUUUUGA | 133 | 23 |
| AAAAUUAUCAAAUGUCCUCAG | 127 | 21 |
| AUCUUGCUGAGGACAUUUGAUA | 97 | 22 |
| UGAAUUAUCUUGCUGAGGAC | 112 | 20 |
| GAGGACAUUUGAUAAUUUUG | 115 | 20 |
| AGGACAUUUGAUAAUUUUGAU | 78 | 21 |
| AAAUUAUCAAAUGUCCUCAG | 61 | 20 |
| AAUUAUCUUGCUGAGGACAUU | 75 | 21 |
| GUGAAUUAUCUUGCUGAGGAC | 64 | 21 |
| CAAUUAUGUGAAUUAUCUUGCU | 72 | 22 |
| AGGACAUUUGAUAAUUUUGAUU | 53 | 22 |
| UGAAUUAUCUUGCUGAGGACA | 61 | 21 |
| UCUUGCUGAGGACAUUUGAUAA | 45 | 22 |
| AUCUUGCUGAGGACAUUUGA | 21 | 20 |
| CUGAGGACAUUUGAUAAUUUUGAU | 23 | 24 |
| AUUAUCAAAUGUCCUCAGC | 30 | 19 |
| AUCUUGCUGAGGACAUUUGAUAA | 29 | 23 |
| UCUUGCUGAGGACAUUUGAU | 29 | 20 |
| UUGCUGAGGACAUUUGAUAA | 43 | 20 |
| AAUUAUGUGAAUUAUCUUGCU | 26 | 21 |
| AUUAUCUUGCUGAGGACAUU | 24 | 20 |
| UUGCUGAGGACAUUUGAUAAUUU | 24 | 23 |
| CAAUUAUGUGAAUUAUCUUG | 34 | 20 |
| AAAUUAUCAAAUGUCCUCAGCA | 17 | 22 |
| ACAAUUAUGUGAAUUAUCUUGCU | 24 | 23 |
| CUGAGGACAUUUGAUAAU | 26 | 18 |
| UAUCAAAUGUCCUCAGCAAGA | 18 | 21 |
| AUUAUCUUGCUGAGGACAUUU | 13 | 21 |
| UUAUCUUGCUGAGGACAUUUG | 10 | 21 |
| AAUUAUCAAAUGUCCUCAG | 11 | 19 |
| AAUUAUCAAAUGUCCUCAGCAA | 8 | 22 |
| AAUUAUGUGAAUUAUCUUGC | 13 | 20 |
| UAUCUUGCUGAGGACAUUUGA | 10 | 21 |
| UCUUGCUGAGGACAUUUGAUAAU | 18 | 23 |
| GCUGAGGACAUUUGAUAAUUUU | 12 | 22 |

#### *D. firmibasis* dfi-mir-1190 continued

| Sequence / structure | count | length |
|---|---|---|
| ((((((((((((((((((((((((((((((((((((((((((((((((((.(((((((((((((((...(((((..........(((((((((.......((((((((...))))))))(((((.(((((.....((((.......))))))))).)))))))))))))))).......)))))..))))))))))))))).))))))))))))))))))))))))))))))))))))))))))))))))) | −142.84 kcal/mol | |
| CAACCAAAUCaAAAUUAUCAAAUGUCCUCAGcAAGAUAAUUCACAUAAUUCUUUAUUAAAUUCUUUUUUAAAUGGGUUAACUCUGAUCUAAAAUAUUAACAAAAUUAUUACUAUAAUAAUUUAUAAAUUGACCAUAAUCAUUUCUUUUUUAAAUGGGUUAUUUUAUAUUUUAGAUCCAAAUAAUAUUUUUAAAGAAUUUAAUAAACAAUUAUGUGAAUUAUCUUGCuGAGGACAUUUGAUAAUUUUGaUUUGGUUG | count | length |
| Libraries combined — Precision: [ total = 76.9% \| 5p-arm = 51.6% \| 3p-arm = 78.1% ] | | |
| GCUGAGGACAUUUGAUAAU | 11 | 19 |
| AGGACAUUUGAUAAUUUUGA | 9 | 20 |
| UCAAAUGUCCUCAGCAAGAUA | 12 | 21 |
| AUCUUGCUGAGGACAUUUGAUAAU | 12 | 24 |
| CUUGCUGAGGACAUUUGAUA | 4 | 20 |
| ACAUUUGAUAAUUUUGAUUUG | 3 | 21 |
| AACAAUUAUGUGAAUUAUCUUGCU | 6 | 24 |
| CAAUUAUGUGAAUUAUCU | 2 | 18 |
| AUUAUGUGAAUUAUCUUGC | 4 | 19 |
| AUGUGAAUUAUCUUGCUGAGG | 17 | 21 |
| UGAAUUAUCUUGCUGAGGACAUU | 5 | 23 |
| GAAUUAUCUUGCUGAGGACAU | 6 | 21 |
| GAAUUAUCUUGCUGAGGACA | 6 | 20 |
| AAUUAUCUUGCUGAGGACAU | 4 | 20 |
| UGCUGAGGACAUUUGAUAA | 6 | 19 |
| GAGGACAUUUGAUAAUUUU | 5 | 19 |
| UUUGAUAAUUUUGAUUUGGUU | 5 | 21 |
| UUGAUAAUUUUGAUUUGGUUG | 4 | 21 |
| UCAAAAUUAUCAAAUGUCCUC | 1 | 21 |
| AAAAUUAUCAAAUGUCCUCAGCA | 3 | 23 |
| AAUUAUCAAAUGUCCUCAGCAAG | 2 | 23 |
| UUAUCAAAUGUCCUCAGC | 1 | 18 |
| UUAUCAAAUGUCCUCAGCA | 3 | 19 |
| UAUCAAAUGUCCUCAGCAAG | 2 | 20 |
| UCCUCAGCAAGAUAAUUCACA | 2 | 21 |
| AAGAUAAUUCACAUAAUUCUU | 1 | 21 |
| AAGAUAAUUCACAUAAUUCU | 4 | 20 |
| AAAUGGGUUAACUCUGAUCUA | 1 | 21 |
| AACAAUUAUGUGAAUUAUCUUGC | 4 | 23 |
| CAAUUAUGUGAAUUAUCUUGCUGA | 1 | 24 |
| AUUAUGUGAAUUAUCUUGCUG | 7 | 21 |
| AUGUGAAUUAUCUUGCUGAGGA | 2 | 22 |
| AUGUGAAUUAUCUUGCUGA | 1 | 19 |
| UGUGAAUUAUCUUGCUGAG | 1 | 19 |
| UGAAUUAUCUUGCUGAGGACAU | 3 | 22 |
| GAAUUAUCUUGCUGAGGACAUU | 2 | 22 |
| AAUUAUCUUGCUGAGGACA | 2 | 19 |
| AUUAUCUUGCUGAGGACAUUUGAU | 1 | 24 |
| UUAUCUUGCUGAGGACAUUUGAU | 2 | 23 |
| UAUCUUGCUGAGGACAUUUGAU | 2 | 22 |
| AUCUUGCUGAGGACAUUUG | 1 | 19 |
| CUUGCUGAGGACAUUUGAUAAU | 3 | 22 |
| UUGCUGAGGACAUUUGAUA | 4 | 19 |
| UGAGGACAUUUGAUAAUUUUGAUUU | 10 | 25 |
| GAGGACAUUUGAUAAUUUUGAUU | 3 | 23 |
| AGGACAUUUGAUAAUUUUG | 5 | 19 |
| GGACAUUUGAUAAUUUUGAUU | 6 | 21 |
| CAUUUGAUAAUUUUGAUUUGG | 1 | 21 |
| UCAAAAUUAUCAAAUGUCCUCAGC | 1 | 24 |
| UCAAAAUUAUCAAAUGUCCU | 1 | 20 |
| AAAUUAUCAAAUGUCCUCAGCAA | 1 | 23 |
| AUUAUCAAAUGUCCUCAG | 3 | 18 |
| AUUAUCAAAUGUCCUCAGCAA | 1 | 21 |
| AUUAUCAAAUGUCCUCAGCA | 2 | 20 |
| UUAUCAAAUGUCCUCAGCAAG | 4 | 21 |
| AUCAAAUGUCCUCAGCAAGAUA | 1 | 22 |
| AUCAAAUGUCCUCAGCAAGAUAAUUCACA | 1 | 29 |
| UCAAAUGUCCUCAGCAAGAU | 1 | 20 |
| CAAAUGUCCUCAGCAAGAUA | 3 | 20 |
| UGUCCUCAGCAAGAUAAUUC | 1 | 20 |
| CUCAGCAAGAUAAUUCACAUA | 1 | 21 |
| CAAGAUAAUUCACAUAAUUCU | 1 | 21 |
| CACAUAAUUCUUUAUUAAAUU | 1 | 21 |

#### *D. firmibasis* dfi-mir-1190 continued

```
((((((((((((((((((((((((((((((((((((((((((((((((((.(((((((((((((((...(((((..........((((((((((.......(((((((((...)))))))))(((((.(((((.....(((((.......))))))))))).))))))))))))))).......))))).)))))))))))))))).))))))))))))))))))))))))))))))))))))))))))))))))))     -142.84 kcal/mol
CAACCAAAUCaAAAUUAUCAAAUGUCCUCAGcAAGAUAAUUCACAUAAUUCUUUAUUAAAUUCUUUUUAAAUGGGUUAACUCUGAUCUAAAAUAUUAACAAAAUUAUUACUAUAAUAAUUUAUAAAUUGACCAUAAUCAUUUCUUUUUAAAUGGGUUAUUUAUAUUUAGAUCCAAAUAAUAUUUUAAAGAAUUUAAUAAACAAUUAUGUGAAUUAUCUUGCuGAGGACAUUUGAUAAUUUUGaUUUGGUUG
```

Libraries combined    Precision: [ total = 76.9% | 5p-arm = 51.6% | 3p-arm = 78.1% ]

| Read | count | length |
|---|---|---|
| CUUUUUAAAUGGGUUAACUC | 1 | 21 |
| AACAAUUAUGUGAAUUAUCUUG | 13 | 22 |
| ACAAUUAUGUGAAUUAUC | 1 | 18 |
| CAAUUAUGUGAAUUAUCUUGCUG | 1 | 23 |
| UUAUGUGAAUUAUCUUGCUGA | 1 | 21 |
| UAUGUGAAUUAUCUUGCUGAG | 2 | 21 |
| UAUGUGAAUUAUCUUGCUGAGG | 2 | 22 |
| AUGUGAAUUAUCUUGCUGAG | 1 | 20 |
| GUGAAUUAUCUUGCUGAGGACA | 1 | 22 |
| UGAAUUAUCUUGCUGAGGA | 1 | 19 |
| GAAUUAUCUUGCUGAGGAC | 1 | 19 |
| AAUUAUCUUGCUGAGGACAUUU | 1 | 22 |
| AUUAUCUUGCUGAGGACAU | 2 | 19 |
| AUUAUCUUGCUGAGGACAUUUGA | 1 | 23 |
| UAUCUUGCUGAGGACAUUUGAUA | 1 | 23 |
| UAUCUUGCUGAGGACAUUUG | 1 | 20 |
| UCUUGCUGAGGACAUUUGAUAAUU | 1 | 24 |
| CUUGCUGAGGACAUUUGAUAA | 6 | 21 |
| UGCUGAGGACAUUUGAUAAUUUU | 1 | 23 |
| GCUGAGGACAUUUGAUAA | 1 | 18 |
| GAGGACAUUUGAUAAUUU | 5 | 18 |
| GAGGACAUUUGAUAAUUUUGAU | 8 | 22 |
| AUUUGAUAAUUUUGAUUUGGU | 1 | 21 |
| AUUUGAUAAUUUUGAUUUGGUU | 1 | 22 |
| UUGAUAAUUUUGAUUUGGU | 1 | 19 |

```
                                                 C                UUU      GGUUAACUCU           AUUAACA          C
CAACCAAAUCaAAAUUAUCAAAUGUCCUCAGcAAGAUAAUUCACAUAAUU UUUAUUAAAUUCUUU    AAAUG          GAUCUAAAAU        AAAUUAUUA U
GUUGGUUUaGUUUUAAUAGUUUACAGGAGuCGUUCUAUUAAGUGUAUUAA AAAUAAUUUAAGAAA   -UUUAU        ---CUAGAUUUUA---\   AUUGGGUAAA /
                                                 C                UU       AAUAAAC                  AU           A
                                                                                                    UA
                                                                                                    UA
                                                                                                    UA       AUAAU      CUU
                                                                                                   U   UUGACC      CAUUU    U
                                                                                                    \--AUUGG-----GUAAA    /
                                                                                                                     UUU
```

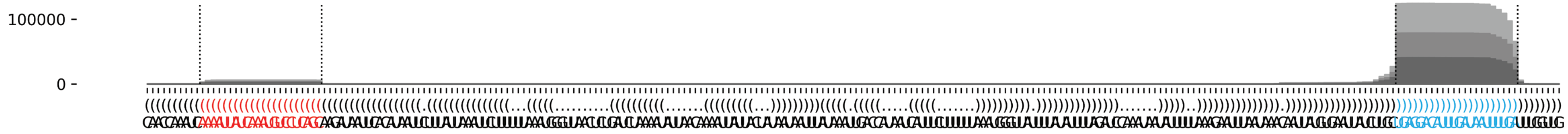

#### *D. firmibasis* dfi-mir-1191

```
((((((((((((((((((.(((.((((((((((.(((((((((((((((((((.(..(((.(((((.(.(((((((.(((((((......)))))))......))))))).))))))...)))..)...)))))).....)))))))))))))).)))))))))).))).)))))))))))))))))) -95.3 kcal/mol
UCAUUAUGCAaACAAGUGCAAUCAGGAAAUaAAUACUAUAACUGAGCAAGUAUCAUUUCACAUCAAACAAACAAGAAGAAUUAAAAAGAGUUAGUUUAUUGGCUCUUGUUGGUUGAUCAUUGAUUUUUAAUACUGCGGUUGUUCAGUUAUAGUUUUUAuUUCCUCGUUACACUUGUUUgCAUAAUGA count length
Libraries combined    Precision: [ total = 97.1% | 5p-arm = 44.6% | 3p-arm = 97.8% ]
--------------------------------------------------------------------------------------------------------------------------------------------------------------UUUCCUCGUUACACUUGUUUG-------- 74616 21
--------------------------------------------------------------------------------------------------------------------------------------------------------------UUUCCUCGUUACACUUGUUUGC------- 11194 22
--------------------------------------------------------------------------------------------------------------------------------------------------------------UUUCCUCGUUACACUUGUUUGCA------ 5823 23
----------AACAAGUGCAAUCAGGAAAUA------------------------------------------------------------------------------------------------------------------------------------------------------------ 464 21
--------------------------------------------------------------------------------------------------------------------------------------------------------------UUUCCUCGUUACACUUGUUU--------- 634 20
--------CAAACAAGUGCAAUCAGGAAAUA------------------------------------------------------------------------------------------------------------------------------------------------------------ 328 23
--------------------------------------------------------------------------------------------------------------------------------------------------------------UUUCCUCGUUACACUUGUUUGCAU----- 526 24
------------------------------------------------------------------------------------------------------------------------------------------------------------UAUUUCCUCGUUACACUUGUU---------- 334 21
---------AAACAAGUGCAAUCAGGAAAUA------------------------------------------------------------------------------------------------------------------------------------------------------------ 228 22
------------------------------------------------------------------------------------------------------------------------------------------------------------UAUUUCCUCGUUACACUUGUUU--------- 337 22
--------------------------------------------------------------------------------------------------------------------------------------------------------------UUUCCUCGUUACACUUGUU---------- 281 19
----------AACAAGUGCAAUCAGGAAAU------------------------------------------------------------------------------------------------------------------------------------------------------------- 89 20
---------------------------------------------------------------------------------------------------------------------------------------------------------------UUCCUCGUUACACUUGUUUGCA------ 156 22
----------------------------------------------------------------------------------------------------------------------------------------------------------------UCCUCGUUACACUUGUUUGCA------ 119 21
--------CAAACAAGUGCAAUCAGGAAAU------------------------------------------------------------------------------------------------------------------------------------------------------------- 83 22
---------------------------------------------------------------------------------------------------------------------------------------------------------------UUCCUCGUUACACUUGUUUG-------- 89 20
-----------------------------------------------------------------------------------------------------------------------------------------------------------UUAUUUCCUCGUUACACUUGU----------- 76 21
---------------------------------------------------------------------------------------------------------------------------------------------------------------UUCCUCGUUACACUUGUUUGC------- 81 21
---------------------------------------------------------------------------------------------------------------------------------------------------------UUUUAUUUCCUCGUUACACUUG------------ 45 22
-------------------------------------------------------------------------------------------------------------------------------------------------------------AUUUCCUCGUUACACUUGUUU--------- 51 21
---------AAACAAGUGCAAUCAGGAAAU------------------------------------------------------------------------------------------------------------------------------------------------------------- 30 21
----------------------------------------------------------------------------------------------------------------------------------------------------------------UCCUCGUUACACUUGUUUGC------- 37 20
---------------------------------------------------------------------------------------------------------------------------------------------------------UUUUAUUUCCUCGUUACACUU------------- 20 21
---------------------------------------------------------------------------------------------------------------------------------------------------------UUUUAUUUCCUCGUUACACU-------------- 31 20
--------------------------------------------------------------------------------------------------------------------------------------------------------------UUUCCUCGUUACACUUGU----------- 30 18
------------------------------------------------------------------------------------------------------------------------------------------------------------UAUUUCCUCGUUACACUUGUUUG-------- 7 23
----------------------------------------------------------------------------------------------------------------------------------------------------------------UCCUCGUUACACUUGUUUG-------- 29 19
--------------------------------------------------------------------------------------------------------------------------------------------------------------UUUCCUCGUUACACUUGUUUGCAUA---- 19 25
--------CAAACAAGUGCAAUCAGGAAA-------------------------------------------------------------------------------------------------------------------------------------------------------------- 7 21
----------AACAAGUGCAAUCAGGAAA-------------------------------------------------------------------------------------------------------------------------------------------------------------- 4 19
-----------------------------------------------------------------------------------------------------------------------------------------------------------UUAUUUCCUCGUUACACUUGUU---------- 10 22
------------------------------------------------------------------------------------------------------------------------------------------------------------UAUUUCCUCGUUACACUUGU----------- 18 20
---------------------------------------------------------------------------------------------------------------------------------------------------------------UUCCUCGUUACACUUGUU---------- 3 18
-----------ACAAGUGCAAUCAGGAAAUA------------------------------------------------------------------------------------------------------------------------------------------------------------ 8 20
---------------------------------------------------------------------------------------------------------------------------------------------------------------UUCCUCGUUACACUUGUUUGCAU----- 11 23
-------GCAAACAAGUGCAAUCAGGAAAU------------------------------------------------------------------------------------------------------------------------------------------------------------- 2 23
----------AACAAGUGCAAUCAGGAAAUAA----------------------------------------------------------------------------------------------------------------------------------------------------------- 3 22
-----------ACAAGUGCAAUCAGGAAAU------------------------------------------------------------------------------------------------------------------------------------------------------------- 1 19
---------------------UCAGGAAAUAAAUACUAUAAC------------------------------------------------------------------------------------------------------------------------------------------------- 1 21
------------------------------------------------------------------------------------------------UAUUGGCUCUUGUUGGUUGAUCAUU------------------------------------------------------------------ 1 25
-------------------------------------------------------------------------------------------------------------------------------------UGCGGUUGUUCAGUUAUAGUU--------------------------------- 4 21
---------------------------------------------------------------------------------------------------------------------------------------CGGUUGUUCAGUUAUAGUUU-------------------------------- 1 20
------------------------------------------------------------------------------------------------------------------------------------------------AGUUAUAGUUUUUAUUUCCUC---------------------- 1 21
--------------------------------------------------------------------------------------------------------------------------------------------------UUAUAGUUUUUAUUUCCUCGUU------------------- 1 22
-------------------------------------------------------------------------------------------------------------------------------------------------------GUUUUUAUUUCCUCGUUACACU-------------- 6 22
----------------------------------------------------------------------------------------------------------------------------------------------------------UUUAUUUCCUCGUUACACUUG------------ 1 21
-----------------------------------------------------------------------------------------------------------------------------------------------------------UUAUUUCCUCGUUACACUUG------------ 13 20
------------------------------------------------------------------------------------------------------------------------------------------------------------UAUUUCCUCGUUACACUUGUUUGC------- 1 24
------------------------------------------------------------------------------------------------------------------------------------------------------------UAUUUCCUCGUUACACUUG------------ 3 19
----------------------------------------------------------------------------------------------------------------------------------------------------------------UCCUCGUUACACUUGUUU--------- 3 18
-----------------------------------------------------------------------------------------------------------------------------------------------------------------CCUCGUUACACUUGUUUGCA------ 5 20
--------CAAACAAGUGCAAUCAGGAAAUAA----------------------------------------------------------------------------------------------------------------------------------------------------------- 1 24
---------AAACAAGUGCAAUCAGGAAAUAA----------------------------------------------------------------------------------------------------------------------------------------------------------- 3 23
```

#### *D. firmibasis* dfi-mir-1191 continued

```
((((((((((((((((((.(((.((((((((((.(((((((((((((((((((.(..(((.(((((.(.(((((((.(((((((......)))))))......))))))).))))))...)))..)...)))))).....)))))))))))))).)))))))))).))).)))))))))))))))))) -95.3 kcal/mol
UCAUUAUGCAaACAAGUGCAAUCAGGAAAUaAAUACUAUAACUGAGCAAGUAUCAUUUCACAUCAAACAAACAAGAAGAAUUAAAAAGAGUUAGUUUAUUGGCUCUUGUUGGUUGAUCAUUGAUUUUUAAUACUGCGGUUGUUCAGUUAUAGUUUUUAuUUCCUCGUUACACUUGUUUgCAUAAUGA count length
Libraries combined    Precision: [ total = 97.1% | 5p-arm = 44.6% | 3p-arm = 97.8% ]
---------AAACAAGUGCAAUCAGGAAA-------------------------------------------------------------------------------------------------------------------------------------------------------------- 3 20
--------------AGUGCAAUCAGGAAAUAAAUA-------------------------------------------------------------------------------------------------------------------------------------------------------- 1 21
----------------------------------------------------------------------------------AAAAAGAGUUAGUUUAUUGGC------------------------------------------------------------------------------------ 1 21
-------------------------------------------------------------------------------------------------------------------------------------UGCGGUUGUUCAGUUAUAGUUU-------------------------------- 1 22
--------------------------------------------------------------------------------------------------------------------------------------GCGGUUGUUCAGUUAUAGUUU-------------------------------- 1 21
-----------------------------------------------------------------------------------------------------------------------------------------------------UAGUUUUUAUUUCCUCGUUAC----------------- 4 21
--------------------------------------------------------------------------------------------------------------------------------------------------------UUUUUAUUUCCUCGUUACACU-------------- 1 21
---------------------------------------------------------------------------------------------------------------------------------------------------------UUUUAUUUCCUCGUUACAC--------------- 1 19
----------------------------------------------------------------------------------------------------------------------------------------------------------UUUAUUUCCUCGUUACACUUGU----------- 1 22
-------------------------------------------------------------------------------------------------------------------------------------------------------------AUUUCCUCGUUACACUUGUUUG-------- 10 22
-------------------------------------------------------------------------------------------------------------------------------------------------------------AUUUCCUCGUUACACUUGUU---------- 3 20
--------------------------------------------------------------------------------------------------------------------------------------------------------------UUUCCUCGUUACACUUGUUUGCAUAA--- 2 26
---------------------------------------------------------------------------------------------------------------------------------------------------------------UUCCUCGUUACACUUGUUU--------- 2 19
---------------------------------------------------------------------------------------------------------------------------------------------------------------UUCCUCGUUACACUUGUUUGCAUA---- 1 24
----------------------------------------------------------------------------------------------------------------------------------------------------------------UCCUCGUUACACUUGUUUGCAU----- 8 22
-----------------------------------------------------------------------------------------------------------------------------------------------------------------CCUCGUUACACUUGUUUG-------- 3 18
-----------------------------------------------------------------------------------------------------------------------------------------------------------------CCUCGUUACACUUGUUUGC------- 1 19
-----------------------------------------------------------------------------------------------------------------------------------------------------------------CCUCGUUACACUUGUUUGCAU----- 4 21
--------------------------------------------------------------------------------------------------------------------------------------------------------------------CGUUACACUUGUUUGCAUAAU-- 1 21
---------------------------------------------------------------------------------------------------------------------------------------------------------------------GUUACACUUGUUUGCAUAAUG- 2 21
```

```
                  C   C          U                        C-- UU   C       A A       A            AA
UCAUUAUGCAaACAAGUG AAU AGGAAAUaAA ACUAUAACUGAGCA-----AGUAU   A  UCA --AUCAA C AACAAGA -----GAAUUAA  A
AGUAAUACgUUUGUUCAC UUG UCCUUuAUUU UGAUAUUGACUUGU     UCAUA   U  AGU   UAGUU-G UUGUUCU      UUUGAUU  G
                  A   C          U              UGGCG     AUU UU   UAC       G       CGGUUA       GA
```

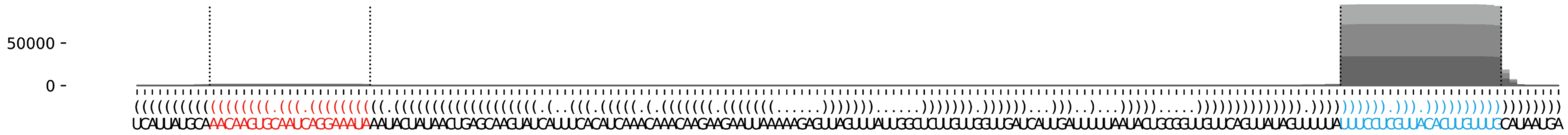

#### *D. firmibasis* dfi-mir-1192

```
((((((((.((.(((((((.((((((((((((((.((((...(((((((((........))))))))).)))).)))))))))))))).))))))).)).))))))))    -69.88 kcal/mol
UUUAUGCAGGuGUACCAAGACAAUCUCCAAgUACGUAUGGUAACUCAAUUGAAAAAUAAUGAUUGAGUACAUAAGUACUuGGAGAUUGGCUUGGUAAACaUGCAUAAA    count    length
Libraries combined          Precision: [ total = 84.5% | 5p-arm = 87.9% | 3p-arm = 76.9% ]
----------UGUACCAAGACAAUCUCCAAG-----------------------------------------------------------------------------    51       21
----UGCAGGUGUACCAAGACAAUC-----------------------------------------------------------------------------------    5        21
-------------------------------------------------------------------------------UGGAGAUUGGCUUGGUAAACA--------    9        21
--------------------------------------------------------------------------------GGAGAUUGGCUUGGUAAACAUG------    2        22
---------------------------------------------------------------------------UACUUGGAGAUUGGCUUGGUA------------    1        21
-------------------------------------------------------------------------------UGGAGAUUGGCUUGGUAAAC---------    10       20
--------------------------------------------------------------------------------GGAGAUUGGCUUGGUAAAC---------    1        19
------------UACCAAGACAAUCUCCAAGUA---------------------------------------------------------------------------    1        21
------------UACCAAGACAAUCUCCAAG-----------------------------------------------------------------------------    1        19
-----------------------------------------------------------------------------CUUGGAGAUUGGCUUGGUAAACA--------    2        23
-------------------------------------------------------------------------------UGGAGAUUGGCUUGGUAAA----------    1        19
Library = 1         Precision: [ total = 68.2% | 5p-arm = 80.0% | 3p-arm = 42.9% ]
----------UGUACCAAGACAAUCUCCAAG-----------------------------------------------------------------------------    12       21
----UGCAGGUGUACCAAGACAAUC-----------------------------------------------------------------------------------    3        21
-------------------------------------------------------------------------------UGGAGAUUGGCUUGGUAAACA--------    2        21
--------------------------------------------------------------------------------GGAGAUUGGCUUGGUAAACAUG------    2        22
---------------------------------------------------------------------------UACUUGGAGAUUGGCUUGGUA------------    1        21
-------------------------------------------------------------------------------UGGAGAUUGGCUUGGUAAAC---------    1        20
--------------------------------------------------------------------------------GGAGAUUGGCUUGGUAAAC---------    1        19
Library = 2         Precision: [ total = 92.0% | 5p-arm = 94.3% | 3p-arm = 86.7% ]
----------UGUACCAAGACAAUCUCCAAG-----------------------------------------------------------------------------    33       21
-------------------------------------------------------------------------------UGGAGAUUGGCUUGGUAAACA--------    7        21
-------------------------------------------------------------------------------UGGAGAUUGGCUUGGUAAAC---------    5        20
-----------------------------------------------------------------------------CUUGGAGAUUGGCUUGGUAAACA--------    2        23
----UGCAGGUGUACCAAGACAAUC-----------------------------------------------------------------------------------    1        21
------------UACCAAGACAAUCUCCAAGUA---------------------------------------------------------------------------    1        21
-------------------------------------------------------------------------------UGGAGAUUGGCUUGGUAAA----------    1        19
Library = 3         Precision: [ total = 83.3% | 5p-arm = 75.0% | 3p-arm = 100.0% ]
----------UGUACCAAGACAAUCUCCAAG-----------------------------------------------------------------------------    6        21
-------------------------------------------------------------------------------UGGAGAUUGGCUUGGUAAAC---------    4        20
----UGCAGGUGUACCAAGACAAUC-----------------------------------------------------------------------------------    1        21
------------UACCAAGACAAUCUCCAAG-----------------------------------------------------------------------------    1        19
```

```
        G  G       A               G    GUA         AAA
UUUAUGCA Gu UACCAAG CAAUCUCCAAgUAC UAUG   ACUCAAUUG   A
AAAUACGU CA AUGGUUC GUUAGAGGuUCAUG AUAC --UGAGUUAGU   A
        a  A       G               A    A           AAU
```

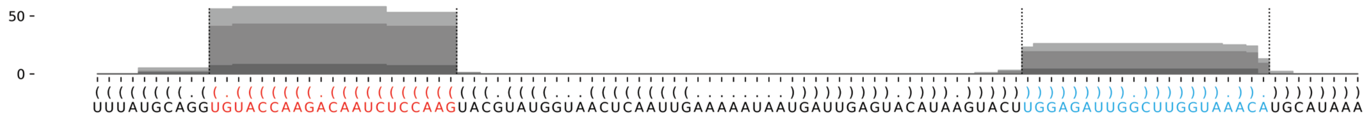

#### *D. firmibasis* dfi-mir-1195

```
(((.(((.(((((((((.(((((((((((((((((((..(((.(((((..((((((((....))))))))...........))))).)))..))))))))))))))))))).))))))))).))).)))      -69.84 kcal/mol
AGGUGAUCGUuGAUGUCGCCAAUGUAAUAAcAGAGGUUAAUUAUAUUAAAUAAUAUUUCAAUAAAUAUUAACAUUAACUUCUAAUACAAUAUACCUCUGUuAUUACAUUGGGGAUAUCAAcCGUCCCCU      count    length
Libraries combined         Precision: [ total = 97.7% | 5p-arm = 97.8% | 3p-arm = 34.5% ]
----------UGAUGUCGCCAAUGUAAUAAC--------------------------------------------------------------------------------------------------      530751   21
----------UGAUGUCGCCAAUGUAAUAA---------------------------------------------------------------------------------------------------      6469     20
---UGAUCGUUGAUGUCGCCAAUG---------------------------------------------------------------------------------------------------------      6127     21
----------UGAUGUCGCCAAUGUAAUAACA-------------------------------------------------------------------------------------------------      3143     22
---------UUGAUGUCGCCAAUGUAAUAAC--------------------------------------------------------------------------------------------------      1647     22
-----------GAUGUCGCCAAUGUAAUAAC--------------------------------------------------------------------------------------------------      1735     20
---------UUGAUGUCGCCAAUGUAAUAA---------------------------------------------------------------------------------------------------      701      21
------------AUGUCGCCAAUGUAAUAACAG------------------------------------------------------------------------------------------------      587      21
---------UUGAUGUCGCCAAUGUAAU-----------------------------------------------------------------------------------------------------      477      19
----------UGAUGUCGCCAAUGUAAU-----------------------------------------------------------------------------------------------------      483      18
-----------------------------------------------------------------------------------------------------AUUACAUUGGGGAUAUCAACC-------      398      21
------------AUGUCGCCAAUGUAAUAAC--------------------------------------------------------------------------------------------------      340      19
----------UGAUGUCGCCAAUGUAAUA----------------------------------------------------------------------------------------------------      305      19
----------------------------------------------------------------------------------------------------UAUUACAUUGGGGAUAUCAAC--------      188      21
---------UUGAUGUCGCCAAUGUAAUA----------------------------------------------------------------------------------------------------      165      20
-----------GAUGUCGCCAAUGUAAUAACA-------------------------------------------------------------------------------------------------      170      21
----------------------------------------------------------------------------------------------------UAUUACAUUGGGGAUAUCAA---------      77       20
------UCGUUGAUGUCGCCAAUGUA-------------------------------------------------------------------------------------------------------      68       20
-----------------------------------------------------------------------------------------------------AUUACAUUGGGGAUAUCAAC--------      55       20
-----------GAUGUCGCCAAUGUAAUAA---------------------------------------------------------------------------------------------------      59       19
-----AUCGUUGAUGUCGCCAAUGUA-------------------------------------------------------------------------------------------------------      39       21
----------UGAUGUCGCCAAUGUAAUAACAG------------------------------------------------------------------------------------------------      40       23
---UGAUCGUUGAUGUCGCCAA-----------------------------------------------------------------------------------------------------------      31       19
--------------GUCGCCAAUGUAAUAACAGAG----------------------------------------------------------------------------------------------      16       21
------------AUGUCGCCAAUGUAAUAACAGA-----------------------------------------------------------------------------------------------      27       22
-----------------------------------------------------------------------------------------------UCUGUUAUUACAUUGGGGAU--------------      12       20
---UGAUCGUUGAUGUCGCCAAU----------------------------------------------------------------------------------------------------------      20       20
-------------UGUCGCCAAUGUAAUAAC--------------------------------------------------------------------------------------------------      9        18
-----------------------------------------------------------------------------------------------------AUUACAUUGGGGAUAUCAA---------      18       19
---UGAUCGUUGAUGUCGCCA------------------------------------------------------------------------------------------------------------      9        18
----------------------------------------------------------------------------------------------------UAUUACAUUGGGGAUAUC-----------      9        18
-----AUCGUUGAUGUCGCCAAUG---------------------------------------------------------------------------------------------------------      12       19
-------------UGUCGCCAAUGUAAUAACAGAG----------------------------------------------------------------------------------------------      5        22
-----------GAUGUCGCCAAUGUAAUAACAG------------------------------------------------------------------------------------------------      8        22
--------------GUCGCCAAUGUAAUAACAG------------------------------------------------------------------------------------------------      5        19
--------------------------------------------------------------------------------------------------GUUAUUACAUUGGGGAUAUC-----------      4        20
---------------------------------------------------------------------------------------------------UUAUUACAUUGGGGAUAUCAA---------      4        21
-----------------------------------------------------------------------------------------------------AUUACAUUGGGGAUAUCAACCG------      7        22
---UGAUCGUUGAUGUCGCCAAUGU--------------------------------------------------------------------------------------------------------      11       22
------UCGUUGAUGUCGCCAAUG---------------------------------------------------------------------------------------------------------      2        18
----------UGAUGUCGCCAAUGUAAUAACAGA-----------------------------------------------------------------------------------------------      4        24
-----------GAUGUCGCCAAUGUAAUA----------------------------------------------------------------------------------------------------      6        18
------------AUGUCGCCAAUGUAAUAACA-------------------------------------------------------------------------------------------------      3        20
---------------UCGCCAAUGUAAUAACAGAGG---------------------------------------------------------------------------------------------      8        21
-------------------------------------------------------------------------------------------------UGUUAUUACAUUGGGGAUAUC-----------      8        21
----------------------------------------------------------------------------------------------------UAUUACAUUGGGGAUAUCAACC-------      5        22
------------------------------------------------------------------------------------------------------UUACAUUGGGGAUAUCAA---------      2        18
------------------------------------------------------------------------------------------------------UUACAUUGGGGAUAUCAAC--------      3        19
------------------------------------------------------------------------------------------------------UUACAUUGGGGAUAUCAACCG------      3        21
---UGAUCGUUGAUGUCGCCAAUGUA-------------------------------------------------------------------------------------------------------      3        23
----GAUCGUUGAUGUCGCCAAUG---------------------------------------------------------------------------------------------------------      7        20
------UCGUUGAUGUCGCCAAUGUAAU-----------------------------------------------------------------------------------------------------      1        22
-----------------------------------------------------------------------------------------------UCUGUUAUUACAUUGGGGAUA-------------      5        21
```

#### *D. firmibasis* dfi-mir-1195 continued

```
(((.(((.(((((((((.(((((((((((((((((((..(((.(((((..((((((((....))))))))...........))))).)))..))))))))))))))))))).))))))))).))).)))   -69.84 kcal/mol
AGGUGAUCGUuGAUGUCGCCAAUGUAAUAAcAGAGGUUAAUUAUAUUAAAUAAUAUUUCAAUAAAUAUUAACAUUAACUUCUAAUACAAUAUACCUCUGUuAUUACAUUGGGGAUAUCAAcCGUCCCCU   count   length
-----------------------------------------------------------------------------------------------------AUUACAUUGGGGAUAUCAACCGU-----   1       23
-----------------------------------------------------------------------------------------------------AUUACAUUGGGGAUAUCA----------   2       18
AGGUGAUCGUUGAUGUCGC--------------------------------------------------------------------------------------------------------------   1       19
--GUGAUCGUUGAUGUCGCCAAUG---------------------------------------------------------------------------------------------------------   1       22
-----AUCGUUGAUGUCGCCAAUGU--------------------------------------------------------------------------------------------------------   2       20
------UCGUUGAUGUCGCCAAUGUAA------------------------------------------------------------------------------------------------------   3       21
-------CGUUGAUGUCGCCAAUGUAAU-----------------------------------------------------------------------------------------------------   2       21
--------GUUGAUGUCGCCAAUGUAAU-----------------------------------------------------------------------------------------------------   1       20
--------GUUGAUGUCGCCAAUGUAAUA----------------------------------------------------------------------------------------------------   1       21
---------UUGAUGUCGCCAAUGUAA------------------------------------------------------------------------------------------------------   1       18
------------AUGUCGCCAAUGUAAUAA---------------------------------------------------------------------------------------------------   1       18
-------------UGUCGCCAAUGUAAUAACAGA-----------------------------------------------------------------------------------------------   4       21
--------------GUCGCCAAUGUAAUAACAGA-----------------------------------------------------------------------------------------------   1       20
---------------UCGCCAAUGUAAUAACAGAG----------------------------------------------------------------------------------------------   1       20
--------------------------------------------------------------------------------------CAAUAUACCUCUGUUAUUACAU---------------------   1       22
-----------------------------------------------------------------------------------------UAUACCUCUGUUAUUACAUUG-------------------   2       21
------------------------------------------------------------------------------------------AUACCUCUGUUAUUACAUUG-------------------   1       20
-----------------------------------------------------------------------------------------------UCUGUUAUUACAUUGGGGA---------------   1       19
-----------------------------------------------------------------------------------------------UCUGUUAUUACAUUGGGGAUAU------------   1       22
----------------------------------------------------------------------------------------------------UAUUACAUUGGGGAUAUCAACCGUC----   1       25
------------------------------------------------------------------------------------------------------UUACAUUGGGGAUAUCAACC-------   1       20
```

```
   U   C          G                   UA    A      AA                  C
AGG GAU GUuGAUGUC CCAAUGUAAUAAcAGAGGU  AUU UAUUA   ---------UAAUAUUU A
UCC CUG cAACUAUAG GGUUACAUUAuUGUCUCCA  UAA AUAAU            AUUAUAAA A
   C   C          G                   UA    C      CUUCAAUUACA         U
```

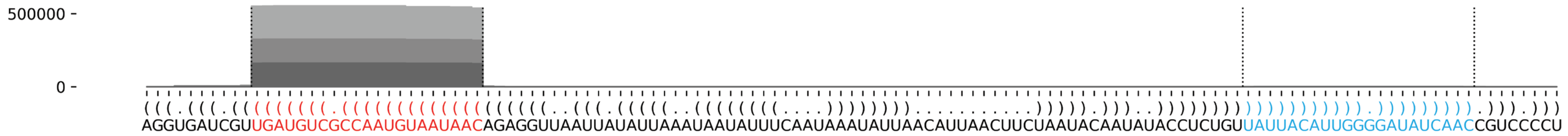

#### *D. firmibasis* dfi-mir-1196

```
(((((((((((((((((((((.((((((((((((((((((((((((...((((........)))).))))).....))))))))))))))))))).)))))))))))))))))))))   -75.37 kcal/mol
AAAAAAAACAuAGUGAAGAUGGGUAGAAGAuAAAAGCUGGUUUUGAAAAUUCAGAGAAUGGUGGGGUUAGAAAAUUACCAGCUUUUAUcUUCUACACAUCUUCACUAUgUUUUUUUU   count    length
Libraries combined         Precision: [ total = 76.8% | 5p-arm = 73.1% | 3p-arm = 79.1% ]
----------UAGUGAAGAUGGGUAGAAGAU--------------------------------------------------------------------------------------   18       21
----------------------------------------------------------------------------------------CUUCUACACAUCUUCACUAUG--------   34       21
--------------------------------------------------------------------------------------AUCUUCUACACAUCUUCACUA----------   2        21
-----------------------------------------------------------------------------------------UUCUACACAUCUUCACUAUGU-------   3        21
----------UAGUGAAGAUGGGUAGAAGAUA-------------------------------------------------------------------------------------   1        22
-----------------GAUGGGUAGAAGAUAAAAGCU-------------------------------------------------------------------------------   3        21
------------------AUGGGUAGAAGAUAAAAGCU-------------------------------------------------------------------------------   1        20
-----------AGUGAAGAUGGGUAGAAGAU--------------------------------------------------------------------------------------   1        20
---------------AGAUGGGUAGAAGAUAAAAGC---------------------------------------------------------------------------------   2        21
------------------------------------------------------------------------AAUUACCAGCUUUUAUCUUCUA-----------------------   1        22
-------------------------------------------------------------------------------------UAUCUUCUACACAUCUUCACU-----------   2        21
---------------------------------------------------------------------------------------UCUUCUACACAUCUUCACUAUG--------   1        22
Library = 1        Precision: [ total = 64.7% | 5p-arm = 75.0% | 3p-arm = 55.6% ]
----------UAGUGAAGAUGGGUAGAAGAU--------------------------------------------------------------------------------------   5        21
----------------------------------------------------------------------------------------CUUCUACACAUCUUCACUAUG--------   5        21
--------------------------------------------------------------------------------------AUCUUCUACACAUCUUCACUA----------   2        21
-----------------------------------------------------------------------------------------UUCUACACAUCUUCACUAUGU-------   2        21
----------UAGUGAAGAUGGGUAGAAGAUA-------------------------------------------------------------------------------------   1        22
-----------------GAUGGGUAGAAGAUAAAAGCU-------------------------------------------------------------------------------   1        21
------------------AUGGGUAGAAGAUAAAAGCU-------------------------------------------------------------------------------   1        20
Library = 2        Precision: [ total = 87.5% | 5p-arm = 71.4% | 3p-arm = 100.0% ]
----------------------------------------------------------------------------------------CUUCUACACAUCUUCACUAUG--------   9        21
----------UAGUGAAGAUGGGUAGAAGAU--------------------------------------------------------------------------------------   5        21
---------------AGAUGGGUAGAAGAUAAAAGC---------------------------------------------------------------------------------   1        21
-----------------GAUGGGUAGAAGAUAAAAGCU-------------------------------------------------------------------------------   1        21
Library = 3        Precision: [ total = 77.8% | 5p-arm = 72.7% | 3p-arm = 80.0% ]
----------------------------------------------------------------------------------------CUUCUACACAUCUUCACUAUG--------   20       21
----------UAGUGAAGAUGGGUAGAAGAU--------------------------------------------------------------------------------------   8        21
-------------------------------------------------------------------------------------UAUCUUCUACACAUCUUCACU-----------   2        21
-----------AGUGAAGAUGGGUAGAAGAU--------------------------------------------------------------------------------------   1        20
---------------AGAUGGGUAGAAGAUAAAAGC---------------------------------------------------------------------------------   1        21
-----------------GAUGGGUAGAAGAUAAAAGCU-------------------------------------------------------------------------------   1        21
------------------------------------------------------------------------AAUUACCAGCUUUUAUCUUCUA-----------------------   1        22
---------------------------------------------------------------------------------------UCUUCUACACAUCUUCACUAUG--------   1        22
-----------------------------------------------------------------------------------------UUCUACACAUCUUCACUAUGU-------   1        21
```

```
                     G                             AAA    GAG
AAAAAAAACAuAGUGAAGAUG GUAGAAGAuAAAAGCUGGU-----UUUGA   UUCA   A
UUUUUUUUgUAUCACUUCUAC CAUCUUcUAUUUUCGACCA     AGAUU --GGGU   A
                     A                   UUAAA     G      GGU
```

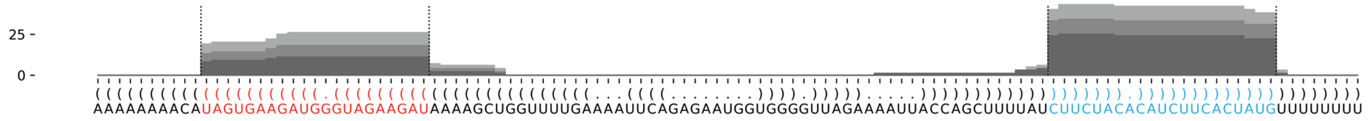

#### *D. lacteum* dla-mir-1197

```
(((((((((((((((((((..(((((((((((((((((((((((.(((((..(((....((((((((........))))).))....)))..))))).))))))...)))))))))))))))))))))))))))))))))))    -93.54 kcal/mol
AUGAGAUGGUcUGAGAUAUAUGUGAGUAAUGAuAGGUAUCAUUCCUGCUUACUAGAAUUCUUAUUAAAUACAAAAUAAUGAUAGAUGGGUAGCAGUAAUGAUUGAACCUAUcAUUACUCAUAUAUCUCAGAcCAUCUCAU    count    length
Libraries combined          Precision: [ total = 94.3% | 5p-arm = 83.3% | 3p-arm = 96.6% ]
--------------------------------------------------------------------------------------------------------------CAUUACUCAUAUAUCUCAGAC--------    21    21
----------CUGAGAUAUAUGUGAGUAAUGA----------------------------------------------------------------------------------------------------------    2     22
---------UCUGAGAUAUAUGUGAGUAAUGAU---------------------------------------------------------------------------------------------------------    1     24
----------CUGAGAUAUAUGUGAGUAAUGAU---------------------------------------------------------------------------------------------------------    2     23
---------------------------------------------------------------------------------------------------------------AUUACUCAUAUAUCUCAGACC-------    1     21
----------CUGAGAUAUAUGUGAGUAAUG-----------------------------------------------------------------------------------------------------------    1     21
--------------------------------------------------------------------------------------------------------------CAUUACUCAUAUAUCUCAGACC-------    7     22
Library = 1       Precision: [ total = 80.0% | 5p-arm = 75.0% | 3p-arm = 83.3% ]
--------------------------------------------------------------------------------------------------------------CAUUACUCAUAUAUCUCAGAC--------    5     21
----------CUGAGAUAUAUGUGAGUAAUGA----------------------------------------------------------------------------------------------------------    2     22
---------UCUGAGAUAUAUGUGAGUAAUGAU---------------------------------------------------------------------------------------------------------    1     24
----------CUGAGAUAUAUGUGAGUAAUGAU---------------------------------------------------------------------------------------------------------    1     23
---------------------------------------------------------------------------------------------------------------AUUACUCAUAUAUCUCAGACC-------    1     21
Library = 2       Precision: [ total = 100.0% | 5p-arm = 100.0% | 3p-arm = 100.0% ]
--------------------------------------------------------------------------------------------------------------CAUUACUCAUAUAUCUCAGAC--------    9     21
--------------------------------------------------------------------------------------------------------------CAUUACUCAUAUAUCUCAGACC-------    5     22
----------CUGAGAUAUAUGUGAGUAAUGAU---------------------------------------------------------------------------------------------------------    1     23
Library = 3       Precision: [ total = 100.0% | 5p-arm = 100.0% | 3p-arm = 100.0% ]
--------------------------------------------------------------------------------------------------------------CAUUACUCAUAUAUCUCAGAC--------    7     21
--------------------------------------------------------------------------------------------------------------CAUUACUCAUAUAUCUCAGACC-------    2     22
----------CUGAGAUAUAUGUGAGUAAUG-----------------------------------------------------------------------------------------------------------    1     21
```

```
                   AU                           C     UA   GAAU        AAA
AUGAGAUGGUcUGAGAUAU  GUGAGUAAUGAuAGGU---AUCAUU CUGCU  CUA    UC-UUAUU   U
UACUCUACcAGACUCUAUA--UACUCAUUAcUAUCCA   UAGUAA GACGA  GGU    AG AAUAA   A
                                     AGU      U     UG   AGAU  U     AAC
```

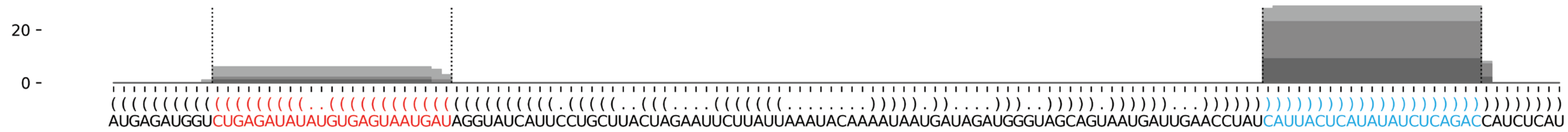


#### *D. lacteum* dla-mir-1198

```
(((((((((((((((.((.((((((((..(((..(((((..((((((....))))))...)))))..)))..))))))))..)).))))))))))))))) -52.92 kcal/mol
AAGAAUCAAAgACAAUUUGUGACCAACACAcAAUUGAUACAAUAUAUUUCUAAAUAUACAGGUAUCUUUUGAaUGUUGGUCUUCAUGUUGUUUuUGAUUUUU count    length
Libraries combined          Precision: [ total = 99.8% | 5p-arm = 90.2% | 3p-arm = 99.9% ]
------------------------------------------------------------------------AUGUUGGUCUUCAUGUUGUUUU-------- 98216    22
------------------------------------------------------------------------AUGUUGGUCUUCAUGUUGUUUUU------- 12225    23
------------------------------------------------------------------------AUGUUGGUCUUCAUGUUGUUU--------- 12730    21
------------------------------------------------------------------------AUGUUGGUCUUCAUGUUGUU---------- 2703     20
------------------------------------------------------------------------AUGUUGGUCUUCAUGUUGU----------- 484      19
------------------------------------------------------------------------AUGUUGGUCUUCAUGUUG------------ 284      18
----------GACAAUUUGUGACCAACACA------------------------------------------------------------------------ 128      20
----------GACAAUUUGUGACCAACACAC----------------------------------------------------------------------- 84       21
-------------------------------------------------------------------------UGUUGGUCUUCAUGUUGUUUUU------- 56       22
-------------------------------------------------------------------------UGUUGGUCUUCAUGUUGUUUU-------- 58       21
----------GACAAUUUGUGACCAACAC------------------------------------------------------------------------- 52       19
---------AGACAAUUUGUGACCAACACA------------------------------------------------------------------------ 24       21
-------------------------------------------------------------------------UGUUGGUCUUCAUGUUGUUU--------- 9        20
----------GACAAUUUGUGACCAACA-------------------------------------------------------------------------- 10       18
--------------------------------------------------------------UAUCUUUUGAAUGUUGGUCU-------------------- 6        20
```

#### *D. lacteum* dla-mir-1198 continued

```
((((((((((((((((.((.((((((((..(((..(((((..((((((....))))))...)))))..)))..))))))))..)).)))))))))))))))) -52.92 kcal/mol
AAGAAUCAAAgACAAUUUGUGACCAACACAcAAUUGAUACAAUAUAUUUCUAAAUAUACAGGUAUCUUUUGAaUGUUGGUCUUCAUGUUGUUUuUGAUUUUU count    length
--------------------------------------------------------------UAUCUUUUGAAUGUUGGUCUU------------------- 6        21
-----------------------------------------------------------------------AAUGUUGGUCUUCAUGUUGUUUU-------- 5        23
------------------------------------------------------------------------AUGUUGGUCUUCAUGUUGUUUUUG------ 5        24
--------------------------------------------------------------------------GUUGGUCUUCAUGUUGUUUU-------- 10       20
---------AGACAAUUUGUGACCAACACAC----------------------------------------------------------------------- 1        22
----------GACAAUUUGUGACCAACACACA---------------------------------------------------------------------- 1        22
--------------------GACCAACACACAAUUGAUACA------------------------------------------------------------- 3        21
-----------------------------------------------------------------------AAUGUUGGUCUUCAUGUUGUUU--------- 2        22
---------------------------------------------------------------------------UUGGUCUUCAUGUUGUUUU-------- 4        19
-----------------------------------------------------------------------------GGUCUUCAUGUUGUUUUU------- 1        18
---------AGACAAUUUGUGACCAACAC------------------------------------------------------------------------- 1        20
---------AGACAAUUUGUGACCAACA-------------------------------------------------------------------------- 1        19
--------------------------------------------------------------UAUCUUUUGAAUGUUGGUCUUC------------------ 1        22
--------------------------------------------------------------UAUCUUUUGAAUGUUGGUC--------------------- 1        19
-----------------------------------------------------------------------AAUGUUGGUCUUCAUGUUGUUUUU------- 2        24
------------------------------------------------------------------------AUGUUGGUCUUCAUGUUGUUUUUGAUU--- 1        27
-------------------------------------------------------------------------UGUUGGUCUUCAUGUUGUU---------- 2        19
---------------------------------------------------------------------------UUGGUCUUCAUGUUGUUUUUG------ 1        21
Library = 1       Precision: [ total = 99.9% | 5p-arm = 89.1% | 3p-arm = 99.9% ]
------------------------------------------------------------------------AUGUUGGUCUUCAUGUUGUUUU-------- 45957    22
------------------------------------------------------------------------AUGUUGGUCUUCAUGUUGUUUUU------- 6198     23
------------------------------------------------------------------------AUGUUGGUCUUCAUGUUGUUU--------- 6126     21
------------------------------------------------------------------------AUGUUGGUCUUCAUGUUGUU---------- 1191     20
------------------------------------------------------------------------AUGUUGGUCUUCAUGUUGU----------- 233      19
------------------------------------------------------------------------AUGUUGGUCUUCAUGUUG------------ 124      18
----------GACAAUUUGUGACCAACACA------------------------------------------------------------------------ 62       20
----------GACAAUUUGUGACCAACACAC----------------------------------------------------------------------- 44       21
-------------------------------------------------------------------------UGUUGGUCUUCAUGUUGUUUUU------- 26       22
-------------------------------------------------------------------------UGUUGGUCUUCAUGUUGUUUU-------- 23       21
----------GACAAUUUGUGACCAACAC------------------------------------------------------------------------- 21       19
---------AGACAAUUUGUGACCAACACA------------------------------------------------------------------------ 14       21
-------------------------------------------------------------------------UGUUGGUCUUCAUGUUGUUU--------- 7        20
----------GACAAUUUGUGACCAACA-------------------------------------------------------------------------- 3        18
--------------------------------------------------------------UAUCUUUUGAAUGUUGGUCU-------------------- 3        20
--------------------------------------------------------------UAUCUUUUGAAUGUUGGUCUU------------------- 2        21
-----------------------------------------------------------------------AAUGUUGGUCUUCAUGUUGUUUU-------- 2        23
------------------------------------------------------------------------AUGUUGGUCUUCAUGUUGUUUUUG------ 2        24
--------------------------------------------------------------------------GUUGGUCUUCAUGUUGUUUU-------- 2        20
---------AGACAAUUUGUGACCAACACAC----------------------------------------------------------------------- 1        22
----------GACAAUUUGUGACCAACACACA---------------------------------------------------------------------- 1        22
--------------------GACCAACACACAAUUGAUACA------------------------------------------------------------- 1        21
-----------------------------------------------------------------------AAUGUUGGUCUUCAUGUUGUUU--------- 1        22
---------------------------------------------------------------------------UUGGUCUUCAUGUUGUUUU-------- 1        19
-----------------------------------------------------------------------------GGUCUUCAUGUUGUUUUU------- 1        18
Library = 2       Precision: [ total = 99.8% | 5p-arm = 89.4% | 3p-arm = 99.9% ]
------------------------------------------------------------------------AUGUUGGUCUUCAUGUUGUUUU-------- 28380    22
------------------------------------------------------------------------AUGUUGGUCUUCAUGUUGUUU--------- 3721     21
------------------------------------------------------------------------AUGUUGGUCUUCAUGUUGUUUUU------- 3480     23
------------------------------------------------------------------------AUGUUGGUCUUCAUGUUGUU---------- 802      20
------------------------------------------------------------------------AUGUUGGUCUUCAUGUUGU----------- 130      19
------------------------------------------------------------------------AUGUUGGUCUUCAUGUUG------------ 86       18
----------GACAAUUUGUGACCAACACA------------------------------------------------------------------------ 34       20
----------GACAAUUUGUGACCAACACAC----------------------------------------------------------------------- 22       21
-------------------------------------------------------------------------UGUUGGUCUUCAUGUUGUUUU-------- 19       21
```

#### *D. lacteum* dla-mir-1198 continued

```
((((((((((((((((.((.((((((((..(((..(((((..((((((....))))))...)))))..)))..))))))))..)).)))))))))))))))) -52.92 kcal/mol
AAGAAUCAAAgACAAUUUGUGACCAACACAcAAUUGAUACAAUAUAUUUCUAAAUAUACAGGUAUCUUUUGAaUGUUGGUCUUCAUGUUGUUUuUGAUUUUU count    length
-------------------------------------------------------------------------UGUUGGUCUUCAUGUUGUUUUU------- 18       22
----------GACAAUUUGUGACCAACAC------------------------------------------------------------------------- 15       19
---------AGACAAUUUGUGACCAACACA------------------------------------------------------------------------ 7        21
----------GACAAUUUGUGACCAACA-------------------------------------------------------------------------- 5        18
--------------------------------------------------------------UAUCUUUUGAAUGUUGGUCUU------------------- 4        21
--------------------------------------------------------------------------GUUGGUCUUCAUGUUGUUUU-------- 4        20
------------------------------------------------------------------------AUGUUGGUCUUCAUGUUGUUUUUG------ 3        24
-----------------------------------------------------------------------AAUGUUGGUCUUCAUGUUGUUUUU------- 2        24
---------AGACAAUUUGUGACCAACAC------------------------------------------------------------------------- 1        20
--------------------GACCAACACACAAUUGAUACA------------------------------------------------------------- 1        21
--------------------------------------------------------------UAUCUUUUGAAUGUUGGUCU-------------------- 1        20
--------------------------------------------------------------UAUCUUUUGAAUGUUGGUCUUC------------------ 1        22
--------------------------------------------------------------UAUCUUUUGAAUGUUGGUC--------------------- 1        19
-----------------------------------------------------------------------AAUGUUGGUCUUCAUGUUGUUUU-------- 1        23
-------------------------------------------------------------------------UGUUGGUCUUCAUGUUGUUU--------- 1        20
---------------------------------------------------------------------------UUGGUCUUCAUGUUGUUUU-------- 1        19
Library = 3       Precision: [ total = 99.8% | 5p-arm = 93.2% | 3p-arm = 99.9% ]
------------------------------------------------------------------------AUGUUGGUCUUCAUGUUGUUUU-------- 23879    22
------------------------------------------------------------------------AUGUUGGUCUUCAUGUUGUUU--------- 2883     21
------------------------------------------------------------------------AUGUUGGUCUUCAUGUUGUUUUU------- 2547     23
------------------------------------------------------------------------AUGUUGGUCUUCAUGUUGUU---------- 710      20
------------------------------------------------------------------------AUGUUGGUCUUCAUGUUGU----------- 121      19
------------------------------------------------------------------------AUGUUGGUCUUCAUGUUG------------ 74       18
----------GACAAUUUGUGACCAACACA------------------------------------------------------------------------ 32       20
----------GACAAUUUGUGACCAACACAC----------------------------------------------------------------------- 18       21
----------GACAAUUUGUGACCAACAC------------------------------------------------------------------------- 16       19
-------------------------------------------------------------------------UGUUGGUCUUCAUGUUGUUUU-------- 16       21
-------------------------------------------------------------------------UGUUGGUCUUCAUGUUGUUUUU------- 12       22
--------------------------------------------------------------------------GUUGGUCUUCAUGUUGUUUU-------- 4        20
---------AGACAAUUUGUGACCAACACA------------------------------------------------------------------------ 3        21
----------GACAAUUUGUGACCAACA-------------------------------------------------------------------------- 2        18
--------------------------------------------------------------UAUCUUUUGAAUGUUGGUCU-------------------- 2        20
-----------------------------------------------------------------------AAUGUUGGUCUUCAUGUUGUUUU-------- 2        23
-------------------------------------------------------------------------UGUUGGUCUUCAUGUUGUU---------- 2        19
---------------------------------------------------------------------------UUGGUCUUCAUGUUGUUUU-------- 2        19
---------AGACAAUUUGUGACCAACA-------------------------------------------------------------------------- 1        19
--------------------GACCAACACACAAUUGAUACA------------------------------------------------------------- 1        21
-----------------------------------------------------------------------AAUGUUGGUCUUCAUGUUGUUU--------- 1        22
------------------------------------------------------------------------AUGUUGGUCUUCAUGUUGUUUUUGAUU--- 1        27
-------------------------------------------------------------------------UGUUGGUCUUCAUGUUGUUU--------- 1        20
---------------------------------------------------------------------------UUGGUCUUCAUGUUGUUUUUG------ 1        21
```

```
               U  U          CA    UU     AA       U
AAGAAUCAAAgACAAU UG -GACCAACA  cAA   GAUAC  -UAUAUU C
UUUUUAGUuUUUGUUG AC  CUGGUUGU  GUU   CUAUG   AUAUAA U
               U  UU         aA    UU     GAC      A
```

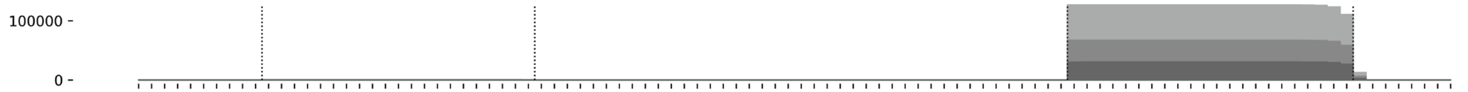


```
((((((((((((((((.((.((((((((..(((..(((((..((((((....))))))...)))))..)))..))))))))..)).))))))))))))))))
AAGAAUCAAAGACAAUUUGUGACCAACACACAAUUGAUACAAUAUAUUUCUAAAUAUACAGGUAUCUUUUGAAUGUUGGUCUUCAUGUUGUUUUUGAUUUUU
```

#### *P. pallidum* ppa-mir-1199

```
.......((((((((((((((((((((((((((((((((((((((((.........)))))))))))))))))))))))))))))))))))))))).......          -65.05 kcal/mol
UAAUUUUUUGaACAGACAUCUUGAAAAAAUAaAGUUAAAUUAAAGUUCAAAAAAUUAACUUUAAUUUAACUUUaUUUUUUCAAGAUGUCUGUUCaACUUAGGU          count    length
Libraries combined          Precision: [ total = 83.6% | 5p-arm = 72.6% | 3p-arm = 84.5% ]
-------------------------------------------------------------------------AUUUUUUCAAGAUGUCUGUUC---------          2199     21
----------AACAGACAUCUUGAAAAAAUA------------------------------------------------------------------------          153      21
--------------------------------------------------------------------------UUUUUUCAAGAUGUCUGUUCA--------          176      21
-------------------------------------------------------------------------AUUUUUUCAAGAUGUCUGUU----------          145      20
---------------------------------------------------------------------------------AAGAUGUCUGUUCAACUUAGG-          61       21
----------------------------------------------------------------------------------AGAUGUCUGUUCAACUUAGGU          65       21
--------------------------------------------------------------------------UUUUUUCAAGAUGUCUGUUC---------          63       20
--------UGAACAGACAUCUUGAAAAAA--------------------------------------------------------------------------          25       21
----------AACAGACAUCUUGAAAAAAU-------------------------------------------------------------------------          10       20
-------------------------------------------------------------------------AUUUUUUCAAGAUGUCUGUUCA--------          18       22
-------UUGAACAGACAUCUUGAAAAA---------------------------------------------------------------------------          9        21
-------------------------------------------------------------------AACUUUAUUUUUUCAAGAUGUC--------------          7        22
----------------------------------------------------------------------------------AGAUGUCUGUUCAACUUAGG-          4        20
-----------ACAGACAUCUUGAAAAAAUA------------------------------------------------------------------------          8        20
------------------------------------------------------------------UAACUUUAUUUUUUCAAGAUG----------------          3        21
----------------------------------------------------------------------UUUAUUUUUUCAAGAUGUCUGUU----------          5        23
------------------------------------------------------------------------UAUUUUUUCAAGAUGUCUGUUC---------          11       22
-------------------------------------------------------------------------AUUUUUUCAAGAUGUCUGU-----------          5        19
-----------------------------------------------------------------------------UUUCAAGAUGUCUGUUCAACU-----          4        21
-----------ACAGACAUCUUGAAAAAAUAA-----------------------------------------------------------------------          5        21
--------------------------------------------------------------------------UUUUUUCAAGAUGUCUGUU----------          9        19
-------UUGAACAGACAUCUUGAAAAAAU-------------------------------------------------------------------------          1        23
--------UGAACAGACAUCUUGAAAAA---------------------------------------------------------------------------          5        20
-------------------------------------------------------------------AACUUUAUUUUUUCAAGAUGUCUGUU----------          2        26
-------------------------------------------------------------------------AUUUUUUCAAGAUGUCUGUUCAA-------          1        23
---------------------------------------------------------------------------------AAGAUGUCUGUUCAACUUAGGU          3        22
-------UUGAACAGACAUCUUGAAAAAA--------------------------------------------------------------------------          3        22
--------UGAACAGACAUCUUGAAAAAAU-------------------------------------------------------------------------          2        22
---------GAACAGACAUCUUGAAAAAAUA------------------------------------------------------------------------          1        22
----------AACAGACAUCUUGAAAAAAUAA-----------------------------------------------------------------------          1        22
-----------ACAGACAUCUUGAAAAAA--------------------------------------------------------------------------          1        18
-----------ACAGACAUCUUGAAAAAAUAAA----------------------------------------------------------------------          1        22
-----------ACAGACAUCUUGAAAAAAU-------------------------------------------------------------------------          1        19
------------------------------------------------------------------UAACUUUAUUUUUUCAAGAUGU---------------          1        22
-------------------------------------------------------------------AACUUUAUUUUUUCAAGAUGUCUGU-----------          2        25
-------------------------------------------------------------------AACUUUAUUUUUUCAAGAUGUCUG------------          3        24
----------------------------------------------------------------------UUUAUUUUUUCAAGAUGUCUGU-----------          1        22
-----------------------------------------------------------------------UUAUUUUUUCAAGAUGUCUGU-----------          1        21
-----------------------------------------------------------------------UUAUUUUUUCAAGAUGUCUGUU----------          2        22
------------------------------------------------------------------------UAUUUUUUCAAGAUGUCUGUU----------          6        21
------------------------------------------------------------------------UAUUUUUUCAAGAUGUCUGUUCA--------          1        23
-------------------------------------------------------------------------AUUUUUUCAAGAUGUCUG------------          2        18
--------------------------------------------------------------------------UUUUUUCAAGAUGUCUGUUCAA-------          1        22
---------------------------------------------------------------------------UUUUUCAAGAUGUCUGUUCAA-------          1        21
-------------------------------------------------------------------------------UCAAGAUGUCUGUUCAACUUAGGU          1        24
--------------------------------------------------------------------------------CAAGAUGUCUGUUCAACUUAGG-          1        22
---------------------------------------------------------------------------------AAGAUGUCUGUUCAACUUAG--          1        20
Library = 1        Precision: [ total = 85.2% | 5p-arm = 77.9% | 3p-arm = 85.8% ]
-------------------------------------------------------------------------AUUUUUUCAAGAUGUCUGUUC---------          1005     21
----------AACAGACAUCUUGAAAAAAUA------------------------------------------------------------------------          74       21
--------------------------------------------------------------------------UUUUUUCAAGAUGUCUGUUCA--------          63       21
-------------------------------------------------------------------------AUUUUUUCAAGAUGUCUGUU----------          57       20
---------------------------------------------------------------------------------AAGAUGUCUGUUCAACUUAGG-          35       21
```

#### *P. pallidum* ppa-mir-1199 continued

```
.......((((((((((((((((((((((((((((((((((((((.............)))))))))))))))))))))))))))))))))))))).......   -65.05 kcal/mol
UAAUUUUUUGaACAGACAUCUUGAAAAAAUAaAGUUAAAUUAAAGUUCAAAAAAUUAACUUUAAUUUAACUUUaUUUUUUCAAGAUGUCUGUUCaACUUAGGU   count   length
----------------------------------------------------------------------------------AGAUGUCUGUUCAACUUAGGU   29      21
--------------------------------------------------------------------------UUUUUUCAAGAUGUCUGUUC---------   27      20
--------UGAACAGACAUCUUGAAAAAA--------------------------------------------------------------------------   11      21
----------AACAGACAUCUUGAAAAAAU-------------------------------------------------------------------------   7       20
-------------------------------------------------------------------------AUUUUUUCAAGAUGUCUGUUCA--------   6       22
-------UUGAACAGACAUCUUGAAAAA---------------------------------------------------------------------------   5       21
-------------------------------------------------------------------AACUUUAUUUUUUCAAGAUGUC--------------   4       22
----------------------------------------------------------------------------------AGAUGUCUGUUCAACUUAGG-   4       20
-----------ACAGACAUCUUGAAAAAAUA------------------------------------------------------------------------   3       20
------------------------------------------------------------------UAACUUUAUUUUUUCAAGAUG----------------   3       21
----------------------------------------------------------------------UUUAUUUUUUCAAGAUGUCUGUU----------   3       23
------------------------------------------------------------------------UAUUUUUUCAAGAUGUCUGUUC---------   3       22
-------------------------------------------------------------------------AUUUUUUCAAGAUGUCUGU-----------   3       19
-----------------------------------------------------------------------------UUUCAAGAUGUCUGUUCAACU-----   3       21
-----------ACAGACAUCUUGAAAAAAUAA-----------------------------------------------------------------------   2       21
--------------------------------------------------------------------------UUUUUUCAAGAUGUCUGUU----------   2       19
-------UUGAACAGACAUCUUGAAAAAAU-------------------------------------------------------------------------   1       23
--------UGAACAGACAUCUUGAAAAA---------------------------------------------------------------------------   1       20
-------------------------------------------------------------------AACUUUAUUUUUUCAAGAUGUCUGUU----------   1       26
-------------------------------------------------------------------------AUUUUUUCAAGAUGUCUGUUCAA-------   1       23
---------------------------------------------------------------------------------AAGAUGUCUGUUCAACUUAGGU   1       22
Library = 2        Precision: [ total = 78.7% | 5p-arm = 63.4% | 3p-arm = 80.5% ]
-------------------------------------------------------------------------AUUUUUUCAAGAUGUCUGUUC---------   526     21
--------------------------------------------------------------------------UUUUUUCAAGAUGUCUGUUCA--------   73      21
----------AACAGACAUCUUGAAAAAAUA------------------------------------------------------------------------   49      21
-------------------------------------------------------------------------AUUUUUUCAAGAUGUCUGUU----------   36      20
---------------------------------------------------------------------------------AAGAUGUCUGUUCAACUUAGG-   17      21
--------UGAACAGACAUCUUGAAAAAA--------------------------------------------------------------------------   11      21
-------------------------------------------------------------------------AUUUUUUCAAGAUGUCUGUUCA--------   10      22
--------------------------------------------------------------------------UUUUUUCAAGAUGUCUGUUC---------   9       20
----------------------------------------------------------------------------------AGAUGUCUGUUCAACUUAGGU   7       21
------------------------------------------------------------------------UAUUUUUUCAAGAUGUCUGUUC---------   5       22
------------------------------------------------------------------------UAUUUUUUCAAGAUGUCUGUU----------   5       21
-----------ACAGACAUCUUGAAAAAAUA------------------------------------------------------------------------   4       20
--------------------------------------------------------------------------UUUUUUCAAGAUGUCUGUU----------   4       19
-------UUGAACAGACAUCUUGAAAAAA--------------------------------------------------------------------------   3       22
--------UGAACAGACAUCUUGAAAAA---------------------------------------------------------------------------   3       20
-----------ACAGACAUCUUGAAAAAAUAA-----------------------------------------------------------------------   3       21
--------UGAACAGACAUCUUGAAAAAAU-------------------------------------------------------------------------   2       22
----------AACAGACAUCUUGAAAAAAU-------------------------------------------------------------------------   2       20
-------------------------------------------------------------------AACUUUAUUUUUUCAAGAUGUCUGU-----------   2       25
-------------------------------------------------------------------AACUUUAUUUUUUCAAGAUGUCUG------------   2       24
----------------------------------------------------------------------UUUAUUUUUUCAAGAUGUCUGUU----------   2       23
-----------------------------------------------------------------------UUAUUUUUUCAAGAUGUCUGUU----------   2       22
-------------------------------------------------------------------------AUUUUUUCAAGAUGUCUG------------   2       18
-------UUGAACAGACAUCUUGAAAAA---------------------------------------------------------------------------   1       21
----------AACAGACAUCUUGAAAAAAUAA-----------------------------------------------------------------------   1       22
-----------ACAGACAUCUUGAAAAAA--------------------------------------------------------------------------   1       18
-----------ACAGACAUCUUGAAAAAAUAAA----------------------------------------------------------------------   1       22
-----------ACAGACAUCUUGAAAAAAU-------------------------------------------------------------------------   1       19
------------------------------------------------------------------UAACUUUAUUUUUUCAAGAUGU---------------   1       22
-------------------------------------------------------------------AACUUUAUUUUUUCAAGAUGUC--------------   1       22
-------------------------------------------------------------------AACUUUAUUUUUUCAAGAUGUCUGUU----------   1       26
-----------------------------------------------------------------------UUAUUUUUUCAAGAUGUCUGU-----------   1       21
```

#### *P. pallidum* ppa-mir-1199 continued

```
.......((((((((((((((((((((((((((((((((((((((((.........))))))))))))))))))))))))))))))))))))))))).......     -65.05 kcal/mol
UAAUUUUUUGaACAGACAUCUUGAAAAAAUAaAGUUAAAUUAAAGUUCAAAAAAUUAACUUUAAUUUAACUUUaUUUUUUCAAGAUGUCUGUUCaACUUAGGU     count   length
------------------------------------------------------------------------UAUUUUUUCAAGAUGUCUGUUCA--------     1       23
---------------------------------------------------------------------------UUUUUCAAGAUGUCUGUUCAA-------     1       21
-----------------------------------------------------------------------------UUUCAAGAUGUCUGUUCAACU-----     1       21
-------------------------------------------------------------------------------UCAAGAUGUCUGUUCAACUUAGGU     1       24
--------------------------------------------------------------------------------CAAGAUGUCUGUUCAACUUAGG-     1       22
---------------------------------------------------------------------------------AAGAUGUCUGUUCAACUUAGGU     1       22
---------------------------------------------------------------------------------AAGAUGUCUGUUCAACUUAG--     1       20
Library = 3        Precision: [ total = 85.6% | 5p-arm = 77.5% | 3p-arm = 86.0% ]
-------------------------------------------------------------------------AUUUUUUCAAGAUGUCUGUUC---------     668     21
-------------------------------------------------------------------------AUUUUUUCAAGAUGUCUGUU----------     52      20
--------------------------------------------------------------------------UUUUUUCAAGAUGUCUGUUCA--------     40      21
----------AACAGACAUCUUGAAAAAAUA------------------------------------------------------------------------     30      21
----------------------------------------------------------------------------------AGAUGUCUGUUCAACUUAGGU     29      21
--------------------------------------------------------------------------UUUUUUCAAGAUGUCUGUUC---------     27      20
---------------------------------------------------------------------------------AAGAUGUCUGUUCAACUUAGG-     9       21
-------UUGAACAGACAUCUUGAAAAA---------------------------------------------------------------------------     3       21
--------UGAACAGACAUCUUGAAAAAA--------------------------------------------------------------------------     3       21
------------------------------------------------------------------------UAUUUUUUCAAGAUGUCUGUUC---------     3       22
--------------------------------------------------------------------------UUUUUUCAAGAUGUCUGUU----------     3       19
-------------------------------------------------------------------AACUUUAUUUUUUCAAGAUGUC--------------     2       22
-------------------------------------------------------------------------AUUUUUUCAAGAUGUCUGUUCA--------     2       22
-------------------------------------------------------------------------AUUUUUUCAAGAUGUCUGU-----------     2       19
--------UGAACAGACAUCUUGAAAAA---------------------------------------------------------------------------     1       20
---------GAACAGACAUCUUGAAAAAAUA------------------------------------------------------------------------     1       22
----------AACAGACAUCUUGAAAAAAU-------------------------------------------------------------------------     1       20
-----------ACAGACAUCUUGAAAAAAUA------------------------------------------------------------------------     1       20
-------------------------------------------------------------------AACUUUAUUUUUUCAAGAUGUCUG------------     1       24
----------------------------------------------------------------------UUUAUUUUUUCAAGAUGUCUGU-----------     1       22
------------------------------------------------------------------------UAUUUUUUCAAGAUGUCUGUU----------     1       21
--------------------------------------------------------------------------UUUUUUCAAGAUGUCUGUUCAA-------     1       22
---------------------------------------------------------------------------------AAGAUGUCUGUUCAACUUAGGU     1       22
```

```
UAAUUUU                                       CAAA
       UUGaACAGACAUCUUGAAAAAAUAaAGUUAAAUUAAAGUU    A
       AaCUUGUCUGUAGAACUUUUUUaUUUCAAUUUAAUUUCAA    /
UGGAUUC                                       UUAA
```

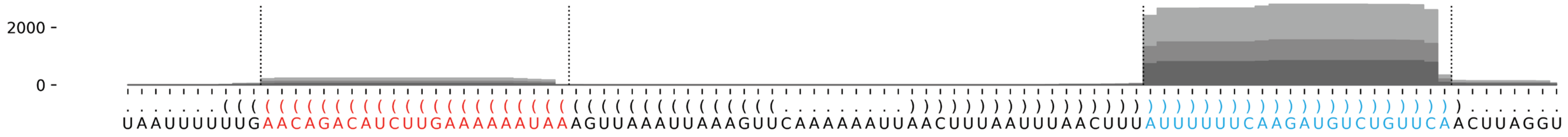

#### *P. palliddum* ppa-mir-1200

```
.....(((((((.(((((((((((((.(((((.((....(((((.((((((.....)))))).)))))...)).))))).))))))))))))).))))))).....        -35.21 kcal/mol
CUAAUUAAUUuGCUAAAAUCAAAAAUAGACaAAUUCUAACUAAGUAUUUUUAUUUAAAAUAAUUAGUAUUAUCUGuCUCUUUUUGGUUUUAAUAAaUUAUUAUU       count    length
Libraries combined         Precision: [ total = 100.0% | 5p-arm = 100.0% | 3p-arm = 100.0% ]
----------UGCUAAAAUCAAAAAUAGACA-----------------------------------------------------------------------     21       21
----------UGCUAAAAUCAAAAAUAGAC------------------------------------------------------------------------     22       20
----------------------------------------------------------------------------UCUCUUUUUGGUUUUAAUAAA--------  5        21
Library = 1        Precision: [ total = 100.0% | 5p-arm = 100.0% | 3p-arm = 100.0% ]
----------UGCUAAAAUCAAAAAUAGACA-----------------------------------------------------------------------     15       21
----------UGCUAAAAUCAAAAAUAGAC------------------------------------------------------------------------     11       20
----------------------------------------------------------------------------UCUCUUUUUGGUUUUAAUAAA--------  4        21
Library = 2        Precision: [ total = 100.0% | 5p-arm = 100.0% | 3p-arm = 0% ]
----------UGCUAAAAUCAAAAAUAGACA-----------------------------------------------------------------------     4        21
----------UGCUAAAAUCAAAAAUAGAC------------------------------------------------------------------------     3        20
Library = 3        Precision: [ total = 100.0% | 5p-arm = 100.0% | 3p-arm = 100.0% ]
----------UGCUAAAAUCAAAAAUAGAC------------------------------------------------------------------------     8        20
----------UGCUAAAAUCAAAAAUAGACA-----------------------------------------------------------------------     2        21
----------------------------------------------------------------------------UCUCUUUUUGGUUUUAAUAAA--------  1        21

CUAAU       C            U     A  UCUA     G      UA
     UAAUUuG UAAAAUCAAAAA AGACa AU    ACUAA UAUUUU  U
     AUUaAAU AUUUUGGUUUUU UCuGU UA   -UGAUU AUAAAA  /
UUAUU       A            C     C  UUA      A      UU
```

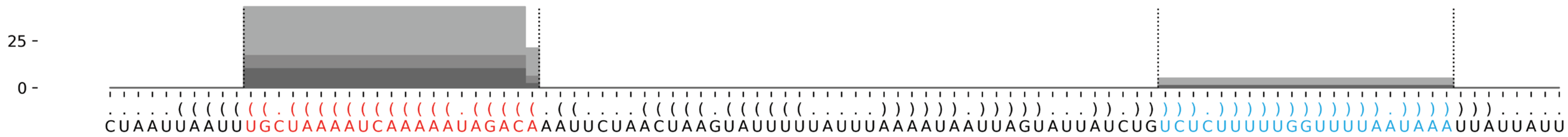

#### *P. pallidum* ppa-mir-1201

```
((.((((((((((.(((((((.((((((((((((((((..((...))..))))).(((((................))))).))))))))))).))))))).)))))))))).))   -54.86 kcal/mol
ACUCGGAUUAuCUUACACUUUCUCAUUCUAuCCGAAGCCAGUUCAACAAGCUUUGAGAUACCAAGAAUACAGAUCAUGUCUAGGAUaGAAUGGUAAAGUGUCAGAUaAUCUGUGU   count    length
Libraries combined          Precision: [ total = 99.1% | 5p-arm = 100.0% | 3p-arm = 99.0% ]
--------------------------------------------------------------------------------------AGAAUGGUAAAGUGUCAGAUA--------   479      21
--------------------------------------------------------------------------------------AGAAUGGUAAAGUGUCAGAU---------   35       20
--------------------------------------------------------------------------------------AGAAUGGUAAAGUGUCAGAUAAU------   2        23
----------UCUUACACUUUCUCAUUCUAU------------------------------------------------------------------------------------   4        21
--------------------------------------------------------------------------------------AGAAUGGUAAAGUGUCAGAUAA-------   1        22
-----------------------------------------------------------------------------------------AUGGUAAAGUGUCAGAUAAUC-----   2        21
------------------------------------------------------------------AUACAGAUCAUGUCUAGGAUA----------------------------   1        21
--------------------------------------------------------------------------------------AGAAUGGUAAAGUGUCAGA----------   2        19
---------------------------------------------------------------------------------------GAAUGGUAAAGUGUCAGAUA--------   1        20
-----------------------------------------------------------------------------------------AUGGUAAAGUGUCAGAUA--------   1        18
Library = 1        Precision: [ total = 99.6% | 5p-arm = 100.0% | 3p-arm = 99.6% ]
--------------------------------------------------------------------------------------AGAAUGGUAAAGUGUCAGAUA--------   247      21
--------------------------------------------------------------------------------------AGAAUGGUAAAGUGUCAGAU---------   21       20
--------------------------------------------------------------------------------------AGAAUGGUAAAGUGUCAGAUAAU------   2        23
----------UCUUACACUUUCUCAUUCUAU------------------------------------------------------------------------------------   1        21
--------------------------------------------------------------------------------------AGAAUGGUAAAGUGUCAGAUAA-------   1        22
-----------------------------------------------------------------------------------------AUGGUAAAGUGUCAGAUAAUC-----   1        21
Library = 2        Precision: [ total = 98.4% | 5p-arm = 100.0% | 3p-arm = 98.4% ]
--------------------------------------------------------------------------------------AGAAUGGUAAAGUGUCAGAUA--------   115      21
--------------------------------------------------------------------------------------AGAAUGGUAAAGUGUCAGAU---------   9        20
----------UCUUACACUUUCUCAUUCUAU------------------------------------------------------------------------------------   1        21
--------------------------------------------------------------------------------------AGAAUGGUAAAGUGUCAGA----------   1        19
-----------------------------------------------------------------------------------------AUGGUAAAGUGUCAGAUAAUC-----   1        21
-----------------------------------------------------------------------------------------AUGGUAAAGUGUCAGAUA--------   1        18
Library = 3        Precision: [ total = 98.4% | 5p-arm = 100.0% | 3p-arm = 98.4% ]
--------------------------------------------------------------------------------------AGAAUGGUAAAGUGUCAGAUA--------   117      21
--------------------------------------------------------------------------------------AGAAUGGUAAAGUGUCAGAU---------   5        20
----------UCUUACACUUUCUCAUUCUAU------------------------------------------------------------------------------------   2        21
------------------------------------------------------------------AUACAGAUCAUGUCUAGGAUA----------------------------   1        21
--------------------------------------------------------------------------------------AGAAUGGUAAAGUGUCAGA----------   1        19
---------------------------------------------------------------------------------------GAAUGGUAAAGUGUCAGAUA--------   1        20
```

```
                                 C-
                                U  A
                                 UA
                                 GC
                                A  A
                                C  A
                                 CG
                                 GC
                                 AU
                                 AU
  U          U       C           GUG      CCAAGAA
AC CGGAUUAuCU ACACUUU UCAUUCUAuCC   AGAUA       U
UG GUCUAaUAGA UGUGAAA GGUAAGaUAGG --UCUGU       A
  U          C       U           A        ACUAGAC
```

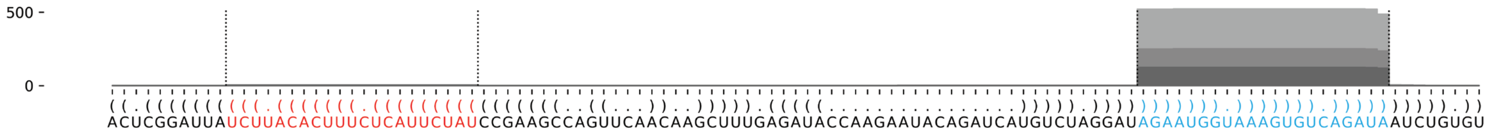

#### *P. pallidum* ppa-mir-1202

```
...(((((((.((.(((((((((.(((((.(((((((((.(((.((..((((((.((...(((.((((.(((((.....)))))..))))...))).))))))))..)).))).)..((((((((...((((..(((...)))..))))....))))))))))))).)))))).)))))))))).)).))))))))...   -101.16 kcal/mol
CCAUCGGAUGaUGCUUAACAGAUUUAAUUUcAAUUGGUACCAAGCUGACAUGUCUCUUUGUUUUGAAAGGGUAUCUUUAGACCAAAGACCACCCCUCUUCUACAAGAGAAUUGACAGGUCUGAUGGCUAUCUACAGAAUAGUCGCUAAGCUGUCUCUUAGUCGUCUGGUGUCAAGUGAaAUAGAAUCUGUUGUGCUUCaUCCGAGUA   count   length
Libraries combined      Precision: [ total = 99.2% | 5p-arm = 99.3% | 3p-arm = 99.2% ]
----------------------------------------------------------------------------------------------------------------------------------------------------------------------------------AAUAGAAUCUGUUGUGCUUCA--------   1794   21
----------------------------------------------------------------------------------------------------------------------------------------------------------------------------------AAUAGAAUCUGUUGUGCUUC---------   868   20
----------AUGCUUAACAGAUUUAAUUUC--------------------------------------------------------------------------------------------------------------------------------------------------------------------------------   534   21
----------------------------------------------------------------------------------------------------------------------------------------------------------------------------------AAUAGAAUCUGUUGUGCUUCAU-------   281   22
----------------------------------------------------------------------------------------------------------------------------------------------------------------------------------AAUAGAAUCUGUUGUGCUU----------   79   19
----------------------------------------------------------------------------------------------------------------------------------------------------------------------------------AAUAGAAUCUGUUGUGCU-----------   13   18
------------------------------------------------------------------------------------------------------------------------------------------------------UGUCUCUUAGUCGUCUGGUGUCAAGUGA-----------------------------   4   28
-------------------------------------------------------CUUUGUUUUGAAAGGGUAUCU-----------------------------------------------------------------------------------------------------------------------------------   2   21
----------------------------------------------------------------------------------------------------------------------------------------------------------------------------------AAUAGAAUCUGUUGUGCUUCAUC------   8   23
-------------------------------------UACCAAGCUGACAUGUCUCUU-----------------------------------------------------------------------------------------------------------------------------------------------------   2   21
-------------------------------------------------------------------------------------------------------AAGAGAAUUGACAGGUCUGAUG----------------------------------------------------------------------------------   1   22
-------------------------------------------------------------------------------------------------------AAGAGAAUUGACAGGUCUGAU-----------------------------------------------------------------------------------   3   21
-------------------------------------------------------------------------------------------------------------------------------------------------UAAGCUGUCUCUUAGUCGUC------------------------------------------   4   20
----------------------------------------------------------------------------------------------------------------------------------------------------------------------------------AAUAGAAUCUGUUGUGCUUCAUCCGAGUA   1   29
-----------------------------------------------------------------------------------------------------------------------------------------------------------------------------------AUAGAAUCUGUUGUGCUUCA--------   1   20
---------------------------------------------------------------------------------------------------CUACAAGAGAAUUGACAGGUC---------------------------------------------------------------------------------------   1   21
------------------------------------------------------------------------------------------------------CAAGAGAAUUGACAGGUCUGA------------------------------------------------------------------------------------   4   21
-------------------------------------------------------------------------------------------------------------------------------------------------UAAGCUGUCUCUUAGUCGU-------------------------------------------   2   19
-------------------------------------------------------------------------------------------------------------------------------------------------UAAGCUGUCUCUUAGUCGUCU-----------------------------------------   3   21
-------------------------------------------------------------------------------------------------------------------------------------------------------GUCUCUUAGUCGUCUGGUGUCAAGUGA-----------------------------   1   27
------------------------------------------------------------------------------------------------------------------------------------------------------------------------------------UAGAAUCUGUUGUGCUUCAU-------   1   20
Library = 1 Precision: [ total = 98.9% | 5p-arm = 98.0% | 3p-arm = 99.1% ]
----------------------------------------------------------------------------------------------------------------------------------------------------------------------------------AAUAGAAUCUGUUGUGCUUCA--------   484   21
----------------------------------------------------------------------------------------------------------------------------------------------------------------------------------AAUAGAAUCUGUUGUGCUUC---------   248   20
----------AUGCUUAACAGAUUUAAUUUC--------------------------------------------------------------------------------------------------------------------------------------------------------------------------------   145   21
----------------------------------------------------------------------------------------------------------------------------------------------------------------------------------AAUAGAAUCUGUUGUGCUUCAU-------   102   22
----------------------------------------------------------------------------------------------------------------------------------------------------------------------------------AAUAGAAUCUGUUGUGCUU----------   22   19
----------------------------------------------------------------------------------------------------------------------------------------------------------------------------------AAUAGAAUCUGUUGUGCU-----------   5   18
------------------------------------------------------------------------------------------------------------------------------------------------------UGUCUCUUAGUCGUCUGGUGUCAAGUGA-----------------------------   3   28
-------------------------------------------------------CUUUGUUUUGAAAGGGUAUCU-----------------------------------------------------------------------------------------------------------------------------------   2   21
----------------------------------------------------------------------------------------------------------------------------------------------------------------------------------AAUAGAAUCUGUUGUGCUUCAUC------   2   23
-------------------------------------UACCAAGCUGACAUGUCUCUU-----------------------------------------------------------------------------------------------------------------------------------------------------   1   21
-------------------------------------------------------------------------------------------------------AAGAGAAUUGACAGGUCUGAUG----------------------------------------------------------------------------------   1   22
-------------------------------------------------------------------------------------------------------AAGAGAAUUGACAGGUCUGAU-----------------------------------------------------------------------------------   1   21
-------------------------------------------------------------------------------------------------------------------------------------------------UAAGCUGUCUCUUAGUCGUC------------------------------------------   1   20
----------------------------------------------------------------------------------------------------------------------------------------------------------------------------------AAUAGAAUCUGUUGUGCUUCAUCCGAGUA   1   29
-----------------------------------------------------------------------------------------------------------------------------------------------------------------------------------AUAGAAUCUGUUGUGCUUCA--------   1   20
Library = 2 Precision: [ total = 99.1% | 5p-arm = 99.6% | 3p-arm = 99.0% ]
----------------------------------------------------------------------------------------------------------------------------------------------------------------------------------AAUAGAAUCUGUUGUGCUUCA--------   733   21
----------------------------------------------------------------------------------------------------------------------------------------------------------------------------------AAUAGAAUCUGUUGUGCUUC---------   351   20
----------AUGCUUAACAGAUUUAAUUUC--------------------------------------------------------------------------------------------------------------------------------------------------------------------------------   268   21
----------------------------------------------------------------------------------------------------------------------------------------------------------------------------------AAUAGAAUCUGUUGUGCUUCAU-------   101   22
----------------------------------------------------------------------------------------------------------------------------------------------------------------------------------AAUAGAAUCUGUUGUGCUU----------   38   19
----------------------------------------------------------------------------------------------------------------------------------------------------------------------------------AAUAGAAUCUGUUGUGCU-----------   5   18
----------------------------------------------------------------------------------------------------------------------------------------------------------------------------------AAUAGAAUCUGUUGUGCUUCAUC------   4   23
-------------------------------------------------------------------------------------------------------------------------------------------------UAAGCUGUCUCUUAGUCGUC------------------------------------------   3   20
-------------------------------------------------------------------------------------------------------AAGAGAAUUGACAGGUCUGAU-----------------------------------------------------------------------------------   2   21
-------------------------------------------------------------------------------------------------------------------------------------------------UAAGCUGUCUCUUAGUCGU-------------------------------------------   2   19
-------------------------------------------------------------------------------------------------------------------------------------------------UAAGCUGUCUCUUAGUCGUCU-----------------------------------------   2   21
-------------------------------------UACCAAGCUGACAUGUCUCUU-----------------------------------------------------------------------------------------------------------------------------------------------------   1   21
---------------------------------------------------------------------------------------------------CUACAAGAGAAUUGACAGGUC---------------------------------------------------------------------------------------   1   21
------------------------------------------------------------------------------------------------------CAAGAGAAUUGACAGGUCUGA------------------------------------------------------------------------------------   1   21
------------------------------------------------------------------------------------------------------------------------------------------------------UGUCUCUUAGUCGUCUGGUGUCAAGUGA-----------------------------   1   28
-------------------------------------------------------------------------------------------------------------------------------------------------------GUCUCUUAGUCGUCUGGUGUCAAGUGA-----------------------------   1   27
Library = 3 Precision: [ total = 99.5% | 5p-arm = 100.0% | 3p-arm = 99.5% ]
----------------------------------------------------------------------------------------------------------------------------------------------------------------------------------AAUAGAAUCUGUUGUGCUUCA--------   577   21
----------------------------------------------------------------------------------------------------------------------------------------------------------------------------------AAUAGAAUCUGUUGUGCUUC---------   269   20
----------AUGCUUAACAGAUUUAAUUUC--------------------------------------------------------------------------------------------------------------------------------------------------------------------------------   121   21
----------------------------------------------------------------------------------------------------------------------------------------------------------------------------------AAUAGAAUCUGUUGUGCUUCAU-------   78   22
----------------------------------------------------------------------------------------------------------------------------------------------------------------------------------AAUAGAAUCUGUUGUGCUU----------   19   19
------------------------------------------------------------------------------------------------------CAAGAGAAUUGACAGGUCUGA------------------------------------------------------------------------------------   3   21
----------------------------------------------------------------------------------------------------------------------------------------------------------------------------------AAUAGAAUCUGUUGUGCU-----------   3   18
```

#### *P. pallidum* ppa-mir-1202

```
...(((((((.((.(((((((((.(((((.(((((((((.(((.((..((((((.((...(((.((((.(((((.....)))))..))))...))).)))))))))..)).))).)..((((((((...((((..(((...)))..))))....))))))))))))))))).)))))).))))))))).)).))))))))...   -101.16 kcal/mol
CCAUCGGAUGaUGCUUAACAGAUUUAAUUUcAAUUGGUACCAAGCUGACAUGUCUCUUUGUUUUGAAAGGGUAUCUUUAGACCAAAGACCACCCCUCUUCUACAAGAGAAUUGACAGGUCUGAUGGCUAUCUACAGAAUAGUCGCUAAGCUGUCUCUUAGUCGUCUGGUGUCAAGUGAaAUAGAAUCUGUUGUGCUUCaUCCGAGUA   count   length
---------------------------------------------------------------------------------------------------------------------------------------------------------------------AAUAGAAUCUGUUGUGCUUCAUC------   2   23
---------------------------------------------------------------------------------------------------------------------------UAAGCUGUCUCUUAGUCGUCU--------------------------------------------   1   21
------------------------------------------------------------------------------------------------------------------------------------------------------------------------UAGAAUCUGUUGUGCUUCAU-------   1   20
```

```
CCA         U  U          A      A              G   A  UG      U  UUU   A--    A-     AG
   UCGGAUGa GC UAACAGAUUU AUUUcA UUGGUACCA----A CUG CA  UCUCUU GU   GAA   GGGU  UCUUU  A
   AGCCUUaCU CG GUUGUCUAAG UAaAGU AACUGUGGU\    U GAC GU  AGAGAA-CA   CUU   CCCA  AGAAA  /
AUG         U  U          A      G          CGCU G   A  UA        U--   CUC    CC     CC
                                            UA
                                            GU     UCU-     AA   U
                                            CGGCUA     ACAG  UAG C
                                            \-UGAU     UGUC  AUC /
                                                  UCUC     GA   G
```

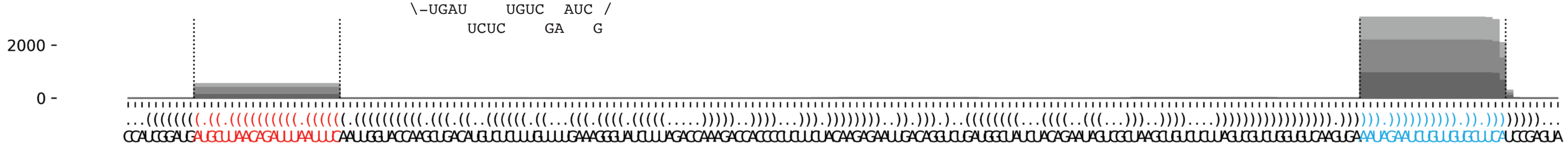

##### *A. subglobosum* asu-mir-1203

```
..((.((((((((((.((((((((((((((((.((.((((((((.(((.((....)).))).)).........)))))).)).)))))))))))))))).)))))))))).))..   -52.88 kcal/mol
AUCGUCUGUAcUUAUCUAAUACCCAAUAAAcAACAGUAAUCAGUAUCUAUCACCAGACAGAAGCAAGUCAAGUUGGUUAAUGAUGUuUAUUGGGUAUUAAAUAAGUAaUGGUUGUC   count   length
Libraries combined         Precision: [ total = 92.0% | 5p-arm = 66.7% | 3p-arm = 92.1% ]
--------------------------------------------------------------------------------------UUAUUGGGUAUUAAAUAAGUA--------   1682    21
--------------------------------------------------------------------------------------UUAUUGGGUAUUAAAUAAGU---------   388     20
--------------------------------------------------------------------------------------UUAUUGGGUAUUAAAUAAGUAU-------   172     22
------------------------------------------------------------------------------------------UGGGUAUUAAAUAAGUAUGGU----   143     21
-----------------------------------------------------------------------------------------UUGGGUAUUAAAUAAGUAUGG-----   44      21
--------------------------------------------------------------------------------------UUAUUGGGUAUUAAAUAAG----------   5       19
-------------------------------------------------------------------------------------UUUAUUGGGUAUUAAAUAAGU---------   2       21
----------CUUAUCUAAUACCCAAUAAAC------------------------------------------------------------------------------------   4       21
-------------------------------------AAUCAGUAUCUAUCACCAGAC---------------------------------------------------------   2       21
--------------------------------------------------------------------------------------UUAUUGGGUAUUAAAUAAGUAUG------   1       23
-----------------------------------------------------------------------------------------UUGGGUAUUAAAUAAGUA--------   2       18
------------------------------------------------------------------------------------------UGGGUAUUAAAUAAGUAUGG-----   3       20
Library = 1       Precision: [ total = 93.7% | 5p-arm = 50.0% | 3p-arm = 93.8% ]
--------------------------------------------------------------------------------------UUAUUGGGUAUUAAAUAAGUA--------   944     21
--------------------------------------------------------------------------------------UUAUUGGGUAUUAAAUAAGU---------   216     20
--------------------------------------------------------------------------------------UUAUUGGGUAUUAAAUAAGUAU-------   86      22
------------------------------------------------------------------------------------------UGGGUAUUAAAUAAGUAUGGU----   57      21
-----------------------------------------------------------------------------------------UUGGGUAUUAAAUAAGUAUGG-----   24      21
--------------------------------------------------------------------------------------UUAUUGGGUAUUAAAUAAG----------   3       19
-------------------------------------------------------------------------------------UUUAUUGGGUAUUAAAUAAGU---------   2       21
----------CUUAUCUAAUACCCAAUAAAC------------------------------------------------------------------------------------   1       21
-------------------------------------AAUCAGUAUCUAUCACCAGAC---------------------------------------------------------   1       21
--------------------------------------------------------------------------------------UUAUUGGGUAUUAAAUAAGUAUG------   1       23
Library = 2       Precision: [ total = 90.3% | 5p-arm = 75.0% | 3p-arm = 90.4% ]
--------------------------------------------------------------------------------------UUAUUGGGUAUUAAAUAAGUA--------   562     21
--------------------------------------------------------------------------------------UUAUUGGGUAUUAAAUAAGU---------   136     20
--------------------------------------------------------------------------------------UUAUUGGGUAUUAAAUAAGUAU-------   79      22
------------------------------------------------------------------------------------------UGGGUAUUAAAUAAGUAUGGU----   61      21
-----------------------------------------------------------------------------------------UUGGGUAUUAAAUAAGUAUGG-----   17      21
----------CUUAUCUAAUACCCAAUAAAC------------------------------------------------------------------------------------   3       21
------------------------------------------------------------------------------------------UGGGUAUUAAAUAAGUAUGG-----   3       20
--------------------------------------------------------------------------------------UUAUUGGGUAUUAAAUAAG----------   2       19
-----------------------------------------------------------------------------------------UUGGGUAUUAAAUAAGUA--------   2       18
-------------------------------------AAUCAGUAUCUAUCACCAGAC---------------------------------------------------------   1       21
Library = 3       Precision: [ total = 88.7% | 5p-arm = 0% | 3p-arm = 88.7% ]
--------------------------------------------------------------------------------------UUAUUGGGUAUUAAAUAAGUA--------   176     21
--------------------------------------------------------------------------------------UUAUUGGGUAUUAAAUAAGU---------   36      20
------------------------------------------------------------------------------------------UGGGUAUUAAAUAAGUAUGGU----   25      21
--------------------------------------------------------------------------------------UUAUUGGGUAUUAAAUAAGUAU-------   7       22
-----------------------------------------------------------------------------------------UUGGGUAUUAAAUAAGUAUGG-----   3       21
```

```
AU  U          C                A  G                 A   A  A
  CG CUGUAcUUAU UAAUACCCAAUAAAcA CA UAAUCA---------GU UCU UC C
  GU GGUaUGAAUA AUUAUGGGUUAUuUGU GU AUUGGU         CG AGA AG C
CU  U          A                A  A      UGAACUGAA A   C  A
```

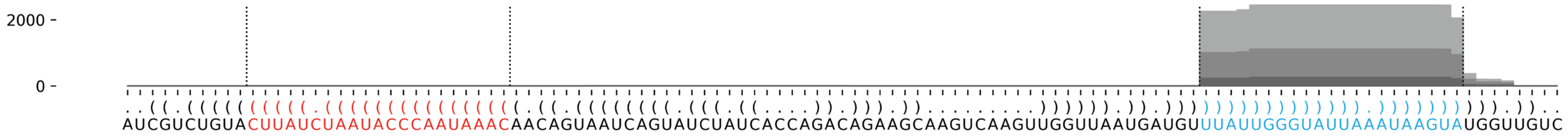

**A. subglobosum** asu-mir-1204-P1-8

```
(((((.((((.((((((..((((((((((((((((.((.(((.((.(((...))))).))).)).))))))))))))))))..)))))).))))))))). -48.86 kcal/mol
AUUGGAUGUUuCAAGUGAGGUAAUGGAUAAuAGAUAUUGAUUGAAAUAGCUUUGCUUUACGGGCACUGUUUAUuAUCUAUUGUAGCACUUGUGaCACCAGUU count    length
Libraries combined         Precision: [ total = 99.5% | 5p-arm = 98.4% | 3p-arm = 99.5% ]
-------------------------------------------------------------------------UAUCUAUUGUAGCACUUGUGA-------- 2237498 21
----------UCAAGUGAGGUAAUGGAUAAU----------------------------------------------------------------------- 9135    21
-------------------------------------------------------------------------UAUCUAUUGUAGCACUUGUG--------- 5991    20
-----------------------------------------------------------------------------UAUUGUAGCACUUGUGACACC---- 4953    21
------------------------------------------------------------------------UUAUCUAUUGUAGCACUUGUGA-------- 3192    22
-------------------------------------------------------------------------UAUCUAUUGUAGCACUUG----------- 2618    18
-------------------------------------------------------------------------UAUCUAUUGUAGCACUUGU---------- 1334    19
-------------------------------------------------------------------------UAUCUAUUGUAGCACUUGUGAC------- 792     22
--------------------------------------------------------------------------AUCUAUUGUAGCACUUGUGA-------- 597     20
-----------------------------------------------------------------------------UAUUGUAGCACUUGUGACAC----- 401     20
-------------------------------------------------------------------------UAUCUAUUGUAGCACUUGUGACA------ 390     23
----------UCAAGUGAGGUAAUGGAUAA------------------------------------------------------------------------ 338     20
-------------------------------------------------------------------------UAUCUAUUGUAGCACUUGUGACACC---- 294     25
-------------------------------------------------------------------------UAUCUAUUGUAGCACUUGUGACACCAG-- 280     27
--------------------------------------------------------------------------AUCUAUUGUAGCACUUGUGAC------- 121     21
----------UCAAGUGAGGUAAUGGAUA------------------------------------------------------------------------- 131     19
-----------------------------------------------------------------------------UAUUGUAGCACUUGUGACA------ 112     19
----------UCAAGUGAGGUAAUGGAUAAUA---------------------------------------------------------------------- 104     22
-------------------------------------------------------------------------UAUCUAUUGUAGCACUUGUGACAC----- 89      24
-------------------------------------------------------------------------UAUCUAUUGUAGCACUUGUGACACCA--- 86      26
-------------------------------------------------------------------------UAUCUAUUGUAGCACUUGUGACACCAGU- 62      28
---------------------------------------------------------------------------UCUAUUGUAGCACUUGUGA-------- 50      19
---------------------------------------------------------------------------UCUAUUGUAGCACUUGUGACACC---- 51      23
----------------------------------------------------------------------------CUAUUGUAGCACUUGUGA-------- 48      18
------UGUUUCAAGUGAGGUAAUGGA--------------------------------------------------------------------------- 38      21
----------UCAAGUGAGGUAAUGGAU-------------------------------------------------------------------------- 51      18
--------UUUCAAGUGAGGUAAUGGAUA------------------------------------------------------------------------- 31      21
---------UUCAAGUGAGGUAAUGGAUAA------------------------------------------------------------------------ 29      21
------------------------------------------------------------------------UUAUCUAUUGUAGCACUUGUG--------- 17      21
--------------------------------------------------------------------------AUCUAUUGUAGCACUUGUGACA------ 23      22
-----------------------------------------------------------------------------UAUUGUAGCACUUGUGACACCA--- 21      22
---------------------------------------------------------------------------UCUAUUGUAGCACUUGUGACA------ 23      21
----------------------------------------------------------------------UAUUAUCUAUUGUAGCACU------------- 12      19
---------------------------------------------------------------------------UCUAUUGUAGCACUUGUGACAC----- 9       22
-----------CAAGUGAGGUAAUGGAUAAU----------------------------------------------------------------------- 10      20
----------------------------------------------------------------------UAUUAUCUAUUGUAGCACUUG----------- 9       21
-----------------------------------------------------------------------------UAUUGUAGCACUUGUGAC------- 12      18
--------------------------------------------------------------------------AUCUAUUGUAGCACUUGUGACACC---- 6       24
---------------------------------------------------------------------------UCUAUUGUAGCACUUGUGAC------- 7       20
------UGUUUCAAGUGAGGUAAUGGAU-------------------------------------------------------------------------- 8       22
---------UUCAAGUGAGGUAAUGGAUAAU----------------------------------------------------------------------- 5       22
--------------------------------------------------------------------------AUCUAUUGUAGCACUUGUGACACCAG-- 6       26
----------------------------------------------------------------------------CUAUUGUAGCACUUGUGACACC---- 10      22
AUUGGAUGUUUCAAGUGAGGUAAUGGAUAAU----------------------------------------------------------------------- 3       31
------UGUUUCAAGUGAGGUAAUG----------------------------------------------------------------------------- 5       19
--------UUUCAAGUGAGGUAAUGGAUAA------------------------------------------------------------------------ 7       22
--------------------UAAUGGAUAAUAGAUAUUGAU------------------------------------------------------------- 2       21
---------------------AAUGGAUAAUAGAUAUUGAU------------------------------------------------------------- 2       20
-----------------------------------------------------------------------AUUAUCUAUUGUAGCACUUGU---------- 3       21
------------------------------------------------------------------------UUAUCUAUUGUAGCACUUGUGAC------- 5       23
------------------------------------------------------------------------UUAUCUAUUGUAGCACUUGUGACA------ 2       24
------------------------------------------------------------------------UUAUCUAUUGUAGCACUUG----------- 3       19
---GGAUGUUUCAAGUGAGGUA-------------------------------------------------------------------------------- 1       19
```

#### *A. subglobosum* asu-mir-1204 continued

```
(((((.((((.((((((..(((((((((((((((((.((.(((.((.(((...))))).))).)).)))))))))))))))))..)))))).))))))))). -48.86 kcal/mol
AUUGGAUGUUuCAAGUGAGGUAAUGGAUAAuAGAUAUUGAUUGAAAUAGCUUUGCUUUACGGGCACUGUUUAUuAUCUAUUGUAGCACUUGUGaCACCAGUU count    length
Libraries combined          Precision: [ total = 99.5% | 5p-arm = 98.4% | 3p-arm = 99.5% ]
---GGAUGUUUCAAGUGAGGUAA------------------------------------------------------------------------------- 1        20
-------GUUUCAAGUGAGGUAAUGGAUA------------------------------------------------------------------------- 1        22
--------UUUCAAGUGAGGUAAUGGAUAAU----------------------------------------------------------------------- 1        23
---------UUCAAGUGAGGUAAUGGAUA------------------------------------------------------------------------- 3        20
------------AAGUGAGGUAAUGGAUAAU----------------------------------------------------------------------- 1        19
----------------GAGGUAAUGGAUAAUAGAU------------------------------------------------------------------- 1        19
-------------------------------------------------------UUUACGGGCACUGUUUAU----------------------------- 1        18
--------------------------------------------------------UUACGGGCACUGUUUAUUAUC------------------------- 6        21
----------------------------------------------------------ACGGGCACUGUUUAUUAUC------------------------- 1        19
----------------------------------------------------------ACGGGCACUGUUUAUUAUCUA----------------------- 1        21
-------------------------------------------------------------GGCACUGUUUAUUAUCUAUUG-------------------- 1        21
----------------------------------------------------------------------UAUUAUCUAUUGUAGCACUUGU---------- 2        22
------------------------------------------------------------------------UUAUCUAUUGUAGCACUUGU---------- 5        20
------------------------------------------------------------------------UUAUCUAUUGUAGCACUUGUGACACC---- 2        26
-------------------------------------------------------------------------UAUCUAUUGUAGCACUUGUGACACCAGUU 3        29
--------------------------------------------------------------------------AUCUAUUGUAGCACUUGUGACACCA--- 2        25
------------------------------------------------------------------------------AUUGUAGCACUUGUGACACC---- 1        20
-------------------------------------------------------------------------------UUGUAGCACUUGUGACACC---- 1        19
-------------------------------------------------------------------------------UUGUAGCACUUGUGACACCAG-- 3        21
----------------------------------------------------------------------------------UAGCACUUGUGACACCAGUU 2        20
AUUGGAUGUUUCAAGUGAGGUAAUGGAUAA------------------------------------------------------------------------ 1        30
---GGAUGUUUCAAGUGAGGUAAUGGA--------------------------------------------------------------------------- 1        24
------UGUUUCAAGUGAGGUAAUGG---------------------------------------------------------------------------- 1        20
-----------CAAGUGAGGUAAUGGAUAA------------------------------------------------------------------------ 2        19
----------------GAGGUAAUGGAUAAUAGAUAUU---------------------------------------------------------------- 1        22
-----------------------------------AUUGAUUGAAAUAGCUUUGCU---------------------------------------------- 1        21
-------------------------------------------AAAUAGCUUUGCUUUACGGGC-------------------------------------- 1        21
--------------------------------------------------------------------UUUAUUAUCUAUUGUAGCACU------------- 2        21
----------------------------------------------------------------------UAUUAUCUAUUGUAGCACUU------------ 2        20
------------------------------------------------------------------------UUAUCUAUUGUAGCACUUGUGACAC----- 1        25
--------------------------------------------------------------------------AUCUAUUGUAGCACUUGUG--------- 1        19
---------------------------------------------------------------------------UCUAUUGUAGCACUUGUGACACCA--- 1        24
---------------------------------------------------------------------------UCUAUUGUAGCACUUGUGACACCAG-- 2        25
--------------------------------------------------------------------------------UGUAGCACUUGUGACACCAGU- 1        21
----------------------------------------------------------------------------------UAGCACUUGUGACACCAGU- 1        19
```

```
      A    u      AG                 U  A   A  U   U
 AUUGG UGUU CAAGUG  GUAAUGGAUAAuAGAUA UG UUG AA AGC U
 UGACC-ACaG GUUCAC  UGUUAUCUAuUAUUUGU AC GGC UU-UCG /
U          U      GA                 C  G   A      U
```

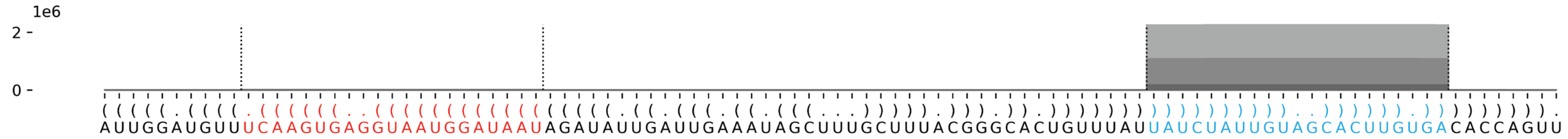

#### *A. subglobosum* asu-mir-1205

```
....(((((((((((((((..(.((((.((((((((..((((((((..((.((((((....(........)....)))))))).))..)))))))).)))).)))).)))).).)).)))))))))))))))))....   -70.5 kcal/mol
UAUUAUGUCUuGAUGAACAUUUGAGUGGCGuUGAAUGGAAGUGUCUCAAUUAUGUUGACAUUCAUUCAAUCGUUCAUUGACAAGAGAGAGAUACUACAUUGCAGCuCCACACUUAUGUUCAUCAAgACAUCGAC   count    length
Libraries combined           Precision: [ total = 98.5% | 5p-arm = 98.6% | 3p-arm = 85.7% ]
----------UGAUGAACAUUUGAGUGGCGU-------------------------------------------------------------------------------------------------------   8502     21
----------UGAUGAACAUUUGAGUGGCG--------------------------------------------------------------------------------------------------------   584      20
----------UGAUGAACAUUUGAGUGGC---------------------------------------------------------------------------------------------------------   153      19
----------UGAUGAACAUUUGAGUGGCGUU------------------------------------------------------------------------------------------------------   123      22
----------UGAUGAACAUUUGAGUGG----------------------------------------------------------------------------------------------------------   68       18
---------UUGAUGAACAUUUGAGUGGCGU-------------------------------------------------------------------------------------------------------   49       22
---------UUGAUGAACAUUUGAGUGGCG--------------------------------------------------------------------------------------------------------   27       21
-----------GAUGAACAUUUGAGUGGCGU-------------------------------------------------------------------------------------------------------   30       20
---------UUGAUGAACAUUUGAGUGG----------------------------------------------------------------------------------------------------------   9        19
---------UUGAUGAACAUUUGAGUGGC---------------------------------------------------------------------------------------------------------   13       20
---------------------------------------------------------------------------------------------------------UCCACACUUAUGUUCAUCAAG--------   6        21
---------UUGAUGAACAUUUGAGUG-----------------------------------------------------------------------------------------------------------   3        18
---UAUGUCUUGAUGAACAUUUGAGUGGCGU-------------------------------------------------------------------------------------------------------   1        28
---------UUGAUGAACAUUUGAGUGGCGUU------------------------------------------------------------------------------------------------------   2        23
----------UGAUGAACAUUUGAGUGGCGUUG-----------------------------------------------------------------------------------------------------   1        23
------------------------------------------------------------UUCAUUCAAUCGUUCAUUGACAAGA-------------------------------------------------   1        25
-----------GAUGAACAUUUGAGUGGCG--------------------------------------------------------------------------------------------------------   3        19
-------------UGAACAUUUGAGUGGCGU-------------------------------------------------------------------------------------------------------   1        18
Library = 1         Precision: [ total = 98.6% | 5p-arm = 98.7% | 3p-arm = 75.0% ]
----------UGAUGAACAUUUGAGUGGCGU-------------------------------------------------------------------------------------------------------   3976     21
----------UGAUGAACAUUUGAGUGGCG--------------------------------------------------------------------------------------------------------   298      20
----------UGAUGAACAUUUGAGUGGC---------------------------------------------------------------------------------------------------------   78       19
----------UGAUGAACAUUUGAGUGGCGUU------------------------------------------------------------------------------------------------------   63       22
----------UGAUGAACAUUUGAGUGG----------------------------------------------------------------------------------------------------------   31       18
---------UUGAUGAACAUUUGAGUGGCGU-------------------------------------------------------------------------------------------------------   21       22
---------UUGAUGAACAUUUGAGUGGCG--------------------------------------------------------------------------------------------------------   18       21
-----------GAUGAACAUUUGAGUGGCGU-------------------------------------------------------------------------------------------------------   10       20
---------UUGAUGAACAUUUGAGUGG----------------------------------------------------------------------------------------------------------   4        19
---------UUGAUGAACAUUUGAGUGGC---------------------------------------------------------------------------------------------------------   3        20
---------------------------------------------------------------------------------------------------------UCCACACUUAUGUUCAUCAAG--------   3        21
---------UUGAUGAACAUUUGAGUG-----------------------------------------------------------------------------------------------------------   2        18
---UAUGUCUUGAUGAACAUUUGAGUGGCGU-------------------------------------------------------------------------------------------------------   1        28
---------UUGAUGAACAUUUGAGUGGCGUU------------------------------------------------------------------------------------------------------   1        23
----------UGAUGAACAUUUGAGUGGCGUUG-----------------------------------------------------------------------------------------------------   1        23
------------------------------------------------------------UUCAUUCAAUCGUUCAUUGACAAGA-------------------------------------------------   1        25
Library = 2         Precision: [ total = 98.5% | 5p-arm = 98.5% | 3p-arm = 100.0% ]
----------UGAUGAACAUUUGAGUGGCGU-------------------------------------------------------------------------------------------------------   3687     21
----------UGAUGAACAUUUGAGUGGCG--------------------------------------------------------------------------------------------------------   243      20
----------UGAUGAACAUUUGAGUGGC---------------------------------------------------------------------------------------------------------   67       19
----------UGAUGAACAUUUGAGUGGCGUU------------------------------------------------------------------------------------------------------   47       22
----------UGAUGAACAUUUGAGUGG----------------------------------------------------------------------------------------------------------   33       18
---------UUGAUGAACAUUUGAGUGGCGU-------------------------------------------------------------------------------------------------------   24       22
-----------GAUGAACAUUUGAGUGGCGU-------------------------------------------------------------------------------------------------------   14       20
---------UUGAUGAACAUUUGAGUGGCG--------------------------------------------------------------------------------------------------------   9        21
---------UUGAUGAACAUUUGAGUGGC---------------------------------------------------------------------------------------------------------   8        20
---------UUGAUGAACAUUUGAGUGG----------------------------------------------------------------------------------------------------------   3        19
-----------GAUGAACAUUUGAGUGGCG--------------------------------------------------------------------------------------------------------   3        19
---------------------------------------------------------------------------------------------------------UCCACACUUAUGUUCAUCAAG--------   2        21
---------UUGAUGAACAUUUGAGUG-----------------------------------------------------------------------------------------------------------   1        18
---------UUGAUGAACAUUUGAGUGGCGUU------------------------------------------------------------------------------------------------------   1        23
-------------UGAACAUUUGAGUGGCGU-------------------------------------------------------------------------------------------------------   1        18
Library = 3         Precision: [ total = 98.5% | 5p-arm = 98.5% | 3p-arm = 100.0% ]
----------UGAUGAACAUUUGAGUGGCGU-------------------------------------------------------------------------------------------------------   839      21
```

##### *A. subglobosum* asu-mir-1205 continued

```
....((((((((((((((((..(.((((.((((((((..((((((((..((.((((((....(........)....)))))).))..)))))))).)))).)))).)))).)..))))))))))))))))....   -70.5 kcal/mol
UAUUAUGUCUuGAUGAACAUUUGAGUGGCGuUGAAUGGAAGUGUCUCAAUUAUGUUGACAUUCAUUCAAUCGUUCAUUGACAAGAGAGAGAUACUACAUUGCAGCuCCACACUUAUGUUCAUCAAgACAUCGAC   count    length
----------UGAUGAACAUUUGAGUGGCG--------------------------------------------------------------------------------------------------------   43       20
----------UGAUGAACAUUUGAGUGGCGUU------------------------------------------------------------------------------------------------------   13       22
----------UGAUGAACAUUUGAGUGGC---------------------------------------------------------------------------------------------------------   8        19
-----------GAUGAACAUUUGAGUGGCGU-------------------------------------------------------------------------------------------------------   6        20
---------UUGAUGAACAUUUGAGUGGCGU-------------------------------------------------------------------------------------------------------   4        22
----------UGAUGAACAUUUGAGUGG----------------------------------------------------------------------------------------------------------   4        18
---------UUGAUGAACAUUUGAGUGG----------------------------------------------------------------------------------------------------------   2        19
---------UUGAUGAACAUUUGAGUGGC---------------------------------------------------------------------------------------------------------   2        20
---------------------------------------------------------------------------------------------------------UCCACACUUAUGUUCAUCAAG--------   1        21
```

```
UAUU                UU A    C         GA        AA  A      CAUU AUU
    AUGUCUuGAUGAACAU  G GUGG GuUG-AAUG  AGUGUCUC  UU UGUUGA    C   C
    UACAgAACUACUUGUA  C CACC CGAC UUAC  UCAUAGAG  AG ACAGUU    G   A
CAGC                UU A    u    G    A-        AG  A      ACUU CUA
```

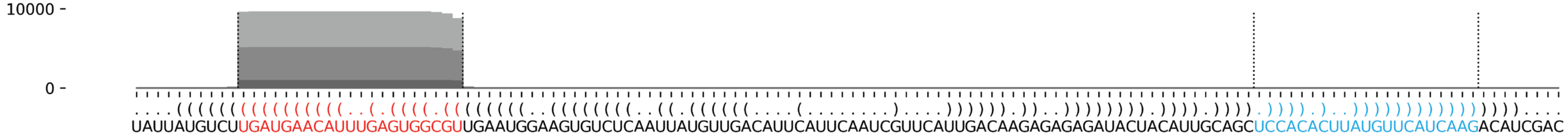


##### *A. subglobosum* asu-mir-1206

```
...(..(.((((((((.(((((((((((.(((...((((.((((((((((((.(((((((((((((..((.......))..))))))).)))))).)))))).)))...))).))))...))).))))))))))).)))))))).)..)...   -85.3 kcal/mol
UAUGUUAUCGaAUCGAGUGAGAUGAUGCUUaCCAAUGGAAUGACCAAGUGUCAGUGUUCAAACUGCAAUGUUAUAAUCACCGCAGUUUGUAAUACCGACGCUCUGGUAUUCAAUCCACCUGUAuGCAUCAUCUCGAUCGAUUCgCUUUCUAC   count    length
Libraries combined         Precision: [ total = 99.3% | 5p-arm = 50.0% | 3p-arm = 99.3% ]
---------------------------------------------------------------------------------------------------------------------------UGCAUCAUCUCGAUCGAUUCG--------   23374    21
----------------------------------------------------------------------------------------------------------------------------GCAUCAUCUCGAUCGAUUCG--------   131      20
---------------------------------------------------------------------------------------------------------------------------UGCAUCAUCUCGAUCGAUU----------   14       19
---------------------------------------------------------------------------------------------------------------------------UGCAUCAUCUCGAUCGAUUCGC-------   23       22
---------------------------------------------------------------------------------------------------------------------------UGCAUCAUCUCGAUCGAUUCGCU------   14       23
---------------------------------------------------------------------------------------------------------------------------UGCAUCAUCUCGAUCGAUUC---------   16       20
-------------------------------------------------------------------------------------------------------------------------------UCAUCUCGAUCGAUUCGCU------   8        19
----UUAUCGAAUCGAGUGAGAUGA-------------------------------------------------------------------------------------------------------------------------------   1        21
----UUAUCGAAUCGAGUGAGAU---------------------------------------------------------------------------------------------------------------------------------   1        19
-----UAUCGAAUCGAGUGAGAUGA-------------------------------------------------------------------------------------------------------------------------------   1        20
------AUCGAAUCGAGUGAGAUGAU------------------------------------------------------------------------------------------------------------------------------   1        20
----------AAUCGAGUGAGAUGAUGCUU--------------------------------------------------------------------------------------------------------------------------   1        20
---------------------------------------------AAGUGUCAGUGUUCAAACUGC--------------------------------------------------------------------------------------   1        21
-------------------------------------------------------------------------------------UUUGUAAUACCGACGCUCUGGU---------------------------------------------   1        22
--------------------------------------------------------------------------------------UUGUAAUACCGACGCUCUGGUAU-------------------------------------------   2        23
-------------------------------------------------------------------------------------------------------------------------UAUGCAUCAUCUCGAUCGAUUCGC-------   1        24
--------------------------------------------------------------------------------------------------------------------------AUGCAUCAUCUCGAUCGAUUCG--------   1        22
---------------------------------------------------------------------------------------------------------------------------UGCAUCAUCUCGAUCGAU-----------   1        18
-----------------------------------------------------------------------------------------------------------------------------CAUCAUCUCGAUCGAUUCGCU------   1        21
-------------------------------------------------------------------------------------------------------------------------------UCAUCUCGAUCGAUUCGCUUU----   2        21
-------------------------------------------------------------------------------------------------------------------------------UCAUCUCGAUCGAUUCGCUUUCU--   2        23
-------------------------------------------------------------------------------------------------------------------------------UCAUCUCGAUCGAUUCGCUUUC---   2        22
----------AAUCGAGUGAGAUGAUGCUUA-------------------------------------------------------------------------------------------------------------------------   4        21
-------------------------------------------------------------------------UAAUCACCGCAGUUUGUAAUAC---------------------------------------------------------   1        22
--------------------------------------------------------------------------------------UUGUAAUACCGACGCUCUGGU---------------------------------------------   1        21
--------------------------------------------------------------------------------------------------------------------------AUGCAUCAUCUCGAUCGAUUC---------   1        21
```

#### *A. subglobosum* asu-mir-1206 continued

```
...(..(.((((((((.((((((((((.(((...((((.((((((((((((.((((((((((((..((.......))..)))))))).))))).)))))).)))...))).))))...))).)))))))))))).)))))))).)..)...   -85.3 kcal/mol
UAUGUUAUCGaAUCGAGUGAGAUGAUGCUUaCCAAUGGAAUGACCAAGUGUCAGUGUUCAAACUGCAAUGUUAUAAUCACCGCAGUUUGUAAUACCGACGCUCUGGUAUUCAAUCCACCUGUAuGCAUCAUCUCGAUCGAUUCgCUUUCUAC   count   length
Library = 1      Precision: [ total = 99.3% | 5p-arm = 16.7% | 3p-arm = 99.3% ]
---------------------------------------------------------------------------------------------------------------------------UGCAUCAUCUCGAUCGAUUCG--------   11768   21
----------------------------------------------------------------------------------------------------------------------------GCAUCAUCUCGAUCGAUUCG--------   72      20
---------------------------------------------------------------------------------------------------------------------------UGCAUCAUCUCGAUCGAUU----------   8       19
---------------------------------------------------------------------------------------------------------------------------UGCAUCAUCUCGAUCGAUUCGC-------   8       22
---------------------------------------------------------------------------------------------------------------------------UGCAUCAUCUCGAUCGAUUCGCU------   4       23
---------------------------------------------------------------------------------------------------------------------------UGCAUCAUCUCGAUCGAUUC---------   3       20
-------------------------------------------------------------------------------------------------------------------------------UCAUCUCGAUCGAUUCGCU------   2       19
----UUAUCGAAUCGAGUGAGAUGA-------------------------------------------------------------------------------------------------------------------------------   1       21
----UUAUCGAAUCGAGUGAGAU---------------------------------------------------------------------------------------------------------------------------------   1       19
-----UAUCGAAUCGAGUGAGAUGA-------------------------------------------------------------------------------------------------------------------------------   1       20
------AUCGAAUCGAGUGAGAUGAU------------------------------------------------------------------------------------------------------------------------------   1       20
----------AAUCGAGUGAGAUGAUGCUU--------------------------------------------------------------------------------------------------------------------------   1       20
---------------------------------------------AAGUGUCAGUGUUCAAACUGC--------------------------------------------------------------------------------------   1       21
-------------------------------------------------------------------------------------UUUGUAAUACCGACGCUCUGGU---------------------------------------------   1       22
--------------------------------------------------------------------------------------UUGUAAUACCGACGCUCUGGUAU-------------------------------------------   1       23
-------------------------------------------------------------------------------------------------------------------------UAUGCAUCAUCUCGAUCGAUUCGC-------   1       24
--------------------------------------------------------------------------------------------------------------------------AUGCAUCAUCUCGAUCGAUUCG--------   1       22
---------------------------------------------------------------------------------------------------------------------------UGCAUCAUCUCGAUCGAU-----------   1       18
-----------------------------------------------------------------------------------------------------------------------------CAUCAUCUCGAUCGAUUCGCU------   1       21
-------------------------------------------------------------------------------------------------------------------------------UCAUCUCGAUCGAUUCGCUUU----   1       21
-------------------------------------------------------------------------------------------------------------------------------UCAUCUCGAUCGAUUCGCUUUCU--   1       23
-------------------------------------------------------------------------------------------------------------------------------UCAUCUCGAUCGAUUCGCUUUC---   1       22
Library = 2      Precision: [ total = 99.4% | 5p-arm = 100.0% | 3p-arm = 99.4% ]
---------------------------------------------------------------------------------------------------------------------------UGCAUCAUCUCGAUCGAUUCG--------   9733    21
----------------------------------------------------------------------------------------------------------------------------GCAUCAUCUCGAUCGAUUCG--------   50      20
---------------------------------------------------------------------------------------------------------------------------UGCAUCAUCUCGAUCGAUUC---------   12      20
---------------------------------------------------------------------------------------------------------------------------UGCAUCAUCUCGAUCGAUUCGC-------   11      22
---------------------------------------------------------------------------------------------------------------------------UGCAUCAUCUCGAUCGAUUCGCU------   8       23
---------------------------------------------------------------------------------------------------------------------------UGCAUCAUCUCGAUCGAUU----------   6       19
----------AAUCGAGUGAGAUGAUGCUUA-------------------------------------------------------------------------------------------------------------------------   4       21
-------------------------------------------------------------------------------------------------------------------------------UCAUCUCGAUCGAUUCGCU------   4       19
-------------------------------------------------------------------------UAAUCACCGCAGUUUGUAAUAC---------------------------------------------------------   1       22
--------------------------------------------------------------------------------------UUGUAAUACCGACGCUCUGGUAU-------------------------------------------   1       23
--------------------------------------------------------------------------------------UUGUAAUACCGACGCUCUGGU---------------------------------------------   1       21
--------------------------------------------------------------------------------------------------------------------------AUGCAUCAUCUCGAUCGAUUC---------   1       21
-------------------------------------------------------------------------------------------------------------------------------UCAUCUCGAUCGAUUCGCUUU----   1       21
-------------------------------------------------------------------------------------------------------------------------------UCAUCUCGAUCGAUUCGCUUUCU--   1       23
-------------------------------------------------------------------------------------------------------------------------------UCAUCUCGAUCGAUUCGCUUUC---   1       22
Library = 3      Precision: [ total = 99.4% | 5p-arm = 0% | 3p-arm = 99.4% ]
---------------------------------------------------------------------------------------------------------------------------UGCAUCAUCUCGAUCGAUUCG--------   1873    21
----------------------------------------------------------------------------------------------------------------------------GCAUCAUCUCGAUCGAUUCG--------   9       20
---------------------------------------------------------------------------------------------------------------------------UGCAUCAUCUCGAUCGAUUCGC-------   4       22
---------------------------------------------------------------------------------------------------------------------------UGCAUCAUCUCGAUCGAUUCGCU------   2       23
-------------------------------------------------------------------------------------------------------------------------------UCAUCUCGAUCGAUUCGCU------   2       19
---------------------------------------------------------------------------------------------------------------------------UGCAUCAUCUCGAUCGAUUC---------   1       20
```

```
UAU UU U         G            U    CAA     A                  A               AA   UUA
   G  A CGaAUCGA UGAGAUGAUGC UaC   UGGA UGA---CCA-AGUGUC GUGUU-CAAACUGC  UG    U
   C  U gCUUAGCU GCUCUACUACG AUG   ACCU ACU    GGU UCGCAG CAUAA GUUUGACG  AC    /
CAU UU C         A            u    UCC     A  UAU    C       C      U         CC   UAA
```

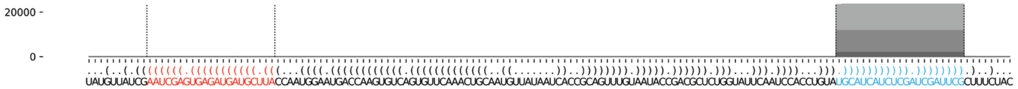

### *A. subglobosum* asu-mir-1207

```
.....((((.(((((...((((((((((((((((.(((((((((.......((..((.(((((((((((((.(.....(((.(((....))))))).....).)))))))))))))))...))))))))))).))))))))))))))))...))))).)))).....   -87.6 kcal/mol
AACACACACUuGCGCUAUCUGUUCAAACAAuCGGAGUUACCGAGUACAACUCCUACGAUUAUAAGUGUCACGGACUACUCACUCAGCAAUGAUGAAUCAAUGGUGAUACUUGUGACGCAUGGCUCGGUAGUAUCGAUuGUUUGGACAGCUUGCGUAUgUGUUUACC   count   length
Libraries combined   Precision: [ total = 99.9% | 5p-arm = 87.5% | 3p-arm = 99.9% ]
-----------------------------------------------------------------------------------------------------------------------------------------UGUUUGGACAGCUUGCGUAUG--------   10264   21
-----------------------------------------------------------------------------------------------------------------------------------------UGUUUGGACAGCUUGCGUAUGU-------   354     22
-----------------------------------------------------------------------------------------------------------------------------------------UGUUUGGACAGCUUGCGUAU---------   220     20
-----------------------------------------------------------------------------------------------------------------------------------------UGUUUGGACAGCUUGCGUA----------   81      19
-----------------------------------------------------------------------------------------------------------------------------------------UGUUUGGACAGCUUGCGU-----------   26      18
----------------------------------------------------------------------------------------------------------------------------------------UUGUUUGGACAGCUUGCGUAUG--------   6       22
----------UGCGCUAUCUGUUCAAACAAU---------------------------------------------------------------------------------------------------------------------------------------   6       21
---------------------------------------------------------AUUAUAAGUGUCACGGACU------------------------------------------------------------------------------------------   1       19
-----------------------------------------------------------------------------------------------------------------------UGGCUCGGUAGUAUCGAUUGU--------------------------   1       21
--------------------------------------------------------------------------------------------------------------------------------------GAUUGUUUGGACAGCUUGCGU-----------   1       21
----------------------------------------------------------------------------------------------------------------------------------------UUGUUUGGACAGCUUGCGUAU---------   2       21
------------------------------------------------------------------------------------------------------------------------------------------GUUUGGACAGCUUGCGUAUG--------   1       20
-------------------------------------------------------------------------------------------------------------------------------------------UUUGGACAGCUUGCGUAUG--------   1       19
----------UGCGCUAUCUGUUCAAACAAUC--------------------------------------------------------------------------------------------------------------------------------------   1       22
-----------------------------------------------------------------------------------------------------------------------------------------UGUUUGGACAGCUUGCGUAUGUG------   1       23
Library = 1          Precision: [ total = 99.8% | 5p-arm = 80.0% | 3p-arm = 99.8% ]
-----------------------------------------------------------------------------------------------------------------------------------------UGUUUGGACAGCUUGCGUAUG--------   4784    21
-----------------------------------------------------------------------------------------------------------------------------------------UGUUUGGACAGCUUGCGUAUGU-------   171     22
-----------------------------------------------------------------------------------------------------------------------------------------UGUUUGGACAGCUUGCGUAU---------   104     20
-----------------------------------------------------------------------------------------------------------------------------------------UGUUUGGACAGCUUGCGUA----------   39      19
-----------------------------------------------------------------------------------------------------------------------------------------UGUUUGGACAGCUUGCGU-----------   13      18
----------------------------------------------------------------------------------------------------------------------------------------UUGUUUGGACAGCUUGCGUAUG--------   5       22
----------UGCGCUAUCUGUUCAAACAAU---------------------------------------------------------------------------------------------------------------------------------------   4       21
---------------------------------------------------------AUUAUAAGUGUCACGGACU------------------------------------------------------------------------------------------   1       19
-----------------------------------------------------------------------------------------------------------------------UGGCUCGGUAGUAUCGAUUGU--------------------------   1       21
--------------------------------------------------------------------------------------------------------------------------------------GAUUGUUUGGACAGCUUGCGU-----------   1       21
----------------------------------------------------------------------------------------------------------------------------------------UUGUUUGGACAGCUUGCGUAU---------   1       21
------------------------------------------------------------------------------------------------------------------------------------------GUUUGGACAGCUUGCGUAUG--------   1       20
-------------------------------------------------------------------------------------------------------------------------------------------UUUGGACAGCUUGCGUAUG--------   1       19
Library = 2          Precision: [ total = 100.0% | 5p-arm = 100.0% | 3p-arm = 100.0% ]
-----------------------------------------------------------------------------------------------------------------------------------------UGUUUGGACAGCUUGCGUAUG--------   4393    21
-----------------------------------------------------------------------------------------------------------------------------------------UGUUUGGACAGCUUGCGUAUGU-------   145     22
-----------------------------------------------------------------------------------------------------------------------------------------UGUUUGGACAGCUUGCGUAU---------   97      20
-----------------------------------------------------------------------------------------------------------------------------------------UGUUUGGACAGCUUGCGUA----------   34      19
-----------------------------------------------------------------------------------------------------------------------------------------UGUUUGGACAGCUUGCGU-----------   12      18
----------UGCGCUAUCUGUUCAAACAAU---------------------------------------------------------------------------------------------------------------------------------------   2       21
----------UGCGCUAUCUGUUCAAACAAUC--------------------------------------------------------------------------------------------------------------------------------------   1       22
----------------------------------------------------------------------------------------------------------------------------------------UUGUUUGGACAGCUUGCGUAUG--------   1       22
-----------------------------------------------------------------------------------------------------------------------------------------UGUUUGGACAGCUUGCGUAUGUG------   1       23
Library = 3          Precision: [ total = 99.9% | 5p-arm = 0% | 3p-arm = 99.9% ]
-----------------------------------------------------------------------------------------------------------------------------------------UGUUUGGACAGCUUGCGUAUG--------   1087    21
-----------------------------------------------------------------------------------------------------------------------------------------UGUUUGGACAGCUUGCGUAUGU-------   38      22
-----------------------------------------------------------------------------------------------------------------------------------------UGUUUGGACAGCUUGCGUAU---------   19      20
-----------------------------------------------------------------------------------------------------------------------------------------UGUUUGGACAGCUUGCGUA----------   8       19
----------------------------------------------------------------------------------------------------------------------------------------UUGUUUGGACAGCUUGCGUAU---------   1       21
-----------------------------------------------------------------------------------------------------------------------------------------UGUUUGGACAGCUUGCGU-----------   1       18
```

```
AACAC    U     UAU                A         UACAACU  UA-  A             G ACUAC   C   G
     ACAC uGCGC   CUGUUCAAACAAuCGG GUUACCGAG       CC   CG UUAUAAGUGUCAC G     UCA UCA C
     UGUg AUGCG   GACAGGUUUGuUAGCU UGAUGGCUC-------GG   GC-AGUGUUCAUAGUG U     AGU-AGU A
CCAUU    U     UUC                A                  UAC                G AACUA       A
```

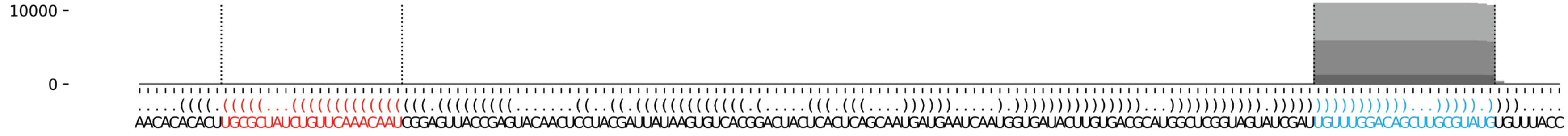

#### *A. subglobosum* asu-mir-1208

```
(((...((.(((((..(((((.(((((.((((.((.((((.....((((((((..(((((....)))))))))))))...((((.((((............)))).)))))))).)).)))).))))).)))))))))).))..))).     -68.63 kcal/mol
GCGACUAGUGuGCUGUGAGAUUGGUCACCGAGcUGAUAGUAAGAAGGAGGUUGACGCACUGUAGAGUGCCAGUCUCCAACAUAACCAACAUUUAAUCUCAGUUGUUUAUACUACCAUCuCGUUGACUUAUCUCAGCACCcCUUACGCG     count    length
Libraries combined          Precision: [ total = 97.8% | 5p-arm = 97.9% | 3p-arm = 89.8% ]
----------UGCUGUGAGAUUGGUCACCGAGC-------------------------------------------------------------------------------------------------------------------     4493     23
----------UGCUGUGAGAUUGGUCACCG----------------------------------------------------------------------------------------------------------------------     3444     20
----------UGCUGUGAGAUUGGUCACCGAG--------------------------------------------------------------------------------------------------------------------     1724     22
----------UGCUGUGAGAUUGGUCACCGA---------------------------------------------------------------------------------------------------------------------     1025     21
----------------------------------------------------------------------------------------------------------------------UCGUUGACUUAUCUCAGCACC---------     211      21
----------UGCUGUGAGAUUGGUCACC-----------------------------------------------------------------------------------------------------------------------     135      19
----------UGCUGUGAGAUUGGUCAC------------------------------------------------------------------------------------------------------------------------     132      18
-----------GCUGUGAGAUUGGUCACCGAGC-------------------------------------------------------------------------------------------------------------------     78       22
------------CUGUGAGAUUGGUCACCGAGCUG-----------------------------------------------------------------------------------------------------------------     43       23
-----------GCUGUGAGAUUGGUCACCGAG--------------------------------------------------------------------------------------------------------------------     47       21
----------UGCUGUGAGAUUGGUCACCGAGCU------------------------------------------------------------------------------------------------------------------     29       24
-------------UGUGAGAUUGGUCACCGAGCUG-----------------------------------------------------------------------------------------------------------------     16       22
---------------UGAGAUUGGUCACCGAGCUG-----------------------------------------------------------------------------------------------------------------     18       20
--------------------------------------------------------------------------------------------------------------------UCUCGUUGACUUAUCUCAGC------------     17       20
-----------GCUGUGAGAUUGGUCACCG----------------------------------------------------------------------------------------------------------------------     14       19
----------UGCUGUGAGAUUGGUCACCGAGCUG-----------------------------------------------------------------------------------------------------------------     14       25
-----------GCUGUGAGAUUGGUCACCGA---------------------------------------------------------------------------------------------------------------------     10       20
--------------------------------------------------------------------------------------------------------------------UCUCGUUGACUUAUCUCAGCA-----------     5        21
-----------GCUGUGAGAUUGGUCACCGAGCU------------------------------------------------------------------------------------------------------------------     2        23
--------------------------------------------------------------------------------------------------------------------------UGACUUAUCUCAGCACCCU-------     1        19
----------UGCUGUGAGAUUGGUCACCGAGCUGA----------------------------------------------------------------------------------------------------------------     1        26
-----------GCUGUGAGAUUGGUCACCGAGCUG-----------------------------------------------------------------------------------------------------------------     1        24
------------CUGUGAGAUUGGUCACCGAG--------------------------------------------------------------------------------------------------------------------     1        20
-------------UGUGAGAUUGGUCACCGAGC-------------------------------------------------------------------------------------------------------------------     1        20
---------------UGAGAUUGGUCACCGAGC-------------------------------------------------------------------------------------------------------------------     1        18
--------------------------------------------------------------------------------------------------------------------UCUCGUUGACUUAUCUCAG-------------     1        19
Library = 1         Precision: [ total = 97.9% | 5p-arm = 98.0% | 3p-arm = 90.3% ]
----------UGCUGUGAGAUUGGUCACCGAGC-------------------------------------------------------------------------------------------------------------------     2246     23
----------UGCUGUGAGAUUGGUCACCG----------------------------------------------------------------------------------------------------------------------     1687     20
----------UGCUGUGAGAUUGGUCACCGAG--------------------------------------------------------------------------------------------------------------------     865      22
----------UGCUGUGAGAUUGGUCACCGA---------------------------------------------------------------------------------------------------------------------     505      21
----------------------------------------------------------------------------------------------------------------------UCGUUGACUUAUCUCAGCACC---------     112      21
----------UGCUGUGAGAUUGGUCACC-----------------------------------------------------------------------------------------------------------------------     69       19
----------UGCUGUGAGAUUGGUCAC------------------------------------------------------------------------------------------------------------------------     57       18
-----------GCUGUGAGAUUGGUCACCGAGC-------------------------------------------------------------------------------------------------------------------     29       22
------------CUGUGAGAUUGGUCACCGAGCUG-----------------------------------------------------------------------------------------------------------------     25       23
-----------GCUGUGAGAUUGGUCACCGAG--------------------------------------------------------------------------------------------------------------------     23       21
----------UGCUGUGAGAUUGGUCACCGAGCU------------------------------------------------------------------------------------------------------------------     18       24
-------------UGUGAGAUUGGUCACCGAGCUG-----------------------------------------------------------------------------------------------------------------     11       22
---------------UGAGAUUGGUCACCGAGCUG-----------------------------------------------------------------------------------------------------------------     8        20
--------------------------------------------------------------------------------------------------------------------UCUCGUUGACUUAUCUCAGC------------     8        20
-----------GCUGUGAGAUUGGUCACCG----------------------------------------------------------------------------------------------------------------------     7        19
----------UGCUGUGAGAUUGGUCACCGAGCUG-----------------------------------------------------------------------------------------------------------------     6        25
-----------GCUGUGAGAUUGGUCACCGA---------------------------------------------------------------------------------------------------------------------     6        20
--------------------------------------------------------------------------------------------------------------------UCUCGUUGACUUAUCUCAGCA-----------     3        21
-----------GCUGUGAGAUUGGUCACCGAGCU------------------------------------------------------------------------------------------------------------------     1        23
--------------------------------------------------------------------------------------------------------------------------UGACUUAUCUCAGCACCCU-------     1        19
Library = 2         Precision: [ total = 97.6% | 5p-arm = 97.8% | 3p-arm = 87.5% ]
----------UGCUGUGAGAUUGGUCACCGAGC-------------------------------------------------------------------------------------------------------------------     1762     23
----------UGCUGUGAGAUUGGUCACCG----------------------------------------------------------------------------------------------------------------------     1553     20
----------UGCUGUGAGAUUGGUCACCGAG--------------------------------------------------------------------------------------------------------------------     674      22
----------UGCUGUGAGAUUGGUCACCGA---------------------------------------------------------------------------------------------------------------------     438      21
----------------------------------------------------------------------------------------------------------------------UCGUUGACUUAUCUCAGCACC---------     77       21
```

#### *A. subglobosum* asu-mir-1208 continued

```
(((...((.(((((..(((((.(((((.((((.((.((((.....((((((((..(((((....)))))))))))))...((((.((((...........)))).)))))))).)).)))).))))).)))))))))).))..))).  -68.63 kcal/mol
GCGACUAGUGuGCUGUGAGAUUGGUCACCGAGcUGAUAGUAAGAAGGAGGUUGACGCACUGUAGAGUGCCAGUCUCCAACAUAACCAACAUUUAAUCUCAGUUGUUUAUACUACCAUCuCGUUGACUUAUCUCAGCACCcCUUACGCG  count  length
----------UGCUGUGAGAUUGGUCAC-----------------------------------------------------------------------------------------------------------------------  68  18
----------UGCUGUGAGAUUGGUCACC----------------------------------------------------------------------------------------------------------------------  61  19
-----------GCUGUGAGAUUGGUCACCGAGC------------------------------------------------------------------------------------------------------------------  38  22
-----------GCUGUGAGAUUGGUCACCGAG-------------------------------------------------------------------------------------------------------------------  22  21
------------CUGUGAGAUUGGUCACCGAGCUG----------------------------------------------------------------------------------------------------------------  13  23
---------------UGAGAUUGGUCACCGAGCUG----------------------------------------------------------------------------------------------------------------  10  20
--------------------------------------------------------------------------------------------------------------------UCUCGUUGACUUAUCUCAGC-----------  8  20
----------UGCUGUGAGAUUGGUCACCGAGCU-----------------------------------------------------------------------------------------------------------------  7  24
----------UGCUGUGAGAUUGGUCACCGAGCUG----------------------------------------------------------------------------------------------------------------  6  25
-----------GCUGUGAGAUUGGUCACCG---------------------------------------------------------------------------------------------------------------------  5  19
-------------UGUGAGAUUGGUCACCGAGCUG----------------------------------------------------------------------------------------------------------------  5  22
-----------GCUGUGAGAUUGGUCACCGA--------------------------------------------------------------------------------------------------------------------  3  20
--------------------------------------------------------------------------------------------------------------------UCUCGUUGACUUAUCUCAGCA----------  2  21
----------UGCUGUGAGAUUGGUCACCGAGCUGA---------------------------------------------------------------------------------------------------------------  1  26
-----------GCUGUGAGAUUGGUCACCGAGCU-----------------------------------------------------------------------------------------------------------------  1  23
-----------GCUGUGAGAUUGGUCACCGAGCUG----------------------------------------------------------------------------------------------------------------  1  24
------------CUGUGAGAUUGGUCACCGAG-------------------------------------------------------------------------------------------------------------------  1  20
-------------UGUGAGAUUGGUCACCGAGC------------------------------------------------------------------------------------------------------------------  1  20
--------------------------------------------------------------------------------------------------------------------UCUCGUUGACUUAUCUCAG------------  1  19
Library = 3      Precision: [ total = 97.7% | 5p-arm = 97.8% | 3p-arm = 95.7% ]
----------UGCUGUGAGAUUGGUCACCGAGC------------------------------------------------------------------------------------------------------------------  485  23
----------UGCUGUGAGAUUGGUCACCG---------------------------------------------------------------------------------------------------------------------  204  20
----------UGCUGUGAGAUUGGUCACCGAG-------------------------------------------------------------------------------------------------------------------  185  22
----------UGCUGUGAGAUUGGUCACCGA--------------------------------------------------------------------------------------------------------------------  82  21
----------------------------------------------------------------------------------------------------------------------UCGUUGACUUAUCUCAGCACC--------  22  21
-----------GCUGUGAGAUUGGUCACCGAGC------------------------------------------------------------------------------------------------------------------  11  22
----------UGCUGUGAGAUUGGUCAC-----------------------------------------------------------------------------------------------------------------------  7  18
----------UGCUGUGAGAUUGGUCACC----------------------------------------------------------------------------------------------------------------------  5  19
------------CUGUGAGAUUGGUCACCGAGCUG----------------------------------------------------------------------------------------------------------------  5  23
----------UGCUGUGAGAUUGGUCACCGAGCU-----------------------------------------------------------------------------------------------------------------  4  24
----------UGCUGUGAGAUUGGUCACCGAGCUG----------------------------------------------------------------------------------------------------------------  2  25
-----------GCUGUGAGAUUGGUCACCGAG-------------------------------------------------------------------------------------------------------------------  2  21
-----------GCUGUGAGAUUGGUCACCG---------------------------------------------------------------------------------------------------------------------  2  19
-----------GCUGUGAGAUUGGUCACCGA--------------------------------------------------------------------------------------------------------------------  1  20
---------------UGAGAUUGGUCACCGAGC------------------------------------------------------------------------------------------------------------------  1  18
--------------------------------------------------------------------------------------------------------------------UCUCGUUGACUUAUCUCAGC-----------  1  20
```

```
    ACU  U     GU     U     C    c  A    AAGAA         AC    G
 GCG   AG GuGCU  GAGAU GGUCA CGAG UG UAGU     GGAGGUUG  GCACU U
 CGC   UC CACGA--CUCUA UCAGU GCuC AC AUCA-\   /CCUCUGAC--CGUGA A
G   AU-  c             U     U    U  C      UA                G
                                            AU C    AUUUA
                                            UAA CAAC     A
                                            \-U GUUG     /
                                               U    ACUCU
```

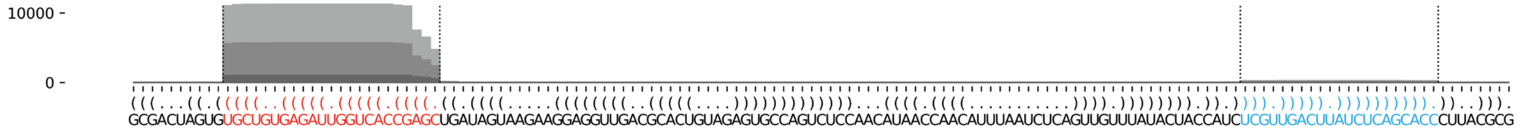

#### *A. subglobosum* asu-mir-1209

```
......(((...((((((((((((((((((((((((((.((((.(((.......((.((((((((...)))))))))))))..)))).))).)))))))))))))))))))))))...)))......  -73.19 kcal/mol
AUUGUUCAUUuGAACGAUUUUCACCAAAAUcAGGGAAUGGUCACUGGUUACUCAAGAUGUUCGGUCCGACCGAACACUCCGAUUGACGAUUCCCCUGAuUUUGGUGAAAAUCGUUAAUaUGUCGUAU  count  length
Libraries combined         Precision: [ total = 99.9% | 5p-arm = 99.9% | 3p-arm = 82.4% ]
----------UGAACGAUUUUCACCAAAAUC------------------------------------------------------------------------------------------------  19394  21
----------UGAACGAUUUUCACCAAAAU-------------------------------------------------------------------------------------------------  555    20
----------UGAACGAUUUUCACCAAAAUCA-----------------------------------------------------------------------------------------------  11     22
--------------------------------------------------------------------------------------------------UUUUGGUGAAAAUCGUUAAUA--------  14     21
-----------GAACGAUUUUCACCAAAAUC------------------------------------------------------------------------------------------------  14     20
--------UUUGAACGAUUUUCACCAAAAU-------------------------------------------------------------------------------------------------  1      22
---------UUGAACGAUUUUCACCAAAAUC------------------------------------------------------------------------------------------------  1      22
----------UGAACGAUUUUCACCAAAA--------------------------------------------------------------------------------------------------  4      19
-------------------------------AGGGAAUGGUCACUGGUUACUCAAG-----------------------------------------------------------------------  1      25
-------------------------------AGGGAAUGGUCACUGGUUACUC--------------------------------------------------------------------------  1      22
----------------------------------------------------CAAGAUGUUCGGUCCGACCGA------------------------------------------------------  1      21
--------------------------------------------------------------------------------------CGAUUCCCCUGAUUUUGGU----------------------  1      19
--------UUUGAACGAUUUUCACCAAAA--------------------------------------------------------------------------------------------------  1      21
---------UUGAACGAUUUUCACCAAAAU-------------------------------------------------------------------------------------------------  1      21
-------------------------------AGGGAAUGGUCACUGGUUACUCAAGAUGUUC-----------------------------------------------------------------  4      31
-------------------------------------------------------------CGGUCCGACCGAACACUCCGA---------------------------------------------  1      21
---------------------------------------------------------------------------------------------------UUUGGUGAAAAUCGUUAAUA--------  1      20
Library = 1        Precision: [ total = 99.9% | 5p-arm = 99.9% | 3p-arm = 85.7% ]
----------UGAACGAUUUUCACCAAAAUC------------------------------------------------------------------------------------------------  9493   21
----------UGAACGAUUUUCACCAAAAU-------------------------------------------------------------------------------------------------  276    20
----------UGAACGAUUUUCACCAAAAUCA-----------------------------------------------------------------------------------------------  6      22
--------------------------------------------------------------------------------------------------UUUUGGUGAAAAUCGUUAAUA--------  6      21
-----------GAACGAUUUUCACCAAAAUC------------------------------------------------------------------------------------------------  5      20
--------UUUGAACGAUUUUCACCAAAAU-------------------------------------------------------------------------------------------------  1      22
---------UUGAACGAUUUUCACCAAAAUC------------------------------------------------------------------------------------------------  1      22
----------UGAACGAUUUUCACCAAAA--------------------------------------------------------------------------------------------------  1      19
-------------------------------AGGGAAUGGUCACUGGUUACUCAAG-----------------------------------------------------------------------  1      25
-------------------------------AGGGAAUGGUCACUGGUUACUC--------------------------------------------------------------------------  1      22
----------------------------------------------------CAAGAUGUUCGGUCCGACCGA------------------------------------------------------  1      21
--------------------------------------------------------------------------------------CGAUUCCCCUGAUUUUGGU----------------------  1      19
Library = 2        Precision: [ total = 99.8% | 5p-arm = 99.8% | 3p-arm = 88.9% ]
----------UGAACGAUUUUCACCAAAAUC------------------------------------------------------------------------------------------------  8160   21
----------UGAACGAUUUUCACCAAAAU-------------------------------------------------------------------------------------------------  253    20
-----------GAACGAUUUUCACCAAAAUC------------------------------------------------------------------------------------------------  9      20
--------------------------------------------------------------------------------------------------UUUUGGUGAAAAUCGUUAAUA--------  8      21
----------UGAACGAUUUUCACCAAAAUCA-----------------------------------------------------------------------------------------------  4      22
-------------------------------AGGGAAUGGUCACUGGUUACUCAAGAUGUUC-----------------------------------------------------------------  3      31
---------UUGAACGAUUUUCACCAAAAU-------------------------------------------------------------------------------------------------  1      21
----------UGAACGAUUUUCACCAAAA--------------------------------------------------------------------------------------------------  1      19
-------------------------------------------------------------CGGUCCGACCGAACACUCCGA---------------------------------------------  1      21
Library = 3        Precision: [ total = 99.8% | 5p-arm = 99.9% | 3p-arm = 0% ]
----------UGAACGAUUUUCACCAAAAUC------------------------------------------------------------------------------------------------  1741   21
----------UGAACGAUUUUCACCAAAAU-------------------------------------------------------------------------------------------------  26     20
----------UGAACGAUUUUCACCAAAA--------------------------------------------------------------------------------------------------  2      19
--------UUUGAACGAUUUUCACCAAAA--------------------------------------------------------------------------------------------------  1      21
----------UGAACGAUUUUCACCAAAAUCA-----------------------------------------------------------------------------------------------  1      22
-------------------------------AGGGAAUGGUCACUGGUUACUCAAGAUGUUC-----------------------------------------------------------------  1      31
---------------------------------------------------------------------------------------------------UUUGGUGAAAAUCGUUAAUA--------  1      20
```

```
AUUGUU    UuG                        G    C    UUACUCA  A         C
      CAU    AACGAUUUUCACCAAAAUcAGGG-AAU GUCA -UGG       AG UGUUCGGU C
      GUa    UUGCUAAAAGUGGUUUuAGUCCC UUA CAGU GCC-------UC-ACAAGCCA /
UAUGCU    UAA                      C    G    UA                    G
```

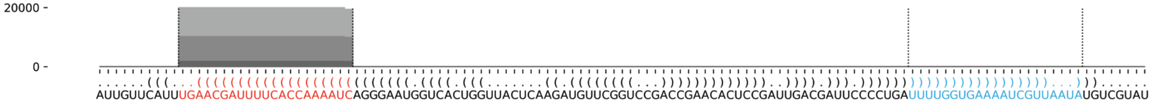

##### *A. subglobosum* asu-mir-1210

```
(((((((.(..((((.((((((((((.(((((..((((((((((.......((((((((...))))))))).....))))))))))..)))))...)))))))))).))))..).)))))))       -76.03 kcal/mol
GAUGAUGCGUuGAGCCGUACCAUUCUCUGUcUCUGUACAGAGUGCCCAUCUGGCAAGUGGACUACUUGCCCAUUACACUCUGUACUUGGAcACUUAGGAUGGUGCUGCUUCUcACAUUAUU       count    length
Libraries combined          Precision: [ total = 95.8% | 5p-arm = 96.2% | 3p-arm = 95.4% ]
----------UGAGCCGUACCAUUCUCUGUCU-----------------------------------------------------------------------------------------       116      22
------------------------------------------------------------------------------------------CACUUAGGAUGGUGCUGCUUC----------       92       21
------------------------------------------------------------------------------------------CACUUAGGAUGGUGCUGCUU-----------       59       20
----------UGAGCCGUACCAUUCUCUGUC------------------------------------------------------------------------------------------       67       21
------------------------------------------------------------------------------------------CACUUAGGAUGGUGCUGCUUCU---------       26       22
------------------------------------------------------------------------------------------CACUUAGGAUGGUGCUGCUUCUC--------       18       23
----------UGAGCCGUACCAUUCUCUG--------------------------------------------------------------------------------------------       16       19
------------------------------------------------------------------------------------------CACUUAGGAUGGUGCUGCU------------       9        19
------------AGCCGUACCAUUCUCUGUC------------------------------------------------------------------------------------------       3        19
------------AGCCGUACCAUUCUCUGUCU-----------------------------------------------------------------------------------------       5        20
----------UGAGCCGUACCAUUCUCUGU-------------------------------------------------------------------------------------------       4        20
--------------------------------------------------------------------------------UGUACUUGGACACUUAGGAUGG-------------------       6        22
-------------------------------------------------------------------------------CUGUACUUGGACACUUAGGAU---------------------       1        21
------------------------------------------------------------------------------------------CACUUAGGAUGGUGCUGC-------------       3        18
-------------------------------------------------------------------------------------------ACUUAGGAUGGUGCUGCUUC----------       1        20
--------------------------------------------------------------------------------UGUACUUGGACACUUAGGAUG--------------------       1        21
--------------------------------------------------------------------------------------------CUUAGGAUGGUGCUGCUUCUC--------       1        21
Library = 1        Precision: [ total = 95.1% | 5p-arm = 94.0% | 3p-arm = 96.2% ]
----------UGAGCCGUACCAUUCUCUGUCU-----------------------------------------------------------------------------------------       51       22
------------------------------------------------------------------------------------------CACUUAGGAUGGUGCUGCUUC----------       41       21
------------------------------------------------------------------------------------------CACUUAGGAUGGUGCUGCUU-----------       36       20
----------UGAGCCGUACCAUUCUCUGUC------------------------------------------------------------------------------------------       34       21
------------------------------------------------------------------------------------------CACUUAGGAUGGUGCUGCUUCU---------       9        22
------------------------------------------------------------------------------------------CACUUAGGAUGGUGCUGCUUCUC--------       9        23
----------UGAGCCGUACCAUUCUCUG--------------------------------------------------------------------------------------------       7        19
------------------------------------------------------------------------------------------CACUUAGGAUGGUGCUGCU------------       4        19
------------AGCCGUACCAUUCUCUGUC------------------------------------------------------------------------------------------       3        19
------------AGCCGUACCAUUCUCUGUCU-----------------------------------------------------------------------------------------       3        20
----------UGAGCCGUACCAUUCUCUGU-------------------------------------------------------------------------------------------       2        20
--------------------------------------------------------------------------------UGUACUUGGACACUUAGGAUGG-------------------       2        22
-------------------------------------------------------------------------------CUGUACUUGGACACUUAGGAU---------------------       1        21
------------------------------------------------------------------------------------------CACUUAGGAUGGUGCUGC-------------       1        18
-------------------------------------------------------------------------------------------ACUUAGGAUGGUGCUGCUUC----------       1        20
Library = 2        Precision: [ total = 96.0% | 5p-arm = 97.6% | 3p-arm = 94.4% ]
----------UGAGCCGUACCAUUCUCUGUCU-----------------------------------------------------------------------------------------       51       22
------------------------------------------------------------------------------------------CACUUAGGAUGGUGCUGCUUC----------       38       21
----------UGAGCCGUACCAUUCUCUGUC------------------------------------------------------------------------------------------       24       21
------------------------------------------------------------------------------------------CACUUAGGAUGGUGCUGCUU-----------       19       20
------------------------------------------------------------------------------------------CACUUAGGAUGGUGCUGCUUCU---------       13       22
------------------------------------------------------------------------------------------CACUUAGGAUGGUGCUGCUUCUC--------       7        23
----------UGAGCCGUACCAUUCUCUG--------------------------------------------------------------------------------------------       6        19
------------------------------------------------------------------------------------------CACUUAGGAUGGUGCUGCU------------       5        19
--------------------------------------------------------------------------------UGUACUUGGACACUUAGGAUGG-------------------       4        22
----------UGAGCCGUACCAUUCUCUGU-------------------------------------------------------------------------------------------       2        20
------------AGCCGUACCAUUCUCUGUCU-----------------------------------------------------------------------------------------       2        20
------------------------------------------------------------------------------------------CACUUAGGAUGGUGCUGC-------------       2        18
--------------------------------------------------------------------------------------------CUUAGGAUGGUGCUGCUUCUC--------       1        21
Library = 3        Precision: [ total = 98.0% | 5p-arm = 100.0% | 3p-arm = 95.8% ]
----------UGAGCCGUACCAUUCUCUGUCU-----------------------------------------------------------------------------------------       14       22
------------------------------------------------------------------------------------------CACUUAGGAUGGUGCUGCUUC----------       13       21
----------UGAGCCGUACCAUUCUCUGUC------------------------------------------------------------------------------------------       9        21
------------------------------------------------------------------------------------------CACUUAGGAUGGUGCUGCUU-----------       4        20
------------------------------------------------------------------------------------------CACUUAGGAUGGUGCUGCUUCU---------       4        22
```

#### *A. subglobosum* asu-mir-1210 continued

```
(((((((.(..((((.((((((((((.(((((..((((((((((.......((((((((...)))))))).....))))))))))..)))))...)))))))))).))))..).)))))))   -76.03 kcal/mol
GAUGAUGCGUuGAGCCGUACCAUUCUCUGUcUCUGUACAGAGUGCCCAUCUGGCAAGUGGACUACUUGCCCAUUACACUCUGUACUUGGAcACUUAGGAUGGUGCUGCUUCUcACAUUAUU   count  length
----------UGAGCCGUACCAUUCUCUG--------------------------------------------------------------------------------------------   3      19
------------------------------------------------------------------------------------------CACUUAGGAUGGUGCUGCUUCUC--------   2      23
--------------------------------------------------------------------------------UGUACUUGGACACUUAGGAUG--------------------   1      21
```

```
       C Uu    C          C       CU          CCCAUCU        G
GAUGAUG G  GAGC GUACCAUUCU --UGUcU  GUACAGAGUG       GGCAAGUG A
UUAUUAC c  UUCG CGUGGUAGGA   AcAGG  CAUGUCUCAC     --CCGUUCAU /
       A UC    U          UUC     UU          AUUAC          C
```

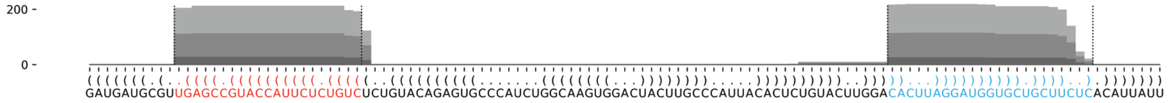


#### *A. subglobosum* asu-mir-1211

```
....((((((((((..(((.((((((((((((((.(((((((((((.(.......).)).))))))))).)))))))))))))))))...))))))))))....   -55.45 kcal/mol
UCAUUAAGAAcCCGAUACAUAUGUCAUAUUGaUAUAAUGAAUAGGACGACUUUAACUUCAUUGUUCAUUCUAUCaAUAUGACAUUGUUAACGGGUuCUUACUUG   count  length
Libraries combined         Precision: [ total = 99.9% | 5p-arm = 66.7% | 3p-arm = 99.9% ]
--------------------------------------------------------------------------AAUAUGACAUUGUUAACGGGU---------   3469   21
--------------------------------------------------------------------------AAUAUGACAUUGUUAACGGG----------   152    20
--------------------------------------------------------------------------AAUAUGACAUUGUUAACGGGUU--------   161    22
--------------------------------------------------------------------------AAUAUGACAUUGUUAACGG-----------   43     19
--------------------------------------------------------------------------AAUAUGACAUUGUUAACG------------   5      18
----------CCCGAUACAUAUGUCAUAUUG-------------------------------------------------------------------------   4      21
-----------CCGAUACAUAUGUCAUAUUGA------------------------------------------------------------------------   2      21
-------------------------------------------------------------------------CAAUAUGACAUUGUUAACGGGU---------   1      22
--------------------------------------------------------------------------AAUAUGACAUUGUUAACGGGUUC-------   1      23
---------------------------------------------------------------------------AUAUGACAUUGUUAACGGGU---------   1      20
Library = 1        Precision: [ total = 99.9% | 5p-arm = 50.0% | 3p-arm = 100.0% ]
--------------------------------------------------------------------------AAUAUGACAUUGUUAACGGGU---------   1677   21
--------------------------------------------------------------------------AAUAUGACAUUGUUAACGGG----------   81     20
--------------------------------------------------------------------------AAUAUGACAUUGUUAACGGGUU--------   79     22
--------------------------------------------------------------------------AAUAUGACAUUGUUAACGG-----------   21     19
--------------------------------------------------------------------------AAUAUGACAUUGUUAACG------------   3      18
----------CCCGAUACAUAUGUCAUAUUG-------------------------------------------------------------------------   1      21
-----------CCGAUACAUAUGUCAUAUUGA------------------------------------------------------------------------   1      21
Library = 2        Precision: [ total = 99.9% | 5p-arm = 75.0% | 3p-arm = 100.0% ]
--------------------------------------------------------------------------AAUAUGACAUUGUUAACGGGU---------   1490   21
--------------------------------------------------------------------------AAUAUGACAUUGUUAACGGGUU--------   74     22
--------------------------------------------------------------------------AAUAUGACAUUGUUAACGGG----------   62     20
--------------------------------------------------------------------------AAUAUGACAUUGUUAACGG-----------   20     19
----------CCCGAUACAUAUGUCAUAUUG-------------------------------------------------------------------------   3      21
--------------------------------------------------------------------------AAUAUGACAUUGUUAACG------------   2      18
-----------CCGAUACAUAUGUCAUAUUGA------------------------------------------------------------------------   1      21
--------------------------------------------------------------------------AAUAUGACAUUGUUAACGGGUUC-------   1      23
Library = 3        Precision: [ total = 99.4% | 5p-arm = 0% | 3p-arm = 99.4% ]
--------------------------------------------------------------------------AAUAUGACAUUGUUAACGGGU---------   302    21
--------------------------------------------------------------------------AAUAUGACAUUGUUAACGGG----------   9      20
--------------------------------------------------------------------------AAUAUGACAUUGUUAACGGGUU--------   8      22
--------------------------------------------------------------------------AAUAUGACAUUGUUAACGG-----------   2      19
-------------------------------------------------------------------------CAAUAUGACAUUGUUAACGGGU---------   1      22
---------------------------------------------------------------------------AUAUGACAUUGUUAACGGGU---------   1      20
```

```
UCAU          AU   U              U            C ACU
    UAAGAAcCCG  ACA AUGUCAUAUUGaUA AAUGAAUAG-GA G   U
    AUUCuUGGGC    UGUUACAGUAUAaCUAU UUACUUGUU CU C   /
GUUC          AAU                 C          A  U AAU
```

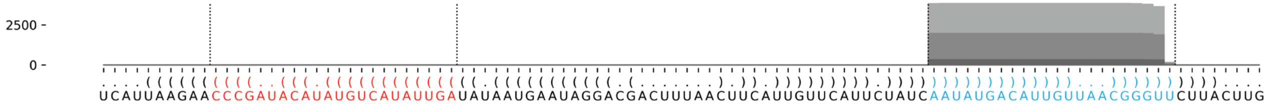

#### *A. subglobosum* asu-mir-1212

```
..((((.((((.(((((..(((((.((((((((.(((((..(((((............)))))...........))))))))))))).)))))..))))).)))).))))..   -51.1 kcal/mol
GUAUAUAUCAuUCAGUAUUCCAUCUGUGAUgUAUAUUGGUGCAGUCACAUUUCUGAUCGACUGGCCAUACUUCUCCAAUUACAuCACCGAUGGCCUACUGCAUgAGAUGUUC   count   length
Libraries combined         Precision: [ total = 99.4% | 5p-arm = 99.7% | 3p-arm = 99.1% ]
-----------------------------------------------------------------------------------UCACCGAUGGCCUACUGCAUG--------   962     21
----------UUCAGUAUUCCAUCUGUGAUG---------------------------------------------------------------------------------   916     21
-----------------------------------------------------------------------------------UCACCGAUGGCCUACUGCA----------   23      19
-----------------------------------------------------------------------------------UCACCGAUGGCCUACUGCAU---------   15      20
-------------------------------------------------------------------------------------ACCGAUGGCCUACUGCAUGAG------   3       21
----------UUCAGUAUUCCAUCUGUGA-----------------------------------------------------------------------------------   2       19
-----------UCAGUAUUCCAUCUGUGAUG---------------------------------------------------------------------------------   2       20
----------------AUUCCAUCUGUGAUGUAUAUUG--------------------------------------------------------------------------   1       22
---------------------------------------------------------------------------------CAUCACCGAUGGCCUACUG------------   1       19
---------------------------------------------------------------------------------CAUCACCGAUGGCCUACUGCA----------   1       21
------------------------------------------------------------------------------------------UGGCCUACUGCAUGAGAUGUUC   1       22
-----------------------------------------------------UGAUCGACUGGCCAUACUUCU--------------------------------------   1       21
------------------------------------------------------------------------------UUACAUCACCGAUGGCCUACU-------------   1       21
-----------------------------------------------------------------------------------UCACCGAUGGCCUACUGCAUGAGAUGU--   1       27
-----------------------------------------------------------------------------------UCACCGAUGGCCUACUGC-----------   1       18
-----------------------------------------------------------------------------------UCACCGAUGGCCUACUGCAUGA-------   1       22
Library = 1        Precision: [ total = 99.3% | 5p-arm = 99.6% | 3p-arm = 99.0% ]
-----------------------------------------------------------------------------------UCACCGAUGGCCUACUGCAUG--------   474     21
----------UUCAGUAUUCCAUCUGUGAUG---------------------------------------------------------------------------------   449     21
-----------------------------------------------------------------------------------UCACCGAUGGCCUACUGCA----------   11      19
-----------------------------------------------------------------------------------UCACCGAUGGCCUACUGCAU---------   8       20
-------------------------------------------------------------------------------------ACCGAUGGCCUACUGCAUGAG------   2       21
----------UUCAGUAUUCCAUCUGUGA-----------------------------------------------------------------------------------   1       19
-----------UCAGUAUUCCAUCUGUGAUG---------------------------------------------------------------------------------   1       20
----------------AUUCCAUCUGUGAUGUAUAUUG--------------------------------------------------------------------------   1       22
---------------------------------------------------------------------------------CAUCACCGAUGGCCUACUG------------   1       19
---------------------------------------------------------------------------------CAUCACCGAUGGCCUACUGCA----------   1       21
------------------------------------------------------------------------------------------UGGCCUACUGCAUGAGAUGUUC   1       22
Library = 2        Precision: [ total = 99.4% | 5p-arm = 99.8% | 3p-arm = 99.0% ]
----------UUCAGUAUUCCAUCUGUGAUG---------------------------------------------------------------------------------   398     21
-----------------------------------------------------------------------------------UCACCGAUGGCCUACUGCAUG--------   398     21
-----------------------------------------------------------------------------------UCACCGAUGGCCUACUGCA----------   12      19
-----------------------------------------------------------------------------------UCACCGAUGGCCUACUGCAU---------   5       20
----------UUCAGUAUUCCAUCUGUGA-----------------------------------------------------------------------------------   1       19
-----------UCAGUAUUCCAUCUGUGAUG---------------------------------------------------------------------------------   1       20
-----------------------------------------------------UGAUCGACUGGCCAUACUUCU--------------------------------------   1       21
------------------------------------------------------------------------------UUACAUCACCGAUGGCCUACU-------------   1       21
-----------------------------------------------------------------------------------UCACCGAUGGCCUACUGCAUGAGAUGU--   1       27
-----------------------------------------------------------------------------------UCACCGAUGGCCUACUGC-----------   1       18
-----------------------------------------------------------------------------------UCACCGAUGGCCUACUGCAUGA-------   1       22
-------------------------------------------------------------------------------------ACCGAUGGCCUACUGCAUGAG------   1       21
Library = 3        Precision: [ total = 100.0% | 5p-arm = 100.0% | 3p-arm = 100.0% ]
-----------------------------------------------------------------------------------UCACCGAUGGCCUACUGCAUG--------   90      21
----------UUCAGUAUUCCAUCUGUGAUG---------------------------------------------------------------------------------   69      21
-----------------------------------------------------------------------------------UCACCGAUGGCCUACUGCAU---------   2       20
```

```
GU     A     U     UU     U           U     UG---------      ACAUU
  AUAU  UCAu  CAGUA  CCAUC  GUGAUgUA  AUUGG            CAGUC     U
  UGUA  AgUA  GUCAU  GGUAG  CACuACAU-UAACC            GUCAG     C
CU     G     C     CC     C                 UCUUCAUACCG      CUAGU
```

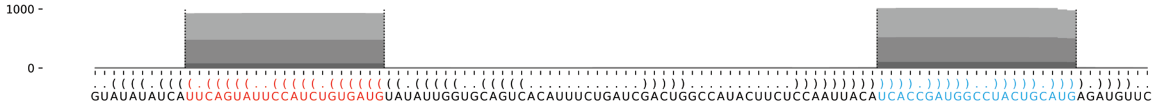

#### *A. subglobosum* asu-mir-1213

```
.....(((((.((((((((((((((((((((((..(((((.(.(((............))).).)))))..)))))))))))))))))))))).))))).....   -67.09 kcal/mol
AGUCAGGACAaCAGUCUAACAUGGUAGCAUuAUACCAAACUAUUGCGAGUAACGAAGGGCAUUCGUUUGUAAUAAuGCUACCAUGUUAGACUGAUgUCCAAUCU   count   length
Libraries combined          Precision: [ total = 96.7% | 5p-arm = 91.9% | 3p-arm = 97.9% ]
---------------------------------------------------------------------------UGCUACCAUGUUAGACUGAUG--------   15455   21
----------ACAGUCUAACAUGGUAGCAUU-------------------------------------------------------------------------   3701    21
---------------------------------------------------------------------------UGCUACCAUGUUAGACUGA----------   321     19
--------------------------------------------------------------------------AUGCUACCAUGUUAGACUGAU---------   222     21
---------------------------------------------------------------------------UGCUACCAUGUUAGACUGAUGU-------   220     22
---------AACAGUCUAACAUGGUAGCAUU-------------------------------------------------------------------------   171     22
---------------------------------------------------------------------------UGCUACCAUGUUAGACUGAU---------   147     20
----------------------------------------------------------------------------GCUACCAUGUUAGACUGAUG--------   67      20
---------AACAGUCUAACAUGGUAGCAU--------------------------------------------------------------------------   54      21
-----------CAGUCUAACAUGGUAGCAUUA------------------------------------------------------------------------   47      21
-----------CAGUCUAACAUGGUAGCAUU-------------------------------------------------------------------------   28      20
----------ACAGUCUAACAUGGUAGCAU--------------------------------------------------------------------------   17      20
---------------------------------------------------------------------------UGCUACCAUGUUAGACUG-----------   28      18
----------------------------------------------------------------------------GCUACCAUGUUAGACUGAUGU-------   24      21
-----------------------------------------------------------------------------CUACCAUGUUAGACUGAUGUC------   13      21
---------AACAGUCUAACAUGGUAGC----------------------------------------------------------------------------   9       19
--------------------------------------------------------------------------AUGCUACCAUGUUAGACUGA----------   4       20
---------AACAGUCUAACAUGGUAGCAUUA------------------------------------------------------------------------   2       23
-----------CAGUCUAACAUGGUAGCAUUAU-----------------------------------------------------------------------   2       22
--------------------------------------------------------------------------AUGCUACCAUGUUAGACU------------   3       18
------------------------------------------------------------------------------UACCAUGUUAGACUGAUG--------   4       18
--------CAACAGUCUAACAUGGUAGCA---------------------------------------------------------------------------   2       21
---------AACAGUCUAACAUGGUAGCA---------------------------------------------------------------------------   8       20
----------ACAGUCUAACAUGGUAGCA---------------------------------------------------------------------------   3       19
----------ACAGUCUAACAUGGUAGCAUUA------------------------------------------------------------------------   7       22
------------AGUCUAACAUGGUAGCAUU-------------------------------------------------------------------------   3       19
-------------------------------------------------------------------------AAUGCUACCAUGUUAGACUGA----------   1       21
---------------------------------------------------------------------------UGCUACCAUGUUAGACUGAUGUC------   3       23
---------------------------------------------------------------------------UGCUACCAUGUUAGACUGAUGUCC-----   1       24
---------------------------------------------------------------------------UGCUACCAUGUUAGACUGAUGUCCA----   1       25
----------------------------------------------------------------------------GCUACCAUGUUAGACUGA----------   2       18
----------------------------------------------------------------------------GCUACCAUGUUAGACUGAU---------   2       19
-----------------------------------------------------------------------------CUACCAUGUUAGACUGAUG--------   2       19
------------------------------------------------------------------------------UACCAUGUUAGACUGAUGUCCAA---   1       23
------------------------------------------------------------------------------UACCAUGUUAGACUGAUGUCC-----   3       21
--------CAACAGUCUAACAUGGUAGC----------------------------------------------------------------------------   1       20
----------ACAGUCUAACAUGGUAGC----------------------------------------------------------------------------   2       18
```

```
AGUCA      a                       AC      U U    GAGUA
     GGACA CAGUCUAACAUGGUAGCAUuAU  CAAAC A UGC      A
     CCUgU GUCAGAUUGUACCAUCGuAAUA  GUUUG U ACG      C
UCUAA      A                       AU      C U    GGAAG
```

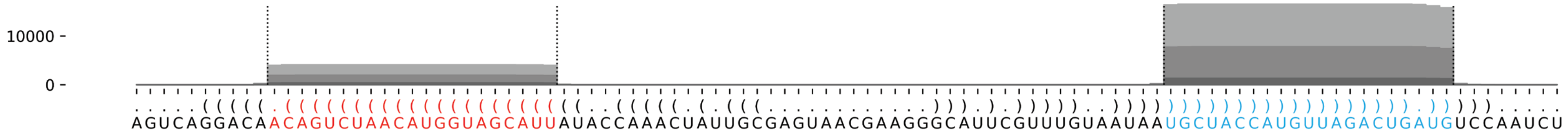

#### *A. subglobosum* asu-mir-1214

```
.(((.((.(((((((((((....((((((((((((.((((.((((..(((....))).)))))))).))))))))))))).....))))))))))).)).))).          -39.78 kcal/mol
CAUUUGUUUUaCAAAUGUUGAACAGAUAAUaAAUAUAUGAAGUAAAGAUACAAAUAUAUUGCUCGUUUAUUUAuUAUCUAUUGAAACAUUUGUGaUACCAGUU          count    length
Libraries combined          Precision: [ total = 99.5% | 5p-arm = 99.7% | 3p-arm = 98.7% ]
----------ACAAAUGUUGAACAGAUAAUA------------------------------------------------------------------------          322      21
-----------------------------------------------------------------------------UUAUCUAUUGAAACAUUUGUGA--------   66       22
-----------------------------------------------------------------------------UUAUCUAUUGAAACAUUUG-----------   5        19
----------ACAAAUGUUGAACAGAUAAU-------------------------------------------------------------------------       6        20
------------AAAUGUUGAACAGAUAAUA------------------------------------------------------------------------       1        19
-----------------------------------------------------------------------------UUAUCUAUUGAAACAUUUGUGAU-------   4        23
----------ACAAAUGUUGAACAGAUAA--------------------------------------------------------------------------       1        19
-----------------------------------------------------------------------------UUAUCUAUUGAAACAUUUGU----------   1        20
------------------------------------------------------------------------------UAUCUAUUGAAACAUUUGUGA--------   1        21
Library = 1        Precision: [ total = 99.5% | 5p-arm = 99.4% | 3p-arm = 100.0% ]
----------ACAAAUGUUGAACAGAUAAUA------------------------------------------------------------------------          157      21
-----------------------------------------------------------------------------UUAUCUAUUGAAACAUUUGUGA--------   32       22
-----------------------------------------------------------------------------UUAUCUAUUGAAACAUUUG-----------   4        19
----------ACAAAUGUUGAACAGAUAAU-------------------------------------------------------------------------       3        20
------------AAAUGUUGAACAGAUAAUA------------------------------------------------------------------------       1        19
-----------------------------------------------------------------------------UUAUCUAUUGAAACAUUUGUGAU-------   1        23
Library = 2        Precision: [ total = 99.4% | 5p-arm = 100.0% | 3p-arm = 97.1% ]
----------ACAAAUGUUGAACAGAUAAUA------------------------------------------------------------------------          130      21
-----------------------------------------------------------------------------UUAUCUAUUGAAACAUUUGUGA--------   30       22
----------ACAAAUGUUGAACAGAUAAU-------------------------------------------------------------------------       2        20
-----------------------------------------------------------------------------UUAUCUAUUGAAACAUUUGUGAU-------   2        23
----------ACAAAUGUUGAACAGAUAA--------------------------------------------------------------------------       1        19
-----------------------------------------------------------------------------UUAUCUAUUGAAACAUUUG-----------   1        19
-----------------------------------------------------------------------------UUAUCUAUUGAAACAUUUGU----------   1        20
------------------------------------------------------------------------------UAUCUAUUGAAACAUUUGUGA--------   1        21
Library = 3        Precision: [ total = 100.0% | 5p-arm = 100.0% | 3p-arm = 100.0% ]
----------ACAAAUGUUGAACAGAUAAUA------------------------------------------------------------------------          35       21
-----------------------------------------------------------------------------UUAUCUAUUGAAACAUUUGUGA--------   4        22
----------ACAAAUGUUGAACAGAUAAU-------------------------------------------------------------------------       1        20
-----------------------------------------------------------------------------UUAUCUAUUGAAACAUUUGUGAU-------   1        23
```

```
C   U  U                GAAC-               U     A    AG    C
 AUU GU UUaCAAAUGUU     AGAUAAUaAAUA AUGA GUAA  AUA A
 UGA CA aGUGUUUACAA     UCUAUUuAUUUAU UGCU-CGUU -UAU A
U   C  U               AGUUA                U          A     A
```

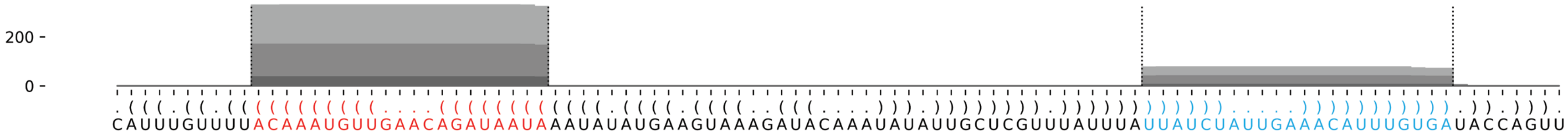

#### *A. subglobosum* asu-mir-1215

```
((((((((((((((((((.(((((((((((((((((((((((((((((((((.(((.((.......((((......))))......)).))).)))))(((((((((.((((.........)))).)))......)))).)).)))))))))))))))))))))))))))))).))))))))))))))))))   -129.41 kcal/mol
UGAGCUUGGGaAUGGCACAACUGUAAAGAGgAUAUGAAACAUGAGUUGGUCAUUCAUUCUCUCCAAGUUAUUACAUUGACGACAUUGAUUGAUUGAGUGAGCGCGCGAUACUAACCCUCGCCCGCCAAUGAUCACAUCAGACUCAUGUUUCAUAUCCuCUUUACAGUAGUGCCAUUCcCAAGCUCA   count   length
Libraries combined   Precision: [ total = 93.3% | 5p-arm = 86.6% | 3p-arm = 96.9% ]
-----------------------------------------------------------------------------------------------------------------------------------------------------------------UCUUUACAGUAGUGCCAUUCC--------   94   21
----------AAUGGCACAACUGUAAAGAGG---------------------------------------------------------------------------------------------------------------------------------------------------------------   54   21
-----------------------------------------------------------------------------------------------------------------------------------------------------------------UCUUUACAGUAGUGCCAUUC---------   24   20
-------------------------AAGAGGAUAUGAAACAUGAGU------------------------------------------------------------------------------------------------------------------------------------------------   7   21
----------AAUGGCACAACUGUAAAGAGGAU-------------------------------------------------------------------------------------------------------------------------------------------------------------   3   23
-----------------------------------------------------------------------------------------------------------------------------------------------------------------UCUUUACAGUAGUGCCAUU----------   4   19
------------------------------------------------------------------------------------------------------------------------------------------------------------------CUUUACAGUAGUGCCAUUCCC-------   2   21
-----------------------------------------------------------------------------------------------------------------------------------------------------------------UCUUUACAGUAGUGCCAU-----------   1   18
----------AAUGGCACAACUGUAAAGAGGA--------------------------------------------------------------------------------------------------------------------------------------------------------------   1   22
-----------AUGGCACAACUGUAAAGAGGA--------------------------------------------------------------------------------------------------------------------------------------------------------------   1   21
--------------------------------UAUGAAACAUGAGUUGGUCAU-----------------------------------------------------------------------------------------------------------------------------------------   1   21
----------------------------------------------------------------------------------------------------------------------------------------CACAUCAGACUCAUGUUUCAUAUC------------------------------   1   24
-------------------------------------------------------------------------------------------------------------------------------------------------------------------UUUACAGUAGUGCCAUUCC--------   1   19
Library = 1          Precision: [ total = 95.0% | 5p-arm = 93.0% | 3p-arm = 96.5% ]
-----------------------------------------------------------------------------------------------------------------------------------------------------------------UCUUUACAGUAGUGCCAUUCC--------   42   21
----------AAUGGCACAACUGUAAAGAGG---------------------------------------------------------------------------------------------------------------------------------------------------------------   38   21
-----------------------------------------------------------------------------------------------------------------------------------------------------------------UCUUUACAGUAGUGCCAUUC---------   10   20
-------------------------AAGAGGAUAUGAAACAUGAGU------------------------------------------------------------------------------------------------------------------------------------------------   3   21
----------AAUGGCACAACUGUAAAGAGGAU-------------------------------------------------------------------------------------------------------------------------------------------------------------   2   23
-----------------------------------------------------------------------------------------------------------------------------------------------------------------UCUUUACAGUAGUGCCAUU----------   2   19
------------------------------------------------------------------------------------------------------------------------------------------------------------------CUUUACAGUAGUGCCAUUCCC-------   2   21
-----------------------------------------------------------------------------------------------------------------------------------------------------------------UCUUUACAGUAGUGCCAU-----------   1   18
Library = 2          Precision: [ total = 89.5% | 5p-arm = 70.0% | 3p-arm = 96.4% ]
-----------------------------------------------------------------------------------------------------------------------------------------------------------------UCUUUACAGUAGUGCCAUUCC--------   40   21
----------AAUGGCACAACUGUAAAGAGG---------------------------------------------------------------------------------------------------------------------------------------------------------------   13   21
-----------------------------------------------------------------------------------------------------------------------------------------------------------------UCUUUACAGUAGUGCCAUUC---------   13   20
-------------------------AAGAGGAUAUGAAACAUGAGU------------------------------------------------------------------------------------------------------------------------------------------------   4   21
----------AAUGGCACAACUGUAAAGAGGA--------------------------------------------------------------------------------------------------------------------------------------------------------------   1   22
-----------AUGGCACAACUGUAAAGAGGA--------------------------------------------------------------------------------------------------------------------------------------------------------------   1   21
--------------------------------UAUGAAACAUGAGUUGGUCAU-----------------------------------------------------------------------------------------------------------------------------------------   1   21
----------------------------------------------------------------------------------------------------------------------------------------CACAUCAGACUCAUGUUUCAUAUC------------------------------   1   24
-----------------------------------------------------------------------------------------------------------------------------------------------------------------UCUUUACAGUAGUGCCAUU----------   1   19
-------------------------------------------------------------------------------------------------------------------------------------------------------------------UUUACAGUAGUGCCAUUCC--------   1   19
Library = 3          Precision: [ total = 100.0% | 5p-arm = 100.0% | 3p-arm = 100.0% ]
-----------------------------------------------------------------------------------------------------------------------------------------------------------------UCUUUACAGUAGUGCCAUUCC--------   12   21
----------AAUGGCACAACUGUAAAGAGG---------------------------------------------------------------------------------------------------------------------------------------------------------------   3   21
----------AAUGGCACAACUGUAAAGAGGAU-------------------------------------------------------------------------------------------------------------------------------------------------------------   1   23
-----------------------------------------------------------------------------------------------------------------------------------------------------------------UCUUUACAGUAGUGCCAUUC---------   1   20
-----------------------------------------------------------------------------------------------------------------------------------------------------------------UCUUUACAGUAGUGCCAUU----------   1   19
```

```
                  A                            G---    U   U  UCUCCAA    UU
UGAGCUUGGGaAUGGCAC ACUGUAAAGAGgAUAUGAAACAUGAGUU    GUCA UCA UC       GUUA  A
ACUCGAACcCUUACCGUG UGACAUUUCuCCUAUACUUUGUACUCAG\  /UAGU AGU AG       CAGU  C
                  A                             AU     U   U  UUACAG-    UA
                                                CG
                                                UA             C    UACU
                                                A GUGA------GCG GCGA    A
                                                 \CACU      CGC CGCU    /
                                                      AGUAAC   C    CCCA
```

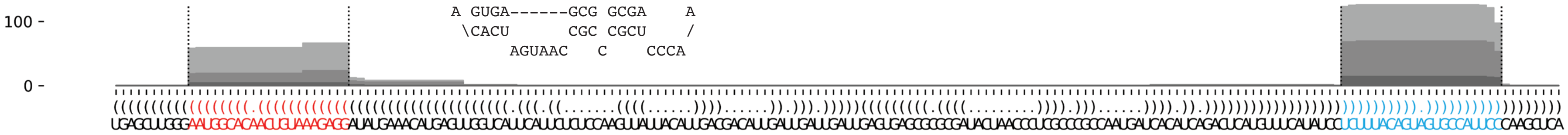

#### *A. subglobosum* asu-mir-1216

```
.....(.(((((((.((.((((((((((.(..(.(((((((((((((........))))))))))))).)..).)))))))))).)).))))))).)..... -65.75 kcal/mol
UCUACCCAGCcAAUCACCUAGCUACACCCCaCCUGGCCAAUGGCACAGCUACCUGUGUGUCAUUGGCUUGACGaGGUGUAGCUAUGUCAUUGGcUUGAUGAG count    length
Libraries combined          Precision: [ total = 99.7% | 5p-arm = 59.4% | 3p-arm = 99.7% ]
-------------------------------------------------------------------------AGGUGUAGCUAUGUCAUUGGC-------- 97242    21
-------------------------------------------------------------------------AGGUGUAGCUAUGUCAUUGG--------- 17621    20
-------------------------------------------------------------------------AGGUGUAGCUAUGUCAUUG---------- 4455     19
-------------------------------------------------------------------------AGGUGUAGCUAUGUCAUU----------- 273      18
-------------------------------------------------------------------------AGGUGUAGCUAUGUCAUUGGCU------- 196      22
--------------------------------------------------------------------------GGUGUAGCUAUGUCAUUGGC-------- 99       20
-----------------------------------------------------------------------CGAGGUGUAGCUAUGUCAUUG---------- 83       21
----------------------------------------------------CUGUGUGUCAUUGGCUUGACG----------------------------- 43       21
-------------------------------------------------------------------------AGGUGUAGCUAUGUCAUUGGCUU------ 30       23
-----------------------------------------------------------------------CGAGGUGUAGCUAUGUCAUUGG--------- 25       22
----------------------------------------------------CUGUGUGUCAUUGGCUUGA------------------------------- 17       19
-----------------------------------------------------------------------CGAGGUGUAGCUAUGUCAUU----------- 16       20
----------CAAUCACCUAGCUACACCCCA----------------------------------------------------------------------- 13       21
--------------------------------------------------------------------------GGUGUAGCUAUGUCAUUGG--------- 8        19
----------------------------------------------------CUGUGUGUCAUUGGCUUGAC------------------------------ 6        20
----------------------------------------------------CUGUGUGUCAUUGGCUUG-------------------------------- 5        18
------------------------------------------------------------------------GAGGUGUAGCUAUGUCAUUGG--------- 4        21
----------------------------------------------------------------------------UGUAGCUAUGUCAUUGGC-------- 7        18
----------CAAUCACCUAGCUACACCCC------------------------------------------------------------------------ 5        20
------------AUCACCUAGCUACACCCCACC--------------------------------------------------------------------- 2        21
-------------------------------CCUGGCCAAUGGCACAGCU---------------------------------------------------- 3        19
------------------------------------------------------------------------GAGGUGUAGCUAUGUCAUUG---------- 3        20
--------------------------------------------------------------------------GGUGUAGCUAUGUCAUUGGCU------- 4        21
---------------------------------------------------------------------------GUGUAGCUAUGUCAUUGGC-------- 2        19
--------------------------------------------------------------------------------GCUAUGUCAUUGGCUUGAU--- 2        19
---------CCAAUCACCUAGCUACACCCC------------------------------------------------------------------------ 2        21
-----------------------ACACCCCACCUGGCCAAUGGC---------------------------------------------------------- 1        21
--------------------------CCCCACCUGGCCAAUGGCAC-------------------------------------------------------- 1        20
-----------------------------------------------------UGUGUGUCAUUGGCUUGACG----------------------------- 1        20
-----------------------------------------------------UGUGUGUCAUUGGCUUGAC------------------------------ 1        19
--------------------------------------------------------------------------GGUGUAGCUAUGUCAUUG---------- 4        18
----------------------------------------------------------------------------UGUAGCUAUGUCAUUGGCUUG----- 1        21
----------CAAUCACCUAGCUACACCC------------------------------------------------------------------------- 1        19
--------------CACCUAGCUACACCCCACCUGG------------------------------------------------------------------ 2        22
----------------------UACACCCCACCUGGCCAAUGG----------------------------------------------------------- 2        21
---------------------------------------------------CCUGUGUGUCAUUGGCUUGACG----------------------------- 3        22
------------------------------------------------------------------------GAGGUGUAGCUAUGUCAUUGGC-------- 2        22
-------------------------------------------------------------------------AGGUGUAGCUAUGUCAUUGGCUUG----- 1        24
----------------------------------------------------------------------------------UAUGUCAUUGGCUUGAUG-- 1        18
```

```
UCUAC C         C   C             C  aC  U                 GCU
      C  AGCcAAU  AC  UAGCUACACC  C   C  GGCCAAUGGCACA    A
      G  UcGGUUA  UG  AUCGAUGUGG  G   G  UCGGUUACUGUGU    C
GAGUA U         C   U             a  CA  U                 GUC
```

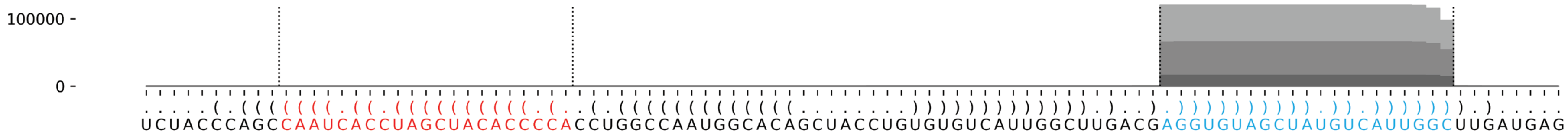

##### *A. subglobosum* asu-mir-1217

```
.......(((((((.(((.((((((((((((((((...............(((((((((....))))))))))))))))))))))))).))).)))))))....... 	-55.33 kcal/mol
UAAUUAAUGAaCGUAUCACCUAUGUCUGAUaCUCAUUCUAAAUUAAUCUUAUAUUGAUCAUUAGAUUAAUAUUGAGUAuCAGACAUAGUUGACACGUUcACUCAUUC 	count	length
Libraries combined	Precision: [ total = 98.7% | 5p-arm = 55.6% | 3p-arm = 98.7% ]
------------------------------------------------------------------------------UCAGACAUAGUUGACACGUUC-------- 	11779	21
------------------------------------------------------------------------------UCAGACAUAGUUGACACGUU--------- 	206	20
-----------------------------------------------------------------------------AUCAGACAUAGUUGACACGUU--------- 	62	21
------------------------------------------------------------------------------UCAGACAUAGUUGACACGUUCACU----- 	41	24
------------------------------------------------------------------------------UCAGACAUAGUUGACACGU---------- 	41	19
------------------------------------------------------------------------------UCAGACAUAGUUGACACGUUCACUC---- 	11	25
------------------------------------------------------------------------------UCAGACAUAGUUGACACGUUCA------- 	6	22
------------------------------------------------------------------------------UCAGACAUAGUUGACACG----------- 	9	18
---------------------------------------------------------------------------------GACAUAGUUGACACGUUC-------- 	4	18
----------ACGUAUCACCUAUGUCUGAUA---------------------------------------------------------------------------- 	8	21
-----------------------------------------------------------------------------AUCAGACAUAGUUGACACGU---------- 	4	20
--------------------------------------------------------------------------------AGACAUAGUUGACACGUUC-------- 	7	19
---------AACGUAUCACCUAUGUCUGAU----------------------------------------------------------------------------- 	6	21
----------ACGUAUCACCUAUGUCUGAU----------------------------------------------------------------------------- 	2	20
--------------------------------------------------------------------AUAUUGAGUAUCAGACAUAGU------------------ 	3	21
--------------------------------------------------------------------------AGUAUCAGACAUAGUUGACACG----------- 	6	22
--------------------------------------------------------------------------------AGACAUAGUUGACACGUUCACUC---- 	2	23
---------AACGUAUCACCUAUGUCUGA------------------------------------------------------------------------------ 	1	20
--------------------------------------------------------------------AUAUUGAGUAUCAGACAUAG------------------- 	1	20
--------------------------------------------------------------------------AGUAUCAGACAUAGUUGACAC------------ 	3	21
----------------------------------------------------------------------------UAUCAGACAUAGUUGACACGU---------- 	2	21
----------------------------------------------------------------------------UAUCAGACAUAGUUGACACGUUCACU----- 	1	26
-----------------------------------------------------------------------------AUCAGACAUAGUUGACACG----------- 	2	19
-------------------------------------------------------------------------------CAGACAUAGUUGACACGUUCA------- 	2	21
-------------------------------------------------------------------------------CAGACAUAGUUGACACGUUCACUC---- 	1	24
--------------AUCACCUAUGUCUGAUACUC------------------------------------------------------------------------- 	1	20
-------------------------------------------------------------------------------CAGACAUAGUUGACACGUUCACU----- 	1	23
-------------------------------------------------------------------------------CAGACAUAGUUGACACGUUC-------- 	1	20
UAAUUAA       A   C                UUCUAAAUUAAUCUU          A
       UGAaCGU UCA CUAUGUCUGAUaCUCA               AUAUUGAUC U
       AcUUGCA AGU GAUACAGACuAUGAGU---------------UAUAAUUAG U
CUUACUC       C   U                                         A
```

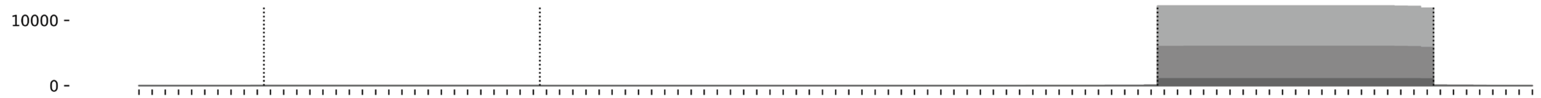


```
.......(((((((.(((.((((((((((((((((...............(((((((((....))))))))))))))))))))))))).))).)))))))).......
UAAUUAAUGAACGUAUCACCUAUGUCUGAUACUCAUUCUAAAUUAAUCUUAUAUUGAUCAUUAGAUUAAUAUUGAGUAUCAGACAUAGUUGACACGUUCACUCAUUC
```

#### *D. fasciculatum* dfa-mir-1218

```
((((..((((((((((.((((((((((((((((((((((.((((.(.((..(....(((......)))....)..))).)))))))))))))))))))))))))).)))))))))).)))).   -64.67 kcal/mol
UGGACUGUUCuUUUCACCAUUUAUCUCUUCcAAUAAGAUCAUUUUCUCUCUUUCUCUGGCUUCAACCAAAUUAUAAGGUAGAUAUCUUAUUGGaAGAGAUGAAUGCUGAAAAGaAUUUUCAA   count   length
Libraries combined         Precision: [ total = 95.3% | 5p-arm = 100.0% | 3p-arm = 94.6% ]
---------------------------------------------------------------------------------------------AAGAGAUGAAUGCUGAAAAGA--------   26   21
---------------------------------------------------------------------------------------------AAGAGAUGAAUGCUGAAAAGAA-------   7    22
----------UUUUCACCAUUUAUCUCUUCC-------------------------------------------------------------------------------------------   5    21
----------UUUUCACCAUUUAUCUCUUC--------------------------------------------------------------------------------------------   1    20
---------------------------------------------------------------------------------------UAUUGGAAGAGAUGAAUGCUG--------------   1    21
--------------------------------------------------------------------------------------------GAAGAGAUGAAUGCUGAAAAG---------   1    21
---------------------------------------------------------------------------------------------AAGAGAUGAAUGCUGAAAAGAAU------   2    23
Library = 1       Precision: [ total = 100.0% | 5p-arm = 0% | 3p-arm = 100.0% ]
---------------------------------------------------------------------------------------------AAGAGAUGAAUGCUGAAAAGA--------   9    21
---------------------------------------------------------------------------------------------AAGAGAUGAAUGCUGAAAAGAA-------   3    22
Library = 2       Precision: [ total = 90.9% | 5p-arm = 100.0% | 3p-arm = 87.5% ]
----------UUUUCACCAUUUAUCUCUUCC-------------------------------------------------------------------------------------------   3    21
---------------------------------------------------------------------------------------------AAGAGAUGAAUGCUGAAAAGA--------   3    21
---------------------------------------------------------------------------------------------AAGAGAUGAAUGCUGAAAAGAA-------   3    22
--------------------------------------------------------------------------------------------GAAGAGAUGAAUGCUGAAAAG---------   1    21
---------------------------------------------------------------------------------------------AAGAGAUGAAUGCUGAAAAGAAU------   1    23
Library = 3       Precision: [ total = 95.0% | 5p-arm = 100.0% | 3p-arm = 94.1% ]
---------------------------------------------------------------------------------------------AAGAGAUGAAUGCUGAAAAGA--------   14   21
----------UUUUCACCAUUUAUCUCUUCC-------------------------------------------------------------------------------------------   2    21
----------UUUUCACCAUUUAUCUCUUC--------------------------------------------------------------------------------------------   1    20
---------------------------------------------------------------------------------------UAUUGGAAGAGAUGAAUGCUG--------------   1    21
---------------------------------------------------------------------------------------------AAGAGAUGAAUGCUGAAAAGAA-------   1    22
---------------------------------------------------------------------------------------------AAGAGAUGAAUGCUGAAAAGAAU------   1    23
```

```
    CU          C                      C    U U  CU UCUC   CU
UGGA  GUUCuUUUCA CAUUUAUCUCUUCcAAUAAGAU AUUU C CU  U    UGG  U
 ACUU UAaGAAAAGU GUAAGUAGAGAaGGUUAUUCUA-UAGA G-GA  A    ACC  C
A    U          C                           U    AU UUAA   AA
```

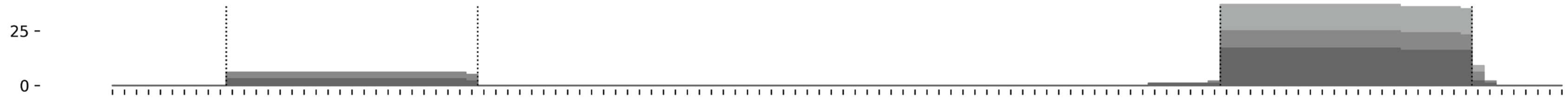


```
((((..((((((((((.((((((((((((((((((((((.((((.(.((..(....(((......)))....)..))).)))))))))))))))))))))))))).)))))))))).)))).
UGGACUGUUCUUUUCACCAUUUAUCUCUUCCAAUAAGAUCAUUUUCUCUCUUUCUCUGGCUUCAACCAAAUUAUAAGGUAGAUAUCUUAUUGGAAGAGAUGAAUGCUGAAAAGAAUUUUCAA
```

#### *D. fasciculatum* dfa-mir-1219

```
(((((((((((((((((.((((((((.((((((((.((((((((..((((...))))...)))))))).........(((.((((.....)))).)))...(((((((.(((...)))))))))))))))))).)))))))).)))))))))))))))))   -84.55 kcal/mol
UGCAAUGUUAuAAUGAUCAACGAGUUCAAUaCCUCAAAUUGAAUUCGAUUGUUGGUCUAUGUUCAAUUCUCUUUUGCCAAGCAUUUUGAUGAUGGUUGUUCACCUUUGACGUGUGGCGCAGAGGUGAGGUAuUAAACUCGUUUAUCAUUAUaACAUUGUA   count   length
Libraries combined         Precision: [ total = 92.8% | 5p-arm = 87.7% | 3p-arm = 95.8% ]
-----------------------------------------------------------------------------------------------------------------------------------UUAAACUCGUUUAUCAUUAUA--------   512   21
----------UAAUGAUCAACGAGUUCAAUA---------------------------------------------------------------------------------------------------------------------------------   250   21
----------UAAUGAUCAACGAGUUCAAU----------------------------------------------------------------------------------------------------------------------------------   21    20
----------UAAUGAUCAACGAGUUCAAUAC--------------------------------------------------------------------------------------------------------------------------------   10    22
------------AUGAUCAACGAGUUCAAUACC-------------------------------------------------------------------------------------------------------------------------------   6     21
-------------UGAUCAACGAGUUCAAUACCU------------------------------------------------------------------------------------------------------------------------------   7     21
----------------------------------------------------------------------------------------------------------UGACGUGUGGCGCAGAGGU-----------------------------------   2     19
-----------------------------------------------------------------------------------------------------------------------------------UUAAACUCGUUUAUCAUUAU---------   6     20
UGCAAUGUUAUAAUGAUCAAC-------------------------------------------------------------------------------------------------------------------------------------------   4     21
----------UAAUGAUCAACGAGUUCAA-----------------------------------------------------------------------------------------------------------------------------------   3     19
-----------AAUGAUCAACGAGUUCAAUAC--------------------------------------------------------------------------------------------------------------------------------   6     21
----------------------------------------------------------------------------------------------------------------------CAGAGGUGAGGUAUUAAACUC---------------------   5     21
---------------------------------------------------------------------------------------------------------------------------------UAUUAAACUCGUUUAUCAUUA----------   7     21
-------UUAUAAUGAUCAACGAGUUC-------------------------------------------------------------------------------------------------------------------------------------   4     20
-------UUAUAAUGAUCAACGAGUUCA------------------------------------------------------------------------------------------------------------------------------------   6     21
----------UAAUGAUCAACGAGUUCAAUACC-------------------------------------------------------------------------------------------------------------------------------   1     23
--------------GAUCAACGAGUUCAAUACCUC-----------------------------------------------------------------------------------------------------------------------------   1     21
------------------------------ACCUCAAAUUGAAUUCGAUUG-------------------------------------------------------------------------------------------------------------   1     21
------------------------------------AAUUGAAUUCGAUUGUUGGUC-------------------------------------------------------------------------------------------------------   1     21
-------------------------------------AUUGAAUUCGAUUGUUGGUCU------------------------------------------------------------------------------------------------------   1     21
--------------------------------------UUGAAUUCGAUUGUUGGUCUA-----------------------------------------------------------------------------------------------------   1     21
----------------------------------------------------------AUGUUCAAUUCUCUUUUGCCA---------------------------------------------------------------------------------   2     21
--------------------------------------------------------------------------------------------------------------------CGCAGAGGUGAGGUAUUAAAC-----------------------   2     21
----------------------------------------------------------------------------------------------------------------------CAGAGGUGAGGUAUUAAACU----------------------   1     20
-----------------------------------------------------------------------------------------------------------------------AGAGGUGAGGUAUUAAAC-----------------------   1     18
-----------------------------------------------------------------------------------------------------------------------AGAGGUGAGGUAUUAAACUC---------------------   1     20
---------------------------------------------------------------------------------------------------------------------------------UAUUAAACUCGUUUAUCAUUAU---------   1     22
---------------------------------------------------------------------------------------------------------------------------------UAUUAAACUCGUUUAUCA-------------   1     18
----------------------------------------------------------------------------------------------------------------------------------AUUAAACUCGUUUAUCAUUAU---------   1     21
-----------------------------------------------------------------------------------------------------------------------------------UUAAACUCGUUUAUCAUUA----------   8     19
-----------------------------------------------------------------------------------------------------------------------------------UUAAACUCGUUUAUCAUUAUAA-------   1     22
------------------------------------------------------------------------------------------------------------------------------------UAAACUCGUUUAUCAUUAUAAC------   1     22
Library = 1       Precision: [ total = 94.4% | 5p-arm = 92.8% | 3p-arm = 95.7% ]
-----------------------------------------------------------------------------------------------------------------------------------UUAAACUCGUUUAUCAUUAUA--------   88    21
----------UAAUGAUCAACGAGUUCAAUA---------------------------------------------------------------------------------------------------------------------------------   67    21
----------UAAUGAUCAACGAGUUCAAU----------------------------------------------------------------------------------------------------------------------------------   6     20
----------UAAUGAUCAACGAGUUCAAUAC--------------------------------------------------------------------------------------------------------------------------------   3     22
------------AUGAUCAACGAGUUCAAUACC-------------------------------------------------------------------------------------------------------------------------------   2     21
-------------UGAUCAACGAGUUCAAUACCU------------------------------------------------------------------------------------------------------------------------------   2     21
----------------------------------------------------------------------------------------------------------UGACGUGUGGCGCAGAGGU-----------------------------------   2     19
-----------------------------------------------------------------------------------------------------------------------------------UUAAACUCGUUUAUCAUUAU---------   2     20
UGCAAUGUUAUAAUGAUCAAC-------------------------------------------------------------------------------------------------------------------------------------------   1     21
----------UAAUGAUCAACGAGUUCAA-----------------------------------------------------------------------------------------------------------------------------------   1     19
-----------AAUGAUCAACGAGUUCAAUAC--------------------------------------------------------------------------------------------------------------------------------   1     21
----------------------------------------------------------------------------------------------------------------------CAGAGGUGAGGUAUUAAACUC---------------------   1     21
---------------------------------------------------------------------------------------------------------------------------------UAUUAAACUCGUUUAUCAUUA----------   1     21
Library = 2       Precision: [ total = 86.5% | 5p-arm = 77.2% | 3p-arm = 92.6% ]
-----------------------------------------------------------------------------------------------------------------------------------UUAAACUCGUUUAUCAUUAUA--------   105   21
----------UAAUGAUCAACGAGUUCAAUA---------------------------------------------------------------------------------------------------------------------------------   56    21
----------UAAUGAUCAACGAGUUCAAU----------------------------------------------------------------------------------------------------------------------------------   4     20
----------------------------------------------------------------------------------------------------------------------CAGAGGUGAGGUAUUAAACUC---------------------   4     21
-----------------------------------------------------------------------------------------------------------------------------------UUAAACUCGUUUAUCAUUA----------   4     19
------------AUGAUCAACGAGUUCAAUACC-------------------------------------------------------------------------------------------------------------------------------   3     21
```

#### *D. fasciculatum* dfa-mir-1219 continued

```
(((((((((((((((((.((((((((.((((((((.((((((((..((((...))))...)))))))).........(((.((((.....)))).)))...(((((((.(((...))))))))))))))))))).)))))))).))))))))))))))))))   -84.55 kcal/mol
UGCAAUGUUAuAAUGAUCAACGAGUUCAAUaCCUCAAAUUGAAUUCGAUUGUUGGUCUAUGUUCAAUUCUCUUUUGCCAAGCAUUUUGAUGAUGGUUGUUCACCUUUGACGUGUGGCGCAGAGGUGAGGUAuUAAACUCGUUUAUCAUUAUaACAUUGUA   count    length
UGCAAUGUUAUAAUGAUCAAC-------------------------------------------------------------------------------------------------------------------------------------------   2        21
-------UUAUAAUGAUCAACGAGUUC-------------------------------------------------------------------------------------------------------------------------------------   2        20
-----------AAUGAUCAACGAGUUCAAUAC--------------------------------------------------------------------------------------------------------------------------------   2        21
----------------------------------------------------------AUGUUCAAUUCUCUUUUGCCA---------------------------------------------------------------------------------   2        21
--------------------------------------------------------------------------------------------------------------------CGCAGAGGUGAGGUAUUAAAC-----------------------   2        21
-----------------------------------------------------------------------------------------------------------------------------------UUAAACUCGUUUAUCAUUAU--------   2        20
-------UUAUAAUGAUCAACGAGUUCA------------------------------------------------------------------------------------------------------------------------------------   1        21
----------UAAUGAUCAACGAGUUCAAUAC--------------------------------------------------------------------------------------------------------------------------------   1        22
-------------UGAUCAACGAGUUCAAUACCU------------------------------------------------------------------------------------------------------------------------------   1        21
--------------GAUCAACGAGUUCAAUACCUC-----------------------------------------------------------------------------------------------------------------------------   1        21
------------------------------ACCUCAAAUUGAAUUCGAUUG-------------------------------------------------------------------------------------------------------------   1        21
------------------------------------AAUUGAAUUCGAUUGUUGGUC-------------------------------------------------------------------------------------------------------   1        21
-------------------------------------AUUGAAUUCGAUUGUUGGUCU------------------------------------------------------------------------------------------------------   1        21
--------------------------------------UUGAAUUCGAUUGUUGGUCUA-----------------------------------------------------------------------------------------------------   1        21
-----------------------------------------------------------------------------------------------------------------------AGAGGUGAGGUAUUAAAC-----------------------   1        18
---------------------------------------------------------------------------------------------------------------------------------UAUUAAACUCGUUUAUCAUUA----------   1        21
----------------------------------------------------------------------------------------------------------------------------------AUUAAACUCGUUUAUCAUUAU---------   1        21
-----------------------------------------------------------------------------------------------------------------------------------UUAAACUCGUUUAUCAUUAUAA-------   1        22
Library = 3        Precision: [ total = 94.8% | 5p-arm = 90.2% | 3p-arm = 97.0% ]
-----------------------------------------------------------------------------------------------------------------------------------UUAAACUCGUUUAUCAUUAUA--------   319      21
----------UAAUGAUCAACGAGUUCAAUA---------------------------------------------------------------------------------------------------------------------------------   127      21
----------UAAUGAUCAACGAGUUCAAU----------------------------------------------------------------------------------------------------------------------------------   11       20
----------UAAUGAUCAACGAGUUCAAUAC--------------------------------------------------------------------------------------------------------------------------------   6        22
-------UUAUAAUGAUCAACGAGUUCA------------------------------------------------------------------------------------------------------------------------------------   5        21
---------------------------------------------------------------------------------------------------------------------------------UAUUAAACUCGUUUAUCAUUA----------   5        21
-------------UGAUCAACGAGUUCAAUACCU------------------------------------------------------------------------------------------------------------------------------   4        21
-----------------------------------------------------------------------------------------------------------------------------------UUAAACUCGUUUAUCAUUA----------   4        19
-----------AAUGAUCAACGAGUUCAAUAC--------------------------------------------------------------------------------------------------------------------------------   3        21
-------UUAUAAUGAUCAACGAGUUC-------------------------------------------------------------------------------------------------------------------------------------   2        20
----------UAAUGAUCAACGAGUUCAA-----------------------------------------------------------------------------------------------------------------------------------   2        19
-----------------------------------------------------------------------------------------------------------------------------------UUAAACUCGUUUAUCAUUAU--------   2        20
UGCAAUGUUAUAAUGAUCAAC-------------------------------------------------------------------------------------------------------------------------------------------   1        21
----------UAAUGAUCAACGAGUUCAAUACC-------------------------------------------------------------------------------------------------------------------------------   1        23
------------AUGAUCAACGAGUUCAAUACC-------------------------------------------------------------------------------------------------------------------------------   1        21
----------------------------------------------------------------------------------------------------------------------CAGAGGUGAGGUAUUAAACU----------------------   1        20
-----------------------------------------------------------------------------------------------------------------------AGAGGUGAGGUAUUAAACUC---------------------   1        20
---------------------------------------------------------------------------------------------------------------------------------UAUUAAACUCGUUUAUCAUUAU---------   1        22
---------------------------------------------------------------------------------------------------------------------------------UAUUAAACUCGUUUAUCA-------------   1        18
------------------------------------------------------------------------------------------------------------------------------------UAAACUCGUUUAUCAUUAUAAC------   1        22
```

```
                                           CU-    G
                                     /-GAAU   GAUU U
                                     UACUUG   CUGG /
                                     UA    UAU    U
                                     AU
                 C        C         AAUCUCUUUUGC    G    UU
UGCAAUGUUAuAAUGAU AACGAGUU AAUaCCUC/            \CAA CAUU   G
AUGUUACAaUAUUACUA UUGCUCAA UuAUGGAG-\     /------GUU GUAG   /
                 U        A          UACUU          G    UA
                                     GC
                                     GC   A   G
                                     AUUUG CGU U
                                     \-GAC-GCG /
                                              G
```

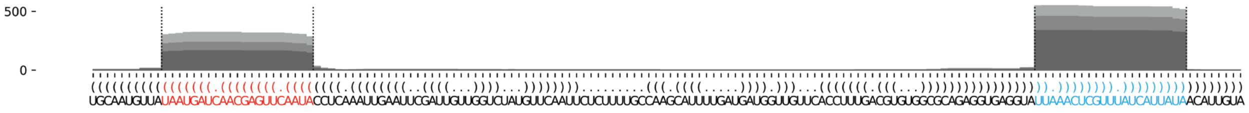

#### *D. fasciculatum* dfa-mir-1220-P1

```
.....(.((((((((.((((((((((..((.(((((((..(((((((..........((.(((((..((((((...(((..............))).))))))..))))).))........)))))))..))))))).))..)))))))))).)))))))).).....   -61.47 kcal/mol
AUACUCUAGAuCAUUCCCAUUGUCACCGAAaUCAUUAUUUAAUGAAUCACAAACCAUGCCAUUGUCAAGAAGAAACCCAACCAAUCAAUCAACUGGCUCUUUUAAAUAAUCGCUUCAUCUCAUUCAUUUUAUAAUGACUuAAGUGAUGAUGGUAAUGAUcUUGUUUUA   count     length
Libraries combined           Precision: [ total = 99.9% | 5p-arm = 95.2% | 3p-arm = 99.9% ]
-------------------------------------------------------------------------------------------------------------------------------------------UAAGUGAUGAUGGUAAUGAUC--------   77764     21
-------------------------------------------------------------------------------------------------------------------------------------------UAAGUGAUGAUGGUAAUGAU---------   31349     20
-------------------------------------------------------------------------------------------------------------------------------------------UAAGUGAUGAUGGUAAUGAUCU-------   2904      22
-------------------------------------------------------------------------------------------------------------------------------------------UAAGUGAUGAUGGUAAUGA----------   1326      19
-------------------------------------------------------------------------------------------------------------------------------------------UAAGUGAUGAUGGUAAUG-----------   620       18
-------------------------------------------------------------------------------------------------------------------------------------------UAAGUGAUGAUGGUAAUGAUCUU------   681       23
------------------------------------------------------------------------------------------------------------------------------------------UUAAGUGAUGAUGGUAAUGAUC--------   24        22
--------------------------------------------------------------------------------------------------------------------------------------------AAGUGAUGAUGGUAAUGAUC--------   10        20
----------UCAUUCCCAUUGUCACCG--------------------------------------------------------------------------------------------------------------------------------------------   7         18
-----------------------------------------------------------------------------------------------------------------------------------------CUUAAGUGAUGAUGGUAAUGA----------   7         21
-----------------------------------------------------------------------------------------------------------------------------------------CUUAAGUGAUGAUGGUAAUGAUC--------   9         23
------------------------------------------------------------------------------------------------------------------------------------------UUAAGUGAUGAUGGUAAUGAU---------   36        21
--------------------------------------------------------------------------------------------------------------------------------------------AAGUGAUGAUGGUAAUGAUCU-------   3         21
----------UCAUUCCCAUUGUCACCGAAA-----------------------------------------------------------------------------------------------------------------------------------------   13        21
----------------------------------------------------------------------------------------------------------------------------------------ACUUAAGUGAUGAUGGUAAUGAU---------   3         23
-----------------------------------------------------------------------------------------------------------------------------------------CUUAAGUGAUGAUGGUAAUGAU---------   3         22
-----------------------------------------------------------------------------------------------------------------------------------------CUUAAGUGAUGAUGGUAAUG-----------   5         20
-----------------------------------------------------------------------------------------------------------------------------------------CUUAAGUGAUGAUGGUAAUGAUCU-------   1         24
----------------------------------------------------------------------------------------------------------------------------------------------GUGAUGAUGGUAAUGAUC--------   1         18
--------------------------------------UUAAUGAAUCACAAACCAUG--------------------------------------------------------------------------------------------------------------   1         20
----------------------------------------------------------------------------------------------------------------CUUCAUCUCAUUCAUUUUAUA-----------------------------------   1         21
--------------------------------------------------------------------------------------------------------------------------------UUAUAAUGACUUAAGUGAUG--------------------   1         20
--------------------------------------------------------------------------------------------------------------------------------UUAUAAUGACUUAAGUGAU---------------------   2         19
------------------------------------------------------------------------------------------------------------------------------------AAUGACUUAAGUGAUGAUGGUAAUGAUC--------   1         28
----------------------------------------------------------------------------------------------------------------------------------------ACUUAAGUGAUGAUGGUAAUGA----------   1         22
----------------------------------------------------------------------------------------------------------------------------------------ACUUAAGUGAUGAUGGUAAUG-----------   1         21
------------------------------------------------------------------------------------------------------------------------------------------UUAAGUGAUGAUGGUAAUGAUCU-------   3         23
------------------------------------------------------------------------------------------------------------------------------------------UUAAGUGAUGAUGGUAAUG-----------   1         19
-------------------------------------------------------------------------------------------------------------------------------------------UAAGUGAUGAUGGUAAUGAUCUUGU----   1         25
--------------------------------------------------------------------------------------------------------------------------------------------AAGUGAUGAUGGUAAUGAU---------   8         19
---------------------------------------------------------------------------------------------------------------------------------------------AGUGAUGAUGGUAAUGAU---------   1         18
---------------------------------------------------------------------------------------------------------------------------------------------AGUGAUGAUGGUAAUGAUC--------   1         19
```

```
AUACU U        C          CG  a       UU       CACAAACCAU  C     CA      AAC   ACCAAU
     C AGAuCAUU CCAUUGUCAC  AA UCAUUAU  AAUGAAU          GC AUUGU  AGAAGA   CCA      C
     G UcUAGUAA GGUAGUAGUG  uU AGUAAUA  UUACUUA          CG UAAUA  UUUUCU   GGU      A
AUUUU U        U          AA  C       UU       CUCUACUU--  C     AA      C--   CAACUA
```

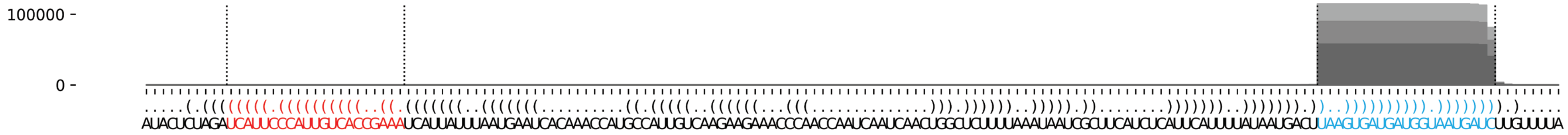

#### *D. fasciculatum* dfa-mir-1220-P2

```
.....(.((((((((.((((((((((..(((((((.(((..((((((((...((((.......((.((((.((((..((((((((....)))))))).)))).)))))).))))..)))))))).))).)))))))..)))))))))).)))))))).).....   -66.48 kcal/mol
AUACUCUAGAuCAUUCCCAUUGUCACCCAAaUCAUCAUUAUUUAUUUAUUUAUGUCUCUCAUCAGUGUGUCGACACAGUAUAUAUACAUAUAUAUAUAUGUUUACACCUCGGCAGUGUGGAUAACGAUCAUGAUUuAAGUGAUGAUGGUAAUGAUcUUGUUUUA   count   length
Libraries combined             Precision: [ total = 99.9% | 5p-arm = 87.5% | 3p-arm = 99.9% ]
---------------------------------------------------------------------------------------------------------------------------------------UAAGUGAUGAUGGUAAUGAUC--------   77764   21
---------------------------------------------------------------------------------------------------------------------------------------UAAGUGAUGAUGGUAAUGAU---------   31349   20
---------------------------------------------------------------------------------------------------------------------------------------UAAGUGAUGAUGGUAAUGAUCU-------   2904    22
---------------------------------------------------------------------------------------------------------------------------------------UAAGUGAUGAUGGUAAUGA----------   1326    19
---------------------------------------------------------------------------------------------------------------------------------------UAAGUGAUGAUGGUAAUG-----------   620     18
---------------------------------------------------------------------------------------------------------------------------------------UAAGUGAUGAUGGUAAUGAUCUU------   681     23
--------------------------------------------------------------------------------------------------------------------------------------UUAAGUGAUGAUGGUAAUGAUC--------   24      22
----------------------------------------------------------------------------------------------------------------------------------------AAGUGAUGAUGGUAAUGAUC--------   10      20
--------------------------------------------------------------------------------------------------------------------------------------UUAAGUGAUGAUGGUAAUGAU---------   36      21
----------------------------------------------------------------------------------------------------------------------------------------AAGUGAUGAUGGUAAUGAUCU-------   3       21
----------UCAUUCCCAUUGUCACCCAAA-------------------------------------------------------------------------------------------------------------------------------------   13      21
--------------------------------------------------------------------------------------------UAUAUAUGUUUACACCUCGGC---------------------------------------------------   1       21
------------------------------------------------------------------------------------------------------------UCGGCAGUGUGGAUAACGAUC-----------------------------------   8       21
------------------------------------------------------------------------------------------------------------------GUGUGGAUAACGAUCAUGAUU-----------------------------   1       21
-------------------------------------------------------------------------------------------------------------------UGUGGAUAACGAUCAUGAUUU----------------------------   1       21
------------------------------------------------------------------------------------------------------------------------------------AUUUAAGUGAUGAUGGUAAUGAU---------   1       23
-------------------------------------------------------------------------------------------------------------------------------------UUUAAGUGAUGAUGGUAAUG-----------   1       20
------------------------------------------------------------------------------------------------------------------------------------------GUGAUGAUGGUAAUGAUC--------   1       18
----------UCAUUCCCAUUGUCACCCAAAU------------------------------------------------------------------------------------------------------------------------------------   1       22
-------------UUCCCAUUGUCACCCAAAUCA----------------------------------------------------------------------------------------------------------------------------------   1       21
----------------------------------------------------UGUCUCUCAUCAGUGUGUCGA-------------------------------------------------------------------------------------------   1       21
----------------------------------------------------------------------------------------------UAUAUGUUUACACCUCGGCAG-------------------------------------------------   2       21
---------------------------------------------------------------------------------------------------------ACCUCGGCAGUGUGGAUAACGAUCAUGAU------------------------------   1       29
-----------------------------------------------------------------------------------------------------------CUCGGCAGUGUGGAUAACGAUCAUGAUU-----------------------------   1       28
------------------------------------------------------------------------------------------------------------UCGGCAGUGUGGAUAACGAUCAUGAU------------------------------   1       26
-------------------------------------------------------------------------------------------------------------CGGCAGUGUGGAUAACGAUCA----------------------------------   3       21
-------------------------------------------------------------------------------------------------------------------------------------UUUAAGUGAUGAUGGUAAUGAU---------   2       22
-------------------------------------------------------------------------------------------------------------------------------------UUUAAGUGAUGAUGGUAAUGAUCU-------   1       24
-------------------------------------------------------------------------------------------------------------------------------------UUUAAGUGAUGAUGGUAAUGAUC--------   5       23
-------------------------------------------------------------------------------------------------------------------------------------UUUAAGUGAUGAUGGUAAUGA----------   1       21
--------------------------------------------------------------------------------------------------------------------------------------UUAAGUGAUGAUGGUAAUGAUCU-------   3       23
--------------------------------------------------------------------------------------------------------------------------------------UUAAGUGAUGAUGGUAAUG-----------   1       19
---------------------------------------------------------------------------------------------------------------------------------------UAAGUGAUGAUGGUAAUGAUCUUGU----   1       25
----------------------------------------------------------------------------------------------------------------------------------------AAGUGAUGAUGGUAAUGAU---------   8       19
-----------------------------------------------------------------------------------------------------------------------------------------AGUGAUGAUGGUAAUGAU---------   1       18
-----------------------------------------------------------------------------------------------------------------------------------------AGUGAUGAUGGUAAUGAUC--------   1       19
```

```
AUACU U        C          CC       C   AU        UUA    UCUCAUC  U    C    CA        A
     C AGAuCAUU CCAUUGUCAC  AAaUCAU AUU  UUAUUUAU   UGUC       AG GUGU GACA  GUAUAUAU C
     G UcUAGUAA GGUAGUAGUG  uUUAGUA UAG  AAUAGGUG   ACGG       UC-CACA UUGU  UAUAUAUA A
AUUUU U        U          AA       C   C-        UG-    C------       U    A-        U
```

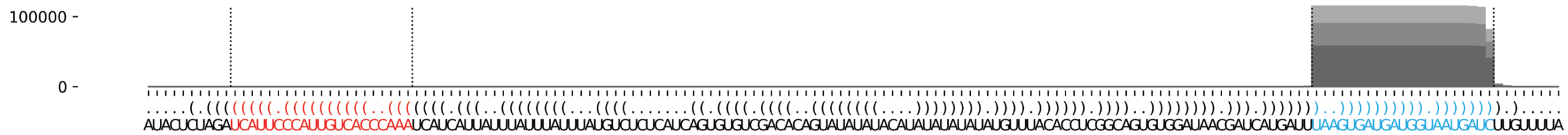

**Supplementary Data 1: Detailed depiction of identified miRNAs in dictyostelids.** Predicted miRNA hairpin sequences and structure displayed in bracket notion. Identified miRNA-5p is indicated in red, and the miRNA-3p in cyan. Lowercase nucleotides represent the 5’ and 3’ ends of the miRNA-5p and miRNA-3p. Below the sequence, all mapped small RNA reads are aligned to the miRNA hairpin and the number reads and their length are shown to the right. Results are shown for all small RNA sequencing libraries combined (denoted Libraries combined). For miRNAs that have fewer than 50,000 reads mapped, the results are shown for the individual libraries as well (denoted Library = 1 to 3). The percentage of reads that map to the exact 5’ nucleotide of either the miRNA-5p or miRNA-3p is shown as ‘precision’ and calculated on the entire miRNA-hairpin (denoted total), and on the 5’ and 3’ arms of the miRNA hairpin (denoted 5p-arm and 3p-arm respectively). The folded miRNA hairpin structure, shown below the mapped small RNA, was predicted using the ViennaRNA RNAlib-2.6.2 python package, with 22°C folding temperature. At the bottom, a graph shows the read mapping density on the miRNA hairpin, with the miRNA-5p and miRNA-3p indicated in red and cyan respectively. The different shades of grey in the graph represent the different small RNA sequencing libraries. ddi-mir-1186 was named ddi_mir_can_D1 when it was first identified (Meier *et al*., 2016), but we have here renamed it to ddi-mir-1186.
