## Supplementary Data 2 for "Evolution of microRNAs in Amoebozoa and implications for the origin of multicellularity"

### *P. polycephalum* ppo-mir-1221

```
((((((((((((((((((((((((.((((((((((((((((.((((....................(((......................)))....)))).)))))))))))))))).)))))))))))))))))))))))   -81.92 kcal/mol
CUUCUUUUGGaGGAUUUCAAAGUGUAUGACgAGUAUCACUACCUAAUUUUCAAACUUACAAACAAACCUAAUUACUUCUGCGCACAAUAUGGUACACUAGGAAGUGAUACUCGuCAUAAACUUUGAAAUCCUCcAAAAGGAG   count   length
Libraries combined         Precision: [ total = 99.5% | 5p-arm = 97.7% | 3p-arm = 99.6% ]
-----------------------------------------------------------------------------------------------------------------UCAUAAACUUUGAAAUCCUCC--------   2979   21
-----------------------------------------------------------------------------------------------------------------UCAUAAACUUUGAAAUCCU----------   1272   19
-----------------------------------------------------------------------------------------------------------------UCAUAAACUUUGAAAUCCUC---------   118    20
----------AGGAUUUCAAAGUGUAUGACG---------------------------------------------------------------------------------------------------------------   28     21
-------------------------------------------------------------------------------------------------------------------AUAAACUUUGAAAUCCUCC--------   9      19
----------AGGAUUUCAAAGUGUAUGAC----------------------------------------------------------------------------------------------------------------   14     20
---------------------------------------------------------------------------------------------ACACUAGGAAGUGAUACUCG-----------------------------   2      20
-----------------------------------------------------------------------------------------------------------------UCAUAAACUUUGAAAUCC-----------   10     18
--------------------------------------------------------------------------------------------------------------------UAAACUUUGAAAUCCUCC--------   4      18
------------------------------------------------------------------------------------------------------------------CAUAAACUUUGAAAUCCUCC--------   3      20
------------------------------------------------------------------------------------------------------------------CAUAAACUUUGAAAUCCU----------   1      18
-----UUUGGAGGAUUUCAAAGUG----------------------------------------------------------------------------------------------------------------------   1      19
-----------------------------------------------------------------------------------------------------------------UCAUAAACUUUGAAAUCCUCCA-------   1      22
Library = 1       Precision: [ total = 99.4% | 5p-arm = 100.0% | 3p-arm = 99.4% ]
-----------------------------------------------------------------------------------------------------------------UCAUAAACUUUGAAAUCCUCC--------   1107   21
-----------------------------------------------------------------------------------------------------------------UCAUAAACUUUGAAAUCCU----------   409    19
-----------------------------------------------------------------------------------------------------------------UCAUAAACUUUGAAAUCCUC---------   46     20
----------AGGAUUUCAAAGUGUAUGACG---------------------------------------------------------------------------------------------------------------   8      21
-------------------------------------------------------------------------------------------------------------------AUAAACUUUGAAAUCCUCC--------   3      19
----------AGGAUUUCAAAGUGUAUGAC----------------------------------------------------------------------------------------------------------------   2      20
---------------------------------------------------------------------------------------------ACACUAGGAAGUGAUACUCG-----------------------------   2      20
-----------------------------------------------------------------------------------------------------------------UCAUAAACUUUGAAAUCC-----------   2      18
--------------------------------------------------------------------------------------------------------------------UAAACUUUGAAAUCCUCC--------   2      18
------------------------------------------------------------------------------------------------------------------CAUAAACUUUGAAAUCCUCC--------   1      20
------------------------------------------------------------------------------------------------------------------CAUAAACUUUGAAAUCCU----------   1      18
Library = 2       Precision: [ total = 99.6% | 5p-arm = 100.0% | 3p-arm = 99.6% ]
-----------------------------------------------------------------------------------------------------------------UCAUAAACUUUGAAAUCCUCC--------   1579   21
-----------------------------------------------------------------------------------------------------------------UCAUAAACUUUGAAAUCCU----------   746    19
-----------------------------------------------------------------------------------------------------------------UCAUAAACUUUGAAAUCCUC---------   66     20
----------AGGAUUUCAAAGUGUAUGACG---------------------------------------------------------------------------------------------------------------   16     21
----------AGGAUUUCAAAGUGUAUGAC----------------------------------------------------------------------------------------------------------------   9      20
-----------------------------------------------------------------------------------------------------------------UCAUAAACUUUGAAAUCC-----------   8      18
-------------------------------------------------------------------------------------------------------------------AUAAACUUUGAAAUCCUCC--------   5      19
------------------------------------------------------------------------------------------------------------------CAUAAACUUUGAAAUCCUCC--------   2      20
--------------------------------------------------------------------------------------------------------------------UAAACUUUGAAAUCCUCC--------   2      18
-----------------------------------------------------------------------------------------------------------------UCAUAAACUUUGAAAUCCUCCA-------   1      22
Library = 3       Precision: [ total = 99.5% | 5p-arm = 87.5% | 3p-arm = 99.8% ]
-----------------------------------------------------------------------------------------------------------------UCAUAAACUUUGAAAUCCUCC--------   293    21
-----------------------------------------------------------------------------------------------------------------UCAUAAACUUUGAAAUCCU----------   117    19
-----------------------------------------------------------------------------------------------------------------UCAUAAACUUUGAAAUCCUC---------   6      20
----------AGGAUUUCAAAGUGUAUGACG---------------------------------------------------------------------------------------------------------------   4      21
----------AGGAUUUCAAAGUGUAUGAC----------------------------------------------------------------------------------------------------------------   3      20
-----UUUGGAGGAUUUCAAAGUG----------------------------------------------------------------------------------------------------------------------   1      19
-------------------------------------------------------------------------------------------------------------------AUAAACUUUGAAAUCCUCC--------   1      19
```

```
                       G                A    AUUUUCAAACUUACAAACAA   UAAUUACUUC
CUUCUUUUGGaGGAUUUCAAAGU UAUGACgAGUAUCACU CCUA                    ACC          U
GAGGAAAAcCUCCUAAAGUUUCA AUACuGCUCAUAGUGA GGAU    ----------------UGG          G
                       A                A    CACA                   UAUAACACGC
```

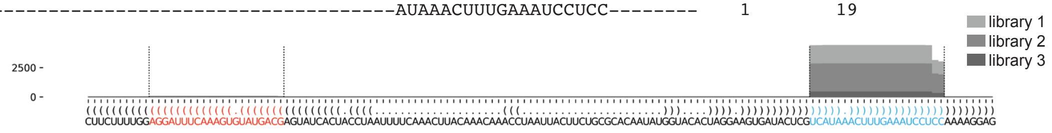

### *P. polycephalum* ppo-mir-1222

```
.((((((((((.(((((.(.(((.((((((((((((((((.((...((((((((((.((((((((((((((.(((((((((((..((.....(((((...((....))....))))))).....)).)))))))))))).))).))))).....))).)))))))))))))...))..))))))))))))))))).))).).))))).)))))))))))).   -128.63 kcal/mol
AGUUGUUCCCuCAGGUUGAGUUGUCCACAGCuGUUGCACUCCUGAAUCUGCACUUGACUACUUGUAGAAAAAUACUUUGCUGAUGGAUAGAUAGUGCCAAGGUUGGCCAUAAGCACUACAAAGCCCUAGCAAAGUAUCUUUCCUACAGCCCAAGUCAGCAAGUGUgGAAUAAGAAAGUGCAACAGCuGUGGCCAAGUGAAUCUAAGgGAACAAUU   count   length
```

Libraries combined Precision: [ total = 98.1% | 5p-arm = 98.5% | 3p-arm = 92.6% ]

| Read | count | length |
|---|---|---|
| UCAGGUUGAGUUGUCCACAGC | 241 | 21 |
| UCAGGUUGAGUUGUCCACA | 41 | 19 |
| UGUUGCACUCCUGAAUCUGC | 58 | 20 |
| UGUGGCCAAGUGAAUCUAAG | 22 | 20 |
| UCAGGUUGAGUUGUCCAC | 17 | 18 |
| UCAGGUUGAGUUGUCCACAG | 26 | 20 |
| UCAGGUUGAGUUGUCCACAGCU | 4 | 22 |
| UGUUGCACUCCUGAAUCUG | 3 | 19 |
| UGUGGAAUAAGAAAGUGCAACAG | 1 | 23 |
| UGUGGCCAAGUGAAUCUAA | 1 | 19 |
| UGUGGCCAAGUGAAUCUAAGG | 1 | 21 |
| CAGGUUGAGUUGUCCACAGC | 2 | 20 |
| UUGCACUCCUGAAUCUGCAC | 1 | 20 |
| ACUUGACUACUUGUAGAAAAAUACUUUGCUGA | 3 | 32 |
| GGAAUAAGAAAGUGCAACAGC | 1 | 21 |
| GAAUAAGAAAGUGCAACAGC | 1 | 20 |

Library = 1 Precision: [ total = 99.3% | 5p-arm = 100.0% | 3p-arm = 92.9% ]

| Read | count | length |
|---|---|---|
| UCAGGUUGAGUUGUCCACAGC | 81 | 21 |
| UCAGGUUGAGUUGUCCACA | 18 | 19 |
| UGUUGCACUCCUGAAUCUGC | 18 | 20 |
| UGUGGCCAAGUGAAUCUAAG | 11 | 20 |
| UCAGGUUGAGUUGUCCAC | 10 | 18 |
| UCAGGUUGAGUUGUCCACAG | 8 | 20 |
| UCAGGUUGAGUUGUCCACAGCU | 2 | 22 |
| UGUUGCACUCCUGAAUCUG | 1 | 19 |
| UGUGGAAUAAGAAAGUGCAACAG | 1 | 23 |
| UGUGGCCAAGUGAAUCUAA | 1 | 19 |
| UGUGGCCAAGUGAAUCUAAGG | 1 | 21 |

Library = 2 Precision: [ total = 97.9% | 5p-arm = 98.2% | 3p-arm = 92.3% ]

| Read | count | length |
|---|---|---|
| UCAGGUUGAGUUGUCCACAGC | 135 | 21 |
| UGUUGCACUCCUGAAUCUGC | 36 | 20 |
| UCAGGUUGAGUUGUCCACA | 20 | 19 |
| UCAGGUUGAGUUGUCCACAG | 17 | 20 |
| UGUGGCCAAGUGAAUCUAAG | 11 | 20 |
| UCAGGUUGAGUUGUCCAC | 5 | 18 |
| ACUUGACUACUUGUAGAAAAAUACUUUGCUGA | 3 | 32 |
| UCAGGUUGAGUUGUCCACAGCU | 2 | 22 |
| UGUUGCACUCCUGAAUCUG | 2 | 19 |
| UUGCACUCCUGAAUCUGCAC | 1 | 20 |
| GGAAUAAGAAAGUGCAACAGC | 1 | 21 |
| GAAUAAGAAAGUGCAACAGC | 1 | 20 |

Library = 3 Precision: [ total = 94.6% | 5p-arm = 94.6% | 3p-arm = 0% ]

| Read | count | length |
|---|---|---|
| UCAGGUUGAGUUGUCCACAGC | 25 | 21 |
| UGUUGCACUCCUGAAUCUGC | 4 | 20 |
| UCAGGUUGAGUUGUCCACA | 3 | 19 |
| UCAGGUUGAGUUGUCCAC | 2 | 18 |
| CAGGUUGAGUUGUCCACAGC | 2 | 20 |
| UCAGGUUGAGUUGUCCACAG | 1 | 20 |

```
A           C     G G   U                 C   GAA          A                          A            AU   AUAGA       CAA    U
 GUUGUUCCCu AGGUU A UUG CCACAGCuGUUGCACU -CU   UCUGcACUUG CU-ACU-----UGUAG-AAA AUACUUUGCUG   GG      UAGUGC    -GG U
 UAACAAGgGA UCUAA U AAC GGUGucGACAACGUGA  GA   AGgUGUGAAC-GA UGA      ACAUC UUU UAUGAAACGAU -CC      AUCACG     CC G
U           A     G G   C                 AA   AUA              C    ACCCG      C    C              C    GAAAC       AAUA   G
```

### *P. polycephalum* ppo-mir-1222 continued

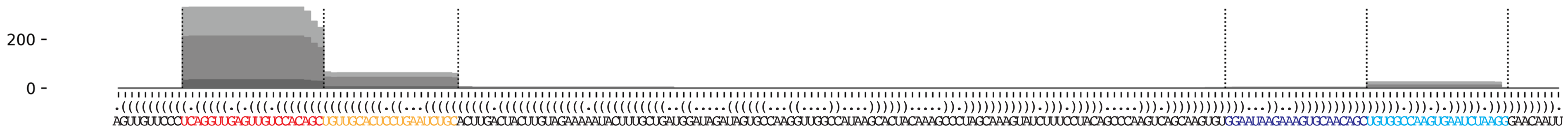


### *P. polycephalum* ppo-mir-1223

```
..((((((.(((((((.((..(((.((.((((((((.....)).)))))).)).)))..)).))))))).))))))..        -32.06 kcal/mol
UAAUAUUUUGuGAAAAAUGCUGGCUGUCAgUGAGAAUCUAUUUCCUCACUaAUUGCUUCCACUUUUCACcAAAUGUGA        count    length
Libraries combined          Precision: [ total = 100.0% | 5p-arm = 100.0% | 3p-arm = 100.0% ]
----------UGAAAAAUGCUGGCUGUCAG------------------------------------------------        100      20
----------UGAAAAAUGCUGGCUGUCA-------------------------------------------------        6        19
----------UGAAAAAUGCUGGCUGUCAGU-----------------------------------------------        6        21
--------------------------------------------------AAUUGCUUCCACUUUUCACC--------        4        20
----------UGAAAAAUGCUGGCUGUC--------------------------------------------------        5        18
Library = 1        Precision: [ total = 100.0% | 5p-arm = 100.0% | 3p-arm = 100.0% ]
----------UGAAAAAUGCUGGCUGUCAG------------------------------------------------        42       20
----------UGAAAAAUGCUGGCUGUCA-------------------------------------------------        2        19
----------UGAAAAAUGCUGGCUGUCAGU-----------------------------------------------        2        21
--------------------------------------------------AAUUGCUUCCACUUUUCACC--------        2        20
----------UGAAAAAUGCUGGCUGUC--------------------------------------------------        1        18
Library = 2        Precision: [ total = 100.0% | 5p-arm = 100.0% | 3p-arm = 100.0% ]
----------UGAAAAAUGCUGGCUGUCAG------------------------------------------------        50       20
----------UGAAAAAUGCUGGCUGUCA-------------------------------------------------        4        19
----------UGAAAAAUGCUGGCUGUCAGU-----------------------------------------------        4        21
----------UGAAAAAUGCUGGCUGUC--------------------------------------------------        3        18
--------------------------------------------------AAUUGCUUCCACUUUUCACC--------        2        20
Library = 3        Precision: [ total = 100.0% | 5p-arm = 100.0% | 3p-arm = 0% ]
----------UGAAAAAUGCUGGCUGUCAG------------------------------------------------        8        20
----------UGAAAAAUGCUGGCUGUC--------------------------------------------------        1        18

UA      U        A  CU   U  C         UC
  AUAUUU GuGAAAA UG  GGC GU AgUGAG-AA   \
  UGUAAA CACUUUU AC  UCG UA UCACUC UU   U
AG      c        C  CU   U  a      C  UA
```

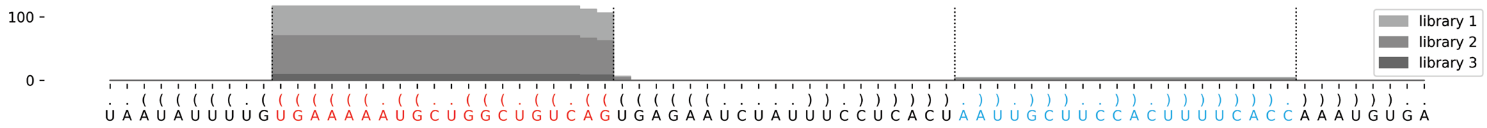

### *P. polycephalum* ppo-mir-1224

```
...((((..((((((((((((((((((((((....(((.((((((((..((((((...))).)))..)))))))).)))....)))))))))))))))))))))).)))).... -68.01 kcal/mol
UCAGGUUUAGuUUAGAACAUGAACUCGGACuCACAGACGCGGGCUUUCAGGUCAAAGUUUGCACUAAGGGGUCCGAGUCCACAAGuCUGAGUUUGUGUUCUAAACcAACCCAGC     count    length
Libraries combined          Precision: [ total = 98.5% | 5p-arm = 98.4% | 3p-arm = 100.0% ]
----------UUUAGAACAUGAACUCGGACU-----------------------------------------------------------------------------------     124      21
-------------------------------------------------------------------------------------UCUGAGUUUGUGUUCUAAACC--------     9        21
------------UAGAACAUGAACUCGGACU-----------------------------------------------------------------------------------     2        19
----------UUUAGAACAUGAACUCGGA-------------------------------------------------------------------------------------     1        19
Library = 1        Precision: [ total = 98.1% | 5p-arm = 98.0% | 3p-arm = 100.0% ]
----------UUUAGAACAUGAACUCGGACU-----------------------------------------------------------------------------------     48       21
-------------------------------------------------------------------------------------UCUGAGUUUGUGUUCUAAACC--------     4        21
------------UAGAACAUGAACUCGGACU-----------------------------------------------------------------------------------     1        19
Library = 2        Precision: [ total = 98.6% | 5p-arm = 98.5% | 3p-arm = 100.0% ]
----------UUUAGAACAUGAACUCGGACU-----------------------------------------------------------------------------------     64       21
-------------------------------------------------------------------------------------UCUGAGUUUGUGUUCUAAACC--------     4        21
------------UAGAACAUGAACUCGGACU-----------------------------------------------------------------------------------     1        19
Library = 3        Precision: [ total = 100.0% | 5p-arm = 100.0% | 3p-arm = 100.0% ]
----------UUUAGAACAUGAACUCGGACU-----------------------------------------------------------------------------------     12       21
----------UUUAGAACAUGAACUCGGA-------------------------------------------------------------------------------------     1        19
-------------------------------------------------------------------------------------UCUGAGUUUGUGUUCUAAACC--------     1        21

UCA    UA                     CACA   G          CA        A
   GGUU  GuUUAGAACAUGAACUCGGACu    GAC CGGGCUUU   GGU-CAA \
    CCAA CAAAUCUUGUGUUUGAGUCuGA    CUG GCCUGGGG   UCA GUU G
CGAC    c                     ACAC   A          AA    C   U
```

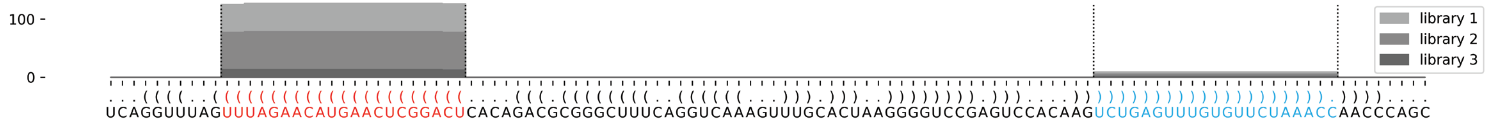

### *P. polycephalum* ppo-mir-1225

```
((((..(((((((.((((.(((((.(((((((((((((((((((((((.(((((((...((.(((((.(((.((.....)).))).)))))))...))))))).))))).)))))))))))))))))).))))).)))).)))))))..)))) -98.02 kcal/mol
UUCAUAaGGAUGGAUGGUAUCGAGCAcaAUUAUGCCAGUGAACCUCUgUGUGUAUGAGCUGUCUCUUUAAAAAUCCAUAAUAUUUUGGGAGCAAUACAUACACCCAgAGUGUUCACUGGUAUAAUUGuaCUUGACACCAACCAUCCUAcUGAA count    length
Libraries combined         Precision: [ total = 94.4% | 5p-arm = 94.8% | 3p-arm = 87.5% ]
---------------------------AAUUAUGCCAGUGAACCUCUG--------------------------------------------------------------------------------------------------------- 153      21
---------------------------AAUUAUGCCAGUGAACCUC----------------------------------------------------------------------------------------------------------- 29       19
---------------------------AAUUAUGCCAGUGAACCUCU---------------------------------------------------------------------------------------------------------- 31       20
------AGGAUGGAUGGUAUCGAGCAC------------------------------------------------------------------------------------------------------------------------------ 22       21
---------------------------AAUUAUGCCAGUGAACCUCUGU-------------------------------------------------------------------------------------------------------- 16       22
--------------------------------------------------------------------------------------------------------------------------------ACUUGACACCAACCAUCCUAC---- 12       21
------AGGAUGGAUGGUAUCGAGC-------------------------------------------------------------------------------------------------------------------------------- 4        19
----------------------------------------------------------------------------------------------------------GAGUGUUCACUGGUAUAAUUGU------------------------- 2        22
-UCAUAAGGAUGGAUGGUAUCGAGCAC------------------------------------------------------------------------------------------------------------------------------ 1        26
----------------------------AUUAUGCCAGUGAACCUCUG--------------------------------------------------------------------------------------------------------- 3        20
---------------------------------------------------------------------------------------------------------AGAGUGUUCACUGGUAUAAUUGU------------------------- 1        23
UUCAUAAGGAUGGAUGGUAUCGAGC-------------------------------------------------------------------------------------------------------------------------------- 1        25
----UAAGGAUGGAUGGUAUCGAGCAC------------------------------------------------------------------------------------------------------------------------------ 1        23
----UAAGGAUGGAUGGUAUCGAGC-------------------------------------------------------------------------------------------------------------------------------- 1        21
---------------------------AAUUAUGCCAGUGAACCU------------------------------------------------------------------------------------------------------------ 1        18
---------------------------AAUUAUGCCAGUGAACCUCUGUGUGUA--------------------------------------------------------------------------------------------------- 1        27
-----------------------------UUAUGCCAGUGAACCUCUGUGUGUA--------------------------------------------------------------------------------------------------- 3        25
------------------------------UAUGCCAGUGAACCUCUGU-------------------------------------------------------------------------------------------------------- 1        19
------------------------------------------------UGUGUAUGAGCUGUCUCUUU------------------------------------------------------------------------------------- 2        20
----------------------------------------------------------------------------------------------------------------------------------UUGACACCAACCAUCCUACUG-- 1        21
Library = 1        Precision: [ total = 97.4% | 5p-arm = 98.2% | 3p-arm = 85.7% ]
---------------------------AAUUAUGCCAGUGAACCUCUG--------------------------------------------------------------------------------------------------------- 65       21
---------------------------AAUUAUGCCAGUGAACCUC----------------------------------------------------------------------------------------------------------- 15       19
---------------------------AAUUAUGCCAGUGAACCUCU---------------------------------------------------------------------------------------------------------- 12       20
------AGGAUGGAUGGUAUCGAGCAC------------------------------------------------------------------------------------------------------------------------------ 7        21
---------------------------AAUUAUGCCAGUGAACCUCUGU-------------------------------------------------------------------------------------------------------- 6        22
--------------------------------------------------------------------------------------------------------------------------------ACUUGACACCAACCAUCCUAC---- 4        21
------AGGAUGGAUGGUAUCGAGC-------------------------------------------------------------------------------------------------------------------------------- 3        19
----------------------------------------------------------------------------------------------------------GAGUGUUCACUGGUAUAAUUGU------------------------- 2        22
-UCAUAAGGAUGGAUGGUAUCGAGCAC------------------------------------------------------------------------------------------------------------------------------ 1        26
----------------------------AUUAUGCCAGUGAACCUCUG--------------------------------------------------------------------------------------------------------- 1        20
---------------------------------------------------------------------------------------------------------AGAGUGUUCACUGGUAUAAUUGU------------------------- 1        23
Library = 2        Precision: [ total = 91.6% | 5p-arm = 91.7% | 3p-arm = 88.9% ]
---------------------------AAUUAUGCCAGUGAACCUCUG--------------------------------------------------------------------------------------------------------- 78       21
---------------------------AAUUAUGCCAGUGAACCUCU---------------------------------------------------------------------------------------------------------- 18       20
------AGGAUGGAUGGUAUCGAGCAC------------------------------------------------------------------------------------------------------------------------------ 14       21
---------------------------AAUUAUGCCAGUGAACCUC----------------------------------------------------------------------------------------------------------- 12       19
---------------------------AAUUAUGCCAGUGAACCUCUGU-------------------------------------------------------------------------------------------------------- 10       22
--------------------------------------------------------------------------------------------------------------------------------ACUUGACACCAACCAUCCUAC---- 8        21
-----------------------------UUAUGCCAGUGAACCUCUGUGUGUA--------------------------------------------------------------------------------------------------- 3        25
----------------------------AUUAUGCCAGUGAACCUCUG--------------------------------------------------------------------------------------------------------- 2        20
------------------------------------------------UGUGUAUGAGCUGUCUCUUU------------------------------------------------------------------------------------- 2        20
UUCAUAAGGAUGGAUGGUAUCGAGC-------------------------------------------------------------------------------------------------------------------------------- 1        25
----UAAGGAUGGAUGGUAUCGAGCAC------------------------------------------------------------------------------------------------------------------------------ 1        23
----UAAGGAUGGAUGGUAUCGAGC-------------------------------------------------------------------------------------------------------------------------------- 1        21
---------------------------AAUUAUGCCAGUGAACCU------------------------------------------------------------------------------------------------------------ 1        18
---------------------------AAUUAUGCCAGUGAACCUCUGUGUGUA--------------------------------------------------------------------------------------------------- 1        27
------------------------------UAUGCCAGUGAACCUCUGU-------------------------------------------------------------------------------------------------------- 1        19
----------------------------------------------------------------------------------------------------------------------------------UUGACACCAACCAUCCUACUG-- 1        21
```

### *P. polycephalum* ppo-mir-1225 continued

```
((((..(((((((.((((.(((((.(((((((((((((((((((((((.(((((((...((.(((((.(((.((.....)).))).)))))))...))))))).))))).)))))))))))))))))).))))).)))).)))))))..)))) -98.02 kcal/mol
UUCAUAaGGAUGGAUGGUAUCGAGCAcaAUUAUGCCAGUGAACCUCUgUGUGUAUGAGCUGUCUCUUUAAAAAUCCAUAAUAUUUUGGGAGCAAUACAUACACCCAgAGUGUUCACUGGUAUAAUUGuaCUUGACACCAACCAUCCUAcUGAA count    length
Library = 3       Precision: [ total = 100.0% | 5p-arm = 100.0% | 3p-arm = 0% ]
---------------------------AAUUAUGCCAGUGAACCUCUG--------------------------------------------------------------------------------------------------------- 10       21
---------------------------AAUUAUGCCAGUGAACCUC----------------------------------------------------------------------------------------------------------- 2        19
------AGGAUGGAUGGUAUCGAGCAC------------------------------------------------------------------------------------------------------------------------------ 1        21
------AGGAUGGAUGGUAUCGAGC-------------------------------------------------------------------------------------------------------------------------------- 1        19
---------------------------AAUUAUGCCAGUGAACCUCU---------------------------------------------------------------------------------------------------------- 1        20

    UA        A    A     C                        U       AGC  U     U   A  CC
UUCA  aGGAUGG UGGU UCGAG AcaAUUAUGCCAGUGAAC-CUCUg GUGUAUG   UG CUCUU AAA AU  \
AAGU  UCCUACC ACCA AGUUC uGUUAAUAUGGUCACUUG GAgAC CACAUAC   AC-GAGGG UUU UA  A
    cA        A    C     a                  U     C       AUA        U   A  AU
```

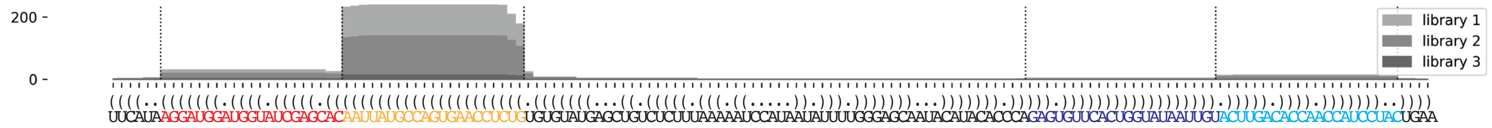

### *P. polycephalum* ppo-mir-1226

`..((((((((((.(((((((((((((((((.(((.(((.(.(((((((.((((((..(((((((((((((((((((((((((((.(.(..((((.((((((((((..((((.(((.((((((((((.(((((((....)))))))..)))..))))))).))).))))..))))..................))))).)))).).).)))))))))))))))))))..))).)))))).))))))))))))))).).))).))))))))))))))))))))).)))))))..)))))` –184.64 kcal/mol

`CCCCCUCUCAaAGAGUACCUUACUUCAGACcCGAUUGCGCGGGUUACUGAGGUAUGUCUCGUGCGGUACAUGCGAGCGCGUGCGCGUGCGUUGCGCGUGUCGUGUUAUGAUGAACUUCAUGUAUCGUAUGUCGCGUACUCGUGACACAACGCAAUGCGUGCAGUGCGUUUCAACAAUAUACCUCACUUCAGCUGACAGUGCGUCUCUUGCGUGUGUGCUCGUGUGCUUGCGCGCGCGGUAUGUACCCAGUGGCACACGCUAUCGGuCUGAAGUAAGGUACCCUUUgAAAGAGGG` count length

Libraries combined Precision: [ total = 98.0% | 5p-arm = 47.6% | 3p-arm = 99.2% ]

| read | count | length |
|---|---|---|
| UCUGAAGUAAGGUACCCUUUG | 2153 | 21 |
| UCUGAAGUAAGGUACCCUUU | 557 | 20 |
| UCUGAAGUAAGGUACCCU | 420 | 18 |
| UCUGAAGUAAGGUACCCUU | 399 | 19 |
| UCUGAAGUAAGGUACCCUUUGA | 61 | 22 |
| AAGAGUACCUUACUUCAGACC | 26 | 21 |
| UUGCGCGGGUUACUGAGGU | 14 | 19 |
| AAGAGUACCUUACUUCAGACCC | 10 | 22 |
| UGAAGUAAGGUACCCUUUG | 7 | 19 |
| UUGCGCGGGUUACUGAGGUA | 13 | 20 |
| UUGCGCGGGUUACUGAGG | 4 | 18 |
| AUGUACCCAGUGGCACACG | 2 | 19 |
| CCCAGUGGCACACGCUAUCGG | 6 | 21 |
| CAGUGGCACACGCUAUCGGUC | 3 | 21 |
| AAGAGUACCUUACUUCAGAC | 2 | 20 |
| UCGUGCGGUACAUGCGAGC | 1 | 19 |
| UUGCGCGCGCGUAUGUACC | 1 | 19 |
| CCAGUGGCACACGCUAUCGGU | 2 | 21 |
| AGUGGCACACGCUAUCGGUC | 3 | 20 |
| GUGGCACACGCUAUCGGUC | 3 | 19 |
| UGAAGUAAGGUACCCUUU | 1 | 18 |
| AAGAGUACCUUACUUCAGA | 2 | 19 |
| CCCGAUUGCGCGGGUUACUGAGG | 1 | 23 |
| CGAUUGCGCGGGUUACUGA | 1 | 19 |
| GAUUGCGCGGGUUACUGAGGUA | 2 | 22 |
| UUGCGCGGGUUACUGAGGUAU | 1 | 21 |
| UAUGUCUCGUGCGGUACAUG | 1 | 20 |
| UAUGUCUCGUGCGGUACAUGCGA | 1 | 23 |
| AUGUCUCGUGCGGUACAUGCGAG | 1 | 23 |
| UGUCUCGUGCGGUACAUG | 2 | 18 |
| UGUCUCGUGCGGUACAUGCGAGCGC | 1 | 25 |
| UGUCUCGUGCGGUACAUGCG | 1 | 20 |
| UGCUUGCGCGCGCGGUAUGUA | 1 | 20 |
| CACACGCUAUCGGUCUGAAG | 1 | 20 |
| UCUGAAGUAAGGUACCCUUUGAA | 3 | 23 |

Library = 1 Precision: [ total = 97.8% | 5p-arm = 53.1% | 3p-arm = 98.9% ]

| read | count | length |
|---|---|---|
| UCUGAAGUAAGGUACCCUUUG | 763 | 21 |
| UCUGAAGUAAGGUACCCUUU | 218 | 20 |
| UCUGAAGUAAGGUACCCU | 201 | 18 |
| UCUGAAGUAAGGUACCCUU | 172 | 19 |
| UCUGAAGUAAGGUACCCUUUGA | 23 | 22 |
| AAGAGUACCUUACUUCAGACC | 10 | 21 |
| UUGCGCGGGUUACUGAGGU | 8 | 19 |
| AAGAGUACCUUACUUCAGACCC | 6 | 22 |
| UGAAGUAAGGUACCCUUUG | 5 | 19 |
| UUGCGCGGGUUACUGAGGUA | 3 | 20 |
| UUGCGCGGGUUACUGAGG | 3 | 18 |
| AUGUACCCAGUGGCACACG | 2 | 19 |
| CCCAGUGGCACACGCUAUCGG | 2 | 21 |
| CAGUGGCACACGCUAUCGGUC | 2 | 21 |
| AAGAGUACCUUACUUCAGAC | 1 | 20 |
| UCGUGCGGUACAUGCGAGC | 1 | 19 |
| UUGCGCGCGCGUAUGUACC | 1 | 19 |
| CCAGUGGCACACGCUAUCGGU | 1 | 21 |
| AGUGGCACACGCUAUCGGUC | 1 | 20 |
| GUGGCACACGCUAUCGGUC | 1 | 19 |
| UGAAGUAAGGUACCCUUU | 1 | 18 |

Library = 2 Precision: [ total = 98.2% | 5p-arm = 38.1% | 3p-arm = 99.5% ]

| read | count | length |
|---|---|---|
| UCUGAAGUAAGGUACCCUUUG | 1192 | 21 |
| UCUGAAGUAAGGUACCCUUU | 299 | 20 |
| UCUGAAGUAAGGUACCCU | 193 | 18 |
| UCUGAAGUAAGGUACCCUU | 186 | 19 |
| UCUGAAGUAAGGUACCCUUUGA | 32 | 22 |

### *P. polycephalum* ppo-mir-1226 continued

```
..((((((((((.((((((((((((((((.(((.(((.(.(((((((.((((((..((((((((((((((((((((((((((.(.(..((((.((((((((..((((.(((.(((((((((.((((((....)))))))..)))..)))))))).))).))))..)))).................))))).)))).).).))))))))))))))))))..))).)))))).))))))))))))).).))).))))))))))))))))))).))))))..)))))   −184.64 kcal/mol
CCCCCUCUCAaAGAGUACCUUACUUCAGACcCGAUUGCGCGGGUUACUGAGGUAUGUCUCGUGCGGUACAUGCCGAGCGCGUGCGCGUGCGUUUGCGGUGUCGUGUUAUGAUGAACUUCAUGUAUCGUAUGUCGGUACUCGUGACACAACGCAAUGCGGUGCAGUGCGUUUCAACAAUAUACCUCACUUCAGCUGACAGUGCGUCUUGCGUGUGUGCUCGUGUGCUUGCGGCGCGGUAUGUACCCAGUGGCACACGGUAUCGGuCUGAAGUAAGGUACCCUUUgAAAGAGGG   count   length
```

Libraries combined — Precision: [ total = 98.0% | 5p-arm = 47.6% | 3p-arm = 99.2% ]

| read | count | length |
| --- | --- | --- |
| AAGAGUACCUUACUUCAGACC | 11 | 21 |
| UUGCGCGGGUUACUGAGGUA | 8 | 20 |
| UUGCGCGGGUUACUGAGGU | 5 | 19 |
| AAGAGUACCUUACUUCAGACCC | 3 | 22 |
| UCUGAAGUAAGGUACCCUUUGAA | 3 | 23 |
| GAUUGCGCGGGUUACUGAGGUA | 2 | 22 |
| UGUCUCGUGCGGUACAUG | 2 | 18 |
| AGUGGCACACGGUAUCGGUC | 2 | 20 |
| UGAAGUAAGGUACCCUUUG | 2 | 19 |
| AAGAGUACCUUACUUCAGAC | 1 | 20 |
| AAGAGUACCUUACUUCAGA | 1 | 19 |
| CCCGAUUGCGCGGGUUACUGAGG | 1 | 23 |
| CGAUUGCGCGGGUUACUGA | 1 | 19 |
| UUGCGCGGGUUACUGAGG | 1 | 18 |
| UUGCGCGGGUUACUGAGGUAU | 1 | 21 |
| UAUGUCUCGUGCGGUACAUG | 1 | 20 |
| UAUGUCUCGUGCGGUACAUGCCGA | 1 | 23 |
| AUGUCUCGUGCGGUACAUGCCGAG | 1 | 23 |
| UGUCUCGUGCGGUACAUGCCGAGCGC | 1 | 25 |
| UGUCUCGUGCGGUACAUGCG | 1 | 20 |
| UGCUUGCGGCGCGGUAUGUA | 1 | 20 |
| CCCAGUGGCACACGGUAUCGG | 1 | 21 |
| CCAGUGGCACACGGUAUCGGU | 1 | 21 |
| CAGUGGCACACGGUAUCGGUC | 1 | 21 |
| GUGGCACACGGUAUCGGUC | 1 | 19 |
| CACACGGUAUCGGUCUGAAG | 1 | 20 |

Library = 3 — Precision: [ total = 97.8% | 5p-arm = 70.0% | 3p-arm = 98.7% ]

| read | count | length |
| --- | --- | --- |
| UCUGAAGUAAGGUACCCUUUG | 198 | 21 |
| UCUGAAGUAAGGUACCCUU | 41 | 19 |
| UCUGAAGUAAGGUACCCUUU | 40 | 20 |
| UCUGAAGUAAGGUACCCU | 26 | 18 |
| UCUGAAGUAAGGUACCCUUUGA | 6 | 22 |
| AAGAGUACCUUACUUCAGACC | 5 | 21 |
| CCCAGUGGCACACGGUAUCGG | 3 | 21 |
| UUGCGCGGGUUACUGAGGUA | 2 | 20 |
| AAGAGUACCUUACUUCAGACCC | 1 | 22 |
| AAGAGUACCUUACUUCAGA | 1 | 19 |
| UUGCGCGGGUUACUGAGGU | 1 | 19 |
| GUGGCACACGGUAUCGGUC | 1 | 19 |

```
CC              A                      C    U    C G        A        CU                                 U C UU     G                       AU    A    U              A          U
  CCCUC--UCAaAG GUACCUUACUUCAGACc GAU GCG G GUUACUG GGUAUGU  CGUGCG-GUA--CAUGCGAGCGCGUGCGCG G G  UGCG UGUCG-----------------UGUU  GAUG ACU CAUGUAU--CGU -UGUCGCG A
  GGGAG  AgUUUC CAUGGAAUGAAGUCuGG-CUA CGC C CGGUGAC-CCAUGUA -GCGCGC CGU  GUGUGCUCGUGUGUGCGU C C -GCGU ACAGU                 ACAA  UUGC UGA GUGCGUA  GCA  ACAGUGC C
       AA       C                   U   A A                U        G   UC                  U U U      G      CGACUUCACUCCAUAUA  CU    G    C         AC   AC         U
```

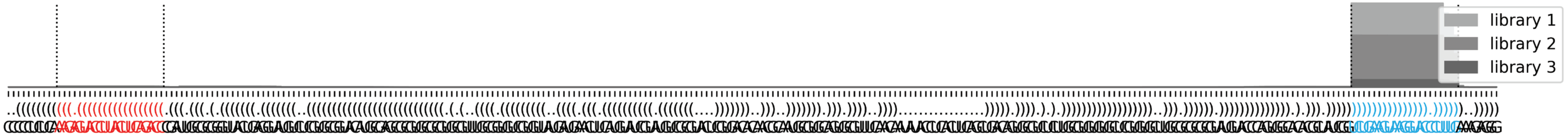

### *P. polycephalum* ppo-mir-1227

```
..((((((.((((((.(((((((.(((((((((.......(((((.(((((((.(..((((.(((.((.(((........))).)))))...))))..).)).))))).)))))...))))))))).))))))).)))))).))..))))   -64.91 kcal/mol
AGACAGUGAGuGAGUAUUUGUGAGGUGCACaGACCAAACAACAAAAACAAAAAAAGGGCGCCAACGUGGUUUUGAAAAUUGAAGCAGUUCCCGCGCAAUAUUCUUUGUGUUUGUGCUUCUGuGCACAUCAUGGACACUCACCcCAGAUUGU   count   length
Libraries combined         Precision: [ total = 97.3% | 5p-arm = 100.0% | 3p-arm = 97.2% ]
-------------------------------------------------------------------------------------------------------------------------UGCACAUCAUGGACACUCACC--------   374   21
----------UGAGUAUUUGUGAGGUGC--------------------------------------------------------------------------------------------------------------------------   13    18
--------------------------------------------------------------------------------------------------------------------------GCACAUCAUGGACACUCACC--------   9     20
----------UGAGUAUUUGUGAGGUGCACA-----------------------------------------------------------------------------------------------------------------------   3     21
----------UGAGUAUUUGUGAGGUGCA-------------------------------------------------------------------------------------------------------------------------   2     19
----------UGAGUAUUUGUGAGGUGCAC------------------------------------------------------------------------------------------------------------------------   3     20
----------UGAGUAUUUGUGAGGUGCACAG----------------------------------------------------------------------------------------------------------------------   1     22
-------------------------------------------------------------------------------------------------------------------------UGCACAUCAUGGACACUCAC---------   2     20
-----------------------------------------------------------------------------------------------------------------------UGUGCACAUCAUGGACACU------------   1     19
-----------------------------------------------------------------------------------------------------------------------UGUGCACAUCAUGGACACUC-----------   1     20
Library = 1        Precision: [ total = 98.1% | 5p-arm = 100.0% | 3p-arm = 97.9% ]
-------------------------------------------------------------------------------------------------------------------------UGCACAUCAUGGACACUCACC--------   188   21
----------UGAGUAUUUGUGAGGUGC--------------------------------------------------------------------------------------------------------------------------   9     18
--------------------------------------------------------------------------------------------------------------------------GCACAUCAUGGACACUCACC--------   4     20
----------UGAGUAUUUGUGAGGUGCACA-----------------------------------------------------------------------------------------------------------------------   2     21
----------UGAGUAUUUGUGAGGUGCA-------------------------------------------------------------------------------------------------------------------------   1     19
----------UGAGUAUUUGUGAGGUGCAC------------------------------------------------------------------------------------------------------------------------   1     20
----------UGAGUAUUUGUGAGGUGCACAG----------------------------------------------------------------------------------------------------------------------   1     22
-------------------------------------------------------------------------------------------------------------------------UGCACAUCAUGGACACUCAC---------   1     20
Library = 2        Precision: [ total = 96.7% | 5p-arm = 100.0% | 3p-arm = 96.5% ]
-------------------------------------------------------------------------------------------------------------------------UGCACAUCAUGGACACUCACC--------   165   21
----------UGAGUAUUUGUGAGGUGC--------------------------------------------------------------------------------------------------------------------------   4     18
--------------------------------------------------------------------------------------------------------------------------GCACAUCAUGGACACUCACC--------   4     20
----------UGAGUAUUUGUGAGGUGCAC------------------------------------------------------------------------------------------------------------------------   2     20
----------UGAGUAUUUGUGAGGUGCACA-----------------------------------------------------------------------------------------------------------------------   1     21
----------UGAGUAUUUGUGAGGUGCA-------------------------------------------------------------------------------------------------------------------------   1     19
-----------------------------------------------------------------------------------------------------------------------UGUGCACAUCAUGGACACU------------   1     19
-----------------------------------------------------------------------------------------------------------------------UGUGCACAUCAUGGACACUC-----------   1     20
-------------------------------------------------------------------------------------------------------------------------UGCACAUCAUGGACACUCAC---------   1     20
Library = 3        Precision: [ total = 95.5% | 5p-arm = 0% | 3p-arm = 95.5% ]
-------------------------------------------------------------------------------------------------------------------------UGCACAUCAUGGACACUCACC--------   21    21
--------------------------------------------------------------------------------------------------------------------------GCACAUCAUGGACACUCACC--------   1     20
```

```
AG        A      A       G         CCAAACA     A        A GG    C     G  G   UGA
  ACAG--UG GuGAGU UUUGUGA GUGCACaGA       ACAAA ACAAA-AA A  GCGC --AAC UG UUU   A
  UGUU  AC CACUCA AGGUACU CACGuGUCU   ----UGUUU UGUUU UU U  CGCG   UUG-AC AAG   A
      AG  c      C       A         UCG         G     C  A AA    CCC      G   UUA
```

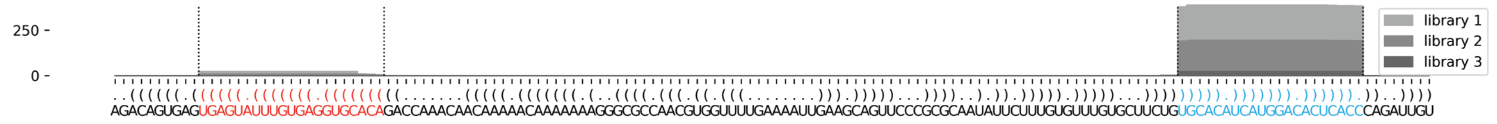

### *A. castellanii* aca-mir-1228

```
.....((((((((((((.(((((.((((((((.((...((....)).)).)))))))).))))).)))))))))))))).....    -56.3 kcal/mol
CCAUUUCAUGUUUAGUGCCUAUGUCCCCUUCUGUCAUACCUAUUGGGGAGAGAAGGGGUCAUAGACACUAAACAUGACUCUG    count    length
Libraries combined          Precision: [ total = 94.8% | 5p-arm = 94.8% | 3p-arm = 94.9% ]
----------UUUAGUGCCUAUGUCCCCUUC---------------------------------------------------    87       21
-----------------------------------------------------AGGGGUCAUAGACACUAAACA--------    40       21
-----------------------------------------------------AGGGGUCAUAGACACUAAAC---------    16       20
---------GUUUAGUGCCUAUGUCCCCUUCU--------------------------------------------------    1        23
---------GUUUAGUGCCUAUGUCCCCUUC---------------------------------------------------    1        22
----------UUUAGUGCCUAUGUCCCCUU----------------------------------------------------    2        20
----------UUUAGUGCCUAUGUCCCCUUCUGUCAUACCUA----------------------------------------    1        32
----------UUUAGUGCCUAUGUCCCCUUCUGUCAUACC------------------------------------------    1        30
----------UUUAGUGCCUAUGUCCCCUUCUG-------------------------------------------------    2        23
-----------UUAGUGCCUAUGUCCCCUUC---------------------------------------------------    1        20
-------------------------------------ACCUAUUGGGGAGAGAAGGGGUCAUAGACAC--------------    1        31
-----------------------------------------AUUGGGGAGAGAAGGGGUCAUAGACACUAAACA--------    1        33
------------------------------------------UUGGGGAGAGAAGGGGUCAUAGACACUAAACA--------    1        32
Library = 1        Precision: [ total = 93.0% | 5p-arm = 92.3% | 3p-arm = 93.9% ]
----------UUUAGUGCCUAUGUCCCCUUC---------------------------------------------------    58       21
-----------------------------------------------------AGGGGUCAUAGACACUAAACA--------    32       21
-----------------------------------------------------AGGGGUCAUAGACACUAAAC---------    14       20
---------GUUUAGUGCCUAUGUCCCCUUCU--------------------------------------------------    1        23
---------GUUUAGUGCCUAUGUCCCCUUC---------------------------------------------------    1        22
----------UUUAGUGCCUAUGUCCCCUU----------------------------------------------------    2        20
----------UUUAGUGCCUAUGUCCCCUUCUGUCAUACCUA----------------------------------------    1        32
----------UUUAGUGCCUAUGUCCCCUUCUGUCAUACC------------------------------------------    1        30
-----------UUAGUGCCUAUGUCCCCUUC---------------------------------------------------    1        20
-------------------------------------ACCUAUUGGGGAGAGAAGGGGUCAUAGACAC--------------    1        31
-----------------------------------------AUUGGGGAGAGAAGGGGUCAUAGACACUAAACA--------    1        33
------------------------------------------UUGGGGAGAGAAGGGGUCAUAGACACUAAACA--------    1        32
Library = 2        Precision: [ total = 100.0% | 5p-arm = 100.0% | 3p-arm = 100.0% ]
----------UUUAGUGCCUAUGUCCCCUUC---------------------------------------------------    26       21
-----------------------------------------------------AGGGGUCAUAGACACUAAACA--------    5        21
-----------------------------------------------------AGGGGUCAUAGACACUAAAC---------    1        20
----------UUUAGUGCCUAUGUCCCCUUCUG-------------------------------------------------    2        23
Library = 3        Precision: [ total = 100.0% | 5p-arm = 100.0% | 3p-arm = 100.0% ]
----------UUUAGUGCCUAUGUCCCCUUC---------------------------------------------------    3        21
-----------------------------------------------------AGGGGUCAUAGACACUAAACA--------    3        21
-----------------------------------------------------AGGGGUCAUAGACACUAAAC---------    1        20
```

```
CCAUU            C     U         G   AUA   U
     UCAUGUUUAGUG CUAUG CCCCUUCU UC    CC  A
     AGUACAAAUCAC GAUAC GGGGAAGA AG  --GG  U
GUCUC            A     U         G   G     U
```

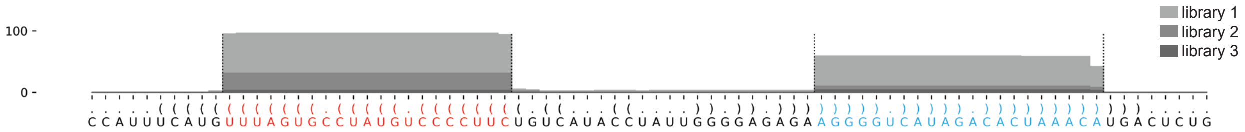

### *A. lenticulata* ale-mir-1228-P1

```
.....((((((((((((.(((((.((((((((((..((.((((........)))))).)))))))))).))))).))))))))))))).....   -58.42 kcal/mol
AUUCUUCAUGUUUAGUGCCUAUGUCCUCUUCACUCAUGAUCGCCAUCACGAGCGGCACAGUGAAGAGGUCAUAGACACUAAACAUGACUUUU   count    length
Libraries combined          Precision: [ total = 99.4% | 5p-arm = 99.3% | 3p-arm = 100.0% ]
----------UUUAGUGCCUAUGUCCUCUUCA------------------------------------------------------------   101      22
----------UUUAGUGCCUAUGUCCUCUUC-------------------------------------------------------------   116      21
----------UUUAGUGCCUAUGUCCUC----------------------------------------------------------------   91       18
----------UUUAGUGCCUAUGUCCUCUU--------------------------------------------------------------   94       20
----------UUUAGUGCCUAUGUCCUCU---------------------------------------------------------------   48       19
--------------------------------------------------------------AAGAGGUCAUAGACACUAAACA--------   9        22
-----------UUAGUGCCUAUGUCCUCUUCA------------------------------------------------------------   2        21
-----------UUAGUGCCUAUGUCCUCUUC-------------------------------------------------------------   1        20
--------------------------------------------------------------AAGAGGUCAUAGACACUAAAC---------   4        21
Library = 1        Precision: [ total = 100.0% | 5p-arm = 100.0% | 3p-arm = 100.0% ]
----------UUUAGUGCCUAUGUCCUCUUCA------------------------------------------------------------   39       22
----------UUUAGUGCCUAUGUCCUCUUC-------------------------------------------------------------   31       21
----------UUUAGUGCCUAUGUCCUC----------------------------------------------------------------   25       18
----------UUUAGUGCCUAUGUCCUCUU--------------------------------------------------------------   22       20
----------UUUAGUGCCUAUGUCCUCU---------------------------------------------------------------   10       19
--------------------------------------------------------------AAGAGGUCAUAGACACUAAACA--------   3        22
Library = 2        Precision: [ total = 98.6% | 5p-arm = 98.6% | 3p-arm = 100.0% ]
----------UUUAGUGCCUAUGUCCUCUUC-------------------------------------------------------------   59       21
----------UUUAGUGCCUAUGUCCUCUU--------------------------------------------------------------   45       20
----------UUUAGUGCCUAUGUCCUC----------------------------------------------------------------   40       18
----------UUUAGUGCCUAUGUCCUCUUCA------------------------------------------------------------   37       22
----------UUUAGUGCCUAUGUCCUCU---------------------------------------------------------------   28       19
-----------UUAGUGCCUAUGUCCUCUUCA------------------------------------------------------------   2        21
--------------------------------------------------------------AAGAGGUCAUAGACACUAAACA--------   2        22
--------------------------------------------------------------AAGAGGUCAUAGACACUAAAC---------   2        21
-----------UUAGUGCCUAUGUCCUCUUC-------------------------------------------------------------   1        20
Library = 3        Precision: [ total = 100.0% | 5p-arm = 100.0% | 3p-arm = 100.0% ]
----------UUUAGUGCCUAUGUCCUCUU--------------------------------------------------------------   27       20
----------UUUAGUGCCUAUGUCCUC----------------------------------------------------------------   26       18
----------UUUAGUGCCUAUGUCCUCUUC-------------------------------------------------------------   26       21
----------UUUAGUGCCUAUGUCCUCUUCA------------------------------------------------------------   25       22
----------UUUAGUGCCUAUGUCCUCU---------------------------------------------------------------   10       19
--------------------------------------------------------------AAGAGGUCAUAGACACUAAACA--------   4        22
--------------------------------------------------------------AAGAGGUCAUAGACACUAAAC---------   2        21
AUUCU            C     U          CA  A    CAU
     UCAUGUUUAGUG CUAUG CCUCUUCACU  UG UCGC   C
     AGUACAAAUCAC GAUAC GGAGAAGUGA -AC-GGCG   A
UUUUC            A     U          C        AGC
```

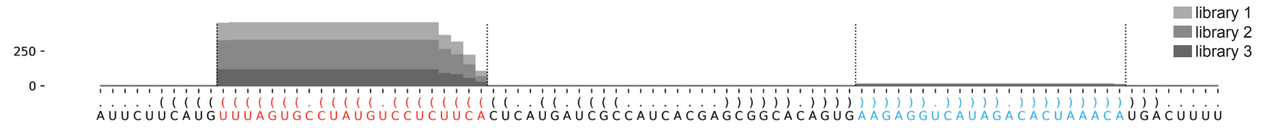

### *A. lenticulata* ale-mir-1229

```
..(.((.(((((((((((((((((.((((((((............(((...(.((.(((((((((((((.((.......)).))))))))).)))).)).)...)))...)))))))).))))))))))))))))).)).)..   -96.8 kcal/mol
ACGAGGAGAGaGCGGGCUGGAGAUGAAGCGGaCGACAGAAGAGAAGACGACGACGCCAGCGUGGCGAUGAUGAGGACGACGACAUCGCCGCUGCUGCCGCCACAGUCCGAGUCcGCUUGGUCUUCAGCUCGCUCuCACCGCAA   count    length
Libraries combined          Precision: [ total = 96.1% | 5p-arm = 95.9% | 3p-arm = 100.0% ]
----------AGCGGGCUGGAGAUGAAGCGGA---------------------------------------------------------------------------------------------------------------   55       22
----------AGCGGGCUGGAGAUGAAGCGG----------------------------------------------------------------------------------------------------------------   6        21
----------AGCGGGCUGGAGAUGAAGC------------------------------------------------------------------------------------------------------------------   6        19
----------AGCGGGCUGGAGAUGAAGCGGAC--------------------------------------------------------------------------------------------------------------   2        23
-------GAGAGCGGGCUGGAGAUGAAGCGGACGAC-----------------------------------------------------------------------------------------------------------   1        29
-------GAGAGCGGGCUGGAGAUGAAGCGGACG-------------------------------------------------------------------------------------------------------------   1        27
----------AGCGGGCUGGAGAUGAAGCG-----------------------------------------------------------------------------------------------------------------   1        20
-----------GCGGGCUGGAGAUGAAGCGG----------------------------------------------------------------------------------------------------------------   1        20
-----------------------------------------------------------------------------------------------------------------CGCUUGGUCUUCAGCUCGCUCU--------   3        22
Library = 1        Precision: [ total = 100.0% | 5p-arm = 100.0% | 3p-arm = 0% ]
----------AGCGGGCUGGAGAUGAAGCGGA---------------------------------------------------------------------------------------------------------------   23       22
----------AGCGGGCUGGAGAUGAAGCGG----------------------------------------------------------------------------------------------------------------   2        21
----------AGCGGGCUGGAGAUGAAGC------------------------------------------------------------------------------------------------------------------   1        19
----------AGCGGGCUGGAGAUGAAGCGGAC--------------------------------------------------------------------------------------------------------------   1        23
Library = 2        Precision: [ total = 92.0% | 5p-arm = 91.7% | 3p-arm = 100.0% ]
----------AGCGGGCUGGAGAUGAAGCGGA---------------------------------------------------------------------------------------------------------------   16       22
----------AGCGGGCUGGAGAUGAAGCGG----------------------------------------------------------------------------------------------------------------   3        21
----------AGCGGGCUGGAGAUGAAGC------------------------------------------------------------------------------------------------------------------   2        19
-------GAGAGCGGGCUGGAGAUGAAGCGGACGAC-----------------------------------------------------------------------------------------------------------   1        29
-------GAGAGCGGGCUGGAGAUGAAGCGGACG-------------------------------------------------------------------------------------------------------------   1        27
----------AGCGGGCUGGAGAUGAAGCG-----------------------------------------------------------------------------------------------------------------   1        20
-----------------------------------------------------------------------------------------------------------------CGCUUGGUCUUCAGCUCGCUCU--------   1        22
Library = 3        Precision: [ total = 95.8% | 5p-arm = 95.5% | 3p-arm = 100.0% ]
----------AGCGGGCUGGAGAUGAAGCGGA---------------------------------------------------------------------------------------------------------------   16       22
----------AGCGGGCUGGAGAUGAAGC------------------------------------------------------------------------------------------------------------------   3        19
-----------------------------------------------------------------------------------------------------------------CGCUUGGUCUUCAGCUCGCUCU--------   2        22
----------AGCGGGCUGGAGAUGAAGCGGAC--------------------------------------------------------------------------------------------------------------   1        23
----------AGCGGGCUGGAGAUGAAGCGG----------------------------------------------------------------------------------------------------------------   1        21
-----------GCGGGCUGGAGAUGAAGCGG----------------------------------------------------------------------------------------------------------------   1        20
```

```
AC A  A                  G        GACAGAAGAGAA   GAC A  C               A  AGG
  G GG GAGaGCGGGCUGGAGAU AAGCGGaC            GAC   G CG CAGC-GUGGCGAUG UG   \
  C CC CuCUCGCUCGACUUCUG UUCGcCUG   ----------CUG   C GC GUCG CGCCGCUAC GC   A
AA G  A                  G        AGC            ACA C  C    U         A  AGC
```

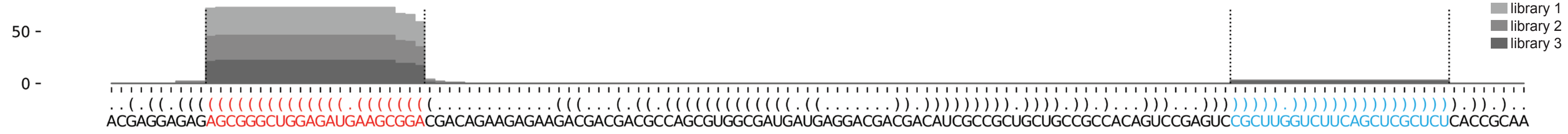

### *A. lenticulata* ale-mir-1228-P2

```
...((.(((((((((((.(((((.((((((((((..((.((((........)))))).)))))))))).))))).)))))))))))))....   -58.46 kcal/mol
AUUGUUCAUGuUUAGUGCCUAUGUCCUCUUCaCUCAUGAUCGCCAUCACGAGCGGCACAGUGaAGAGGUCAUAGACACUAAACaUGACUUUU   count    length
Libraries combined          Precision: [ total = 99.4% | 5p-arm = 99.3% | 3p-arm = 100.0% ]
----------UUUAGUGCCUAUGUCCUCUUCA------------------------------------------------------------   101      22
----------UUUAGUGCCUAUGUCCUCUUC-------------------------------------------------------------   116      21
----------UUUAGUGCCUAUGUCCUC----------------------------------------------------------------   91       18
----------UUUAGUGCCUAUGUCCUCUU--------------------------------------------------------------   94       20
----------UUUAGUGCCUAUGUCCUCU---------------------------------------------------------------   48       19
--------------------------------------------------------------AAGAGGUCAUAGACACUAAACA--------   9        22
-----------UUAGUGCCUAUGUCCUCUUCA------------------------------------------------------------   2        21
-----------UUAGUGCCUAUGUCCUCUUC-------------------------------------------------------------   1        20
--------------------------------------------------------------AAGAGGUCAUAGACACUAAAC---------   4        21
Library = 1        Precision: [ total = 100.0% | 5p-arm = 100.0% | 3p-arm = 100.0% ]
----------UUUAGUGCCUAUGUCCUCUUCA------------------------------------------------------------   39       22
----------UUUAGUGCCUAUGUCCUCUUC-------------------------------------------------------------   31       21
----------UUUAGUGCCUAUGUCCUC----------------------------------------------------------------   25       18
----------UUUAGUGCCUAUGUCCUCUU--------------------------------------------------------------   22       20
----------UUUAGUGCCUAUGUCCUCU---------------------------------------------------------------   10       19
--------------------------------------------------------------AAGAGGUCAUAGACACUAAACA--------   3        22
Library = 2        Precision: [ total = 98.6% | 5p-arm = 98.6% | 3p-arm = 100.0% ]
----------UUUAGUGCCUAUGUCCUCUUC-------------------------------------------------------------   59       21
----------UUUAGUGCCUAUGUCCUCUU--------------------------------------------------------------   45       20
----------UUUAGUGCCUAUGUCCUC----------------------------------------------------------------   40       18
----------UUUAGUGCCUAUGUCCUCUUCA------------------------------------------------------------   37       22
----------UUUAGUGCCUAUGUCCUCU---------------------------------------------------------------   28       19
-----------UUAGUGCCUAUGUCCUCUUCA------------------------------------------------------------   2        21
--------------------------------------------------------------AAGAGGUCAUAGACACUAAACA--------   2        22
--------------------------------------------------------------AAGAGGUCAUAGACACUAAAC---------   2        21
-----------UUAGUGCCUAUGUCCUCUUC-------------------------------------------------------------   1        20
Library = 3        Precision: [ total = 100.0% | 5p-arm = 100.0% | 3p-arm = 100.0% ]
----------UUUAGUGCCUAUGUCCUCUU--------------------------------------------------------------   27       20
----------UUUAGUGCCUAUGUCCUC----------------------------------------------------------------   26       18
----------UUUAGUGCCUAUGUCCUCUUC-------------------------------------------------------------   26       21
----------UUUAGUGCCUAUGUCCUCUUCA------------------------------------------------------------   25       22
----------UUUAGUGCCUAUGUCCUCU---------------------------------------------------------------   10       19
--------------------------------------------------------------AAGAGGUCAUAGACACUAAACA--------   4        22
--------------------------------------------------------------AAGAGGUCAUAGACACUAAAC---------   2        21
```

```
 AUU  U           C     U          CA  A    CAU
    GU CAUGuUUAGUG CUAUG CCUCUUCaCU  UG UCGC   C
    CA-GUaCAAAUCAC GAUAC GGAGAaGUGA -AC-GGCG   A
UUUU              A     U          C        AGC
```

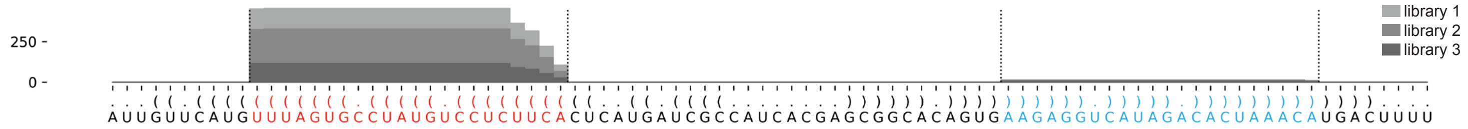

### *A. lenticulata* ale-mir-1230-P1

```
..((((((((.(((..(((((((((((.(((((((.((.(....).))))))))).)))))))))))..))).)))))..)))   -58.43 kcal/mol
CACCCUUUCCcACGCUCUCGGGAGACGAAGcACAAAGCGCAUAAGGGCUUGUGCuCCGUCUCUUGAGCUCGUCGgAAACUGGG   count    length
Libraries combined         Precision: [ total = 93.4% | 5p-arm = 99.2% | 3p-arm = 61.1% ]
----------CACGCUCUCGGGAGACGAAGC----------------------------------------------------   398      21
----------CACGCUCUCGGGAGACGAAG-----------------------------------------------------   158      20
----------CACGCUCUCGGGAGACGAAGCA---------------------------------------------------   61       22
------------------------------------------------------UCCGUCUCUUGAGCUCGUCGG--------   56       21
-----------------------------------------------------CUCCGUCUCUUGAGCUCGUCGG--------   16       22
-----------------------------------------------------CUCCGUCUCUUGAGCUCGUC----------   13       20
------------------------------------------------------UCCGUCUCUUGAGCUCGUCG---------   8        20
----------CACGCUCUCGGGAGACGA-------------------------------------------------------   4        18
-----------------------------------------------------CUCCGUCUCUUGAGCUCGUCG---------   3        21
-----------------------------------------------------CUCCGUCUCUUGAGCUCGU-----------   4        19
----------CACGCUCUCGGGAGACGAAGCAC--------------------------------------------------   2        23
----------------------------------------AUAAGGGCUUGUGCUCCGUCUCU--------------------   1        23
------------------------------------------AAGGGCUUGUGCUCCGUCUCUUGAGC---------------   1        26
------------------------------------------------UUGUGCUCCGUCUCUUGAGCUCGUC----------   1        25
-----------------------------------------------------CUCCGUCUCUUGAGCUCGUCGGA-------   2        23
------------------------------------------------------UCCGUCUCUUGAGCUCGU-----------   2        18
----------CACGCUCUCGGGAGACGAA------------------------------------------------------   7        19
-----------ACGCUCUCGGGAGACGAAGC----------------------------------------------------   4        20
----------------CUCGGGAGACGAAGCACA-------------------------------------------------   1        18
-------------------------------ACAAAGCGCAUAAGGGCUUGUGCUCCGUCUCU--------------------   1        32
----------------------------------------AUAAGGGCUUGUGCUCCGUCUCUUGAGCUCGUC----------   1        33
-------------------------------------------------UGUGCUCCGUCUCUUGAGCUC-------------   1        21
------------------------------------------------------UCCGUCUCUUGAGCUCGUC----------   3        19
Library = 1        Precision: [ total = 92.1% | 5p-arm = 100.0% | 3p-arm = 48.6% ]
----------CACGCUCUCGGGAGACGAAGC----------------------------------------------------   140      21
----------CACGCUCUCGGGAGACGAAG-----------------------------------------------------   44       20
----------CACGCUCUCGGGAGACGAAGCA---------------------------------------------------   17       22
------------------------------------------------------UCCGUCUCUUGAGCUCGUCGG--------   13       21
-----------------------------------------------------CUCCGUCUCUUGAGCUCGUCGG--------   6        22
-----------------------------------------------------CUCCGUCUCUUGAGCUCGUC----------   5        20
------------------------------------------------------UCCGUCUCUUGAGCUCGUCG---------   4        20
----------CACGCUCUCGGGAGACGA-------------------------------------------------------   2        18
-----------------------------------------------------CUCCGUCUCUUGAGCUCGUCG---------   2        21
-----------------------------------------------------CUCCGUCUCUUGAGCUCGU-----------   2        19
----------CACGCUCUCGGGAGACGAAGCAC--------------------------------------------------   1        23
----------------------------------------AUAAGGGCUUGUGCUCCGUCUCU--------------------   1        23
------------------------------------------AAGGGCUUGUGCUCCGUCUCUUGAGC---------------   1        26
------------------------------------------------UUGUGCUCCGUCUCUUGAGCUCGUC----------   1        25
-----------------------------------------------------CUCCGUCUCUUGAGCUCGUCGGA-------   1        23
------------------------------------------------------UCCGUCUCUUGAGCUCGU-----------   1        18
Library = 2        Precision: [ total = 95.0% | 5p-arm = 98.6% | 3p-arm = 68.4% ]
----------CACGCUCUCGGGAGACGAAGC----------------------------------------------------   164      21
----------CACGCUCUCGGGAGACGAAG-----------------------------------------------------   77       20
----------CACGCUCUCGGGAGACGAAGCA---------------------------------------------------   34       22
------------------------------------------------------UCCGUCUCUUGAGCUCGUCGG--------   21       21
-----------------------------------------------------CUCCGUCUCUUGAGCUCGUCGG--------   5        22
----------CACGCUCUCGGGAGACGAA------------------------------------------------------   4        19
-----------------------------------------------------CUCCGUCUCUUGAGCUCGUC----------   4        20
-----------ACGCUCUCGGGAGACGAAGC----------------------------------------------------   3        20
------------------------------------------------------UCCGUCUCUUGAGCUCGUC----------   3        19
----------CACGCUCUCGGGAGACGAAGCAC--------------------------------------------------   1        23
----------CACGCUCUCGGGAGACGA-------------------------------------------------------   1        18
```

### *A. lenticulata* ale-mir-1230-P1 continued

```
..((((((((.(((..(((((((((((.(((((((.((.(....).))))))))).)))))))))))..))).)))))..)))   -58.43 kcal/mol
CACCCUUUCCcACGCUCUCGGGAGACGAAGcACAAAGCGCAUAAGGGCUUGUGCuCCGUCUCUUGAGCUCGUCGgAAACUGGG   count    length
----------------CUCGGGAGACGAAGCACA-------------------------------------------------   1        18
-------------------------------------------------UGUGCUCCGUCUCUUGAGCUC-------------   1        21
-----------------------------------------------------CUCCGUCUCUUGAGCUCGUCG---------   1        21
-----------------------------------------------------CUCCGUCUCUUGAGCUCGUCGGA-------   1        23
------------------------------------------------------UCCGUCUCUUGAGCUCGUCG---------   1        20
------------------------------------------------------UCCGUCUCUUGAGCUCGU-----------   1        18
Library = 3       Precision: [ total = 92.4% | 5p-arm = 99.3% | 3p-arm = 65.8% ]
----------CACGCUCUCGGGAGACGAAGC----------------------------------------------------   94       21
----------CACGCUCUCGGGAGACGAAG-----------------------------------------------------   37       20
------------------------------------------------------UCCGUCUCUUGAGCUCGUCGG--------   22       21
----------CACGCUCUCGGGAGACGAAGCA---------------------------------------------------   10       22
-----------------------------------------------------CUCCGUCUCUUGAGCUCGUCGG--------   5        22
-----------------------------------------------------CUCCGUCUCUUGAGCUCGUC----------   4        20
----------CACGCUCUCGGGAGACGAA------------------------------------------------------   3        19
------------------------------------------------------UCCGUCUCUUGAGCUCGUCG---------   3        20
-----------------------------------------------------CUCCGUCUCUUGAGCUCGU-----------   2        19
----------CACGCUCUCGGGAGACGA-------------------------------------------------------   1        18
-----------ACGCUCUCGGGAGACGAAGC----------------------------------------------------   1        20
-------------------------------ACAAAGCGCAUAAGGGCUUGUGCUCCGUCUCU--------------------   1        32
----------------------------------------AUAAGGGCUUGUGCUCCGUCUCUUGAGCUCGUC----------   1        33

CA          c   CU          A       A  G A
  CCC--UUUCC ACG  CUCGGGAGACG AGcACAA GC C U
  GGG  AAAgG UGC  GAGUUCUCUGC uCGUGUU-CG G A
     UC      C    UC          C          G A
```

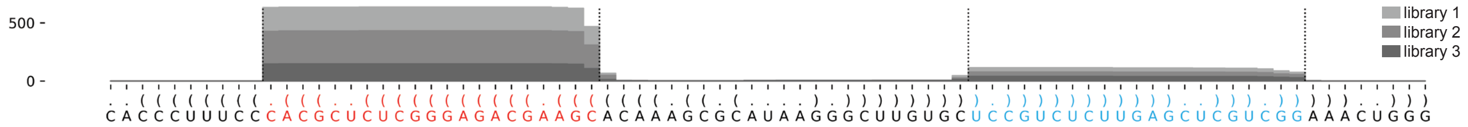

### *A. lenticulata* ale-mir-1230-P2

```
..((((((((.(((..(((((((((((.(((((((.((((((....))))))))))))).)))))))))))..))).)))))..)))    -65.24 kcal/mol
CACCCUUUCCcACGCUCUCGGGAGACGAAGcACAAAGCCCUUAUAUAAGGGCUUGUGCuCCGUCUCUUGAGCUCGUCGgAAACUGGG    count    length
Libraries combined         Precision: [ total = 93.3% | 5p-arm = 98.9% | 3p-arm = 61.6% ]
----------CACGCUCUCGGGAGACGAAGC--------------------------------------------------------    398      21
----------CACGCUCUCGGGAGACGAAG---------------------------------------------------------    158      20
----------CACGCUCUCGGGAGACGAAGCA-------------------------------------------------------    61       22
----------------------------------------------------------UCCGUCUCUUGAGCUCGUCGG--------    56       21
---------------------------------------------------------CUCCGUCUCUUGAGCUCGUCGG--------    16       22
---------------------------------------------------------CUCCGUCUCUUGAGCUCGUC----------    13       20
----------------------------------------------------------UCCGUCUCUUGAGCUCGUCG---------    8        20
----------CACGCUCUCGGGAGACGA-----------------------------------------------------------    4        18
---------------------------------------------------------CUCCGUCUCUUGAGCUCGUCG---------    3        21
---------------------------------------------------------CUCCGUCUCUUGAGCUCGU-----------    4        19
----------CACGCUCUCGGGAGACGAAGCAC------------------------------------------------------    2        23
--------------------------------------------AUAAGGGCUUGUGCUCCGUCUCU--------------------    1        23
----------------------------------------------AAGGGCUUGUGCUCCGUCUCUUGAGC---------------    1        26
----------------------------------------------------UUGUGCUCCGUCUCUUGAGCUCGUC----------    1        25
---------------------------------------------------------CUCCGUCUCUUGAGCUCGUCGGA-------    2        23
----------------------------------------------------------UCCGUCUCUUGAGCUCGU-----------    2        18
----------CACGCUCUCGGGAGACGAA----------------------------------------------------------    7        19
-----------ACGCUCUCGGGAGACGAAGC--------------------------------------------------------    4        20
----------------CUCGGGAGACGAAGCACA-----------------------------------------------------    1        18
----------------CUCGGGAGACGAAGCACAAAGCCC-----------------------------------------------    1        24
---------------------GAGACGAAGCACAAAGCCC-----------------------------------------------    1        19
--------------------------------------------AUAAGGGCUUGUGCUCCGUCUCUUGAGCUCGUC----------    1        33
-----------------------------------------------------UGUGCUCCGUCUCUUGAGCUC-------------    1        21
----------------------------------------------------------UCCGUCUCUUGAGCUCGUC----------    3        19
Library = 1       Precision: [ total = 92.1% | 5p-arm = 100.0% | 3p-arm = 48.6% ]
----------CACGCUCUCGGGAGACGAAGC--------------------------------------------------------    140      21
----------CACGCUCUCGGGAGACGAAG---------------------------------------------------------    44       20
----------CACGCUCUCGGGAGACGAAGCA-------------------------------------------------------    17       22
----------------------------------------------------------UCCGUCUCUUGAGCUCGUCGG--------    13       21
---------------------------------------------------------CUCCGUCUCUUGAGCUCGUCGG--------    6        22
---------------------------------------------------------CUCCGUCUCUUGAGCUCGUC----------    5        20
----------------------------------------------------------UCCGUCUCUUGAGCUCGUCG---------    4        20
----------CACGCUCUCGGGAGACGA-----------------------------------------------------------    2        18
---------------------------------------------------------CUCCGUCUCUUGAGCUCGUCG---------    2        21
---------------------------------------------------------CUCCGUCUCUUGAGCUCGU-----------    2        19
----------CACGCUCUCGGGAGACGAAGCAC------------------------------------------------------    1        23
--------------------------------------------AUAAGGGCUUGUGCUCCGUCUCU--------------------    1        23
----------------------------------------------AAGGGCUUGUGCUCCGUCUCUUGAGC---------------    1        26
----------------------------------------------------UUGUGCUCCGUCUCUUGAGCUCGUC----------    1        25
---------------------------------------------------------CUCCGUCUCUUGAGCUCGUCGGA-------    1        23
----------------------------------------------------------UCCGUCUCUUGAGCUCGU-----------    1        18
Library = 2       Precision: [ total = 94.8% | 5p-arm = 98.3% | 3p-arm = 68.4% ]
----------CACGCUCUCGGGAGACGAAGC--------------------------------------------------------    164      21
----------CACGCUCUCGGGAGACGAAG---------------------------------------------------------    77       20
----------CACGCUCUCGGGAGACGAAGCA-------------------------------------------------------    34       22
----------------------------------------------------------UCCGUCUCUUGAGCUCGUCGG--------    21       21
---------------------------------------------------------CUCCGUCUCUUGAGCUCGUCGG--------    5        22
----------CACGCUCUCGGGAGACGAA----------------------------------------------------------    4        19
---------------------------------------------------------CUCCGUCUCUUGAGCUCGUC----------    4        20
-----------ACGCUCUCGGGAGACGAAGC--------------------------------------------------------    3        20
----------------------------------------------------------UCCGUCUCUUGAGCUCGUC----------    3        19
----------CACGCUCUCGGGAGACGAAGCAC------------------------------------------------------    1        23
----------CACGCUCUCGGGAGACGA-----------------------------------------------------------    1        18
```

### *A. lenticulata* ale-mir-1230-P2 continued

```
..((((((((.(((..(((((((((((.(((((((.((((((....))))))))))))).)))))))))))..))).)))))..)))      -65.24 kcal/mol
CACCCUUUCCcACGCUCUCGGGAGACGAAGcACAAAGCCCUUAUAUAAGGGCUUGUGCuCCGUCUCUUGAGCUCGUCGgAAACUGGG      count    length
----------------CUCGGGAGACGAAGCACA-----------------------------------------------------      1        18
----------------CUCGGGAGACGAAGCACAAAGCCC-----------------------------------------------      1        24
-----------------------------------------------------UGUGCUCCGUCUCUUGAGCUC-------------      1        21
---------------------------------------------------------CUCCGUCUCUUGAGCUCGUCG---------      1        21
---------------------------------------------------------CUCCGUCUCUUGAGCUCGUCGGA-------      1        23
----------------------------------------------------------UCCGUCUCUUGAGCUCGUCG---------      1        20
----------------------------------------------------------UCCGUCUCUUGAGCUCGU-----------      1        18
Library = 3       Precision: [ total = 92.4% | 5p-arm = 98.6% | 3p-arm = 67.6% ]
----------CACGCUCUCGGGAGACGAAGC--------------------------------------------------------      94       21
----------CACGCUCUCGGGAGACGAAG---------------------------------------------------------      37       20
----------------------------------------------------------UCCGUCUCUUGAGCUCGUCGG--------      22       21
----------CACGCUCUCGGGAGACGAAGCA-------------------------------------------------------      10       22
---------------------------------------------------------CUCCGUCUCUUGAGCUCGUCGG--------      5        22
---------------------------------------------------------CUCCGUCUCUUGAGCUCGUC----------      4        20
----------CACGCUCUCGGGAGACGAA----------------------------------------------------------      3        19
----------------------------------------------------------UCCGUCUCUUGAGCUCGUCG---------      3        20
---------------------------------------------------------CUCCGUCUCUUGAGCUCGU-----------      2        19
----------CACGCUCUCGGGAGACGA-----------------------------------------------------------      1        18
-----------ACGCUCUCGGGAGACGAAGC--------------------------------------------------------      1        20
---------------------GAGACGAAGCACAAAGCCC-----------------------------------------------      1        19
--------------------------------------------AUAAGGGCUUGUGCUCCGUCUCUUGAGCUCGUC----------      1        33
```

```
CA          c   CU           A       A      A
  CCC--UUUCC ACG  CUCGGGAGACG AGcACAA GCCCUU U
  GGG  AAAgG UGC  GAGUUCUCUGC uCGUGUU-CGGGAA A
     UC     C   UC           C              U
```

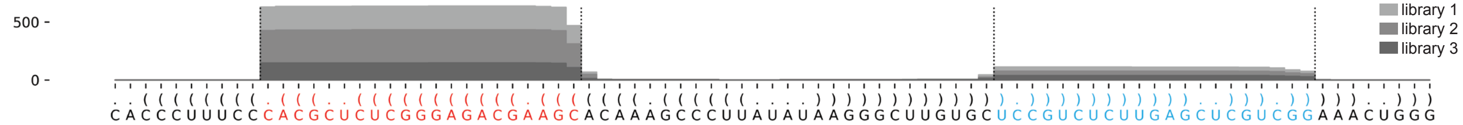

**Supplementary Fig. 7: Detailed depiction of identified miRNAs in P. polycephalum, A. castellanii and A. lenticulata.** Predicted miRNA hairpin sequences and structure displayed in bracket notion. Identified miRNA-5p is indicated in red, and the miRNA-3p in cyan. Lowercase nucleotides represent the 5' and 3' ends of the miRNA-5p and miRNA-3p. Below the sequence, all mapped small RNA reads are aligned to the miRNA hairpin and the number reads and their length are shown to the right. Results are shown for all small RNA sequencing libraries combined (denoted Libraries combined) as well as for the individual libraries (denoted Library = 1 to 3). The percentage of reads that map to the exact 5' nucleotide of either the miRNA-5p or miRNA-3p is shown as 'precision' and calculated on the entire miRNA-hairpin (denoted total), and on the 5p- and 3p-arms of the miRNA hairpin (denoted 5p-arm and 3p-arm respectively). The folded miRNA hairpin structure, shown below the mapped small RNA, was predicted using the ViennaRNA RNAlib-2.6.2 python package, with 22°C folding temperature. At the bottom, a graph shows the read mapping density on the miRNA hairpin, with the miRNA-5p and miRNA-3p indicated in red and cyan respectively. The different shades of grey in the graph represent the different small RNA sequencing libraries. From some miRNA hairpins, two miRNA duplexes might be processed in tandem, in which case the two miRNA-5p sequences are colored red and yellow, and the two miRNA-3p sequences are colored dark blue and cyan.
