## Supplementary Data 3 for "Evolution of microRNAs in Amoebozoa and implications for the origin of multicellularity"

### *D. firmibasis* dfi-mir-1177

```
....((((((.(((..((((((((((((((((((((.((((((((((((((.((((.((((......(((((.(((((..(((....))).))))))))))))))..))))))))))))))))))))))))))))))))))))))..))).))))))....   -69.66 kcal/mol
UAAAAAUAAAcCAGUUAGGGUUUAAUGGUUcUUGUUUAAAUUUUUUUUUUUUGGAUUCAAAAUUUAUUAAUCGAUUUUCCUAAUAAUUUAAAAAAUGAUUAUUUGUUAUCCAAAAAAAAGAAUUUAACAAGAgCCGUUGAGUCCUUUCUGAUuUAUUAAUU   count   length
Libraries combined            Precision: [ total = 96.2% | 5p-arm = 96.4% | 3p-arm = 95.7% ]
----------CCAGUUAGGGUUUAAUGGUUC----------------------------------------------------------------------------------------------------------------------------------   650   21
------------------------------------------------------------------------------------------------------------------------------------GCCGUUGAGUCCUUUCUGAU---------   191   20
------------------------------------------------------------------------------------------------------------------------------------GCCGUUGAGUCCUUUCUGAUU--------   137   21
------------------------------------------------------------------------------------------------------------------------------------GCCGUUGAGUCCUUUCUGA----------   41   19
----------CCAGUUAGGGUUUAAUGGUUCU---------------------------------------------------------------------------------------------------------------------------------   20   22
------------AGUUAGGGUUUAAUGGUUCUUG-------------------------------------------------------------------------------------------------------------------------------   17   22
------------------------------------------------------------------------------------------------------------------------------------GCCGUUGAGUCCUUUCUG-----------   26   18
-----------CAGUUAGGGUUUAAUGGUUC----------------------------------------------------------------------------------------------------------------------------------   3   20
----------------------------------------------------------------------------------------------------------------------------UAACAAGAGCCGUUGAGUCCU----------------   8   21
--------------------------------------------------------------------------------------------------------------------------------------CGUUGAGUCCUUUCUGAUUUA------   3   21
----------CCAGUUAGGGUUUAAUGGUUCUU--------------------------------------------------------------------------------------------------------------------------------   1   23
----------CCAGUUAGGGUUUAAUGGU------------------------------------------------------------------------------------------------------------------------------------   1   19
------------AGUUAGGGUUUAAUGGUUCUU--------------------------------------------------------------------------------------------------------------------------------   1   21
-----------------------------------------------------------------------------------------------------------------------GAAUUUAACAAGAGCCGUUG----------------------   1   20
-----------------------------------------------------------------------------------------------------------------------------AACAAGAGCCGUUGAGUCCU----------------   1   20
------------------------------------------------------------------------------------------------------------------------------------GCCGUUGAGUCCUUUCUGAUUU-------   4   22
----------CCAGUUAGGGUUUAAUGGUU-----------------------------------------------------------------------------------------------------------------------------------   3   20
-----------CAGUUAGGGUUUAAUGGUUCU---------------------------------------------------------------------------------------------------------------------------------   1   21
-------------------------------UUGUUUAAAUUUUUUUUUUUUGG-----------------------------------------------------------------------------------------------------------   3   23
------------------------------------------------------------------------------------------------------------------------------ACAAGAGCCGUUGAGUCCU----------------   1   19
---------------------------------------------------------------------------------------------------------------------------------AGAGCCGUUGAGUCCUUUCUG-----------   1   21
-----------------------------------------------------------------------------------------------------------------------------------AGCCGUUGAGUCCUUUCUGAU---------   1   21
-----------------------------------------------------------------------------------------------------------------------------------AGCCGUUGAGUCCUUUCUGAUU--------   1   22
---------------------------------------------------------------------------------------------------------------------------------------GUUGAGUCCUUUCUGAUU--------   1   18
Library = 1         Precision: [ total = 95.8% | 5p-arm = 96.1% | 3p-arm = 95.3% ]
----------CCAGUUAGGGUUUAAUGGUUC----------------------------------------------------------------------------------------------------------------------------------   238   21
------------------------------------------------------------------------------------------------------------------------------------GCCGUUGAGUCCUUUCUGAU---------   59   20
------------------------------------------------------------------------------------------------------------------------------------GCCGUUGAGUCCUUUCUGAUU--------   41   21
------------------------------------------------------------------------------------------------------------------------------------GCCGUUGAGUCCUUUCUGA----------   13   19
----------CCAGUUAGGGUUUAAUGGUUCU---------------------------------------------------------------------------------------------------------------------------------   8   22
------------AGUUAGGGUUUAAUGGUUCUUG-------------------------------------------------------------------------------------------------------------------------------   7   22
------------------------------------------------------------------------------------------------------------------------------------GCCGUUGAGUCCUUUCUG-----------   7   18
-----------CAGUUAGGGUUUAAUGGUUC----------------------------------------------------------------------------------------------------------------------------------   2   20
----------------------------------------------------------------------------------------------------------------------------UAACAAGAGCCGUUGAGUCCU----------------   2   21
--------------------------------------------------------------------------------------------------------------------------------------CGUUGAGUCCUUUCUGAUUUA------   2   21
----------CCAGUUAGGGUUUAAUGGUUCUU--------------------------------------------------------------------------------------------------------------------------------   1   23
----------CCAGUUAGGGUUUAAUGGU------------------------------------------------------------------------------------------------------------------------------------   1   19
------------AGUUAGGGUUUAAUGGUUCUU--------------------------------------------------------------------------------------------------------------------------------   1   21
-----------------------------------------------------------------------------------------------------------------------GAAUUUAACAAGAGCCGUUG----------------------   1   20
-----------------------------------------------------------------------------------------------------------------------------AACAAGAGCCGUUGAGUCCU----------------   1   20
------------------------------------------------------------------------------------------------------------------------------------GCCGUUGAGUCCUUUCUGAUUU-------   1   22
Library = 2         Precision: [ total = 95.8% | 5p-arm = 96.5% | 3p-arm = 94.5% ]
----------CCAGUUAGGGUUUAAUGGUUC----------------------------------------------------------------------------------------------------------------------------------   209   21
------------------------------------------------------------------------------------------------------------------------------------GCCGUUGAGUCCUUUCUGAU---------   61   20
------------------------------------------------------------------------------------------------------------------------------------GCCGUUGAGUCCUUUCUGAUU--------   38   21
------------------------------------------------------------------------------------------------------------------------------------GCCGUUGAGUCCUUUCUGA----------   17   19
----------CCAGUUAGGGUUUAAUGGUUCU---------------------------------------------------------------------------------------------------------------------------------   8   22
----------------------------------------------------------------------------------------------------------------------------UAACAAGAGCCGUUGAGUCCU----------------   4   21
------------------------------------------------------------------------------------------------------------------------------------GCCGUUGAGUCCUUUCUG-----------   4   18
------------AGUUAGGGUUUAAUGGUUCUUG-------------------------------------------------------------------------------------------------------------------------------   3   22
-------------------------------UUGUUUAAAUUUUUUUUUUUUGG-----------------------------------------------------------------------------------------------------------   3   23
----------CCAGUUAGGGUUUAAUGGUU-----------------------------------------------------------------------------------------------------------------------------------   1   20
-----------CAGUUAGGGUUUAAUGGUUC----------------------------------------------------------------------------------------------------------------------------------   1   20
```

### *D. firmibasis* dfi-mir-1177 continued

```
....((((((.(((..((((((((((((((((((((.((((((((((((((.((((.((((......(((((.(((((..(((....))).))))))))))))))..))))))))))))))))))))))))))))))))))))))..))).))))))....   -69.66 kcal/mol
UAAAAAUAAAcCAGUUAGGGUUUAAUGGUUcUUGUUUAAAUUUUUUUUUUUUGGAUUCAAAAUUUAUUAAUCGAUUUUCCUAAUAAUUUAAAAAAUGAUUAUUUGUUAUCCAAAAAAAAGAAUUUAACAAGAgCCGUUGAGUCCUUUCUGAUuUAUUAAUU   count   length
-----------CAGUUAGGGUUUAAUGGUUCU---------------------------------------------------------------------------------------------------------------------------------   1   21
-----------------------------------------------------------------------------------------------------------------------------------AGCCGUUGAGUCCUUUCUGAU---------   1   21
------------------------------------------------------------------------------------------------------------------------------------GCCGUUGAGUCCUUUCUGAUUU-------   1   22
--------------------------------------------------------------------------------------------------------------------------------------CGUUGAGUCCUUUCUGAUUUA------   1   21
---------------------------------------------------------------------------------------------------------------------------------------GUUGAGUCCUUUCUGAUU--------   1   18
Library = 3          Precision: [ total = 96.8% | 5p-arm = 96.8% | 3p-arm = 96.9% ]
----------CCAGUUAGGGUUUAAUGGUUC----------------------------------------------------------------------------------------------------------------------------------   203   21
------------------------------------------------------------------------------------------------------------------------------------GCCGUUGAGUCCUUUCUGAU---------   71   20
------------------------------------------------------------------------------------------------------------------------------------GCCGUUGAGUCCUUUCUGAUU--------   58   21
------------------------------------------------------------------------------------------------------------------------------------GCCGUUGAGUCCUUUCUG-----------   15   18
------------------------------------------------------------------------------------------------------------------------------------GCCGUUGAGUCCUUUCUGA----------   11   19
------------AGUUAGGGUUUAAUGGUUCUUG-------------------------------------------------------------------------------------------------------------------------------   7   22
----------CCAGUUAGGGUUUAAUGGUUCU---------------------------------------------------------------------------------------------------------------------------------   4   22
----------CCAGUUAGGGUUUAAUGGUU-----------------------------------------------------------------------------------------------------------------------------------   2   20
----------------------------------------------------------------------------------------------------------------------------UAACAAGAGCCGUUGAGUCCU----------------   2   21
------------------------------------------------------------------------------------------------------------------------------------GCCGUUGAGUCCUUUCUGAUUU-------   2   22
------------------------------------------------------------------------------------------------------------------------------ACAAGAGCCGUUGAGUCCU----------------   1   19
---------------------------------------------------------------------------------------------------------------------------------AGAGCCGUUGAGUCCUUUCUG-----------   1   21
-----------------------------------------------------------------------------------------------------------------------------------AGCCGUUGAGUCCUUUCUGAUU--------   1   22
```

```
UAAA      c   UU                        U               U     U-     AUUUAU      G      CC    U
    AAUAAA CAG  AGGGUUUAAUGGUUcUUGUU AAAUUUUUUUUUUU GGAU  CAAA      UAAUC AUUUU   UAA A
    UUAUuU GUC  UCCUGAGUUGCCgAGAACAA-UUUAAGAAAAAAAA-CCUA  GUUU------AUUAG-UAAAA   AUU A
UUAA      A   UU                                        UU                    A-    U
```

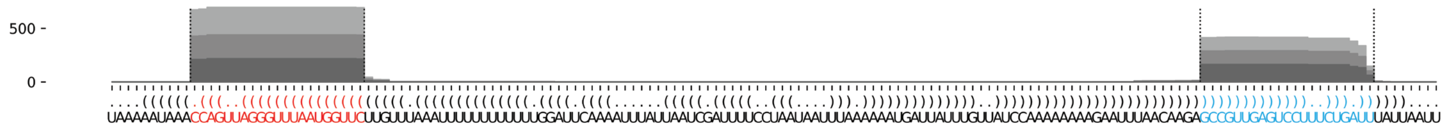


### *D. firmibasis* dfi-mir-1189

```
(((((((((((((((((((((((((((((((((((((((((((((((((......)))))))))))))))))))))))))))))))))))))))))))))))))   -108.21 kcal/mol
GGAAGUAAAGcAUGGAAGAUUAAUGACCGCcUACCAAAGACAAAUAGGAAUCAAAUCCUAUUUGUCUUUGGUAGGcGGUCAUUAAUCUUCCAUGCuUUACUUCC   count   length
Libraries combined          Precision: [ total = 76.3% | 5p-arm = 72.2% | 3p-arm = 96.4% ]

----------CAUGGAAGAUUAAUGACCGCC-------------------------------------------------------------------------   225704   21
-----------AUGGAAGAUUAAUGACCGCCU------------------------------------------------------------------------   84956   21
---------------------------------------------------------------------------CGGUCAUUAAUCUUCCAUGCU--------   63851   21
----------CAUGGAAGAUUAAUGACCGCCU------------------------------------------------------------------------   16011   22
-----------AUGGAAGAUUAAUGACCGCC-------------------------------------------------------------------------   4437   20
---------GCAUGGAAGAUUAAUGACCGCC-------------------------------------------------------------------------   1197   22
---------GCAUGGAAGAUUAAUGACCGC--------------------------------------------------------------------------   743   21
---------------------------------------------------------------------------CGGUCAUUAAUCUUCCAUGCUU-------   1266   22
---------------------------------------------------------------------------CGGUCAUUAAUCUUCCAUGC---------   962   20
----------CAUGGAAGAUUAAUGACCG---------------------------------------------------------------------------   562   19
--------AGCAUGGAAGAUUAAUGACCGC--------------------------------------------------------------------------   520   22
----------------------------------------------------------------------------GGUCAUUAAUCUUCCAUGCUU-------   844   21
--------------------------------------------------------------------------GCGGUCAUUAAUCUUCCAUGC---------   573   21
--------AGCAUGGAAGAUUAAUGACCGCC-------------------------------------------------------------------------   411   23
----------CAUGGAAGAUUAAUGACCGC--------------------------------------------------------------------------   470   20
--------AGCAUGGAAGAUUAAUGACCG---------------------------------------------------------------------------   367   21
--------------------UAAUGACCGCCUACCAAAGAC---------------------------------------------------------------   145   21
----------------------------------------------------------------------------GGUCAUUAAUCUUCCAUGCU--------   223   20
---------GCAUGGAAGAUUAAUGACCGCCU------------------------------------------------------------------------   121   23
```

### *D. firmibasis* dfi-mir-1189 continued

```
(((((((((((((((((((((((((((((((((((((((((((((((((......)))))))))))))))))))))))))))))))))))))))))))))))))       -108.21 kcal/mol
GGAAGUAAAGcAUGGAAGAUUAAUGACCGCcUACCAAAGACAAAUAGGAAUCAAAUCCUAUUUGUCUUUGGUAGGcGGUCAUUAAUCUUCCAUGCuUUACUUCC       count   length
-------------------------------UACCAAAGACAAAUAGGAAUCAAAUCCUAUUU-----------------------------------------       54      32
---------------------------------------------------------CUAUUUGUCUUUGGUAGGC----------------------------       87      19
--------------GAAGAUUAAUGACCGCCUACC---------------------------------------------------------------------       143     21
----------CAUGGAAGAUUAAUGACC----------------------------------------------------------------------------       57      18
-------AAGCAUGGAAGAUUAAUGACC----------------------------------------------------------------------------       63      21
-----------------------------------------------------------------------UAGGCGGUCAUUAAUCUUCCA------------       82      21
--------AGCAUGGAAGAUUAAUGACCGCCU------------------------------------------------------------------------       63      24
---------GCAUGGAAGAUUAAUGACCG---------------------------------------------------------------------------       56      20
-------------------------------------------------------UCCUAUUUGUCUUUGGUAGGC----------------------------       46      21
--------------------------------------------------------------------------GCGGUCAUUAAUCUUCCAUG----------       58      20
-----------AUGGAAGAUUAAUGACCGC--------------------------------------------------------------------------       37      19
-------------------------------UACCAAAGACAAAUAGGAAUCAAAUCCU---------------------------------------------       55      28
-----------------------------------------------------------------------UAGGCGGUCAUUAAUCUUCC-------------       67      20
-----------------------------------------------------------------------------GUCAUUAAUCUUCCAUGCU--------       39      19
------------------AUUAAUGACCGCCUACCAAAG-----------------------------------------------------------------       40      21
-------------------------------UACCAAAGACAAAUAGGAAUCAAAUC-----------------------------------------------       29      26
---------------------------------------------------------------------------CGGUCAUUAAUCUUCCAUG----------       41      19
-------------------------------UACCAAAGACAAAUAGGAAUCAAAUCCUAU-------------------------------------------       46      30
------------------------------------------------------------------UUUGGUAGGCGGUCAUUAAUC-----------------       21      21
--------------GAAGAUUAAUGACCGCCUAC----------------------------------------------------------------------       32      20
-----------------------------------------------------AAUCCUAUUUGUCUUUGGUAG------------------------------       27      21
------------------------------------------------------------------------AGGCGGUCAUUAAUCUUCCAUG----------       61      22
--------------------------------------------------------------------------GCGGUCAUUAAUCUUCCAUGCU--------       42      22
-----------------------------------------------------------AUUUGUCUUUGGUAGGCGGUC------------------------       15      21
-----------AUGGAAGAUUAAUGACCGCCUA-----------------------------------------------------------------------       24      22
-------AAGCAUGGAAGAUUAAUGACCG---------------------------------------------------------------------------       21      22
--------------GAAGAUUAAUGACCGCCU------------------------------------------------------------------------       16      18
------AAAGCAUGGAAGAUUAAUGAC-----------------------------------------------------------------------------       19      21
------------------------------------------------------------------------------UCAUUAAUCUUCCAUGCU--------       8       18
----GUAAAGCAUGGAAGAUUAAUG-------------------------------------------------------------------------------       5       21
-------------------------------UACCAAAGACAAAUAGGAAUCAAAUCCUA--------------------------------------------       9       29
--------------------------------------------------------CCUAUUUGUCUUUGGUAGGCG---------------------------       10      21
---------------------------------------------------------CUAUUUGUCUUUGGUAGGCGG--------------------------       7       21
---------------------------------------------------------------------GGUAGGCGGUCAUUAAUCUUC--------------       10      21
-----------------------------------------------------------------------------------AAUCUUCCAUGCUUUACUUCC       17      21
-------AAGCAUGGAAGAUUAAUGACCGCC-------------------------------------------------------------------------       5       24
------------------------------------------------------------------------AGGCGGUCAUUAAUCUUCCA------------       7       20
----------------------------------------------------------------------------GGUCAUUAAUCUUCCAUGCUUU------       41      22
-----------------------------------------------------------------------------GUCAUUAAUCUUCCAUGCUUU------       6       21
--------------------------------------------------------------------------------AUUAAUCUUCCAUGCUUUACU---       19      21
--------AGCAUGGAAGAUUAAUGACC----------------------------------------------------------------------------       9       20
---------GCAUGGAAGAUUAAUGACC----------------------------------------------------------------------------       5       19
-------------------------------UACCAAAGACAAAUAGGAAUCAAAUCCUAUU------------------------------------------       4       31
-------------------------------------------------------UCCUAUUUGUCUUUGGUA-------------------------------       4       18
GGAAGUAAAGCAUGGAAGAUU-----------------------------------------------------------------------------------       3       21
------AAAGCAUGGAAGAUUAAUGACCG---------------------------------------------------------------------------       5       23
-------AAGCAUGGAAGAUUAAUGAC-----------------------------------------------------------------------------       6       20
----------CAUGGAAGAUUAAUGACCGCCUA-----------------------------------------------------------------------       3       23
-----------AUGGAAGAUUAAUGACCG---------------------------------------------------------------------------       7       18
-------------------UUAAUGACCGCCUACCAAAGA----------------------------------------------------------------       5       21
-------------------UUAAUGACCGCCUACCAAAG-----------------------------------------------------------------       6       20
-------------------------------UACCAAAGACAAAUAGGAAUCAAAUCC----------------------------------------------       17      27
-------------------------------------------------------UCCUAUUUGUCUUUGGUAG------------------------------       4       19
----------------------------------------------------------UAUUUGUCUUUGGUAGGCGG---------------------------       2       20
```

### *D. firmibasis* dfi-mir-1189 continued

```
(((((((((((((((((((((((((((((((((((((((((((((((((......)))))))))))))))))))))))))))))))))))))))))))))))))   -108.21 kcal/mol
GGAAGUAAAGcAUGGAAGAUUAAUGACCGCcUACCAAAGACAAAUAGGAAUCAAAUCCUAUUUGUCUUUGGUAGGcGGUCAUUAAUCUUCCAUGCuUUACUUCC   count   length
-----------------------------------------------------------AUUUGUCUUUGGUAGGCGGU-------------------------   2   20
----------------------------------------------------------------UCUUUGGUAGGCGGUCAUUAA-------------------   3   21
----------------------------------------------------------------UCUUUGGUAGGCGGUCAUUA--------------------   3   20
----------------------------------------------------------------UCUUUGGUAGGCGGUCAUUAAU------------------   3   22
-----------------------------------------------------------------------UAGGCGGUCAUUAAUCUUC--------------   3   19
------------------------------------------------------------------------AGGCGGUCAUUAAUCUUCCAU-----------   6   21
------------------------------------------------------------------------AGGCGGUCAUUAAUCUUCC-------------   8   19
-----UAAAGCAUGGAAGAUUAAUGAC-----------------------------------------------------------------------------   1   22
-----UAAAGCAUGGAAGAUUAAUGACCG---------------------------------------------------------------------------   3   24
------AAAGCAUGGAAGAUUAAUGACC----------------------------------------------------------------------------   3   22
---------GCAUGGAAGAUUAAUGAC-----------------------------------------------------------------------------   1   18
-------------GGAAGAUUAAUGACCGCCUAC----------------------------------------------------------------------   4   21
-------------GGAAGAUUAAUGACCGCCUACC---------------------------------------------------------------------   2   22
-------------GGAAGAUUAAUGACCGCCUA-----------------------------------------------------------------------   2   20
-------------------------------UACCAAAGACAAAUAGGAAUC----------------------------------------------------   1   21
-------------------------------UACCAAAGACAAAUAGGAAUCAAAU------------------------------------------------   4   25
--------------------------------ACCAAAGACAAAUAGGAAUCAAAUCCUAUU------------------------------------------   1   30
-------------------------------------------AUAGGAAUCAAAUCCUAUUUGUCUUUGGUAGG-----------------------------   1   32
--------------------------------------------------------CCUAUUUGUCUUUGGUAG------------------------------   2   18
---------------------------------------------------------CUAUUUGUCUUUGGUAGG-----------------------------   3   18
----------------------------------------------------------UAUUUGUCUUUGGUAGGC----------------------------   1   18
--------------------------------------------------------------UGUCUUUGGUAGGCGGUC------------------------   1   18
--------------------------------------------------------------UGUCUUUGGUAGGCGGUCAUU---------------------   6   21
--------------------------------------------------------------UGUCUUUGGUAGGCGGUCAUUA--------------------   1   22
-----------------------------------------------------------------CUUUGGUAGGCGGUCAUUAAU------------------   7   21
-------------------------------------------------------------------UUGGUAGGCGGUCAUUAAUC-----------------   1   20
--------------------------------------------------------------------UGGUAGGCGGUCAUUAAUCUUC--------------   4   22
------------------------------------------------------------------------AGGCGGUCAUUAAUCUUC--------------   6   18
-------------------------------------------------------------------------GGCGGUCAUUAAUCUUCCAUG----------   3   21
----------------------------------------------------------------------------GGUCAUUAAUCUUCCAUG----------   9   18
---------------------------------------------------------------------------------UUAAUCUUCCAUGCUUUACUU--   1   21
-----UAAAGCAUGGAAGAUUAAUGA------------------------------------------------------------------------------   1   21
-----UAAAGCAUGGAAGAUUAAUG-------------------------------------------------------------------------------   1   20
-------AAGCAUGGAAGAUUAAUGACCGC--------------------------------------------------------------------------   2   23
-------AAGCAUGGAAGAUUAAUGACCGCCU------------------------------------------------------------------------   1   25
--------AGCAUGGAAGAUUAAUGAC-----------------------------------------------------------------------------   1   19
----------CAUGGAAGAUUAAUGACCGCCUACC---------------------------------------------------------------------   1   25
-----------AUGGAAGAUUAAUGACCGCCUACC---------------------------------------------------------------------   2   24
------------UGGAAGAUUAAUGACCGCCU------------------------------------------------------------------------   2   20
-------------GGAAGAUUAAUGACCGCC-------------------------------------------------------------------------   1   18
---------------AAGAUUAAUGACCGCCUAC----------------------------------------------------------------------   1   19
---------------------AAUGACCGCCUACCAAAGACA--------------------------------------------------------------   2   21
---------------------AAUGACCGCCUACCAAAGAC---------------------------------------------------------------   1   20
------------------------------CUACCAAAGACAAAUAGGAAU-----------------------------------------------------   1   21
--------------------------------ACCAAAGACAAAUAGGAAUCAAAU------------------------------------------------   1   24
--------------------------------ACCAAAGACAAAUAGGAAUCAAAUCCUAU-------------------------------------------   1   29
--------------------------------------------UAGGAAUCAAAUCCUAUUUGUCUUUGGUAGG-----------------------------   2   31
---------------------------------------------AGGAAUCAAAUCCUAUUUGUCUUUGGUAGG-----------------------------   2   30
----------------------------------------------GGAAUCAAAUCCUAUUUGUCUUUGGUAGG-----------------------------   1   29
-----------------------------------------------------AAUCCUAUUUGUCUUUGGUA-------------------------------   4   20
-----------------------------------------------------AAUCCUAUUUGUCUUUGG---------------------------------   1   18
------------------------------------------------------AUCCUAUUUGUCUUUGGUAG------------------------------   2   20
-------------------------------------------------------UCCUAUUUGUCUUUGGUAGG-----------------------------   9   20
----------------------------------------------------------UAUUUGUCUUUGGUAGGCGGU-------------------------   5   21
```

#### *D. firmibasis* dfi-mir-1189 continued

```
(((((((((((((((((((((((((((((((((((((((((((((((((......)))))))))))))))))))))))))))))))))))))))))))))))))    -108.21 kcal/mol
GGAAGUAAAGcAUGGAAGAUUAAUGACCGCcUACCAAAGACAAAUAGGAAUCAAAUCCUAUUUGUCUUUGGUAGGcGGUCAUUAAUCUUCCAUGCuUUACUUCC    count    length
------------------------------------------------------------UUUGUCUUUGGUAGGCGGUC------------------------    1        20
------------------------------------------------------------UUUGUCUUUGGUAGGCGGUCA-----------------------    1        21
------------------------------------------------------------UUUGUCUUUGGUAGGCGGUCAU----------------------    1        22
-------------------------------------------------------------UUGUCUUUGGUAGGCGGUCAU----------------------    2        21
--------------------------------------------------------------UGUCUUUGGUAGGCGGUCAU----------------------    1        20
---------------------------------------------------------------GUCUUUGGUAGGCGGUCAUU---------------------    2        20
----------------------------------------------------------------UCUUUGGUAGGCGGUCAUU---------------------    2        19
------------------------------------------------------------------UUUGGUAGGCGGUCAUUAAUCU----------------    3        22
------------------------------------------------------------------UUUGGUAGGCGGUCAUUAAUCUUC--------------    2        24
------------------------------------------------------------------UUUGGUAGGCGGUCAUUAAU------------------    2        20
--------------------------------------------------------------------UGGUAGGCGGUCAUUAAUCUU---------------    3        21
----------------------------------------------------------------------GUAGGCGGUCAUUAAUCUUCC-------------    1        21
-----------------------------------------------------------------------UAGGCGGUCAUUAAUCUU---------------    1        18
------------------------------------------------------------------------AGGCGGUCAUUAAUCUUCCAUGC---------    3        23
------------------------------------------------------------------------AGGCGGUCAUUAAUCUUCCAUGCUU-------    1        25
--------------------------------------------------------------------------GCGGUCAUUAAUCUUCCA------------    1        18
---------------------------------------------------------------------------CGGUCAUUAAUCUUCCAUGCUUU------    12       23
----------------------------------------------------------------------------GGUCAUUAAUCUUCCAUGCUUUA-----    1        23
----------------------------------------------------------------------------GGUCAUUAAUCUUCCAUGC---------    4        19
-----------------------------------------------------------------------------GUCAUUAAUCUUCCAUGC---------    3        18
------------------------------------------------------------------------------UCAUUAAUCUUCCAUGCUUU------    1        20
-------------------------------------------------------------------------------CAUUAAUCUUCCAUGCUUUAC----    1        21
-------------------------------------------------------------------------------CAUUAAUCUUCCAUGCUUUACU---    1        22
-------------------------------------------------------------------------------CAUUAAUCUUCCAUGCUU-------    1        18
----------------------------------------------------------------------------------UAAUCUUCCAUGCUUUACUUC-    1        21
```

```
                                                 AU
GGAAGUAAAGcAUGGAAGAUUAAUGACCGCcUACCAAAGACAAAUAGGA  C
CCUUCAUUuCGUACCUUCUAAUUACUGGcGGAUGGUUUCUGUUUAUCCU  A
                                                 AA
```

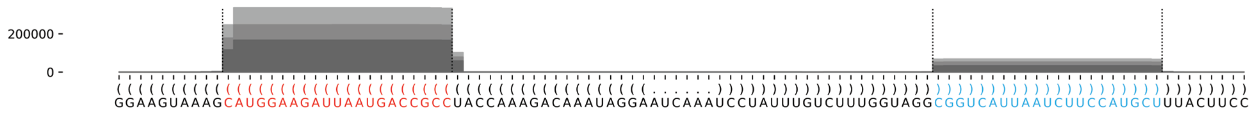

### *D. firmibasis* dfi-mir-1190

((((((((((((((((((((((((((((((((((((((((((((.(((((((((((((((...(((((..........(((((((((.......((((((((...))))))))(((((.(((((.....(((((.......)))))))))).)))))))))))))).......)))))..))))))))))))))).)))))))))))))))))))))))))))))))))))))))))))) −142.84 kcal/mol

CAACCAAAUCaAAAUUAUCAAAUGUCCUCAGcAAGAUAAUUCACAUAAUUCUUUAUUAAAUUCUUUUUAAAUGGGUUAACUCUGAUCUAAAAUAUUAACAAAAUUAUUACUAUAAUAAUUUAUAAAUUGACCAUAAUCAUUUCUUUUUAAAUGGGUUAUUUUAUAUUUUAGAUCCAAAUAAUAUUUUUAAAGAAUUUAAUAAACAAUUAUGUGAAUUAUCUUGCuGAGGACAUUUGAUAAUUUUGaUUUGGUUG

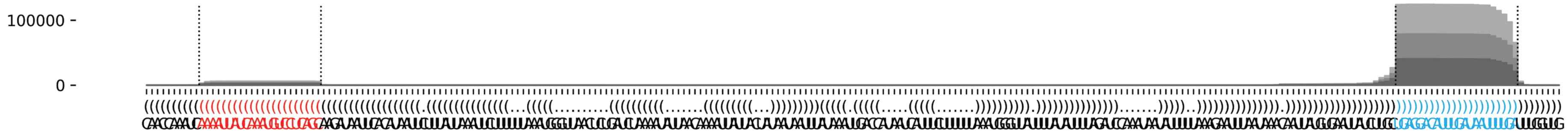

### *D. firmibasis* dfi-mir-1191

```
((((((((((((((((((.(((.((((((((((.(((((((((((((((((((.(..(((.(((((.(.(((((((.(((((((......)))))))......))))))).))))))...)))..)...))))).....)))))))))))))).)))))))))).))).))))))))))))))))))   -95.3 kcal/mol
UCAUUAUGCAaACAAGUGCAAUCAGGAAAUaAAUACUAUAACUGAGCAAGUAUCAUUUCACAUCAAACAAACAAGAAGAAUUAAAAAGAGUUAGUUUAUUGGCUCUUGUUGGUUGAUCAUUGAUUUUUAAUACUGCGGUUGUUCAGUUAUAGUUUUUAuUUCCUCGUUACACUUGUUUgCAUAAUGA   count   length
Libraries combined    Precision: [ total = 97.1% | 5p-arm = 44.6% | 3p-arm = 97.8% ]
--------------------------------------------------------------------------------------------------------------------------------------------------------------UUUCCUCGUUACACUUGUUUG--------   74616   21
--------------------------------------------------------------------------------------------------------------------------------------------------------------UUUCCUCGUUACACUUGUUUGC-------   11194   22
--------------------------------------------------------------------------------------------------------------------------------------------------------------UUUCCUCGUUACACUUGUUUGCA------   5823   23
----------AACAAGUGCAAUCAGGAAAUA------------------------------------------------------------------------------------------------------------------------------------------------------------   464   21
--------------------------------------------------------------------------------------------------------------------------------------------------------------UUUCCUCGUUACACUUGUUU---------   634   20
--------CAAACAAGUGCAAUCAGGAAAUA------------------------------------------------------------------------------------------------------------------------------------------------------------   328   23
--------------------------------------------------------------------------------------------------------------------------------------------------------------UUUCCUCGUUACACUUGUUUGCAU-----   526   24
------------------------------------------------------------------------------------------------------------------------------------------------------------UAUUUCCUCGUUACACUUGUU----------   334   21
---------AAACAAGUGCAAUCAGGAAAUA------------------------------------------------------------------------------------------------------------------------------------------------------------   228   22
------------------------------------------------------------------------------------------------------------------------------------------------------------UAUUUCCUCGUUACACUUGUUU---------   337   22
--------------------------------------------------------------------------------------------------------------------------------------------------------------UUUCCUCGUUACACUUGUU----------   281   19
----------AACAAGUGCAAUCAGGAAAU-------------------------------------------------------------------------------------------------------------------------------------------------------------   89   20
---------------------------------------------------------------------------------------------------------------------------------------------------------------UUCCUCGUUACACUUGUUUGCA------   156   22
----------------------------------------------------------------------------------------------------------------------------------------------------------------UCCUCGUUACACUUGUUUGCA------   119   21
--------CAAACAAGUGCAAUCAGGAAAU-------------------------------------------------------------------------------------------------------------------------------------------------------------   83   22
---------------------------------------------------------------------------------------------------------------------------------------------------------------UUCCUCGUUACACUUGUUUG--------   89   20
-----------------------------------------------------------------------------------------------------------------------------------------------------------UUAUUUCCUCGUUACACUUGU-----------   76   21
---------------------------------------------------------------------------------------------------------------------------------------------------------------UUCCUCGUUACACUUGUUUGC-------   81   21
---------------------------------------------------------------------------------------------------------------------------------------------------------UUUUAUUUCCUCGUUACACUUG------------   45   22
-------------------------------------------------------------------------------------------------------------------------------------------------------------AUUUCCUCGUUACACUUGUUU---------   51   21
---------AAACAAGUGCAAUCAGGAAAU-------------------------------------------------------------------------------------------------------------------------------------------------------------   30   21
----------------------------------------------------------------------------------------------------------------------------------------------------------------UCCUCGUUACACUUGUUUGC-------   37   20
---------------------------------------------------------------------------------------------------------------------------------------------------------UUUUAUUUCCUCGUUACACUU-------------   20   21
---------------------------------------------------------------------------------------------------------------------------------------------------------UUUUAUUUCCUCGUUACACU--------------   31   20
--------------------------------------------------------------------------------------------------------------------------------------------------------------UUUCCUCGUUACACUUGU-----------   30   18
------------------------------------------------------------------------------------------------------------------------------------------------------------UAUUUCCUCGUUACACUUGUUUG--------   7   23
----------------------------------------------------------------------------------------------------------------------------------------------------------------UCCUCGUUACACUUGUUUG--------   29   19
--------------------------------------------------------------------------------------------------------------------------------------------------------------UUUCCUCGUUACACUUGUUUGCAUA----   19   25
--------CAAACAAGUGCAAUCAGGAAA--------------------------------------------------------------------------------------------------------------------------------------------------------------   7   21
----------AACAAGUGCAAUCAGGAAA--------------------------------------------------------------------------------------------------------------------------------------------------------------   4   19
-----------------------------------------------------------------------------------------------------------------------------------------------------------UUAUUUCCUCGUUACACUUGUU----------   10   22
------------------------------------------------------------------------------------------------------------------------------------------------------------UAUUUCCUCGUUACACUUGU-----------   18   20
---------------------------------------------------------------------------------------------------------------------------------------------------------------UUCCUCGUUACACUUGUU----------   3   18
-----------ACAAGUGCAAUCAGGAAAUA------------------------------------------------------------------------------------------------------------------------------------------------------------   8   20
---------------------------------------------------------------------------------------------------------------------------------------------------------------UUCCUCGUUACACUUGUUUGCAU-----   11   23
-------GCAAACAAGUGCAAUCAGGAAAU-------------------------------------------------------------------------------------------------------------------------------------------------------------   2   23
----------AACAAGUGCAAUCAGGAAAUAA-----------------------------------------------------------------------------------------------------------------------------------------------------------   3   22
-----------ACAAGUGCAAUCAGGAAAU-------------------------------------------------------------------------------------------------------------------------------------------------------------   1   19
---------------------UCAGGAAAUAAAUACUAUAAC-------------------------------------------------------------------------------------------------------------------------------------------------   1   21
------------------------------------------------------------------------------------------------UAUUGGCUCUUGUUGGUUGAUCAUU------------------------------------------------------------------   1   25
-------------------------------------------------------------------------------------------------------------------------------------UGCGGUUGUUCAGUUAUAGUU---------------------------------   4   21
---------------------------------------------------------------------------------------------------------------------------------------CGGUUGUUCAGUUAUAGUUU--------------------------------   1   20
------------------------------------------------------------------------------------------------------------------------------------------------AGUUAUAGUUUUUAUUUCCUC----------------------   1   21
--------------------------------------------------------------------------------------------------------------------------------------------------UUAUAGUUUUUAUUUCCUCGUU-------------------   1   22
-------------------------------------------------------------------------------------------------------------------------------------------------------GUUUUUAUUUCCUCGUUACACU--------------   6   22
----------------------------------------------------------------------------------------------------------------------------------------------------------UUUAUUUCCUCGUUACACUUG------------   1   21
-----------------------------------------------------------------------------------------------------------------------------------------------------------UUAUUUCCUCGUUACACUUG------------   13   20
------------------------------------------------------------------------------------------------------------------------------------------------------------UAUUUCCUCGUUACACUUGUUUGC-------   1   24
------------------------------------------------------------------------------------------------------------------------------------------------------------UAUUUCCUCGUUACACUUG------------   3   19
----------------------------------------------------------------------------------------------------------------------------------------------------------------UCCUCGUUACACUUGUUU---------   3   18
-----------------------------------------------------------------------------------------------------------------------------------------------------------------CCUCGUUACACUUGUUUGCA------   5   20
--------CAAACAAGUGCAAUCAGGAAAUAA-----------------------------------------------------------------------------------------------------------------------------------------------------------   1   24
---------AAACAAGUGCAAUCAGGAAAUAA-----------------------------------------------------------------------------------------------------------------------------------------------------------   3   23
```

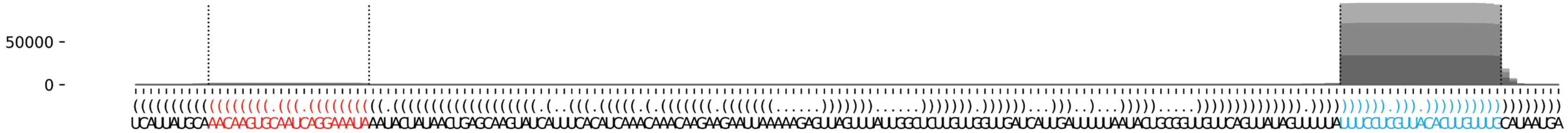

**D. firmibasis dfi-mir-1192**

```
((((((((.((.(((((((.((((((((((((((.((((...(((((((((........))))))))).)))).))))))))))))))).))))))).)).))))))))   -69.88 kcal/mol
UUUAUGCAGGuGUACCAAGACAAUCUCCAAgUACGUAUGGUAACUCAAUUGAAAAAUAAUGAUUGAGUACAUAAGUACUuGGAGAUUGGCUUGGUAAACaUGCAUAAA   count   length
Libraries combined          Precision: [ total = 84.5% | 5p-arm = 87.9% | 3p-arm = 76.9% ]
----------UGUACCAAGACAAUCUCCAAG-----------------------------------------------------------------------------   51      21
----UGCAGGUGUACCAAGACAAUC-----------------------------------------------------------------------------------   5       21
-------------------------------------------------------------------------------UGGAGAUUGGCUUGGUAAACA--------   9       21
--------------------------------------------------------------------------------GGAGAUUGGCUUGGUAAACAUG------   2       22
---------------------------------------------------------------------------UACUUGGAGAUUGGCUUGGUA------------   1       21
-------------------------------------------------------------------------------UGGAGAUUGGCUUGGUAAAC---------   10      20
--------------------------------------------------------------------------------GGAGAUUGGCUUGGUAAAC---------   1       19
------------UACCAAGACAAUCUCCAAGUA---------------------------------------------------------------------------   1       21
------------UACCAAGACAAUCUCCAAG-----------------------------------------------------------------------------   1       19
-----------------------------------------------------------------------------CUUGGAGAUUGGCUUGGUAAACA--------   2       23
-------------------------------------------------------------------------------UGGAGAUUGGCUUGGUAAA----------   1       19
Library = 1         Precision: [ total = 68.2% | 5p-arm = 80.0% | 3p-arm = 42.9% ]
----------UGUACCAAGACAAUCUCCAAG-----------------------------------------------------------------------------   12      21
----UGCAGGUGUACCAAGACAAUC-----------------------------------------------------------------------------------   3       21
-------------------------------------------------------------------------------UGGAGAUUGGCUUGGUAAACA--------   2       21
--------------------------------------------------------------------------------GGAGAUUGGCUUGGUAAACAUG------   2       22
---------------------------------------------------------------------------UACUUGGAGAUUGGCUUGGUA------------   1       21
-------------------------------------------------------------------------------UGGAGAUUGGCUUGGUAAAC---------   1       20
--------------------------------------------------------------------------------GGAGAUUGGCUUGGUAAAC---------   1       19
Library = 2         Precision: [ total = 92.0% | 5p-arm = 94.3% | 3p-arm = 86.7% ]
----------UGUACCAAGACAAUCUCCAAG-----------------------------------------------------------------------------   33      21
-------------------------------------------------------------------------------UGGAGAUUGGCUUGGUAAACA--------   7       21
-------------------------------------------------------------------------------UGGAGAUUGGCUUGGUAAAC---------   5       20
-----------------------------------------------------------------------------CUUGGAGAUUGGCUUGGUAAACA--------   2       23
----UGCAGGUGUACCAAGACAAUC-----------------------------------------------------------------------------------   1       21
------------UACCAAGACAAUCUCCAAGUA---------------------------------------------------------------------------   1       21
-------------------------------------------------------------------------------UGGAGAUUGGCUUGGUAAA----------   1       19
Library = 3         Precision: [ total = 83.3% | 5p-arm = 75.0% | 3p-arm = 100.0% ]
----------UGUACCAAGACAAUCUCCAAG-----------------------------------------------------------------------------   6       21
-------------------------------------------------------------------------------UGGAGAUUGGCUUGGUAAAC---------   4       20
----UGCAGGUGUACCAAGACAAUC-----------------------------------------------------------------------------------   1       21
------------UACCAAGACAAUCUCCAAG-----------------------------------------------------------------------------   1       19
```

```
        G  G       A              G    GUA         AAA
UUUAUGCA Gu UACCAAG CAAUCUCCAAgUAC UAUG   ACUCAAUUG   A
AAAUACGU CA AUGGUUC GUUAGAGGuUCAUG AUAC --UGAGUUAGU   A
        a  A       G              A    A           AAU
```

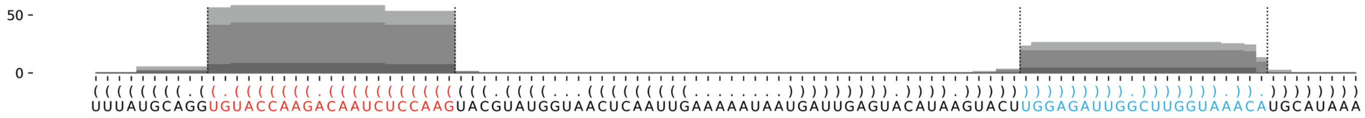

### *D. firmibasis* dfi-mir-1193

```
...(((.(((((((((.((((((((.((.(((((.((((.((((((((........)))))....))))))).))))).)).)))))))).))))))))).)))...   -42.09 kcal/mol
UUUUCUAUUUuUACUCCCAUUAUUAAGGAUgCAAAUUAAAUUUGUAUGCAAAAACACAUAUUUUUAAAUUAAAUUGCAcCCAUAAUAAUGAGAGUAAAAaAAAGAUAA   count    length
Libraries combined          Precision: [ total = 99.6% | 5p-arm = 99.6% | 3p-arm = 100.0% ]
----------UUACUCCCAUUAUUAAGGAUG-----------------------------------------------------------------------------   63315    21
---------UUUACUCCCAUUAUUAAGGAUG-----------------------------------------------------------------------------   137      22
----------UUACUCCCAUUAUUAAGGAU------------------------------------------------------------------------------   86       20
-----------UACUCCCAUUAUUAAGGAUG-----------------------------------------------------------------------------   45       20
------------------------------------------------------------------------------CCCAUAAUAAUGAGAGUAAAA---------   25       21
----------UUACUCCCAUUAUUAAGGA-------------------------------------------------------------------------------   14       19
-------------CUCCCAUUAUUAAGGAUG-----------------------------------------------------------------------------   28       18
------------ACUCCCAUUAUUAAGGAUG-----------------------------------------------------------------------------   17       19
------AUUUUUACUCCCAUUAUUAAGGAUG-----------------------------------------------------------------------------   2        25
------------------------------------------------------------------------------CCCAUAAUAAUGAGAGUAAA----------   6        20
--------UUUUACUCCCAUUAUUAAGGAUG-----------------------------------------------------------------------------   1        23
------------------------------------------------------------------------------CCCAUAAUAAUGAGAGUAAAAA--------   2        22
-------UUUUUACUCCCAUUAUUAAGGAUG-----------------------------------------------------------------------------   1        24
----------UUACUCCCAUUAUUAAGG--------------------------------------------------------------------------------   1        18
Library = 1        Precision: [ total = 99.7% | 5p-arm = 99.7% | 3p-arm = 100.0% ]
----------UUACUCCCAUUAUUAAGGAUG-----------------------------------------------------------------------------   25559    21
---------UUUACUCCCAUUAUUAAGGAUG-----------------------------------------------------------------------------   54       22
----------UUACUCCCAUUAUUAAGGAU------------------------------------------------------------------------------   32       20
-----------UACUCCCAUUAUUAAGGAUG-----------------------------------------------------------------------------   15       20
------------------------------------------------------------------------------CCCAUAAUAAUGAGAGUAAAA---------   8        21
----------UUACUCCCAUUAUUAAGGA-------------------------------------------------------------------------------   7        19
-------------CUCCCAUUAUUAAGGAUG-----------------------------------------------------------------------------   7        18
------------ACUCCCAUUAUUAAGGAUG-----------------------------------------------------------------------------   5        19
------AUUUUUACUCCCAUUAUUAAGGAUG-----------------------------------------------------------------------------   2        25
------------------------------------------------------------------------------CCCAUAAUAAUGAGAGUAAA----------   2        20
--------UUUUACUCCCAUUAUUAAGGAUG-----------------------------------------------------------------------------   1        23
------------------------------------------------------------------------------CCCAUAAUAAUGAGAGUAAAAA--------   1        22
Library = 2        Precision: [ total = 99.6% | 5p-arm = 99.6% | 3p-arm = 100.0% ]
----------UUACUCCCAUUAUUAAGGAUG-----------------------------------------------------------------------------   17531    21
---------UUUACUCCCAUUAUUAAGGAUG-----------------------------------------------------------------------------   36       22
----------UUACUCCCAUUAUUAAGGAU------------------------------------------------------------------------------   22       20
-------------CUCCCAUUAUUAAGGAUG-----------------------------------------------------------------------------   12       18
-----------UACUCCCAUUAUUAAGGAUG-----------------------------------------------------------------------------   11       20
------------ACUCCCAUUAUUAAGGAUG-----------------------------------------------------------------------------   6        19
------------------------------------------------------------------------------CCCAUAAUAAUGAGAGUAAAA---------   6        21
-------UUUUUACUCCCAUUAUUAAGGAUG-----------------------------------------------------------------------------   1        24
----------UUACUCCCAUUAUUAAGGA-------------------------------------------------------------------------------   1        19
------------------------------------------------------------------------------CCCAUAAUAAUGAGAGUAAA----------   1        20
Library = 3        Precision: [ total = 99.6% | 5p-arm = 99.6% | 3p-arm = 100.0% ]
----------UUACUCCCAUUAUUAAGGAUG-----------------------------------------------------------------------------   20225    21
---------UUUACUCCCAUUAUUAAGGAUG-----------------------------------------------------------------------------   47       22
----------UUACUCCCAUUAUUAAGGAU------------------------------------------------------------------------------   32       20
-----------UACUCCCAUUAUUAAGGAUG-----------------------------------------------------------------------------   19       20
------------------------------------------------------------------------------CCCAUAAUAAUGAGAGUAAAA---------   11       21
-------------CUCCCAUUAUUAAGGAUG-----------------------------------------------------------------------------   9        18
----------UUACUCCCAUUAUUAAGGA-------------------------------------------------------------------------------   6        19
------------ACUCCCAUUAUUAAGGAUG-----------------------------------------------------------------------------   6        19
------------------------------------------------------------------------------CCCAUAAUAAUGAGAGUAAA----------   3        20
----------UUACUCCCAUUAUUAAGG--------------------------------------------------------------------------------   1        18
------------------------------------------------------------------------------CCCAUAAUAAUGAGAGUAAAAA--------   1        22
```

```
UUU    A          C         A  A      A    A             CAA
   UCU UUUuUACUC CAUUAUUA GG UgCAA UUAA UUU----GUAUG    A
   AGA AaAAAUGAG GUAAUAAU CC ACGUU AAUU-AAA    UAUAC    A
AAU    A          A         A  c      A         UUUU      ACA
```

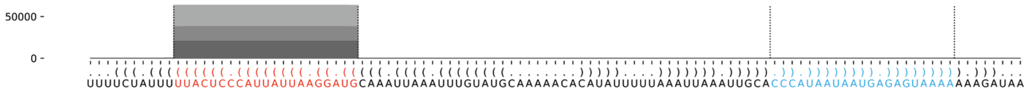

### *D. firmibasis* dfi-mir-1194

```
((((((((((((((((((((((((((((.(((((((((((((.((.(((....))).)).))))))))))))).)))))))))))))))))))))))))))) -101.3 kcal/mol
UUAUUAAUGCuCGGAGCUCUGAUGCGAACUgUGGGUGGACGCUGGAAAUGUUGAUUGCCUGCGUCCACCCACAuUUCGCAUCAGAGCUCCGAGcAUUAAUAA count    length
Libraries combined          Precision: [ total = 98.8% | 5p-arm = 77.1% | 3p-arm = 98.8% ]
-------------------------------------------------------------------------UUUCGCAUCAGAGCUCCGAGC-------- 407849   21
--------------------------------------------------------------------------UUCGCAUCAGAGCUCCGAGC-------- 3289     20
-------------------------------------------------------------------------UUUCGCAUCAGAGCUCCGAG--------- 1908     20
-------------------------------------------------------------------------UUUCGCAUCAGAGCUCCGAGCA------- 1359     22
-------------------------------------------------------------------------UUUCGCAUCAGAGCUCCGAGCAU------ 557      23
----------------------------------------------------------------------------CGCAUCAGAGCUCCGAGCAUU----- 510      21
---------------------------------------------------------------------------UCGCAUCAGAGCUCCGAGC-------- 284      19
-------------------------------------------------------------------------UUUCGCAUCAGAGCUCCGAGCAUU----- 216      24
----------UCGGAGCUCUGAUGCGAACU------------------------------------------------------------------------ 203      20
---------------------------------------------------------------------------UCGCAUCAGAGCUCCGAGCAU------ 242      21
----------UCGGAGCUCUGAUGCGAACUG----------------------------------------------------------------------- 173      21
-------------------------------------------------------------------------UUUCGCAUCAGAGCUCCGA---------- 60       19
----------UCGGAGCUCUGAUGCGAAC------------------------------------------------------------------------- 51       19
-----------------------------------------------------------------------CAUUUCGCAUCAGAGCUCCGA---------- 41       21
---UUAAUGCUCGGAGCUCUGAUG------------------------------------------------------------------------------ 50       21
--------------------------------------------------------------------------UUCGCAUCAGAGCUCCGAGCA------- 43       21
---------------------------------------------------------------------------UCGCAUCAGAGCUCCGAGCAUU----- 39       22
----------------------------------------------------------------------------CGCAUCAGAGCUCCGAGCAU------ 45       20
--------------------------------------------------------------------------------UCAGAGCUCCGAGCAUUAAUA- 34       21
-------------------------------------------------------------------------UUUCGCAUCAGAGCUCCGAGCAUUAA--- 40       26
-------------------------------------------------------------------------UUUCGCAUCAGAGCUCCGAGCAUUA---- 15       25
-------------------------------------------------------------------------UUUCGCAUCAGAGCUCCGAGCAUUAAU-- 25       27
-----------------------------------------------------------------------------GCAUCAGAGCUCCGAGCAUUA---- 15       21
-------------------------------------------------------------------------------AUCAGAGCUCCGAGCAUUAAU-- 20       21
------------------------------------------------------------------------AUUUCGCAUCAGAGCUCCGAGC-------- 22       22
------------------------------------------------------------------------------CAUCAGAGCUCCGAGCAUUAA--- 6        21
--------------------------------------------------------------------------------UCAGAGCUCCGAGCAUUAAU-- 9        20
-----AAUGCUCGGAGCUCUGAUGCG---------------------------------------------------------------------------- 18       21
----------------------------------------------------------------------------CGCAUCAGAGCUCCGAGCAUUA---- 6        22
------AUGCUCGGAGCUCUGAUGCGA--------------------------------------------------------------------------- 3        21
----------UCGGAGCUCUGAUGCGAACUGU---------------------------------------------------------------------- 2        22
---------------------------------------------------------------------CACAUUUCGCAUCAGAGCUCC------------ 4        21
-----------------------------------------------------------------------CAUUUCGCAUCAGAGCUCCG----------- 8        20
--------------------------------------------------------------------------UUCGCAUCAGAGCUCCGAG--------- 4        19
---------------------------------------------------------------------------UCGCAUCAGAGCUCCGAGCAUUA---- 7        23
-----------------------------------------------------------------------------GCAUCAGAGCUCCGAGCAUUAA--- 2        22
------------------------------------------------------------------------------CAUCAGAGCUCCGAGCAUUAAUAA 2        24
------------------------------------------------------------------------------CAUCAGAGCUCCGAGCAUU----- 4        19
---UUAAUGCUCGGAGCUCUGAU------------------------------------------------------------------------------- 3        20
----UAAUGCUCGGAGCUCUGAUGC----------------------------------------------------------------------------- 5        21
-------UGCUCGGAGCUCUGAUGCGAA-------------------------------------------------------------------------- 4        21
-------UGCUCGGAGCUCUGAUGCGAACU------------------------------------------------------------------------ 12       23
--------GCUCGGAGCUCUGAUGCGAAC------------------------------------------------------------------------- 1        21
--------GCUCGGAGCUCUGAUGCGA--------------------------------------------------------------------------- 1        19
---------CUCGGAGCUCUGAUGCGAAC------------------------------------------------------------------------- 1        20
------------GGAGCUCUGAUGCGAACUGUG--------------------------------------------------------------------- 1        21
-------------------UGAUGCGAACUGUGGGUGGACG------------------------------------------------------------- 1        22
---------------------------------------------------UGAUUGCCUGCGUCCACCCA------------------------------- 1        20
-------------------------------------------------------------CGUCCACCCACAUUUCGCAUC-------------------- 1        21
---------------------------------------------------------------------CACAUUUCGCAUCAGAGCUCCG----------- 1        22
----------------------------------------------------------------------ACAUUUCGCAUCAGAGCUCCG----------- 4        21
------------------------------------------------------------------------AUUUCGCAUCAGAGCUCCGAG--------- 2        21
------------------------------------------------------------------------AUUUCGCAUCAGAGCUCCGA---------- 1        20
```

### *D. firmibasis* dfi-mir-1194

```
((((((((((((((((((((((((((((.(((((((((((((.((.(((....))).)).))))))))))))).)))))))))))))))))))))))))))) -101.3 kcal/mol
UUAUUAAUGCuCGGAGCUCUGAUGCGAACUgUGGGUGGACGCUGGAAAUGUUGAUUGCCUGCGUCCACCCACAuUUCGCAUCAGAGCUCCGAGcAUUAAUAA count    length
Libraries combined          Precision: [ total = 98.8% | 5p-arm = 77.1% | 3p-arm = 98.8% ]
--------------------------------------------------------------------------UUCGCAUCAGAGCUCCGAGCAU------ 4        22
---------------------------------------------------------------------------UCGCAUCAGAGCUCCGAGCAUUAA--- 3        24
---------------------------------------------------------------------------UCGCAUCAGAGCUCCGAGCAUUAAU-- 1        25
---------------------------------------------------------------------------UCGCAUCAGAGCUCCGAG--------- 1        18
---------------------------------------------------------------------------UCGCAUCAGAGCUCCGAGCA------- 4        20
----------------------------------------------------------------------------CGCAUCAGAGCUCCGAGCAUUAA--- 2        23
-------------------------------------------------------------------------------AUCAGAGCUCCGAGCAUUAA--- 1        20
--------------------------------------------------------------------------------UCAGAGCUCCGAGCAUUAA--- 1        19
-UAUUAAUGCUCGGAGCUCUGAU------------------------------------------------------------------------------- 1        22
-UAUUAAUGCUCGGAGCUCUGA-------------------------------------------------------------------------------- 1        21
-UAUUAAUGCUCGGAGCUCUG--------------------------------------------------------------------------------- 1        20
--AUUAAUGCUCGGAGCUCUGAU------------------------------------------------------------------------------- 2        21
----UAAUGCUCGGAGCUCUGA-------------------------------------------------------------------------------- 2        18
-----AAUGCUCGGAGCUCUGAUGCGAACU------------------------------------------------------------------------ 1        25
-----AAUGCUCGGAGCUCUGAUGCGA--------------------------------------------------------------------------- 1        22
------AUGCUCGGAGCUCUGAUG------------------------------------------------------------------------------ 1        18
-------UGCUCGGAGCUCUGAUGCGA--------------------------------------------------------------------------- 1        20
--------GCUCGGAGCUCUGAUGCGAACU------------------------------------------------------------------------ 2        22
----------UCGGAGCUCUGAUGCGAACUGUG--------------------------------------------------------------------- 1        23
------------GGAGCUCUGAUGCGAACUG----------------------------------------------------------------------- 1        19
--------------AGCUCUGAUGCGAACUGUGGG------------------------------------------------------------------- 6        21
----------------CUCUGAUGCGAACUGUGGGUG----------------------------------------------------------------- 4        21
-----------------UCUGAUGCGAACUGUGGGUGG---------------------------------------------------------------- 2        21
----------------------UGCGAACUGUGGGUGGACGCU----------------------------------------------------------- 2        21
------------------------------------------------------------------ACCCACAUUUCGCAUCAGAGCU-------------- 1        22
-------------------------------------------------------------------------UUUCGCAUCAGAGCUCCG----------- 6        18
-------------------------------------------------------------------------UUUCGCAUCAGAGCUCCGAGCAUUAAUAA 2        29
--------------------------------------------------------------------------UUCGCAUCAGAGCUCCGAGCAUU----- 1        23
---------------------------------------------------------------------------UCGCAUCAGAGCUCCGAGCAUUAAUA- 1        26
----------------------------------------------------------------------------CGCAUCAGAGCUCCGAGC-------- 8        18
----------------------------------------------------------------------------CGCAUCAGAGCUCCGAGCA------- 2        19
-----------------------------------------------------------------------------GCAUCAGAGCUCCGAGCAUU----- 3        20
-----------------------------------------------------------------------------GCAUCAGAGCUCCGAGCAUUAAU-- 1        23
------------------------------------------------------------------------------CAUCAGAGCUCCGAGCAUUA---- 1        20
------------------------------------------------------------------------------CAUCAGAGCUCCGAGCAUUAAU-- 3        22
------------------------------------------------------------------------------CAUCAGAGCUCCGAGCAUUAAUA- 1        23
--------------------------------------------------------------------------------UCAGAGCUCCGAGCAUUAAUAA 5        22
---------------------------------------------------------------------------------CAGAGCUCCGAGCAUUAAU-- 1        19
```

```
                            C             U  A    G
UUAUUAAUGCuCGGAGCUCUGAUGCGAA UgUGGGUGGACGC GG AAU U
AAUAAUUAcGAGCCUCGAGACUACGCUU ACACCCACCUGCG CC UUA U
                            u             U  G    G
```

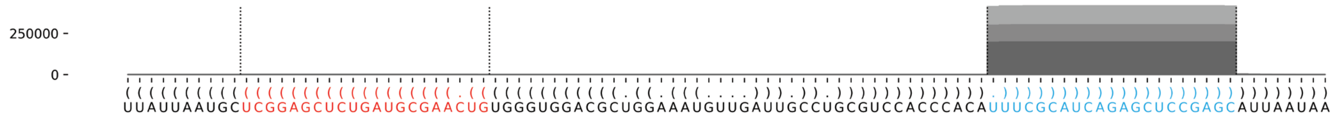

### *D. firmibasis* dfi-mir-1195

```
(((.(((.(((((((((.(((((((((((((((((((..(((.(((((..((((((((....))))))))...........))))).)))..))))))))))))))))))).))))))))).))).)))     -69.84 kcal/mol
AGGUGAUCGUuGAUGUCGCCAAUGUAAUAAcAGAGGUUAAUUAUAUUAAAUAAUAUUUCAAUAAAUAUUAACAUUAACUUCUAAUACAAUAUACCUCUGUuAUUACAUUGGGGAUAUCAAcCGUCCCCU     count    length
Libraries combined          Precision: [ total = 97.7% | 5p-arm = 97.8% | 3p-arm = 34.5% ]
----------UGAUGUCGCCAAUGUAAUAAC--------------------------------------------------------------------------------------------------     530751   21
----------UGAUGUCGCCAAUGUAAUAA---------------------------------------------------------------------------------------------------     6469     20
---UGAUCGUUGAUGUCGCCAAUG---------------------------------------------------------------------------------------------------------     6127     21
----------UGAUGUCGCCAAUGUAAUAACA-------------------------------------------------------------------------------------------------     3143     22
---------UUGAUGUCGCCAAUGUAAUAAC--------------------------------------------------------------------------------------------------     1647     22
-----------GAUGUCGCCAAUGUAAUAAC--------------------------------------------------------------------------------------------------     1735     20
---------UUGAUGUCGCCAAUGUAAUAA---------------------------------------------------------------------------------------------------     701      21
------------AUGUCGCCAAUGUAAUAACAG------------------------------------------------------------------------------------------------     587      21
---------UUGAUGUCGCCAAUGUAAU-----------------------------------------------------------------------------------------------------     477      19
----------UGAUGUCGCCAAUGUAAU-----------------------------------------------------------------------------------------------------     483      18
-----------------------------------------------------------------------------------------------------AUUACAUUGGGGAUAUCAACC-------     398      21
------------AUGUCGCCAAUGUAAUAAC--------------------------------------------------------------------------------------------------     340      19
----------UGAUGUCGCCAAUGUAAUA----------------------------------------------------------------------------------------------------     305      19
----------------------------------------------------------------------------------------------------UAUUACAUUGGGGAUAUCAAC--------     188      21
---------UUGAUGUCGCCAAUGUAAUA----------------------------------------------------------------------------------------------------     165      20
-----------GAUGUCGCCAAUGUAAUAACA-------------------------------------------------------------------------------------------------     170      21
----------------------------------------------------------------------------------------------------UAUUACAUUGGGGAUAUCAA---------     77       20
------UCGUUGAUGUCGCCAAUGUA-------------------------------------------------------------------------------------------------------     68       20
-----------------------------------------------------------------------------------------------------AUUACAUUGGGGAUAUCAAC--------     55       20
-----------GAUGUCGCCAAUGUAAUAA---------------------------------------------------------------------------------------------------     59       19
-----AUCGUUGAUGUCGCCAAUGUA-------------------------------------------------------------------------------------------------------     39       21
----------UGAUGUCGCCAAUGUAAUAACAG------------------------------------------------------------------------------------------------     40       23
---UGAUCGUUGAUGUCGCCAA-----------------------------------------------------------------------------------------------------------     31       19
--------------GUCGCCAAUGUAAUAACAGAG----------------------------------------------------------------------------------------------     16       21
------------AUGUCGCCAAUGUAAUAACAGA-----------------------------------------------------------------------------------------------     27       22
-----------------------------------------------------------------------------------------------UCUGUUAUUACAUUGGGGAU--------------     12       20
---UGAUCGUUGAUGUCGCCAAU----------------------------------------------------------------------------------------------------------     20       20
-------------UGUCGCCAAUGUAAUAAC--------------------------------------------------------------------------------------------------     9        18
-----------------------------------------------------------------------------------------------------AUUACAUUGGGGAUAUCAA---------     18       19
---UGAUCGUUGAUGUCGCCA------------------------------------------------------------------------------------------------------------     9        18
----------------------------------------------------------------------------------------------------UAUUACAUUGGGGAUAUC-----------     9        18
-----AUCGUUGAUGUCGCCAAUG---------------------------------------------------------------------------------------------------------     12       19
-------------UGUCGCCAAUGUAAUAACAGAG----------------------------------------------------------------------------------------------     5        22
-----------GAUGUCGCCAAUGUAAUAACAG------------------------------------------------------------------------------------------------     8        22
--------------GUCGCCAAUGUAAUAACAG------------------------------------------------------------------------------------------------     5        19
--------------------------------------------------------------------------------------------------GUUAUUACAUUGGGGAUAUC-----------     4        20
---------------------------------------------------------------------------------------------------UUAUUACAUUGGGGAUAUCAA---------     4        21
-----------------------------------------------------------------------------------------------------AUUACAUUGGGGAUAUCAACCG------     7        22
---UGAUCGUUGAUGUCGCCAAUGU--------------------------------------------------------------------------------------------------------     11       22
------UCGUUGAUGUCGCCAAUG---------------------------------------------------------------------------------------------------------     2        18
----------UGAUGUCGCCAAUGUAAUAACAGA-----------------------------------------------------------------------------------------------     4        24
-----------GAUGUCGCCAAUGUAAUA----------------------------------------------------------------------------------------------------     6        18
------------AUGUCGCCAAUGUAAUAACA-------------------------------------------------------------------------------------------------     3        20
---------------UCGCCAAUGUAAUAACAGAGG---------------------------------------------------------------------------------------------     8        21
-------------------------------------------------------------------------------------------------UGUUAUUACAUUGGGGAUAUC-----------     8        21
----------------------------------------------------------------------------------------------------UAUUACAUUGGGGAUAUCAACC-------     5        22
------------------------------------------------------------------------------------------------------UUACAUUGGGGAUAUCAA---------     2        18
------------------------------------------------------------------------------------------------------UUACAUUGGGGAUAUCAAC--------     3        19
------------------------------------------------------------------------------------------------------UUACAUUGGGGAUAUCAACCG------     3        21
---UGAUCGUUGAUGUCGCCAAUGUA-------------------------------------------------------------------------------------------------------     3        23
----GAUCGUUGAUGUCGCCAAUG---------------------------------------------------------------------------------------------------------     7        20
------UCGUUGAUGUCGCCAAUGUAAU-----------------------------------------------------------------------------------------------------     1        22
-----------------------------------------------------------------------------------------------UCUGUUAUUACAUUGGGGAUA-------------     5        21
```

```
   U   C         G                   UA   A     AA                 C
AGG GAU GUuGAUGUC CCAAUGUAAUAAcAGAGGU  AUU UAUUA  ---------UAAUAUUU A
UCC CUG cAACUAUAG GGUUACAUUAuUGUCUCCA  UAA AUAAU           AUUAUAAA A
   C   C         G                   UA   C     CUUCAAUUACA        U
```

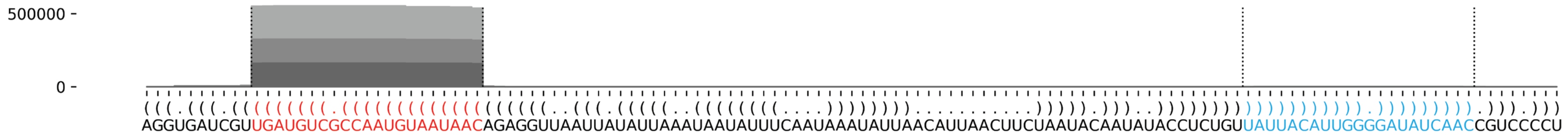

### *D. firmibasis* dfi-mir-1196

```
(((((((((((((((((((((.((((((((((((((((((((((((...((((........)))).))))).....))))))))))))))))))).)))))))))))))))))))))   -75.37 kcal/mol
AAAAAAAACAuAGUGAAGAUGGGUAGAAGAuAAAAGCUGGUUUUGAAAAUUCAGAGAAUGGUGGGGUUAGAAAAUUACCAGCUUUUAUcUUCUACACAUCUUCACUAUgUUUUUUUU   count    length
Libraries combined         Precision: [ total = 76.8% | 5p-arm = 73.1% | 3p-arm = 79.1% ]
----------UAGUGAAGAUGGGUAGAAGAU--------------------------------------------------------------------------------------   18       21
----------------------------------------------------------------------------------------CUUCUACACAUCUUCACUAUG--------   34       21
--------------------------------------------------------------------------------------AUCUUCUACACAUCUUCACUA----------   2        21
-----------------------------------------------------------------------------------------UUCUACACAUCUUCACUAUGU-------   3        21
----------UAGUGAAGAUGGGUAGAAGAUA-------------------------------------------------------------------------------------   1        22
-----------------GAUGGGUAGAAGAUAAAAGCU-------------------------------------------------------------------------------   3        21
------------------AUGGGUAGAAGAUAAAAGCU-------------------------------------------------------------------------------   1        20
-----------AGUGAAGAUGGGUAGAAGAU--------------------------------------------------------------------------------------   1        20
----------------AGAUGGGUAGAAGAUAAAAGC--------------------------------------------------------------------------------   2        21
------------------------------------------------------------------------AAUUACCAGCUUUUAUCUUCUA-----------------------   1        22
-------------------------------------------------------------------------------------UAUCUUCUACACAUCUUCACU-----------   2        21
---------------------------------------------------------------------------------------UCUUCUACACAUCUUCACUAUG--------   1        22
Library = 1        Precision: [ total = 64.7% | 5p-arm = 75.0% | 3p-arm = 55.6% ]
----------UAGUGAAGAUGGGUAGAAGAU--------------------------------------------------------------------------------------   5        21
----------------------------------------------------------------------------------------CUUCUACACAUCUUCACUAUG--------   5        21
--------------------------------------------------------------------------------------AUCUUCUACACAUCUUCACUA----------   2        21
-----------------------------------------------------------------------------------------UUCUACACAUCUUCACUAUGU-------   2        21
----------UAGUGAAGAUGGGUAGAAGAUA-------------------------------------------------------------------------------------   1        22
-----------------GAUGGGUAGAAGAUAAAAGCU-------------------------------------------------------------------------------   1        21
------------------AUGGGUAGAAGAUAAAAGCU-------------------------------------------------------------------------------   1        20
Library = 2        Precision: [ total = 87.5% | 5p-arm = 71.4% | 3p-arm = 100.0% ]
----------------------------------------------------------------------------------------CUUCUACACAUCUUCACUAUG--------   9        21
----------UAGUGAAGAUGGGUAGAAGAU--------------------------------------------------------------------------------------   5        21
----------------AGAUGGGUAGAAGAUAAAAGC--------------------------------------------------------------------------------   1        21
-----------------GAUGGGUAGAAGAUAAAAGCU-------------------------------------------------------------------------------   1        21
Library = 3        Precision: [ total = 77.8% | 5p-arm = 72.7% | 3p-arm = 80.0% ]
----------------------------------------------------------------------------------------CUUCUACACAUCUUCACUAUG--------   20       21
----------UAGUGAAGAUGGGUAGAAGAU--------------------------------------------------------------------------------------   8        21
-------------------------------------------------------------------------------------UAUCUUCUACACAUCUUCACU-----------   2        21
-----------AGUGAAGAUGGGUAGAAGAU--------------------------------------------------------------------------------------   1        20
----------------AGAUGGGUAGAAGAUAAAAGC--------------------------------------------------------------------------------   1        21
-----------------GAUGGGUAGAAGAUAAAAGCU-------------------------------------------------------------------------------   1        21
------------------------------------------------------------------------AAUUACCAGCUUUUAUCUUCUA-----------------------   1        22
---------------------------------------------------------------------------------------UCUUCUACACAUCUUCACUAUG--------   1        22
-----------------------------------------------------------------------------------------UUCUACACAUCUUCACUAUGU-------   1        21
```

```
                     G                             AAA    GAG
AAAAAAAACAuAGUGAAGAUG GUAGAAGAuAAAAGCUGGU-----UUUGA   UUCA   A
UUUUUUUUgUAUCACUUCUAC CAUCUUcUAUUUUCGACCA     AGAUU --GGGU   A
                     A                   UUAAA     G      GGU
```

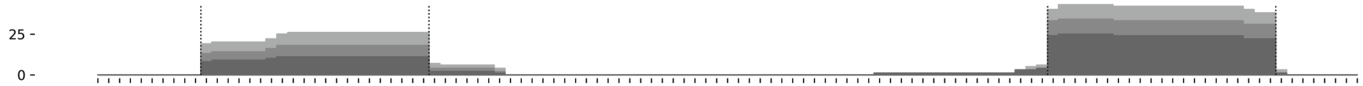


```
(((((((((((((((((((((.((((((((((((((((((((((((...((((........)))).))))).....))))))))))))))))))).)))))))))))))))))))))
AAAAAAAACAUAGUGAAGAUGGGUAGAAGAUAAAAGCUGGUUUUGAAAAUUCAGAGAAUGGUGGGGUUAGAAAAUUACCAGCUUUUAUCUUCUACACAUCUUCACUAUGUUUUUUUU
```

**Supplementary Data 3: Detailed depiction of all identified miRNAs in D. firmibasis following de-novo sequencing of the genome.** Predicted miRNA hairpin sequences and structure displayed in bracket notation. Identified miRNA-5p is indicated in red, and the miRNA-3p in cyan. Lowercase nucleotides represent the 5’ and 3’ ends of the miRNA-5p and miRNA-3p. Below the sequence, all mapped small RNA reads are aligned to the miRNA hairpin and the number reads and their length are shown to the right. Results are shown for all small RNA sequencing libraries combined (denoted Libraries combined). For miRNAs that have fewer than 50,000 reads mapped, the results are shown for the individual libraries as well (denoted Library = 1 to 3). The percentage of reads that map to the exact 5’ nucleotide of either the miRNA-5p or miRNA-3p is shown as ‘precision’ and calculated on the entire miRNA-hairpin (denoted total), and on the 5p- and 3p-arms of the miRNA hairpin (denoted 5p-arm and 3p-arm respectively). The folded miRNA hairpin structure, shown below the mapped small RNA, was predicted using the ViennaRNA RNAlib-2.6.2 python package, with 22°C folding temperature. At the bottom, a graph shows the read mapping density on the miRNA hairpin, with the miRNA-5p and miRNA-3p indicated in red and cyan respectively. The different shades of grey in the graph represent the different small RNA sequencing libraries.
