## Supplementary Table 2 for "Evolution of microRNAs in Amoebozoa and implications for the origin of multicellularity"

**Supplementary Table 2 | Characteristics of the dictyostelid miRNAs.** For each miRNA, the mature miRNA sequence is displayed (the most abundant strand in the duplex). miRNA_counts is the sum of miRNA-5p and miRNA-3p reads, and miRNA_RPM are the normalized reads on the miRNA-hairpin per million mapped total. The precision is the percentage of reads from the miRNA-hairpin that map with their 5' end at the exact 5' end of either the miRNA-5p or miRNA-3p. The genomic_context, genomic_location and strand show where the miRNA is located relative to the annotated ORFs, its genomic location, and whether it is from the Watson strand (+) or the Crick strand (-), respectively. For those miRNAs where we could detect homology, the miRNA_family is indicated, and for identified paralogs, a suffix is included in the miRNA.ID. Folding energy is the computed Gibbs free energy in kcal/mol from folding the miRNA-hairpin in silico at 22°C. The pre-miRNA size (nt) is the length (in nt) of the pre-miRNA, i.e. from the miRNA-5p 5' end to the miRNA-3p 3' end.

| miRNA.ID | miRNA_sequence | miRNA_counts | miRNA_RPM | precision | genomic_context | genomic_location | strand | miRNA_family | folding energy (kcal/mol) | pre-miRNA size (nt) |
|---|---|---|---|---|---|---|---|---|---|---|
| ddi-mir-1179 | UGGGUCUCCAUUCAAAUUAAGUUC | 83 | 13,70 | 98,8 | intergenic | NC_007088.5:1649212..1649317 | - | None | -56,04 | 88 |
| ddi-mir-7097 | UUUUGGCAGAAGUAGAGACGA | 600 | 109,00 | 91,2 | intergenic | NC_007089.4:3739201..3739306 | - | None | -61,87 | 88 |
| ddi-mir-1183 | UAAGUGGUCGUUAAUUGCUGGC | 1543 | 216,00 | 88,1 | intergenic | NC_007090.3:735807..735953 | - | None | -88,47 | 129 |
| ddi-mir-7096 | AAUCUUAAUUGUUCACAGACC | 411 | 53,40 | 99,5 | intergenic | NC_007090.3:1476903..1477007 | - | None | -72,22 | 87 |
| ddi-mir-1186 | AAAUGAAAUUAGAGAAAGGGA | 131 | 17,50 | 97,8 | intergenic | NC_007090.3:1706364..1706468 | + | None | -56,91 | 87 |
| ddi-mir-1176 | CCAAUUUUUAUCAAGGAAAGC | 763 | 93,10 | 98,5 | intergenic | NC_007091.3:172131..172231 | + | None | -70,6 | 83 |
| ddi-mir-1177 | CCAGUUAGGGUUUAAUGGUUC | 777 | 113,00 | 98,7 | intergenic | NC_007091.3:593946..594096 | + | None | -65,48 | 133 |
| dfi-mir-1192 | UGUACCAAGACAAUCUCCAAG | 71 | 8,88 | 84,5 | genome_not_annotated | JH723762.1:365175..365282 | + | None | -69,88 | 90 |
| dfi-mir-1191 | UUUCCUCGUUACACUUGUUUG | 93138 | 10000,00 | 97,1 | genome_not_annotated | JH723798.1:37137..37323 | - | None | -95,3 | 169 |
| dfi-mir-1190 | UGAGGACAUUUGAUAAUUUUGA | 101828 | 18600,00 | 76,9 | genome_not_annotated | JH723798.1:203338..203593 | - | None | -142,84 | 238 |
| dfi-mir-1195 | UGAUGUCGCCAAUGUAAUAAC | 541470 | 52400,00 | 97,7 | genome_not_annotated | JH723896.1:3805..3933 | + | None | -69,84 | 111 |
| dfi-mir-1196 | UAGUGAAGAUGGGUAGAAGAU | 53 | 6,29 | 76,8 | genome_not_annotated | JH724042.1:4390..4506 | + | None | -75,37 | 99 |
| dla-mir-1197 | CAUUACUCAUAUAUCUCAGAC | 33 | 11,60 | 94,3 | intergenic | LODT01000020.1:101326..101465 | - | None | -93,54 | 122 |
| dla-mir-1198 | AUGUUGGUCUUCAUGUUGUUUU | 126922 | 31600,00 | 99,8 | intergenic | LODT01000020.1:565249..565350 | + | None | -52,92 | 84 |
| ppa-mir-1199 | AUUUUUUCAAGAUGUCUGUUC | 2534 | 195,00 | 83,6 | intronic_antisense | NW_008805065.1:51896..51998 | - | None | -65,05 | 85 |
| ppa-mir-1200 | UGCUAAAAUCAAAAAUAGACA | 48 | 2,35 | 100 | intergenic | NW_008805068.1:144909..145012 | + | None | -35,21 | 86 |
| ppa-mir-1201 | AGAAUGGUAAAGUGUCAGAUA | 523 | 27,90 | 99,1 | intronic_antisense | NW_008805072.1:1157625..1157739 | - | None | -54,86 | 97 |
| ppa-mir-1202 | AAUAGAAUCUGUUGUGCUUCA | 3577 | 529,00 | 99,2 | intronic_intergenic | NW_008805100.1:69067..69273 | - | None | -101,16 | 189 |
| asu-mir-1203 | UUAUUGGGUAUUAAAUAAGUA | 2252 | 423,00 | 92 | intergenic | NW_012236446.1:76..190 | - | None | -52,88 | 97 |
| asu-mir-1204-P1 | UAUCUAUUGUAGCACUUGUGA | 2258382 | 50400,00 | 99,5 | intergenic | NW_012236484.1:550..651 | - | mir-1204 | -48,86 | 84 |
| asu-mir-1204-P2 | UAUCUAUUGUAGCACUUGUGA | 2258382 | 50500,00 | 99,5 | intergenic | NW_012236496.1:211486..211587 | + | mir-1204 | -48,86 | 84 |
| asu-mir-1204-P3 | UAUCUAUUGUAGCACUUGUGA | 2258382 | 50200,00 | 99,5 | 3'antisense | NW_012236497.1:34090..34191 | + | mir-1204 | -48,86 | 84 |
| asu-mir-1205 | UGAUGAACAUUUGAGUGGCGU | 9437 | 1570,00 | 98,5 | intergenic | NW_012236508.1:360263..360396 | + | None | -70,5 | 116 |
| asu-mir-1206 | UGCAUCAUCUCGAUCGAUUCG | 23447 | 3680,00 | 99,3 | intergenic | NW_012236512.1:176300..176451 | - | None | -85,3 | 134 |
| asu-mir-1207 | UGUUUGGACAGCUUGCGUAUG | 10953 | 1810,00 | 99,9 | intergenic | NW_012236512.1:427608..427773 | - | None | -87,6 | 148 |
| asu-mir-1208 | UGCUGUGAGAUUGGUCACCGAGC | 11207 | 1900,00 | 97,8 | intergenic | NW_012236514.1:586899..587045 | + | None | -68,63 | 129 |
| asu-mir-1204-P4 | UAUCUAUUGUAGCACUUGUGA | 2258382 | 50600,00 | 99,5 | intergenic | NW_012236515.1:46736..46837 | - | mir-1204 | -48,86 | 84 |
| asu-mir-1209 | UGAACGAUUUUCACCAAAAUC | 19978 | 3140,00 | 99,9 | intergenic | NW_012236516.1:452505..452631 | - | None | -73,19 | 109 |
| asu-mir-1210 | UGAGCCGUACCAUUCUCUGUCU | 410 | 91,70 | 95,8 | intergenic | NW_012236516.1:536175..536295 | - | None | -76,03 | 103 |
| asu-mir-1211 | AAUAUGACAUUGUUAACGGGU | 3835 | 629,00 | 99,9 | intergenic | NW_012236518.1:732968..733071 | - | None | -55,45 | 86 |
| asu-mir-1212 | UCACCGAUGGCCUACUGCAUG | 1920 | 311,00 | 99,4 | exonic_5'sense | NW_012236519.1:489113..489224 | + | None | -51,1 | 94 |
| asu-mir-1213 | UGCUACCAUGUUAGACUGAUG | 19904 | 3910,00 | 96,7 | intergenic | NW_012236521.1:506588..506691 | - | None | -67,09 | 86 |
| asu-mir-1204-P5 | UAUCUAUUGUAGCACUUGUGA | 2258382 | 50100,00 | 99,5 | 3'antisense | NW_012236523.1:1239..1340 | + | mir-1204 | -48,86 | 84 |
| asu-mir-1204-P6 | UAUCUAUUGUAGCACUUGUGA | 2258382 | 50800,00 | 99,5 | 3'antisense | NW_012236523.1:1049681..1049782 | - | mir-1204 | -48,86 | 84 |
| asu-mir-1214 | ACAAAUGUUGAACAGAUAAUA | 405 | 72,50 | 99,5 | intergenic | NW_012236526.1:1087391..1087493 | + | None | -39,78 | 85 |
| asu-mir-1215 | UCUUUACAGUAGUGCCAUUCC | 181 | 36,30 | 93,3 | intergenic | NW_012236527.1:681242..681431 | - | None | -129,41 | 172 |
| asu-mir-1204-P7 | UAUCUAUUGUAGCACUUGUGA | 2258382 | 50200,00 | 99,5 | 3'antisense | NW_012236528.1:129735..129836 | + | mir-1204 | -48,86 | 84 |
| asu-mir-1204-P8 | UAUCUAUUGUAGCACUUGUGA | 2258382 | 50400,00 | 99,5 | 3'antisense | NW_012236530.1:25146..25247 | + | mir-1204 | -48,86 | 84 |
| asu-mir-1216 | AGGUGUAGCUAUGUCAUUGGC | 119836 | 21200,00 | 99,7 | intergenic | NW_012236531.1:1717023..1717124 | - | None | -65,75 | 84 |
| asu-mir-1217 | UCAGACAUAGUUGACACGUUC | 12051 | 1900,00 | 98,7 | intergenic | NW_012236532.1:237071..237177 | + | None | -55,33 | 89 |
| dfa-mir-1218 | AAGAGAUGAAUGCUGAAAAGA | 41 | 22,40 | 95,3 | exonic_antisense | NW_004457690.1:214791..214912 | + | None | -64,67 | 104 |
| dfa-mir-1219 | UUAAACUCGUUUAUCAUUAUA | 812 | 385,00 | 92,8 | exonic_antisense | NW_004457713.1:100858..101017 | + | None | -84,55 | 142 |
| dfa-mir-1220-P1 | UAAGUGAUGAUGGUAAUGAUC | 114664 | 20600,00 | 99,9 | intronic_antisense | NW_004457729.1:1425002..1425169 | - | mir-1220 | -61,47 | 150 |
| dfa-mir-1220-P2 | UAAGUGAUGAUGGUAAUGAUC | 114658 | 20800,00 | 99,9 | intergenic | NW_004457729.1:1445710..1445873 | - | mir-1220 | -66,48 | 146 |
