## Supplementary Table 3 for "Evolution of microRNAs in Amoebozoa and implications for the origin of multicellularity"

**Supplementary Table 3 | miRNA candidates in *Physarum polycephalum* and *Acanthamoebae* which passed at least one of the three sets of miRNA annotation criteria.** Each of the miRNA candidates were analyzed and listed under miRNA.ID, either as their given miRNA name for those that passed our criteria for defining miRNAs or by their cluster annotation for those that did not pass our miRNA definition. All miRNA candidates were subjected to the three sets of miRNA annotation criteria: axtell_pass, mirbase_pass, mirgenedb_pass. If a candidate did not pass all the criteria of a given set, the reason is indicated (see Extended Data Fig. 3). Any candidate that passed at least two out of three sets of criteria, and that passed manual inspection, were identified as *bona fide* miRNAs (passed/failed).

| species | miRNA.ID | axtell_pass | mirbase_pass | mirgenedb_pass | passed/failed |
|---|---|---|---|---|---|
| P. polycephalum | ppo-mir-1221 | 71% precision | TRUE | TRUE | passed |
| P. polycephalum | ppo-mir-1222 | TRUE | TRUE | TRUE | passed |
| P. polycephalum | ppol_Cluster_1701 | 71% precision | TRUE | 81% precision | failed: only one set of criteria |
| P. polycephalum | ppo-mir-1223 | TRUE | 4 miR* reads | TRUE | passed |
| P. polycephalum | ppol_Cluster_2782 | one_library | 33% miR* precision | TRUE | failed: only one set of criteria |
| P. polycephalum | ppol_Cluster_4525 | TRUE | 4 miR* reads | 0 unpaired bases | failed: only one set of criteria |
| P. polycephalum | ppo-mir-1224 | TRUE | 9 miR* reads | TRUE | passed |
| P. polycephalum | ppol_Cluster_6245 | TRUE | 17% miR* precision | TRUE | failed: manual inspection |
| P. polycephalum | ppol_Cluster_8717 | TRUE | 1% miR* precision | TRUE | failed: manual inspection |
| P. polycephalum | ppol_Cluster_8730 | TRUE | 29% miR* precision | TRUE | failed: manual inspection |
| P. polycephalum | ppo-mir-1225 | TRUE | TRUE | TRUE | passed |
| P. polycephalum | ppo-mir-1226 | TRUE | 48% miR* precision | TRUE | passed |
| P. polycephalum | ppol_Cluster_13697 | 2 bulges | TRUE | 52% precision | failed: only one set of criteria |
| P. polycephalum | ppo-mir-1227 | TRUE | TRUE | TRUE | passed |
| A. castellanii | aca-mir-1228 | TRUE | TRUE | TRUE | passed |
| A. lenticulata | ale-mir-1228-P1 | 50% precision | TRUE | TRUE | passed |
| A. lenticulata | ale-mir-1229 | TRUE | 3 miR* reads | TRUE | passed |
| A. lenticulata | alen_Cluster_1703 | TRUE | 4 miR* reads | 88% precision | failed: only one set of criteria |
| A. lenticulata | ale-mir-1228-P2 | 50% precision | TRUE | TRUE | passed |
| A. lenticulata | ale-mir-1230-P1 | TRUE | TRUE | TRUE | passed |
| A. lenticulata | ale-mir-1230-P2 | TRUE | TRUE | TRUE | passed |
| A. lenticulata | alen_Cluster_5165 | 1 bulges | TRUE | 67% precision | failed: only one set of criteria |
