## Supplementary Table 4 for "Evolution of microRNAs in Amoebozoa and implications for the origin of multicellularity"

**Supplementary table 4 | Characteristics of the *Physarum polycephalum* and *Acanthamoebae* miRNAs.** For each miRNA, the mature miRNA sequence is displayed (the most abundant strand in the duplex). miRNA_counts is the sum of miRNA-5p and miRNA-3p reads, and miRNA_RPM are the normalized reads on the miRNA-hairpin per million mapped total. The precision is the percentage of reads from the miRNA-hairpin that map with their 5' end at the exact 5’ end of either the miRNA-5p or miRNA-3p. The genomic_context, genomic_location and strand show where the miRNA is located relative to the annotated ORFs, its genomic location, and whether it is from the Watson strand (+) or the Crick strand (-), respectively. For those miRNAs where we could detect homology, the miRNA_family is indicated, and for identified paralogs, a suffix is included in the miRNA.ID. Folding energy is the computed Gibbs free energy in kcal/mol from folding the miRNA-hairpin in silico at 22 °C. The pre-miRNA size (nt) is the length (in nt) of the pre-miRNA, i.e. from the miRNA-5p 5’ end to the miRNA-3p 3’ end.

| miRNA.ID | miRNA_sequence | miRNA_counts | miRNA_RPM | precision | genomic_context | genomic_location | strand | miRNA_family | folding energy (kcal/mol) | pre-miRNA size (nt) |
|---|---|---|---|---|---|---|---|---|---|---|
| ppo-mir-1221 | UCAUAAACUUUGAAAUCCUCC | 4422 | 797,00 | 99,5 | genome_not_annotated | KL558924.1:33157..33298 | - | None | -81,92 | 124 |
| ppo-mir-1222 | UCAGGUUGAGUUGUCCACAGC | 415 | 74,30 | 98,1 | genome_not_annotated | KL558932.1:231142..231356 | + | None | -128,63 | 197 |
| ppo-mir-1223 | UGAAAAAUGCUGGCUGUCAG | 121 | 21,20 | 100 | genome_not_annotated | KL559005.1:39250..39327 | + | None | -32,06 | 60 |
| ppo-mir-1224 | UUUAGAACAUGAACUCGGACU | 134 | 25,00 | 98,5 | genome_not_annotated | KL559224.1:68684..68797 | - | None | -68,01 | 96 |
| ppo-mir-1225 | AAUUAUGCCAGUGAACCUCUG | 270 | 49,20 | 94,4 | genome_not_annotated | KL560176.1:27473..27625 | + | None | -98,02 | 135 |
| ppo-mir-1226 | UCUGAAGUAAGGUACCCUUUG | 3633 | 676,00 | 98 | genome_not_annotated | KL561177.1:1944..2236 | + | None | -184,64 | 275 |
| ppo-mir-1227 | UGCACAUCAUGGACACUCACC | 398 | 67,10 | 97,3 | genome_not_annotated | KL563403.1:1258..1407 | - | None | -64,91 | 132 |
| aca-mir-1228 | UUUAGUGCCUAUGUCCCCUUC | 147 | 8,23 | 94,8 | intergenic | JAJGAP010000002.1:1842520..1842601 | + | mir-1228 | -56,3 | 64 |
| ale-mir-1228-P1 | UUUAGUGCCUAUGUCCUCUUCA | 463 | 14,90 | 99,4 | genome_not_annotated | NAVB01000078.1:67963..68054 | + | mir-1228 | -58,42 | 74 |
| ale-mir-1229 | AGCGGGCUGGAGAUGAAGCGGA | 73 | 5,98 | 96,1 | genome_not_annotated | NAVB01000357.1:7568..7710 | + | None | -96,8 | 125 |
| ale-mir-1228-P2 | UUUAGUGCCUAUGUCCUCUUCA | 463 | 16,60 | 99,4 | genome_not_annotated | NAVB01000824.1:23909..24000 | + | mir-1228 | -58,46 | 74 |
| ale-mir-1230-P1 | CACGCUCUCGGGAGACGAAGC | 699 | 26,50 | 93,4 | genome_not_annotated | NAVB01000891.1:9327..9409 | - | mir-1230 | -58,43 | 65 |
| ale-mir-1230-P2 | CACGCUCUCGGGAGACGAAGC | 699 | 25,80 | 93,3 | genome_not_annotated | NAVB01003488.1:1461..1547 | - | mir-1230 | -65,24 | 69 |
