## Supplementary Table 5 for "Evolution of microRNAs in Amoebozoa and implications for the origin of multicellularity"

**Supplementary Table 5 | Comparison of the *D. discoideum* and *D. firmibasis* genomes used in this study.** Undetermined bases show the percentage of 'N' bases in the genome. The de-novo sequenced *D. firmibasis* genome is more contiguous as can be seen from the number of contigs and N50 (the length of the contig at the halfway-point of the genome). It is also more complete as can be seen from the BUSCO analysis.

| Species | GenBank assembly accession | GC (%) | Undetermined bases (%) | No chromosomes / contigs | N50 (Mbps) | BUSCO Ids: Single / Duplicated / Fragmented / Missing (%) |
|---|---|---|---|---|---|---|
| Dictyostelium discoideum | GCA_000004695.1 | 22,4 | 0,07 | 8 | 5,13 | 90.2 / 3.5 / 2.0 / 4.3 |
| Dictyostelium firmibasis | GCA_000277485.1 | 21,4 | 13,46 | 997 | 0,18 | 81.2 / 1.6 / 5.1 / 12.1 |
| Dictyostelium firmibasis | This study | 24,4 | 0 | 8 | 4,38 | 92.9 / 0.4 / 2.7 / 4.0 |
