## Supplementary Table 6 for "Evolution of microRNAs in Amoebozoa and implications for the origin of multicellularity"

**Supplementary Table 6 | Characteristics of the *D. firmibasis* miRNAs identified using the previously sequenced genome and the de-novo sequenced genome.** The previously sequenced genome can be accessed with genbank assembly number GCA_000277485.1. For each miRNA, the mature miRNA sequence is displayed (the most abundant strand in the duplex). miRNA_counts is the sum of miRNA-5p and miRNA-3p reads, and miRNA_RPM are the normalized reads on the miRNA-hairpin per million mapped total. The precision is the percentage of reads from the miRNA-hairpin that map with their 5' end at the exact 5' end of either the miRNA-5p or miRNA-3p. The genomic_context, genomic_location and strand show where the miRNA is located relative to the annotated ORFs, its genomic location, and whether it is from the Watson strand (+) or the Crick strand (-), respectively. For those miRNAs where we could detect homology, the miRNA_family is indicated, and for identified paralogs, a suffix is included in the miRNA.ID. Folding energy is the computed Gibbs free energy in kcal/mol from folding the miRNA-hairpin in silico at 22 °C. The pre-miRNA size (nt) is the length (in nt) of the pre-miRNA, i.e. from the miRNA-5p 5' end to the miRNA-3p 3' end.

| Genome | miRNA.ID | miRNA_sequence | miRNA_counts | miRNA_RPM | precision | genomic_context | genomic_location | miRNA_family | strand | folding energy (kcal/mol) | pre-miRNA size (nt) |
|---|---|---|---|---|---|---|---|---|---|---|---|
| GCA_000277485.1 | dfi-mir-1190 | UGAGGACAUUUGAUAAUUUUGA | 101828 | 8,88 | 76,9 | genome_not_annotated | JH723798.1:203338..203593 | None | + | -142,84 | 238 |
| GCA_000277485.1 | dfi-mir-1191 | UUUCCUCGUUACACUUGUUUG | 93138 | 10000,00 | 97,1 | genome_not_annotated | JH723798.1:37137..37323 | None | - | -95,3 | 169 |
| GCA_000277485.1 | dfi-mir-1192 | UGUACCAAGACAAUCUCCAAG | 71 | 18600,00 | 84,5 | genome_not_annotated | JH723762.1:365175..365282 | None | - | -69,88 | 90 |
| GCA_000277485.1 | dfi-mir-1195 | UGAUGUCGCCAAUGUAAUAAC | 541470 | 52400,00 | 97,7 | genome_not_annotated | JH723896.1:3805..3933 | None | + | -69,84 | 111 |
| GCA_000277485.1 | dfi-mir-1196 | UAGUGAAGAUGGGUAGAAGAU | 53 | 6,29 | 76,8 | genome_not_annotated | JH724042.1:4390..4506 | None | + | -75,37 | 99 |
| This study | dfi-mir-1189 | CAUGGAAGAUUAAUGACCGCC | 308939 | 30700,00 | 76,3 | genome_not_annotated | D_firmibasis_chromosome_1:5202378..5202481 | None | - | -108,21 | 86 |
| This study | dfi-mir-1190 | UGAGGACAUUUGAUAAUUUUGA | 101828 | 16400,00 | 76,9 | genome_not_annotated | D_firmibasis_chromosome_1:6559035..6559290 | None | + | -142,84 | 238 |
| This study | dfi-mir-1191 | UUUCCUCGUUACACUUGUUUG | 93138 | 8790,00 | 97,1 | genome_not_annotated | D_firmibasis_chromosome_1:6719430..6719616 | None | + | -95,3 | 169 |
| This study | dfi-mir-1192 | UGUACCAAGACAAUCUCCAAG | 71 | 7,79 | 84,5 | genome_not_annotated | D_firmibasis_chromosome_1:7110680..7110787 | None | - | -69,88 | 90 |
| This study | dfi-mir-1193 | UUACUCCCAUUAUUAAGGAUG | 63449 | 4940,00 | 99,6 | genome_not_annotated | D_firmibasis_chromosome_2:1686826..1686932 | None | + | -42,09 | 89 |
| This study | dfi-mir-1177 | CCAGUUAGGGUUUAAUGGUUC | 1074 | 90,00 | 96,2 | genome_not_annotated | D_firmibasis_chromosome_3:577596..577756 | mir-1177 | - | -69,66 | 143 |
| This study | dfi-mir-1194 | UUUCGCAUCAGAGCUCCGAGC | 412169 | 35300,00 | 98,8 | genome_not_annotated | D_firmibasis_chromosome_3:3078863..3078964 | None | - | -101,3 | 84 |
| This study | dfi-mir-1195 | UGAUGUCGCCAAUGUAAUAAC | 541470 | 46100,00 | 97,7 | genome_not_annotated | D_firmibasis_chromosome_4:3840639..3840767 | None | - | -69,84 | 111 |
| This study | dfi-mir-1196 | UAGUGAAGAUGGGUAGAAGAU | 53 | 5,52 | 76,8 | genome_not_annotated | D_firmibasis_chromosome_6:2524364..2524480 | None | - | -75,37 | 99 |
